# Computation of Antigenicity Predicts SARS-CoV-2 Vaccine Breakthrough Variants

**DOI:** 10.1101/2022.01.19.477009

**Authors:** Ye-fan Hu, Jing-chu Hu, Hua-rui Gong, Antoine Danchin, Ren Sun, Hin Chu, Ivan Fan-Ngai Hung, Kwok Yung Yuen, Kelvin Kai-Wang To, Bao-zhong Zhang, Thomas Yau, Jian-Dong Huang

**Affiliations:** School of Biomedical Sciences, Li Ka Shing Faculty of Medicine, University of Hong Kong, 3/F, Laboratory Block, 21 Sassoon Road, Hong Kong, China; CAS Key Laboratory of Quantitative Engineering Biology, Shenzhen Institute of Synthetic Biology, Shenzhen Institutes of Advanced Technology, Chinese Academy of Sciences, Shenzhen 518055, China; Department of Medicine, Li Ka Shing Faculty of Medicine, University of Hong Kong, 4/F Professional Block, Queen Mary Hospital, 102 Pokfulam Road, Hong Kong, China; Department of Microbiology, Li Ka Shing Faculty of Medicine, University of Hong Kong, 19/F T Block, Queen Mary Hospital, 102 Pokfulam Road, Hong Kong, China; Kodikos Labs / Stellate Therapeutics, Institut Cochin, 24 rue du Faubourg Saint-Jacques, 75014 Paris, France

## Abstract

It has been reported that multiple SARS-CoV-2 variants of concerns (VOCs) including B.1.1.7 (Alpha), B.1.351 (Beta), P.1 (Gamma), and B.1.617.2 (Delta) can reduce neutralisation by antibodies, resulting in vaccine breakthrough infections. Virus-antiserum neutralisation assays are typically performed to monitor potential vaccine breakthrough strains. However, such experimental-based methods are slow and cannot instantly validate whether newly emerging variants can break through current vaccines or therapeutic antibodies. To address this, we sought to establish a computational model to predict the antigenicity of SARS-CoV-2 variants by sequence alone and in real time. In this study, we firstly identified the relationship between the antigenic difference transformed from the amino acid sequence and the antigenic distance from the neutralisation titres. Based on this correlation, we obtained a computational model for the receptor binding domain (RBD) of the spike protein to predict the fold decrease in virus-antiserum neutralisation titres with high accuracy (~0.79). Our predicted results were comparable with experimental neutralisation titres of variants, including B.1.1.7 (Alpha), B.1.351 (Beta), B.1.617.2 (Delta), B.1.429 (Epsilon), P.1 (Gamma), B.1.526 (Iota), B.1.617.1 (Kappa), and C.37 (Lambda), as well as SARS-CoV. Here, we firstly predicted the fold of decrease of B.1.1.529 (Omicron) as 17.4-fold less susceptible to neutralisation. We visualised all 1521 SARS-CoV-2 lineages to indicate variants including B.1.621 (Mu), B.1.630, B.1.633, B.1.649, and C.1.2, which can induce vaccine breakthrough infections in addition to reported VOCs B.1.351 (Beta), P.1 (Gamma), B.1.617.2 (Delta), and B.1.1.529 (Omicron). Our study offers a quick approach to predict the antigenicity of SARS-CoV-2 variants as soon as they emerge. Furthermore, this approach can facilitate future vaccine updates to cover all major variants. An online version can be accessed at http://jdlab.online.

---

Up to January 2022, there have been several SARS-CoV-2 variants including B.1.1.7 (Alpha) ^1–5^, B.1.351 (Beta) ^2,3,6,7^, P.1 (Gamma) ^1,2,8^, and B.1.617.2 (Delta) ^9,10^ that are experimentally tested to lead vaccine breakthrough infections, thus they have been designated as variants of concerns (VOCs) by the world health organization (WHO). There is a concern that other untested emerging variants may lead to vaccine breakthrough infections ^11–16^. The most recent case is the validation of B. 1.1.529 (Omicron). The current virological and epidemiological techniques took several weeks to validate whether the variant is capable of reducing the efficacy of current vaccines ^17,18^, or therapeutic antibodies ^18,19^, even though their viral sequences have been shared in real time via the Global Initiative for Sharing All Influenza Data (GISAID) ^20^. The speed of validation of vaccine breakthrough variants can hardly catch up with the fast-emerging rate of new variants. Thus, it is crucial to develop new approaches for identifying the next potential vaccine breakthrough variant as soon as it is reported.

Here, we established a computational approach for predicting the antigenicity of SARS-CoV-2 variants from viral sequences alone, with the aim to accelerate the identification of potential vaccine breakthrough variants. Our approach is founded on the concept of antigenic mapping, also named antigenic cartography. This method has been used to monitor vaccine breakthrough variants of influenza virus using haemagglutination inhibition (HI) assay data^21,22^, dengue virus and SARS-CoV-2 circulating strains ^24^ using pairwise antisera data. In antigenic mapping, the antigenic distance is calculated from the fold change of the neutralisation titre between the reference virus and its variant, to measure the change of antigenicity between two variants. A computational approach for predicting antigenic distances to indicate vaccine breakthrough variants could theoretically provide much more rapid results once the variant sequence is reported. Past studies proposed a linear relationship between amino acid changes in antigenic sites and neutralisation fold decrease ^25–29^, Computational prediction approaches based on such a relationship could also provide reliable estimates of neutralisation titres for existing antiserum against the vaccine breakthrough variants with similar accuracy to experiment-based approaches used in previous studies ^25–29^. However, these predictions were optimised for influenza virus instead of SARS-CoV-2. For example, the neutralisation titre decrease of any SARS-CoV-2 variant should be less than that of SARS-CoV comparing to the ancestral strain of SARS-CoV-2, because the cross protection between the SARS-CoV-2 variant and the ancestral strain is stronger than that between SARS-CoV and SARS-CoV-2. Thus, it is difficult to use a linear relationship to predict the decrease in neutralisation titre which saturates with the increase in the mutation numbers of variants. A SARS-CoV-2 optimised model for predicting antigenicity is urgently needed.

In this study, we established a computational sequence-based method to predict the antigenicity of SARS-CoV-2 variants to reveal potential vaccine breakthrough variants. This method can also predict the neutralisation titre of VOCs in comparison to the ancestral strain of SARS-CoV-2. Our predicted results were comparable with experimental neutralisation titres of VOCs, including B.1.1.7 (Alpha), B.1.351 (Beta), B.1.617.2 (Delta), B.1.429 (Epsilon), P.1 (Gamma), B.1.526 (Iota), B.1.617.1 (Kappa), and C.37 (Lambda), as well as SARS-CoV. Here, we predicted that B.1.1.529 (Omicron) is 17.4-fold less susceptible to neutralisation, which is consistent with reported decrease folds ranging from 10 to 40 ^17,18^.

## A computational model for predicting antigenicity of SARS-CoV-2 variants

To predict the antigenicity of SARS-CoV-2 variants, we firstly integrated the reported conformational or linear epitopes **(Fig. S1 &Table S1)** on the SARS-CoV-2 Spike protein **(Fig. 1a)** with the reported experimental virus-antiserum neutralisation titres against SARS-CoV-2 variants including B.1.1.7 ^1–5^, B.1.351 ^2,3,6,7^, and P.1 ^1,2,8^ **(Table S2a)**. Considering the distinct assays used in the different studies, we standardised the neutralisation titres of each variant to the titre of the ancestral strain of SARS-CoV-2 (lineage A) using the same assay in each study on a log 2 scale, and thus we got observed antigenic distance (*H_ab_*) from neutralisation titres **(Fig. 1b).** For the antigenic difference (*D_ab_*), we used Poisson distance to represent the difference between two amino acid sequences **(Fig. 1b)**. By comparing the observed antigenic distance with the antigenic difference, we found a relationship between observed antigenic distance and the antigenic difference: *H_ab_*=*T_max_*·*D_ab_*/(*D*_50_+*D_ab_*), where *T_max_* is the maximal fold of decrease and *D_50_* is the antigenic difference which may lead to neutralisation decrease at the 50% level of the maximal decrease (the fold change between SARS-CoV-2 and SARS-CoV). This relationship described that the decrease of neutralisation titre increases with the accumulation of amino acid changes, and then reaches at the maximal decrease **(Figs. 1c-d).** Based on this correlation, we obtained a computational model using the receptor binding domain (RBD) of the spike protein to predict the fold decrease in virus-antiserum neutralisation titres with higher accuracy **(~0.79, the calculation of accuracy in Methods)** compared with other fragments of spike (entire spike, N terminal domain plus RBD, or S1, **Fig. 1d**). With repeated 5-fold or 10-fold cross validation **(Fig. 1d)**, we found that prediction using RBD is relatively robust in terms of root-mean-square error (RMSE), mean absolute error (MAE), coefficient of determination (R^2^) and accuracy.

**Fig. 1.**
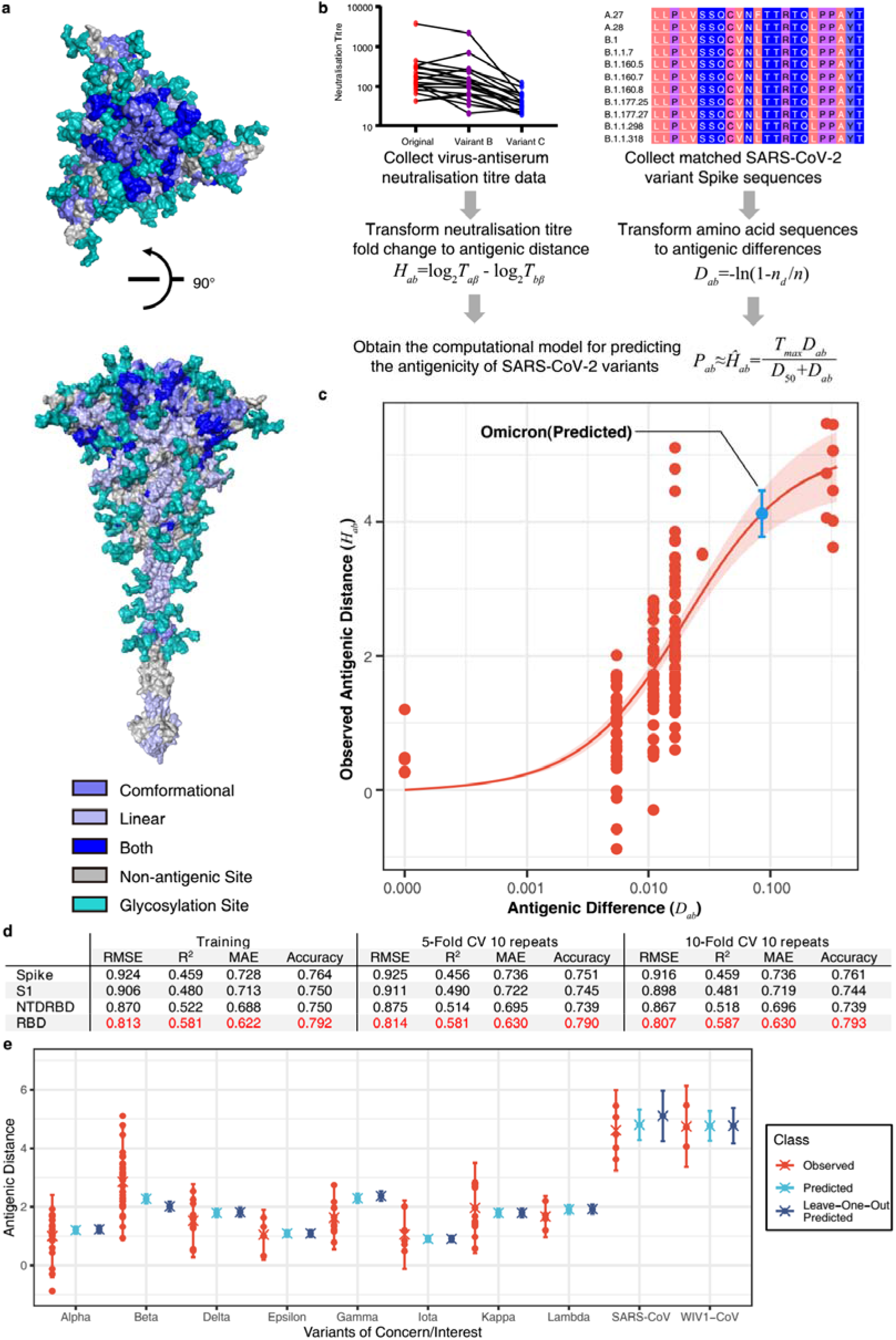
Sequence-based prediction of antigenic distance. **(a)** The top view and the side view of antigenic sites on the full-length Spike protein ^34^. The conformational epitopes are coloured in slate and linear epitopes in light blue. Some antigenic positions in both conformational epitopes and linear epitopes are coloured in blue. All glycosylation sites are in teal. **(b)** A flowchart of the process to establish the sequence-based computational model of SARS-CoV-2 antigenicity. The antigenic distance of variant *a* to reference virus *b* from neutralisation titre was defined as *H_ab_*=log_2_*T_aβ_* - log_2_ *T_bβ_*, where *β*, *T_aβ_*, and *T_bβ_* denote antiserum (referencing virus *b*), the titre of antiserum *β* against virus *b*, and the titre of antiserum *β* against virus *a* ^26^. The antigenic distance of variant *a* to reference virus *b* from amino acid sequences was defined as *D_ab_*=-ln(1-*n_d_/n*), where *n_d_* is the number of amino acid substitutions between variant *a* and reference virus *b*, *n* is the number of antigenic sites. Then, we proposed a relationship between observed antigenic distance and the antigenic difference: *H_ab_=T_max_*·*D_ab_*/(*D*_50_+*D_ab_*), where *T_max_* is the maximal fold of decrease and *D*_50_ is the antigenic difference which may lead to neutralisation decrease at the 50% level of the maximal decrease. **(c)** The relationship between the antigenic difference and the observed antigenic distance. The predicted antigenic distance of B.1.1.529 (Omicron) is marked in cyan. **(d)** The performance of the model in different fragments of the spike protein in terms of root-mean-square error (RMSE), mean absolute error (MAE), coefficient of determination (R-squared R^2^), and accuracy. **(e)** Predicted versus observed antigenic distances of variants of concern. Here, The observed antigenic distances as fold decreases in the neutralisation titres of variants of concern versus the original strain on a log 2 scale. Each point shows the mean of antigenic distances in each assay. Predicted antigenic distances are based on the prediction in (c). Leave-one-out predicted antigenic distances are predicted based on the datasets without the variant that we aim to compare.

To further validate our model, we predicted the fold decreases in neutralisation titres (comparing to the ancestral of SARS-CoV-2) of multiple variants including B.1.1.7 (Alpha), B.1.351 (Beta), B.1.617.2 (Delta), B.1.429 (Epsilon), P.1 (Gamma), B.1.526 (Iota), B.1.617.1 (Kappa), and C.37 (Lambda), as well as SARS-CoV and WIV1-CoV using datasets without the variant that we aimed to validate. Previous studies have reported that VOCs can elicit vaccine breakthrough infections, which correlated with fold decreases in the neutralisation titres from experimental assays was disclosed **(Table S2)**. Our predicted results were highly consistent with the neutralisation assay results **(Fig. 1e)**. We also predicted the fold of decrease in neutralisation titre of the most recent VOC, B. 1.1.529 (Omicron). Considering 15 mutations in the spike of B.1.1.529 (Omicron), the variant is estimated to have a 17.44-fold (95% confidence interval: 13.7, 22.2) decrease in neutralisation titre (shown as a blue point in **Fig. 1c**)., The predicted result is consistent with reported decrease folds ranging from 10 to 40 ^17,18^. This result alarmed the risk of vaccine breakthrough or re-infection of B.1.1.529 (Omicron) due to the dramatic decrease in neutralization.

## The prediction of potential vaccine breakthrough strains

To predict the next potential SARS-CoV-2 vaccine breakthrough variants, we visualised the antigenicity of all available SARS-CoV-2 variants as an indicator of their vaccine breakthrough potential. We firstly selected all 1521 lineage variants using PANGO ^30^ updated on December 6, 2021 (**Table S3**) to predict their antigenicity. Then we calculated the pairwise distances of different variants. For visualising these results, we captured two principal components from the high-dimensional data of antigenic distance ^25^. We used all spike amino acid sequences to plot the ‘genetic map’ of SARS-CoV-2 to represent the genetic difference among different variants **(Fig. 2a-b)**. We then plotted the ‘antigenic map’ using the predicted antigenic distances **(Fig. 2c-d,** online versions available at http://jdlab.online).

**Fig. 2.**
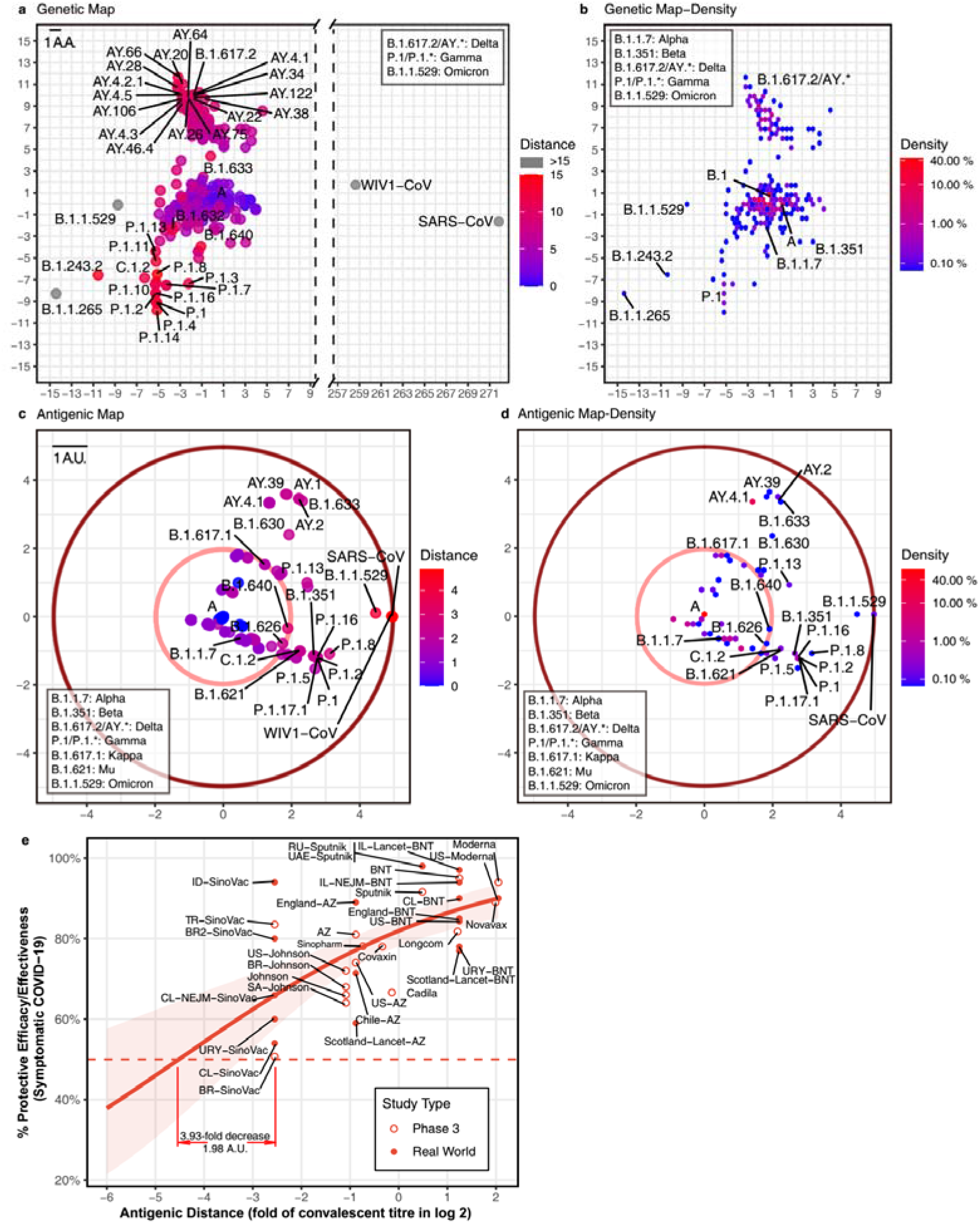
Genetic and antigenic mapping of SARS-CoV-2 variants. **(a)** Genetic map of SARS-CoV-2 variant strains shows amino acid mutation numbers of spike proteins, and **(b)** the density of genetic map shows distribution of variants. The vertical and horizontal axes represent the measured relative genetic distances (1 amino acid/1 A.A. = 1 amino acid difference). **(c)** Antigenic map of SARS-CoV-2 variant strains shows the antigenic distance between variants, and **(d)** the density of antigenic map shows distribution of variants. Variants outside the pink circle are vaccine breakthrough candidates. The red circle suggested the border of antigenic map. The antigenic distance is based on RBD amino acid sequences. The vertical and horizontal axes represent the measured relative antigenic distances (1 arbitrary unit/1 A.U. = 1-fold decrease in the neutralisation titre on a log 2 scale). Colours show the antigenic distance to the SARS-CoV-2 original strain (lineage A). **(e)** Relationship between antigenic distance (mean of neutralisation titres in vaccinees divided by corresponding mean of titres in convalescent patients in log 2) and protection from SARS-CoV-2 infection. The reported mean neutralization level from phase 1 or 2 studies **(Table S4)** and the protective efficacy or effectiveness from phase 3 trials or real-world studies **(Table S5)** for different vaccines. The red line indicates the logistic model, and the red shading indicates the 95% confidence interval of the model. Here, we mark the basis of setting up the cut-off of 3.93-fold decrease (1.98 A.U.).

Based on the relationship between neutralisation titre fold change and protective efficacy ^31^, it was convenient to set up some ‘cut-offs’ in the current vaccine coverage. We included phase 3 and real-world results of vaccine efficacy or effectiveness, as well as neutralisation titre data from phase 1 and 2 studies **(Table S4-5)**. Thus, we got the relationship between neutralisation titre and protective efficacy against a symptomatic COVID-19 **(Fig. 2e)**. A 3.93-fold decrease in neutralisation titres induced by VOCs that can dampened the efficacy of some vaccines to lower than 50%. In this way, one cut-off of 1.98 arbitrary units (A.U.) represented a 3.93-fold decrease in the neutralisation titre (shown as a pink circle in **Fig. 2c-d**). All variants outside this cut-off have the potential to be vaccine breakthrough variants. By comparing the “genetic map” and antigenic map, we can set up the border of antigenic map. Although there are >200 mutations in the SARS-CoV and WIV1-CoV spike (**Fig 2a**), the antigenic distance is around 4.9 A.U. which mean ~ 30-fold decrease in the neutralisation titre (shown as a dark red circle in **Fig. 2c-d**).

To reveal the distribution of variant, we plotted the density of variants on the ‘genetic map’ and antigenic map due to overlapping dots. In the genetic map, hotspots are located at lineage A (>10%) and B.1 (>40%) mainly, as well as AY.* and P.1 (**Fig. 2b**). While in the antigenic map, hotspots are placed at lineage A (>40%) mainly, together with AY.* (**Fig. 2d**). Although most variants were shown to be close to the ancestral strain (**Figs. 2b&d**), multiple variants were found to decrease neutralisation titres significantly (**Fig. 2c**). In addition to reported VOCs including B.1.351 (Beta, containing sub-lineages like B.1.351.2 and B.1.351.5) ^2,3,6,7^, P.1 (Gamma, containing sub-lineages like P.1.11 and P.1.3) ^1,2,8^, B.1.617.2 (Delta, containing sub-lineages AY.*) ^9^, and B.1.621 (Mu, containing sub-lineage B.1.621.1), B.1.1.529 (Omicron) showed over 3.93-fold decrease in the neutralisation titre. Other variants B.1.630, B.1.633, B.1.649, and C.1.2 also have the potential to be vaccine breakthrough variants with more than 3.93-fold decrease **(Fig. 2c)**. Besides the pandemic of B.1.617.2 (Delta) ^9^ and the outbreak of B.1.1.529 (Omicron), multiple variants should be investigated immediately as they have the potential to become tomorrow’s VOCs.

## Discussion

Predicting neutralisation responses against all SARS-CoV-2 variants based on sequences alone is vital for selecting the next vaccine seeds for the development of effective COVID-19 vaccines. We established a computational approach to predict neutralisation titres and validated these predictions using experimental data. Our computational approach could potentially provide the first hints of whether a newly identified variant can break through vaccines just by its sequence information, which would greatly shorten the time for the crucial early warning of emerging vaccine breakthrough strains.

In the prediction of the antigenicity of SARS-CoV-2 variants, we proposed that the limit of neutralisation titre decrease is set by SARS-CoV **(Fig. 1)**. In recent studies, SARS-CoV is ~ 36-fold less susceptible to neutralisation comparing to the ancestral strain of SARS-CoV-2. Based on this result, a non-linear curve was established to describe the relationship between the observed antigenic distance and the antigenic difference. We further performed calculation using different fragments of the Spike protein **(Fig. 1d)**. Among the Spike protein and the RBD, NTD-RBD, and S1 fragments, we found the prediction using amino acid sequences of RBD was able to estimate the neutralisation titre more accurately than the others (**Figs. 1d**). Thus, we used the RBD-based computations to determine the neutralisation titres.

A major concern of our computation of the neutralisation titre is that the data is based on diverse neutralisation assays of serum samples from both patients and vaccinees against both live virus and pseudovirus **(Table S2).** Although the results were consistent qualitatively, the variation of fold change is too large to be ignored **(Fig. 1e)**. Considering the variation in the real world, we set up values 2-fold or less than the experimental values as the criteria based on previous studies ^28^. It is better to establish a convenient and standardised neutralisation pipeline in the future, like the haemagglutination inhibition (HI) assay for influenza virus. Such a pipeline can allow the precise estimation of neutralisation titres. Together with estimating the association of neutralisation with protection, it will help to develop next generation vaccines.

It is crucial to update vaccines to cover all vaccine breakthrough strains that have significant amino acid and glycosylation changes to prevent further infectious outbreaks. However, not all predicted SARS-CoV-2 vaccine breakthrough variants will have the chance to cause an outbreak due to their changed viral fitness ^32^ or by pure luck. Based on previous studies of influenza viruses, it is possible for variants to have alterations that change the antigenicity, but fail to cause outbreaks in the wider population ^33^. Considering immune escape elicited by variants, updating current vaccine seeds with new variants should extend the vaccine coverage. As SARS-CoV-2 showed different variant directions in the antigenic map **(Fig. 2)**, the use of multiple virus seeds based on the different directions might be appropriate to cover all major variants in the long term. Our method could help in the selection of SARS-CoV-2 variants for updating vaccines.

## Methods

### Antigenic footprint

We collected 149 confirmed conformational epitopes with protein structures released in the Protein Data Bank (PDB) (https://www.rcsb.org/) or annotated epitope footprints and 76 linear epitopes published in the literature **(Table S1)**. We plotted the footprint of all Spike protein epitopes from the aforementioned 225 epitopes using R-3.6.6.

### Antigenic distances from neutralisation data

We calculated antigenic distances from the neutralisation data based on previous publications^26^. For virus variant *a,* reference virus *b*, and antiserum *β* (referencing virus *b*), we defined the antigenic distance of variant *a* to reference virus *b* in terms of the standardised log titre as *H_ab_=log_2_T_aβ_* - log_2_*T_bβ_*, where *T*_bβ_ is the titre of antiserum *β* against virus *b*, and *T_aβ_* is the titre of antiserum *β* against virus *a* ^26^. Merged data with reference virus lineage A (the ancestral strain of SARS-CoV-2) were collected from several publications **(Table S2)**.

### Genetic and antigenic difference calculation

We selected 1521 SARS-CoV-2 lineages using PANGO (v.3.1.15) updated on December 6, 2021 (https://cov-lineages.org/). Spike protein amino acid sequences of these lineages were obtained from GISAID, using the earliest collected for each lineage **(Table S3)**. All sequences with neutralisation titres were also included **(Table S3)**. For genetic distances, we used Molecular Evolutionary Genetics Analysis (MEGA) X to calculate the pairwise distances among Spike protein amino acid sequences in the SARS-CoV-2 variants using a Poisson model. For antigenic distance, we used an information theory-based approach *p-all-epitope* ^27,28^ to measure the pairwise distances among amino acid sequences of the antigenic footprint (‘antigenic positions’). The distance is based on the number of different amino acids *n_d_* between two *n*-mer viral sequences of variants *a* and *b*. Under the assumption that the number of amino acid substitutions per site follows a Poisson distribution, we can then calculate the distance between *a* and *b* as *D_ab_*=-ln(1-*n_d_/n*).

### Modelling and performance measurement

A model considering the maximal neutralisation tire decrease was applied to examine the antigenic distance from the neutralisation data *H_ab_* and our computed results *D_ab_* as *H_ab_*=*T_max_*·*D_ab_*/(*D*_50_+*D_ab_*), where *T_max_* is the maximal decrease and *D*_50_ is the antigenic difference which may lead to neutralisation decrease at the 50% level of the maximal decrease. The predicted neutralisation titre is then given as 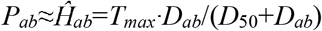. Root-mean-square error (RMSE), mean absolute error (MAE), and coefficient of determination (R-squared R^2^) were used to measure the performance of the linear correlation.

Reproducibility was determined by pairwise sequences and neutralisation titres. Neutralisation titre data were converted into variables by calculating the relative difference in the neutralisation titres between reference virus and variant against the antiserum. Accuracy was the percentage of correctly predicted neutralisation titres using amino acid sequences. Based on previous studies ^28^, computational values 2-fold or less than the experimental values were considered to be similar (correct) and those more than 2-fold lower were considered dissimilar (error). Here, 10-time repeated 5-fold and 10-fold cross validation were applied in terms of root-mean-square error (RMSE), mean absolute error (MAE), coefficient of determination (R-squared R^2^), and accuracy.

### Genetic and antigenic maps

After calculating genetic and antigenic distances, we used classical multidimensional scaling (CMDS) to display the data as a plot using R-3.6.6. We set up SARS-CoV-2 lineage A as the origin and scaled the data in two and three dimensions. We then acquired the genetic and antigenic maps of SARS-CoV-2 lineages. An online version can be obtained at http://jdlab.online.

### Logistic model

Following past studies^31^, we used a logistic model in R-3.6.6 to describe the relationship between antigenic distance (neutralization level) and protective efficacy/effectiveness: *E*=1/(1+ exp(-*k*(*H-H_50_*))). *E* is the protective efficacy/effectiveness at a specific neutralization level *H. H* is the mean of neutralisation titres in vaccinees divided by corresponding mean of titres in convalescent patients, which is the antigenic distance to convalescent patients in log 2. *H_50_* is the antigenic distance at which an individual will have a 50% protective efficacy/effectiveness.

## Acknowledgements

We acknowledge the authors and originating and submitting laboratories of the sequences from GISAID’s EpiCoV Database on which this research is based (GISAID acknowledgments are in Table S3).

## Funding

The work was supported by grants from the Health and Medical Research Fund, the Food and Health Bureau, The Government of the Hong Kong Special Administrative Region (COVID190117, COVID1903010) and Guangdong Science and Technology Department (2020B1212030004) to J.H. J.H. thanks the L & T Charitable Foundation and the Program for Guangdong Introducing Innovative and Entrepreneurial Teams (2019BT02Y198) for their support.

## Author contributions

Y.F.H., B.Z.Z., T.Y., H.C., K.K.W.T., and J.D.H. designed the study. Y.F.H. and J.C.H. analysed the sequences from GISAID and performed the computations and built the online tool. I.F.N.H. and K.K.W.T. recruited the patients and volunteers and collected samples from patients and volunteers. performed the neutralisation assay. Y.F.H., A.D., R.S., K.Y.Y., K.T., H.R.G, and J.D.H. analysed the results. Y.F.H., A.D., T.Y., and J.D.H. wrote the initial draft, and all authors edited the final version.

## Competing interests

All authors declare no competing interests.

## Data and materials availability

All sequence data listed in TableS3 are from GISAID’s EpiCoV Database.

## Supplementary Materials

**Figures S1 to S2**

**Tables S1 to S5**

## Notes

### Competing Interest Statement

The authors have declared no competing interest.

