## Supplementary Materials for "Computation of Antigenicity Predicts SARS-CoV-2 Vaccine Breakthrough Variants"

**Affiliations:**

**Supplementary Methods**

**Sample collection**

This study was approved by the Institutional Review Board of the University of Hong Kong/Hospital Authority Hong Kong West Cluster (UW 13-265, UW 21-429, and UW 21-214), and the Kowloon West Cluster REC (KW/EX-20-038{144-26}). For pseudovirus neutralisation assay, serum samples were collected from six patients infected with original strains, three patients infected with B.1.617, three Sinovac vaccinees and three BioNTech vaccinees. For live virus neutralisation assay, data were collected from eight patients infected with original strains (D614G), and eight BioNTech vaccinees. All patients were sampled under UW 13-265 and KW/EX-20-038[144-26], and written informed consent was obtained before sample collection. All patients were diagnosed by PCR test with sequencing. The initial laboratory confirmation was performed on nasopharyngeal or sputum specimens at the Public Health Laboratory Centre of Hong Kong. All BioNTech vaccinees were sampled >14 days post second dose under UW 21-214, and all Sinovac vaccinees were sampled under UW 21-429.

**Pseudovirus-based** **neutralisation assay**

Pseudovirus-based virus neutralisation test was performed in HEK293T-hACE2 cells. Murine leukaemia virus (MLV)–based SARS-CoV-2 spike-pseudotyped particles of original strain (reference sequence: EPI_ISL_402124) were purchased from eEnzyme (SCV2-PsV-001). MLV-based SARS-CoV-2 spike pseudovirus particles of B.1.617.1 (reference sequence: EPI_ISL_1547802) were purchased from Codex BioSolutions (CB-97100-160). Sera were serially diluted with pseudovirus solution for 60 minutes before adding to the cells in 96-well plate. After incubation at 37°C for 42 hours, firefly luciferase activity indicating pseudovirus entry was determined using the Steady-Glo Luciferase Assay System (Promega, E2520). The fluorescent signal of plate was obtained by Varioskan LUX Multimode Microplate Reader (Thermo Scientific) and exported by Thermo Scientific SkanIt Software.

**Live virus-based neutralisation assay**

Live virus neutralisation test was performed as described previously, with modifications ^1^. Serum specimens were serially diluted in 2-folds from 1:10 to 1:320. Duplicates of each serum dilution was mixed with B.1.36.27 (with D614G) or B.1.617.1 virus isolates for 1 hour, and the serum-virus mixture was then added to TMPRSS2-VeroE6 cells. After incubation for 3 days, cytopathic effect was visually scored for each well by two independent observers.

**Supplementary Figures & Tables**

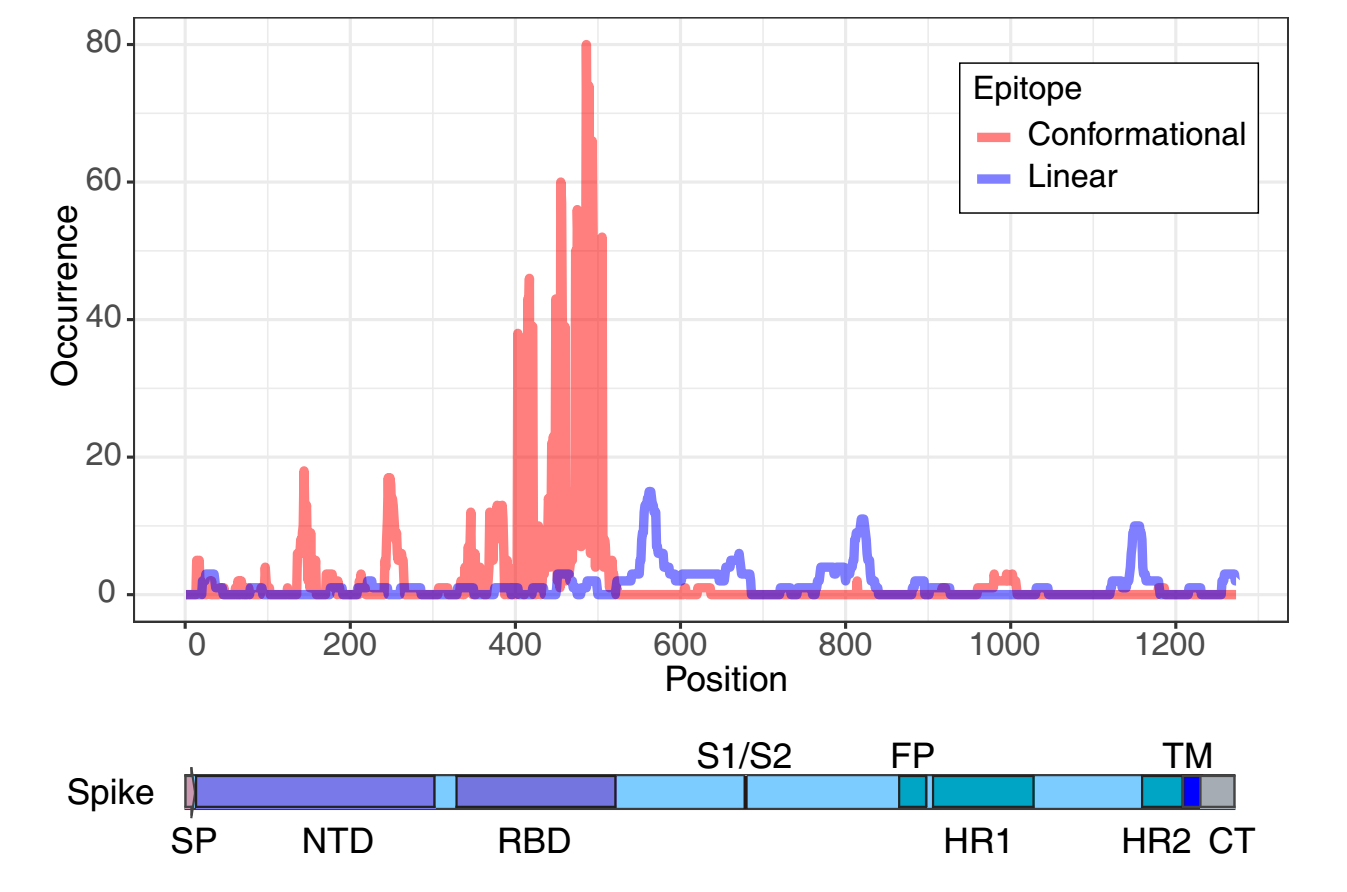

**Figure S1. The epitope mapping of SARS-CoV-2.** Per-residue antigenicity of the Spike protein was plotted by epitope frequency of conformational (N=149) and linear (N=74) epitopes, as well as schematic of the Spike protein coloured by domain. The occurrence showed the number of epitopes containing each position in Table S1. SP, signal peptide; NTD, N-terminal domain; RBD, receptor binding domain; S1/S2, S1/S2 protease cleavage sites; FP, fusion peptide; HR1, heptad repeat 1; HR2, heptad repeat 2; TM, transmembrane domain; CT, cytoplasmic tail.

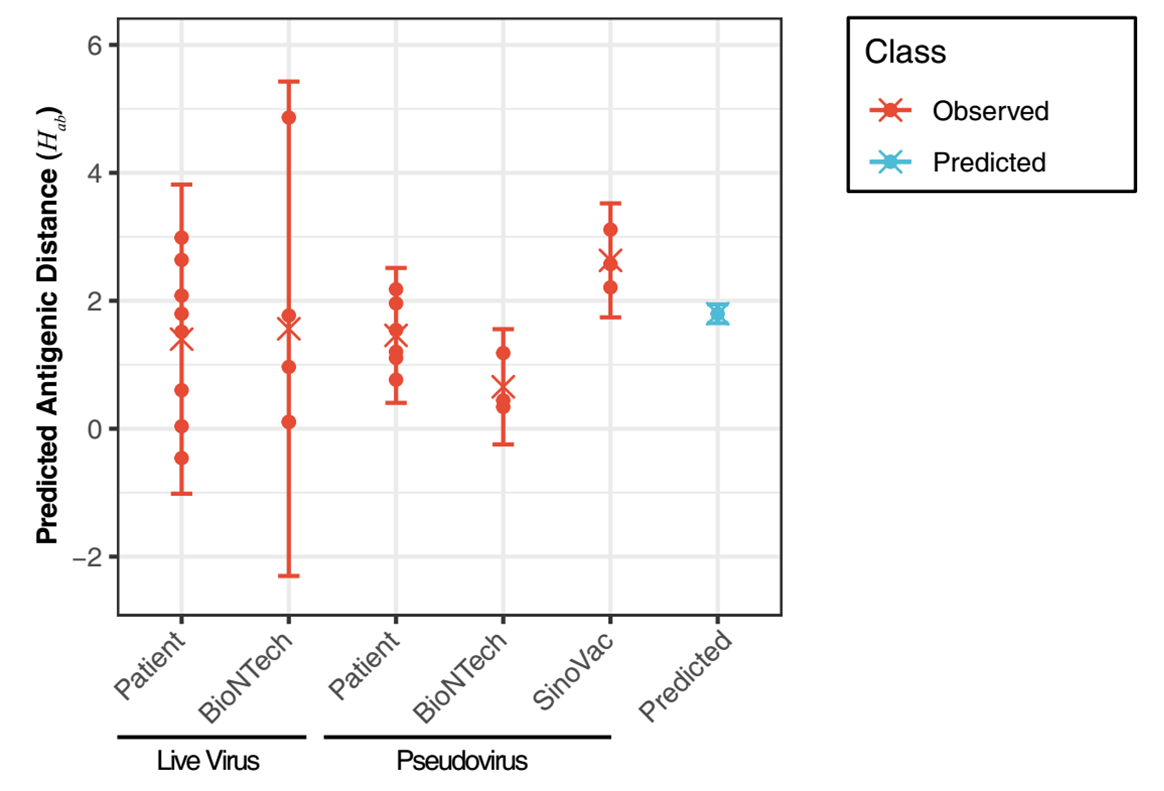

**Figure S2. Predicted versus observed antigenic distances of B.1.617.1 variants in our experiments.** Each point shows the antigenic distance of each individual. Here, we obtained observed antigenic distances by using a pseudovirus neutralisation assay (Detailed results can be found in Fig. S4) and live virus neutralisation data. Error bars show the 95% confidence interval and predicted standard error. “X” shows the mean.

**Table S1. Reported epitopes on SARS-CoV-2 spike protein.**

**Table S2. Observed neutralisation titres of SARS-CoV-2 variants.**

**Table S3. SARS-CoV-2 strains analysed in our study.**

**Table S4. Data sources for Immunogenicity Data**

| **Manufacturer** | **Vaccine** | **NAbFold** | **Reference** |
| --- | --- | --- | --- |
| AstraZeneca | ChAdOx1 nCoV-19 | 0.542 | 10.1016/S0140-6736(20)31604-4 |
| BioNTech | BNT162b2 | 2.372 | 10.1056/NEJMoa2027906 |
| Cadila | ZyCoV-D | 0.907 | 10.1016/j.eclinm.2021.101020 |
| Covaxin | BBV152 | 0.792 | 10.1016/S1473-3099(21)00070-0 |
| Johnson | Ad26.COV2.S | 0.471 | 10.1056/NEJMoa2034201 |
| Longcom | ZF2001 | 2.314 | 10.1016/S1473-3099(21)00127-4 |
| Moderna | mRNA-1273 | 4.139 | 10.1056/NEJMoa2022483 |
| Novavax | NVX-CoV2373 | 3.974 | 10.1056/NEJMoa2026920 |
| Sinopharm | BBIBP-CorV | 0.600 | 10.1016/S1473-3099(20)30831-8 |
| SinoVac | CoronaVac | 0.171 | 10.1016/S1473-3099(20)30843-4 |
| Sputnik | rAd26-S+rAd5-S | 1.400 | 10.1016/S0140-6736(20)31866-3 |

**Table S5. Data sources for Efficacy Data**

**The online tool**

We developed an interactive online tool (<http://jdlab.online>) for the dynamic prediction of the antigenicity of a newly emerging SARS-CoV-2 variants plotted as an antigenic map.

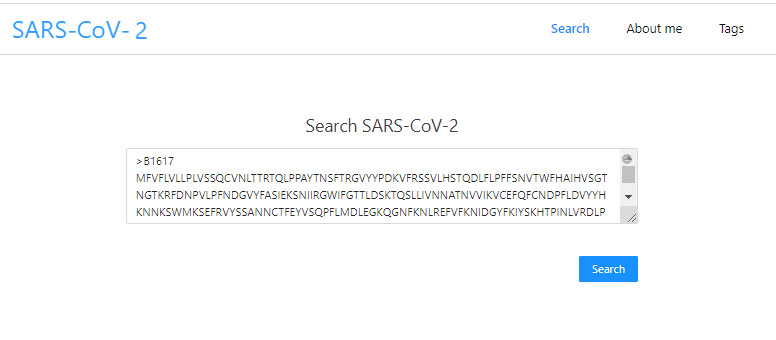

**Figure S3a. The initial interface of the online tool.**

In the online tool, users can search for a variant of interest by inputting one or multiple SARS-CoV-2 Spike amino acid sequences in a fasta format.

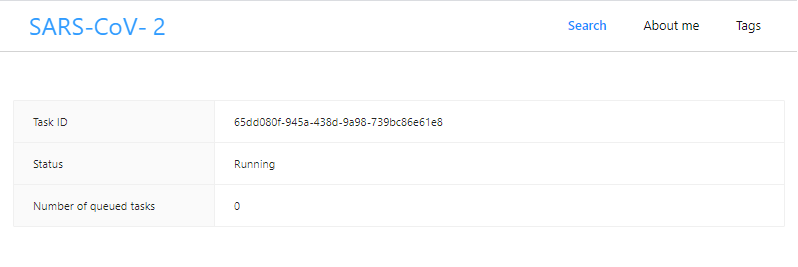

**Figure S3b. The processing interface of the online tool.**

After the search task has completed, the interface then presents the genetic and antigenic maps of the SARS-CoV-2 variants. The two figures above are the genetic distance maps of SARS-CoV-2 variant strains in the Spike protein amino acid sequences and the antigenic distance map of SARS-CoV-2 variant strains in the antigenic epitope amino acid sequences. The other two figures below are the genetic distance maps of SARS-CoV-2 variant strains in the N-terminal domain (NTD) and receptor binding domain (RBD) amino acid sequences and the antigenic distance map of SARS-CoV-2 variant strains in the NTD and RBD amino acid sequences.

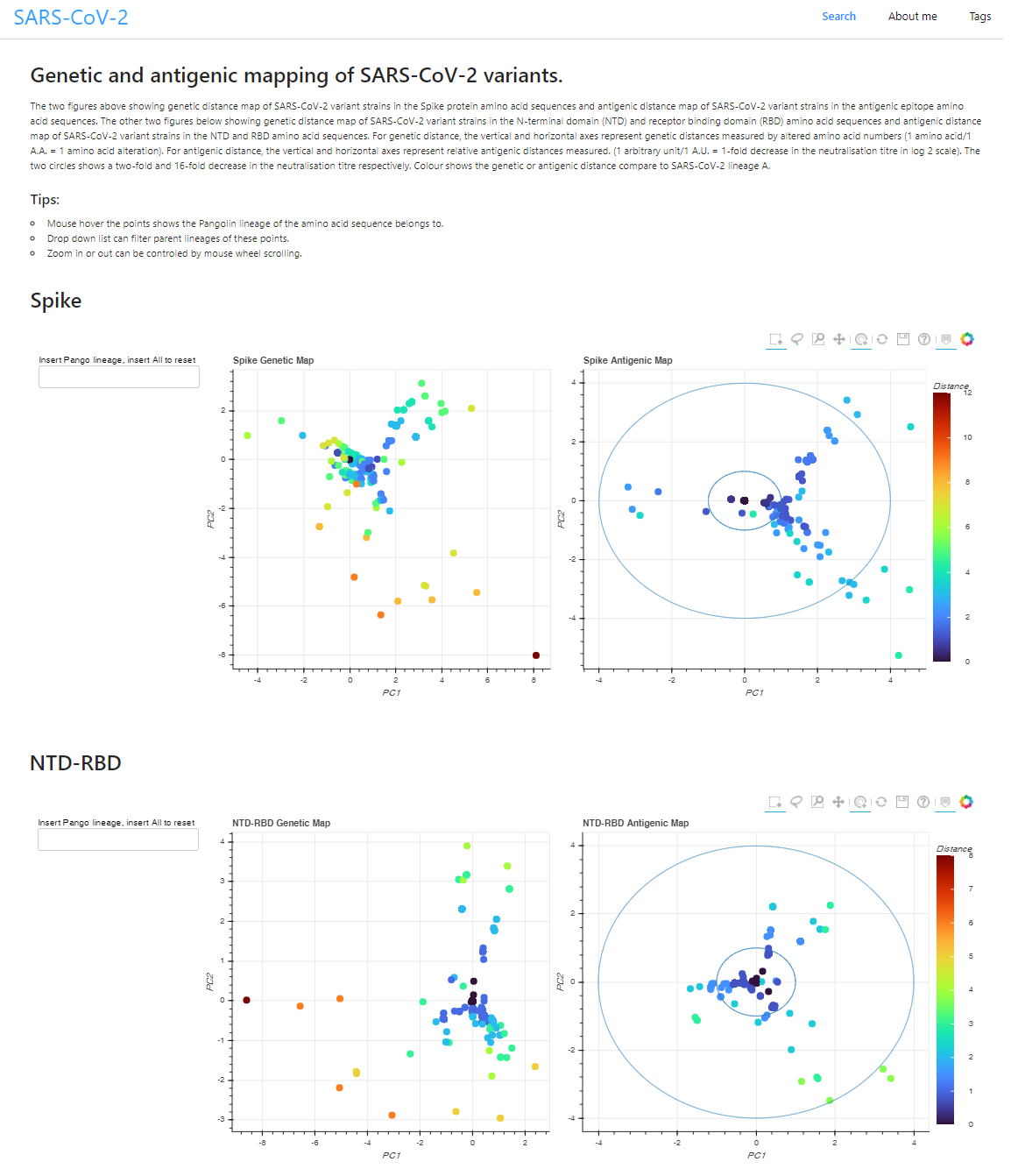

**Figure S3c. The result interface of the online tool.**

For genetic distance, the vertical and horizontal axes represent genetic distances measured by altered amino acid numbers (1 amino acid/1 A.A. = 1 amino acid alteration). For antigenic distance, the vertical and horizontal axes represent the measured relative antigenic distances (1 arbitrary unit/1 A.U. = 1-fold decrease in the neutralization titre in log 2 scale). The two circles show the 2-fold and 16-fold decrease in the neutralization titre, respectively. The colours represent the genetic or antigenic distance compared with SARS-CoV-2 lineage A.

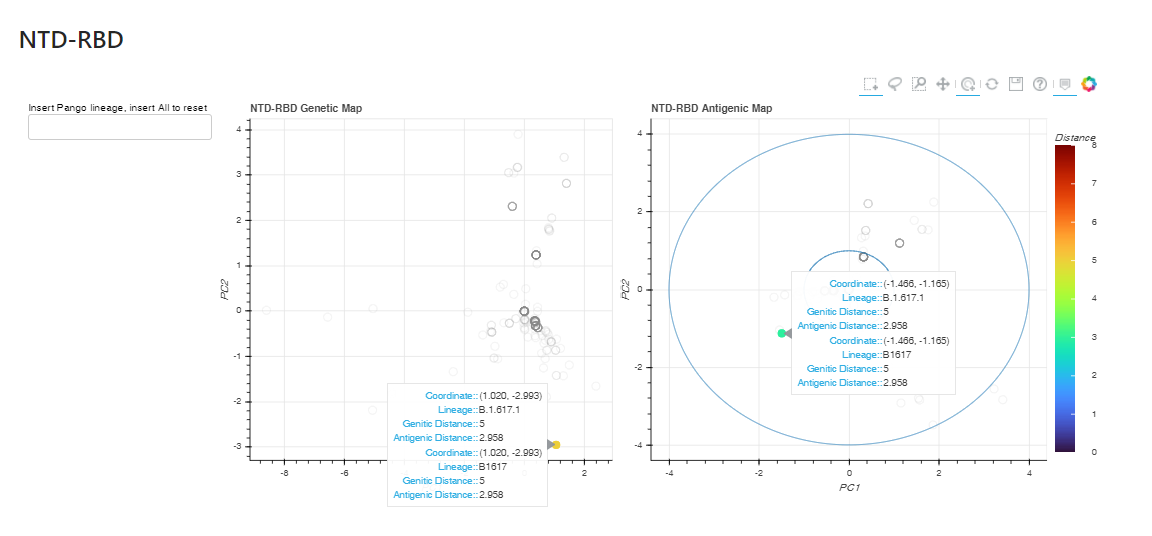

**Figure S3d. The interactive functions of the online tool.**

The Pangolin lineage of the amino acid sequence is shown by hovering the mouse over the points. Strains can be highlighted by searching lineage or user input sequence ID in the text search box. The mouse wheel can be used to zoom in or out.
