## Supplementary material for "Computation of Antigenicity Predicts SARS-CoV-2 Vaccine Breakthrough Variants": Table S1

| Epitope Type | Epitope Position on S Protein | Paired Antibody | Reference DOI |
|---|---|---|---|
| Conformational | Q14,C15,V16,N17,T19,G142,V143,Y144,K147,E156,R158,L244,H245,R246,S247,Y248,L249,T250,P251,G252,S256 | Fab4-18 | 10.1038/s41586-020-2852-1 |
| Conformational | Q14,N17,L18,T76,K77,V143 | Fab2-17 | 10.1101/2021.01.10.426120 |
| Conformational | Q14,Y144,Y145,H146,K147,F157,G252,D253 | COVOX-159 | 10.1016/j.chom.2021.03.005 |
| Conformational | Q14, Y144, H146, K147, N148, N149, W152, M153, E154, E156, F157, R158, R246, Y248, L249, P251, G252, D253 | Fab4-8 | 10.1016/j.cell.2021.02.032 |
| Conformational | Q14,C15,Y144,H146,K147,E154,E156,R158,R246,Y248,L249,T250,P251,D253,S254 | Fab5-24 | 10.1101/2021.01.10.426120 |
| Conformational | V16,N17,T19,Y144,R246,S247,Y248,T250,P251,G252,D253,S254,S255,S256,G257 | S2L28 | 10.1101/2021.01.10.426120 |
| Conformational | V16,N17,T20,Y144,Y145,H146,K147,N148,S155,R158,R246,L249,T250,P251,G252,D253 | S2M28 | 10.1016/j.cell.2021.03.028 |
| Conformational | V16,N17,F140,G142,V143,Y144,Y145,H146,K147,N148,W152,E154,E156,R158,L244,H245,R246,L249,P251 | S2X333 | 10.1016/j.cell.2021.03.028 |
| Conformational | A27,Y28,T29,N30,F32,N61,W64,H66,I68,H69,K97,F186,N211,L212,V213,R214,D215,L216,P217,Q218,S605,N606 | DH1052 | 10.1016/j.cell.2021.03.028 |
| Conformational | N30,F32,W64,H66,I68,K97,N185,K187,N211,V213,R214 | CoV2-2490 | 10.1016/j.cell.2021.06.021 |
| Conformational | V42,F43,R44,S45,V47,L368,Y369,N370,F377,K378,C379,G381,S383,P384,T385,L390,D428,F429,T430 | CR3022-New | 10.1016/j.cell.2021.05.032 |
| Conformational | S71,K97,S98,T124,Y145,H146,K147,K150,S151,W152,E180,G181,K182,Q183,N185,V213,H245,S247,Y248,L249,T259,A260,A262 | P008_056 | 10.1038/s41467-020-19146-5 |
| Conformational | F92,A93,S94,T95,E96,K97,S98,N99,I100,I101,R102,C136,N137,D138,P139,F140,L141,G142,V143,Y144,V171,S172,Q173,P174,F175,L176,M177,D178,L179,L242,A243,L244,H245,R246,S247,Y248,L249,T250,P251,G252,D253,S254,S255,S256,G257,W258,T259,A260,G261,A262,A263,A264 | Ab88 | 10.1126/sciadv.abg7607 |

| | | | |
|---|---|---|---|
| Conformational | C136,N137,D138,P139,F140,L141,G142,V143,Y144,V171,S172,Q173,P174,F175,L176,M177,D178,L242,A243,L244,H245,R246,S247,Y248,L249,T250,P251,G252,D253,S254,S255,S256,G257,W258,T259,A260,G261,A262,A263,A264 | Ab55 | 10.1126/scitranslmed.abf1906 |
| Conformational | C136,N137,D138,P139,F140,L141,G142,V143,Y144,V171,S172,Q173,P174,F175,L176,M177,D178,L179,L242,A243,L244,H245,R246,S247,Y248,L249,T250,P251,G252,D253,S254,S255,S25 | Ab60 | 10.1126/scitranslmed.abf1906 |
| Conformational | C136,N137,D138,P139,F140,L141,G142,V143,Y144,L242,L244,H245,R246,S247,Y248,L249,T250,P251,G252,D253,S254,S255,S256,G257,W258,T259,A260,G261,A262,A263,A264,Y265 | Ab89 | 10.1126/scitranslmed.abf1906 |
| Conformational | C136,N137,D138,P139,F140,L141,G142,V143,Y144,L242,L244,H245,R246,S247,Y248,L249,T250,P251,G252,D253,S254,S255,S256,G257,W258,T259,A260,G261,A262,A263,A264,Y265 | Ab130 | 10.1126/scitranslmed.abf1906 |
| Conformational | F140,G142,V143,Y145,H146,N148,N149,W152,E154,F157,A243,L244,H245 | DH1050.1 | 10.1126/scitranslmed.abf1906 |
| Conformational | Y144,Y145,H146,K147,R246,S247,Y248,L249,T250,P251,G252,S255 | CM25 | 10.1016/j.cell.2021.06.021 |
| Conformational | Y144,Y145,H146,K147,N148,K150,R246,S247,Y248,L249,T250,P251,G252,D253,S254 | Fab2-51 | 10.1126/science.abg5268 |
| Conformational | Y144,Y145,H146,K147,K150,W152,H245,R246,Y248,L249,T250,P251,G252,S254,S255,S256 | Fab1-87 | 10.1101/2021.01.10.426120 |
| Conformational | Y144,Y145,H146,K147,K150,W152,H245,R246,S247,Y248,L249 | 4A8 | 10.1101/2021.01.10.426120 |
| Conformational | Y144,Y145,H146,K147,K150,W152,R246,S247,Y248,L249,S256 | FC05 | 10.1126/science.abc6952 |
| Conformational | Y145,K147,W152,Y248 | CM17 | 10.1038/s41422-020-00446-w |
| Conformational | Y144,W152,R246,Y248 | CM30 | 10.1126/science.abg5268 |
| Conformational | T333,N334,L335,P337,G339,E340,V341,N343,A344,T345,K346,E354,K356,R357,I358,S359,N360,C361,N440,L441,K444,R509 | S309 | 10.1126/science.abg5268 |

| | | | |
|---|---|---|---|
| Conformational | N334,L335,P337,G339,E340,N343,A344,T345,R346,K356,R357,S359,C361,L441 | S309-New | 10.1038/s41586-020-2349-y |
| Conformational | G339,F342,N343,T345,R346,V367,L368,S371,S373,F374,W436,N437,S438,N440,L441,K444,N448,Y449,N450,Q498 | BG10-19 | 10.1016/j.cell.2020.09.037 |
| Conformational | S375,T376,K378,R408,Q409,Q414,K417,Y449,L452,L455,F456,A475,G482,E484,G485,F486,Y4 | BG7-20 | 10.1016/j.cell.2021.04.032 |
| Conformational | G339,F342,N343,V367,S371,A372,S373,F374,Y449,L455,F456,V483,E484,G485,F486,N487,Y489,F490,Q493,S494 | C144 | 10.1016/j.cell.2021.04.032 |
| Conformational | F342,N343,L368,S371,A372,S373,F374,W436,L441,K444,G446,Y449,L452,L455,F456,E484,G485,F486,Y489,F490,L492,Q493,S494 | S2M11-New | 10.1038/s41586-020-2852-1 |
| Conformational | F342,N343,L368,S371,A372,S373,W436,N440,L441 | S2M11 | 10.1016/j.cell.2021.03.028 |
| Conformational | N343,A344,T345,R346,S373,W436,N437,N440,L441,S443,K444,V445,N448,N450,R509 | CV38-142 | 10.1126/science.abe3354 |
| Conformational | T345,R346,S438,N439,N440,L441,P499 | C135 | 10.1016/j.chom.2021.04.005 |
| Conformational | T345,R346,L441,D442,N448,Y449,N450,L452,F490,S494,Q498,P499,T500,R509 | C110 | 10.1038/s41586-020-2852-1 |
| Conformational | T345,N439,N440,S443,K444,V445,G446,G447,N450,Q498,P499,T500,Q506 | Fab2-7 | 10.1038/s41586-020-2852-1 |
| Conformational | R346,F347,S349,Y351,K444,G446,G447,N448,Y449,N450,Y451,L452,T470,E484,F490,L492,Q493,S494 | CV07-270 | 10.1016/j.str.2021.05.014 |
| Conformational | R346,Y351,K444,Y449,N450,L452,T470,I472,N481,G482,V483,E484,F490,L492,S494 | 47D1 | 10.1016/j.cell.2020.09.049 |
| Conformational | R346,N439,N440,S443,K444,V445,G446,G447,N450,Q498,P499,T500,N501,G502,Q506 | Ly-CoV1404 | 10.1016/j.celrep.2021.109109 |
| Conformational | R346,N440,L441,K444,V445,G446,N448,Y449,Q498 | REGN10987 | 10.1101/2021.04.30.442182. |
| Conformational | R346,K444,G446,G447,N448,Y449,N450,L452,V483,E484,G485,F490,S494 | P2B-2F6 | 10.1126/science.abd0827 |
| Conformational | R346,K444,Y449,N450,L452,I472,N481,G482,V483,E484,F490,L492 | BD-368-2-New | 10.1038/s41586-020-2380-z |
| Conformational | Y351,A372,S375,T376,R408,Y449,L452,T470,I472,Y473,G482,V483,E484,G485,F486,Y489,F490,L492,Q493,S494,V503,G504,Y508 | BG1-24 | 10.1038/s41422-021-00514-9 |
| Conformational | Y351,K444,V445,G446,G447,N448,Y449,N450,L452,T470,E484,F490,L492,Q493,S494,Q498 | Fab1-57 | 10.1016/j.cell.2021.04.032 |
| Conformational | Y351,G446,Y449,N450,L452,F456,T470,T478,P479,C480,N481,G482,V483,E484,G485,F486,N487,C488,Y489,F490,P491,L492,Q493,S494 | DH1043 | 10.1016/j.str.2021.05.014 |
| Conformational | Y351,Y449,L455,T470,N481,G482,V483,E484,G485,F486,C488,Y489,F490,L492,Q493,S494 | Ly-CoV555 (Ab169) | 10.1016/j.cell.2021.06.021 |

| | | | |
|---|---|---|---|
| Conformational | W353,N354,R355,K356,R357,S359,N360,N394,Y396,P426,D428,K462,P463,F464,E465,R466,I468,E516,L518,H519,A520,T523 | COVOX-45 | 10.1126/scitranslmed.abf1906 |
| Conformational | W353,R355,R357,Y396,P426,D427,D428,F429,K462,P463,F464,R466,S514,E516,L518,H519,A520,P521 | S2H97 | 10.1016/j.cell.2021.02.032 |
| Conformational | S366,Y369,N370,F374,F377,C379,Y380,G381,V382,S383,P384,T385,K386,N388,L390,F392,P412,D427,D428,F429,L517 | EY6A-New | 10.1101/2021.04.07.438818 |
| Conformational | Y369,N370,S371,A372,F374,S375,T376,F377,K378,C379,Y380,G381,V382,S383,P384,T385,K386,D389,L390,F392,D427,D428,F429,T430,F515,E516,L517,H519 | CR3022 | 10.1016/j.cell.2021.03.055 |
| Conformational | Y369,N370,S371,A372,F374,S375,T376,F377,K378,C379,Y380,V382,S383,P384,T385,G404,D405,R408,T500,N501,G502,V503,G504,Q506 | S2X259 | 10.1126/science.abb7269 |
| Conformational | Y369,N370,S371,A372,F374,S375,T376,F377,K378,C379,S383,P384,T385,R408,Q414 | S2A4 | 10.1101/2021.04.07.438818 |
| Conformational | Y369,N370,S371,A372,F374,S375,T376,F377,K378,C379,S383,P384,D405,R408,Q409,Q414,T415,G416,N501,V503,G504,Y505 | DH1047 | 10.1016/j.cell.2020.09.037 |
| Conformational | Y369,N370,S371,F377,K378,C379,Y380,G381,V382,S383,P384,T385,R408,P412,G413,Q414,T415,G416,D427,D428,F429 | COVA1-16-New | 10.1016/j.cell.2021.06.021 |
| Conformational | Y369,N370,A372,F374,K378,P384 | C126 | 10.1016/j.chom.2021.04.005 |
| Conformational | Y369,N370,F374,S375,T376,F377,K378,C379,Y380,G381,V382,S383,P384,T385,K386,L390,R408,D428,T430,L517,L518 | H11-D4 | 10.1016/j.celrep.2021.109604 |
| Conformational | Y369,N370,F377,K378,C379,Y380,G381,V382,S383,P384,T385,K386,N388,L390,F392,P412,G413,Q414,P426,D427,D428,F429,T430,F515,L517 | S304 | - |
| Conformational | Y369,S371,F377,K378,C379,Y380,G381,V382,S383,P384,T385,R408,P412,G413,Q414,T415,G416,D427,D428,F429 | COVA1-16 | 10.1016/j.cell.2020.09.037;10.1038/s41586-020-2349-y |
| Conformational | Y369,S375,F377,K378,C379,Y380,G381,V382,S383,P384,T385,K386,F392,P412,G413,D427,D428,F429,L517 | EY6A | 10.1016/j.immuni.2020.10.023 |
| Conformational | A372,N440,K444,V445,Y449,N450,L455,T470,E471,N481,G482,V483,E484,G485,F486,Y489,F490,Q493,S494,T500 | C002 | 10.1038/s41594-020-0480-y |
| Conformational | F374,S375,T376,F377,K378,C379,Y380,G381,V382,S383,P384,T385,K386,R408,N437,V503,G504,Y508 | CV2-75 | 10.1038/s41586-020-2852-1 |

| | | | |
|---|---|---|---|
| Conformational | R403,D405,E406,R408,Q409,T415,G416,K417,D420,Y421,Y449,Y453,L455,F456,R457,K458,S459,N460,Y473,Q474,A475,G476,S477,E484,F486,N487,Y489,F490,L492,Q493,S494,Y495,G496,Q498,T500,N501,G502,V503,G504,Y505 | B38-New | 10.1016/j.celrep.2021.109353 |
| Conformational | R403,D405,E406,R408,Q409,T415,G416,K417,D420,Y421,Y449,Y453,L455,F456,R457,K458,S459,N460,Y473,Q474,A475,G476,S477,F486,N487,Y489,F490,Q493,S494,Y495,G496,Q498,T500,N501,G502,G504,Y505 | CB6-New | 10.1038/s41467-020-19231-9 |
| Conformational | R403,D405,E406,R408,Q409,T415,G416,K417,D420,Y421,Y453,L455,F456,R457,K458,N460,Y473,Q474,A475,G476,S477,F486,N487,Y489,Y495,G496,Q498,T500,N501,G502,V503,Y505 | COVOX-158 | 10.1038/s41467-020-19231-9 |
| Conformational | R403,D405,E406,R408,Q409,T415,G416,K417,D420,Y421,L455,F456,R457,K458,N460,Y473,Q474,A475,G476,S477,F486,N487,Y489,Q493,Y495,G502,Y505 | CB6 | 10.1016/j.cell.2021.02.032 |
| Conformational | R403,D405,E406,R408,T415,G416,K417,D420,Y421,Y449,L455,F456,R457,K458,N460,Y473,Q474,A475,G476,S477,N487,Y489,Q493,S494,G496,T500,N501,G502,V503,Y505 | P2B-1A10 | 10.1038/s41586-020-2381-y |
| Conformational | R403,D405,R408,Q409,T415,G416,K417,Y421,Y449,Y453,L455,F456,G485,F486,N487,Y489,Q493,S494,Y495,G496,N501,Y505 | P5A-1B6 | 10.1038/s41422-021-00487-9 |
| Conformational | R403,D405,R408,T415,G416,K417,D420,Y421,Y453,L455,R457,K458,N460,Y473,Q474,A475,G476,S477,F486,N487,Y489,Q493,Y495,G496,Q498,T500,N501,G502,Y505 | COVOX-150 | 10.1038/s41422-021-00487-9 |
| Conformational | R403,D405,T415,G416,K417,D420,Y421,Y453,L455,F456,R457,K458,S459,N460,Y473,A475,G476,S477,F486,N487,Y489,Q493,G496,Q498,T500,N501,G502,Y505 | BD-604-new | 10.1016/j.cell.2021.02.032 |
| Conformational | R403,D405,T415,G416,K417,D420,Y421,Y453,L455,F456,R457,K458,N460,Y473,Q474,A475,G476,S477,T478,F486,N487,Y489,Q493,N501,Y505 | BD-629 | 10.1038/s41422-021-00514-9 |
| Conformational | R403,D405,T415,G416,K417,D420,Y421,L455,F456,R457,K458,S459,N460,Y473,Q474,A475,G476,S477,F486,N487,Y489,N501,G502,Y505 | Ly-CoV488 (Ab133) | 10.1016/j.cell.2020.09.035 |
| Conformational | R403,D405,T415,G416,K417,D420,Y421,Y453,L455,F456,R457,K458,N460,Y473,A475,G476,S477,F486,N487,Y489,Y495,N501,Y505 | CC12.3 | 10.1126/scitranslmed.abf1906 |
| Conformational | R403,D405,T415,G416,K417,D420,Y421,Y453,L455,F456,R457,K458,N460,Y473,A475,G476,S477,F486,N487,Y489,Q493,S494,Y495,G496,T500,N501,G502,Y505 | COVA2-04 | 10.1126/science.abd2321 |

| | | | |
|---|---|---|---|
| Conformational | R403,T415,G416,K417,D420,Y421,Y453,L455,F456,R457,K458,N460,Y473,Q474,A475,G476,F486,N487,Y489,Y495,G496,N501,Y505 | CC12.3-New | 10.1016/j.celrep.2020.108274 |
| Conformational | R403,D405,T415,G416,K417,D420,Y421,Y453,L455,F456,R457,K458,N460,Y473,A475,G476,F486,N487,Y489,Q493,S494,Y495,G496,Q498,T500,N501,G502,G504,Y505 | 910-30 | 10.1126/science.abe6230 |
| Conformational | R403,D405,T415,G416,K417,D420,Y421,Y453,L455,F456,R457,K458,S459,N460,Y473,Q474,A475,G476,S477,F486,N487,Y489,F490,Q493,S494,Y495,G496,Q498,T500,N501,G502,Y505 | CV30 | 10.1101/2020.12.31.424987 |
| Conformational | R403,D405,T415,G416,K417,D420,Y421,L455,F456,R457,K458,N460,Y473,A475,G476,S477,E484,F486,N487,Y489,F490,Q493,S494,G496,Q498,N501,G502,Y505 | BD-508 | 10.1038/s41467-020-19231-9 |
| Conformational | R403,E406,R408,Q409,G416,K417,Y449,Y453,L455,F456,F486,N487,Y489,Q493,S494,Y495,G496,Q498,T500,N501,G502,Y505 | P2B-1A1 | 10.1038/s41422-021-00514-9 |
| Conformational | R403,E406,Q409,T415,G416,K417,D420,Y421,Y453,L455,F456,R457,K458,N460,Y473,Q474,A475,G476,F486,N487,Y489,Q493,S494,Y495,G496,Q498,T500,N501,G502,Y505 | COVOX-269 | 10.1038/s41422-021-00487-9 |
| Conformational | R403,R408,Q409,Q414,T415,G416,K417,D420,Y421,G446,Y449,F456,A475,G476,S477,F486,N487,Y489,Q493,S494,G496,Q498,N501,Y505 | P5A-2G9 | 10.1016/j.cell.2021.02.032 |
| Conformational | R403,R408,T415,G416,K417,D420,Y421,Y453,L455,F456,R457,K458,N460,Y473,Q474,A475,G476,F486,N487,Y489,Q493,S494,Y495,G496,Q498,T500,N501,G502,Y505 | C1A-B3 | 10.1038/s41422-021-00487-9 |
| Conformational | R403,Q409,T415,G416,K417,D420,Y421,L455,F456,R457,K458,S459,N460,Y473,Q474,A475,G476,S477,F486,N487,Y489,F490,Q493,Y495,G496,Q498,N501,G502,Y505 | B38 | 10.1016/j.cell.2021.03.027 |
| Conformational | R403,T415,G416,K417,D420,Y421,Y453,L455,F456,R457,K458,S459,N460,Y473,Q474,A475,G476,S477,F486,N487,Y489,Q493,G502,Y505 | P2C-1F11 | 10.1126/science.abc2241 |
| Conformational | R403,T415,G416,K417,D420,Y421,Y453,L455,F456,R457,K458,S459,N460,Y473,Q474,A475,G476,S477,F486,N487,Y489,Y495,G496,Q498,T500,N501,G502,Y505 | COVOX-40 | 10.1038/s41467-020-20501-9 |
| Conformational | R403,T415,G416,K417,D420,Y421,Y453,L455,F456,R457,K458,N460,Y473,Q474,A475,G476,S477,F486,N487,Y489,Q493,Y505 | BG4-25 | 10.1016/j.cell.2021.02.032 |

| | | | |
|---|---|---|---|
| Conformational | R403,T415,G416,K417,D420,Y421,Y453,L455,F456,R457,K458,N460,Y473,Q474,A475,G476,S477,F486,N487,Y489,Q493,S494,Y495,G496,Q498,T500,N501,G502,Y505 | P4A1 | 10.1016/j.cell.2021.04.032 |
| Conformational | R403,T415,G416,K417,D420,Y421,Y453,L455,F456,R457,K458,N460,Y473,Q474,A475,G476,F486,N487,Y489,Q493,S494,Y495,G496,Q498,T500,N501,G502,Y505 | C1A-C2 | 10.1038/s41467-021-22926-2 |
| Conformational | R403,T415,G416,K417,D420,Y421,Y453,L455,F456,R457,K458,N460,Y473,A475,G476,S477,F486,N487,Y489,Q493,S494,Y495,G496,Q498,T500,N501,G502,V503,Y505 | BD-236 | 10.1016/j.cell.2021.03.027 |
| Conformational | R403,T415,G416,K417,D420,Y421,Y453,L455,F456,R457,K458,N460,Y473,A475,G476,S477,F486,N487,Y489,Q493,Q498,T500,N501,G502,Y505 | C1A-F10 | 10.1016/j.cell.2020.09.035 |
| Conformational | R403,T415,G416,K417,D420,Y421,Y453,L455,F456,R457,K458,N460,Y473,A475,G476,S477,F486,N487,Y489,T500,N501,G502,Y505 | C102 | 10.1016/j.cell.2021.03.027 |
| Conformational | R403,T415,G416,K417,D420,Y421,Y453,L455,F456,R457,K458,N460,Y473,A475,G476,F486,N487,Y489,Q493,S494,Y495,G496,Q498,T500,N501,G502,Y505 | C1A-B12 | 10.1038/s41586-020-2852-1 |
| Conformational | R403,T415,G416,K417,D420,Y421,Y453,L455,R457,K458,S459,N460,Y473,Q474,A475,G476,S477,F486,N487,Y489,Q493,Q498,T500,N501,G502,V503,Y505 | BD-604 | 10.1016/j.cell.2021.03.027 |
| Conformational | R403,T415,G416,K417,D420,Y421,Y453,L455,R457,K458,N460,Y473,A475,G476,S477,F486,N487,Y489,Q493,S494,Y495,G496,Q498,T500,N501,G502,V503,Y505 | Ly-CoV481 (Ab128) | 10.1016/j.cell.2020.09.035 |
| Conformational | R403,T415,G416,K417,D420,Y421,L455,F456,R457,N460,Y473,A475,G476,F486,N487,Y489,Q493,G496,T500,N501,G502,Y505 | P5A-1B8 | 10.1126/scitranslmed.abf1906 |
| Conformational | R403,T415,K417,D420,Y421,L455,R457,K458,N460,Y473,Q474,A475,G476,S477,T478,F486,N487,Y489,N501,G502,Y505 | BD-515 | 10.1038/s41422-021-00487-9 |
| Conformational | R403,K417,Y449,N450,L452,Y453,L455,F456,E484,G485,F486,Y489,F490,L492,Q493,S494,Y495,Y505 | CT-P59 | 10.1038/s41422-021-00514-9 |
| Conformational | R403,K417,Y449,L452,Y453,L455,F456,E484,G485,F486,C488,Y489,F490,L492,Q493,Y505 | P5A-2G7 | 10.1038/s41467-020-20602-5 |
| Conformational | R403,K417,Y453,L455,F456,E484,G485,F486,N487,C488,Y489,Q493,N501,G502,Y505 | COVOX-88 | 10.1038/s41422-021-00487-9 |
| Conformational | R403,V445,G446,Y449,Y453,L455,F456,N487,Y489,Q493,Y495,G496,Q498,P499,T500,N501,G502,Y505 | S2H14 | 10.1016/j.cell.2021.02.032 |

| | | | |
|---|---|---|---|
| Conformational | G404,D405,E406,V407,R408,Q414,T415,Y449,L452,L455,A475,E484,G485,F486,Y489,F490,L492,Q493,S494,V503,G504,Y505,Y508 | C121 | 10.1016/j.cell.2020.09.037 |
| Conformational | D405,R408,T415,G416,Y421,F456,R457,K458,N460,Y473,Q474,A475,G476,F486,N487,T500,N501,G502,Y505 | C105 | 10.1038/s41586-020-2852-1 |
| Conformational | D405,E406,T415,K417,D420,Y453,L455,F456,R457,A475,S477,F486,Y489,N487,Q493,G496,Q498,T500,N501,G502,Y505 | CC12.1 | 10.1016/j.cell.2020.06.025; 10.1038/s41586-020-2456-9 |
| Conformational | D405,K417,D420,L455,F456,N460,I472,Y473,A475,G476,F486,N487,Y489,G504 | Ly-CoV016 | 10.1126/science.abd2321 |
| Conformational | R408,K444,T470,V483,F486,F490,V503,G504 | C104 | 10.1016/j.xcrm.2021.100255 |
| Conformational | T415,G416,K417,D420,Y421,Y453,L455,F456,R457,K458,S459,N460,Y473,Q474,A475,G476,S477,F486,N487,Y489,Q493,S494,Y495,G496,Q498,T500,N501,G502,Y505 | CV30 | 10.1038/s41586-020-2852-1 |
| Conformational | T415,G416,K417,D420,Y421,L455,F456,R457,K458,N460,Y473,Q474,A475,G476,S477,F486,N487,G496,Y505 | P5A-3A1 | 10.1038/s41467-020-19231-9 |
| Conformational | T415,G416,K417,D420,Y432,Y453,L455,R457,K458,N460,Y473,A475,G476,F486,N487,Y489,S494,Y495,G496,Q498,T500,N501,G502,Y504 | SET90-C11 | 10.1038/s41422-021-00487-9 |
| Conformational | T415,Y421,A475,G476,N487,S494,G502 | BG1-22 | 10.1016/j.celrep.2021.109433 |
| Conformational | K417,D420,L455,F456,N460,Y473,A475 | C105-New | 10.1016/j.cell.2021.04.032 |
| Conformational | K417,Y453,L455,F456,E484,G485,F486,N487,C488,Y489,Q493 | REGN10933 | 10.1038/s41467-021-24435-8 |
| Conformational | Y421,F456,R457,Y473,A475,G476,S477,E484,G485,F486,N487,Y489,Q493 | P5A-2F11 | 10.1126/science.abd0827 |
| Conformational | N439,N440,S443,K444,V445,G446,G447,Y449,N450,S494,P499,T500,Q506 | BG7-15 | 10.1038/s41422-021-00487-9 |
| Conformational | N440,L441,S443,K444,V445,G446,G447,N448,Y449,N450,L452,F490,L492,Q493,S494,Y495,G496 | COVOX-75 | 10.1016/j.cell.2021.04.032 |
| Conformational | K444,V445,G446,Y449,N450,E484,Q493,S494,Q498,G504, Y505 | C119 | 10.1016/j.cell.2021.02.032 |
| Conformational | K444,G446,Y449,N450,L452,N481,G482,V483,E484,G485,F490 | BD-368-2 | 10.1038/s41586-020-2852-1 |
| Conformational | K444,G447,N448,Y449,L452,F490,S494 | C110-New | 10.1016/j.cell.2020.09.035 |
| Conformational | V445,G446,Y449,F456,T478,N481,V483,E484,G485,F486,N487,Y489,F490,L492,Q493,S494,Q498,T500 | P2C-1A3 | 10.1038/s41467-021-24435-8 |
| Conformational | G446,G447,Y449,F456,T470,V483,E484,G485,F486,C488,Y489,F490,P491,L492,Q493,S494,Q498 | CV05-163 | 10.1038/s41467-020-20501-9 |

| | | | |
|---|---|---|---|
| Conformational | G446,N448,Y449,L452,E484,G485,F486,N487,Y489,F490,L492,Q493,S494 | P5A-1B9 | 10.1126/science.abh1139 |
| Conformational | G446,Y449,L452,T478,V483,E484,G485,F486,N487,Y489,F490,L492,Q493,S494,G496,Q498 | Fab2-15 | 10.1038/s41422-021-00487-9 |
| Conformational | G446,Y449,N481,G482,V483,E484,G485,F486,F490,S494 | S2H13 | 10.1016/j.celrep.2021.108950 |
| Conformational | G446,Y449,E484,G485,F486,Y489,F490,L492,Q493,S494,G496,Q498,N501,Y505 | BD-23 | 10.1016/j.cell.2020.09.037 |
| Conformational | G446,Y449,Y453,L455,F456,A475,G476,S477,T478,G485,F486,N487,Y489,Q493,Y495,Q498,N501,Y505 | CV07-250 | 10.1016/j.cell.2020.05.025 |
| Conformational | G446,Y449,F456,A475,V483,E484,G485,F486,N487,Y489,Q493 | COVA2-39 | 10.1016/j.cell.2020.09.049 |
| Conformational | Y449,L452,T470,E471,I472,N481,G482,V483,E484,G485,F486,F490,L492,Q493,S494 | DH1041 | 10.1016/j.celrep.2020.108274 |
| Conformational | Y449,Y453,L455,F456,E484,G485,F486,Y489,F490,L492,Q493,S494 | Fab2-4 | 10.1016/j.cell.2021.06.021 |
| Conformational | Y449,L455,F456,V483,E484,G485,F486,Y489,F490,Q493,S494 | H4 | 10.1038/s41586-020-2571-7 |
| Conformational | Y449,L455,F456,V483,E484,G485,F486,Y489,F490,L492,Q493,S494 | COVOX-316 | 10.1016/j.celrep.2021.108950 |
| Conformational | L452,L455,F456,I472,N481,G482,V483,E484,G485,F486,Y489,F490 | COVOX-384 | 10.1016/j.cell.2021.02.032 |
| Conformational | L452,I472,V483,E484,G485,F486,F490,Q493,S494 | Ly-CoV555 | 10.1016/j.cell.2021.02.032 |
| Conformational | L455,F456,K458,Y473,A475,G476,S477,T478,G485,F486,N487,Y489,Q493 | COVOX-253 | 10.1016/j.xcrm.2021.100255 |
| Conformational | L455,K458,Y473,A475,G476,S477,T478,G485,F486,N487,C488,Y489,Q493 | COVOX-253H55L | 10.1016/j.cell.2021.02.032 |
| Conformational | L455,F456,E484,F486,F490,Q493 | C002-New | 10.1016/j.cell.2021.02.032 |
| Conformational | L455,Y473,A475,G476,S477,E484,G485,F486,N487,C488,Y489 | S2E12 | 10.1038/s41467-021-24435-8 |
| Conformational | F456,Y473,A475,G476,S477,T478,V483,E484,G485,F486,N487,C488,Y489 | BD-623 | 10.1126/science.abe3354 |
| Conformational | Y473,A475,T478,F486,N487 | COVOX-253H165L | 10.1038/s41422-021-00514-9 |
| Conformational | V483,E484,F486,Y489 | P5A-3C12 | 10.1016/j.cell.2021.02.032 |
| Conformational | K814,Y917,Q920,K921 | COV2-2002/COV2-2333 | 10.1038/s41422-021-00487-9 |
| Conformational | K814,I980,R995,Q1002 | CnC2t1p1_B10 | 10.1016/j.celrep.2021.109604 |
| Conformational | I472,Y473,Q474,A475,G476,S477,T478,P479,C480,N481,G482,V483,E484,G485,F486,N487,C488,Y489,F490,P491,L492,Q493,S494,Y495 | Ab18 | 10.1016/j.celrep.2021.109604 |

| | | | |
|---|---|---|---|
| Conformational | D467,I468,S469,T470,E471,I472,Y473,Q474,A475,G476,S477,T478,P479,C480,N481,G482,V483,E484,G485,F486,N487,C488,Y489,F490,P491,L492,Q493,S494,Y495,G496,F497,Q498,P499,T500,N501,G502,V503,G504,Y505,Q506,P507,Y508,R509,V510,V511,V512,L513 | Ab104 | 10.1126/scitranslmed.abf1906 |
| Conformational | D467,I468,S469,T470,E471,I472,Y473,Q474,A475,G476,S477,T478,P479,C480,N481,G482,V483,E484,G485,F486,N487,C488,Y489 | Ab116 | 10.1126/scitranslmed.abf1906 |
| Conformational | V433,I434,A435,W436,N437,S438,N439,N440,L441,D442,S443,K444,V445,G446,G447,N448,Y449,N450,Y451,L452,Y453,R454,L455,G496,F497,Q498,P499,T500,N501,G502,V503,G504,Y505,Q506,P507,Y508,R509,V510,V511,V512,L513 | Ab145 | 10.1126/scitranslmed.abf1906 |
| Conformational | C136,N137,D138,P139,F140,L141,G142,V143,T307,V308,E309,K310,G311,I312,Y313,Q314,T315,S316,N317,F318,P621,V622,A623,I624,H625,A626,D627,Q628,L629,T630,P631,T632,W633,R634,V635,Y636 | Ab82 | 10.1126/scitranslmed.abf1906 |
| Conformational | I980,L981,S982,R983,L984,D985,K986,V987,E988,A989,E990,V991,Q992,I993,D994,R995,L996,I997,T998,G999,R1000,L1001,Q1002,S1003,L1004,Q1005,T1006,I1179,Q1180,K1181,E1182,I1183,D1184,R1185,L1186 | Ab127 | 10.1126/scitranslmed.abf1906 |
| Conformational | N960,T961,L962,V963,K964,Q965,L966,S967,S968,N969,F970,G971,A972,I973,S974,S975,V976,L977,N978,D979,I980,L981,S982,R983,L984,D985,K986,V987,E988,A989,E990,V991,Q992,I993,D994,R995,L996,I997,T998,G999,R1000,L1001,Q1002,S1003,L1004,Q1005,T1006,Y1007 | Ab164 | 10.1126/scitranslmed.abf1906 |
| Conformational | K417,I418,A419,D420,Y421,V433,I434,A435,W436,N437,S438,N439,N440,L441,D442,S443,K444,S459,N460,L461,K462,P463,F464,E465,R466,D467,I468,S469,T470,E471,I472,Y473,Q474,A475,G476,S477,T478,P479,C480,N481,G482,V483,E484,G485,F486,N487,C488,Y489,F490,P491,L492,Q493,S494,Y495,G496,F497,Q498,P499,T500,N501,G502,V503,G504,Y505,Q506,P507,Y508,R509,V510,V511,V512,L513 | Ly-CoV488 (Ab133) | 10.1126/scitranslmed.abf1906 |
| Conformational | D467,I468,S469,T470,E471,I472,Y473,Q474,A475,G476,S477,T478,P479,C480,N481,G482,V483,E484,G485,F486,N487,C488,Y489,F490,G496,F497,Q498,P499,T500,N501,G502,V503,G504,Y505,Q506,P507,Y508,R509,V510,V511,V512,L513 | Ly-CoV481 (Ab128) | 10.1126/scitranslmed.abf1906 |

| | | | |
|---|---|---|---|
| Conformational | V433,I434,A435,W436,N437,S438,N439,N440,L441,D442,S443,K444,S459,N460,L461,K462,P463,F464,E465,R466,D467,I468,S469,T470,E471,I472,Y473,Q474,A475,G476,S477,T478,P479,C480,N481,G482,V483,E484,G485,F486,N487,C488,Y489,F490,P491,L492,Q493,S494,Y495, | Ly-CoV555 (Ab169) | 10.1126/scitranslmed.abf1906 |
| Linear | 487-498;553-684;764-829;884-895;1148-1159;1256-1273 | N/A | 10.1038/s41423-020-00523-5 |
| Linear | 209-226;553-570;769-786;809-826 | N/A | 10.1016/j.ebiom.2020.102911 |
| Linear | 562-579;818-835 | N/A | 10.1038/s41467-020-16638-2 |
| Linear | 554-593;654-673;806-825;1146-1165 | N/A | 10.1080/22221751.2020.1815591 |
| Linear | 556-570;675-689;721-733 | N/A | 10.1101/2020.08.27.267716 |
| Linear | 553-564;655-672;787-798;811-822;1123-1134;1147-1158 | N/A | 10.1371/journal.pone.0238089 |
| Linear | 25-36;451-474;523-685;770-829;1148-1159;1256-1273 | N/A | 10.2139/ssrn.3671941 |
| Linear | 21-45;221-245;261-285;330-349;375-394;450-469;480-499;522-586;602-646;902-926 | N/A | 10.1038/s41422-020-0366-x |
| Linear | 551-570;766-785;811-830;1144-1163 | N/A | 10.1126/science.abd4250 |
| Linear | 421-434;742-759 | N/A | 10.3390/microorganisms8121993 |
| Linear | 79-93 | N/A | 10.3390/vaccines9010035 |
| Linear | 560-572;819-824;1150-1156 | N/A | 10.1016/j.xcrm.2020.100189 |
| Linear | 556-570 | N/A | 10.1016/j.celrep.2020.108666 |
| Linear | 397-403;557-567;661-671;789-799;813-823;1145-1159;1259-1271 | N/A | 10.1002/eji.202049101 |
| Linear | 541-579;801-839;1141-1179;1121-1159 | N/A | 10.1016/j.celrep.2021.109164 |
| Linear | 21-35;176-190;451-465;551-565;671-685;811-825;881-895;1146-1160;1166-1180;1216-1230 | N/A | 10.1172/jci.insight.148855. |

| | | | |
|---|---|---|---|
| Linear | 1031-1045 | N/A | 10.1186/s12866-021-02241-y |
