## Supplementary material for "Computation of Antigenicity Predicts SARS-CoV-2 Vaccine Breakthrough Variants": Table S2

**Table S2. Observed neutralisation titres of SARS-CoV-2 variants**

| Serum | VirusRef | VirusVar | WHO | Type | log2FC | Mean | N | Source | DOI |
|---|---|---|---|---|---|---|---|---|---|
| SinoVac | A | B.1.1.7 | Alpha | Pseudovirus ID50 | 1.02 | 2.03 | 75 | Acevedo ML, et al | 10.1101/2021.06.28.21259673 |
| SinoVac | A | P.1 | Gamma | Pseudovirus ID50 | 1.22 | 2.33 | 75 | Acevedo ML, et al | 10.1101/2021.06.28.21259673 |
| SinoVac | A | C.37 | Lambda | Pseudovirus ID50 | 1.61 | 3.05 | 75 | Acevedo ML, et al | 10.1101/2021.06.28.21259673 |
| SinoVac | A | B.1(D614G) | Non-VOC | Pseudovirus ID50 | 0.45 | 1.37 | 75 | Acevedo ML, et al | 10.1101/2021.06.28.21259673 |
| BioNTech/ Mod | A | B.1.1.7(E484K) | Alpha | Live Virus FRNT50 | 1.93 | 3.80 | 30 | Carreno, et al | 10.1101/2021.07.21.21260961 |
| BioNTech/ Mod | A | B.1.351 | Beta | Live Virus FRNT50 | 2.07 | 4.20 | 30 | Carreno, et al | 10.1101/2021.07.21.21260961 |
| BioNTech/Mod | A | B.1.617.2 | Delta | Live Virus FRNT50 | 1.58 | 3.00 | 30 | Carreno, et al | 10.1101/2021.07.21.21260961 |
| BioNTech/Mod | A | B.1.526 | Iota | Live Virus FRNT50 | 0.49 | 1.40 | 30 | Carreno, et al | 10.1101/2021.07.21.21260961 |
| BioNTech/Mod | A | B.1.526.2 | Iota | Live Virus FRNT50 | 1.20 | 2.30 | 30 | Carreno, et al | 10.1101/2021.07.21.21260961 |
| BioNTech/Mod | A | B.1.526(E484K) | Iota | Live Virus FRNT50 | 0.85 | 1.80 | 30 | Carreno, et al | 10.1101/2021.07.21.21260961 |
| BioNTech/Mod | A | C.37 | Lambda | Live Virus FRNT50 | 2.20 | 4.60 | 30 | Carreno, et al | 10.1101/2021.07.21.21260961 |
| Patient | B.1.1.117 | B.1.351 | Beta | Live Virus FRNT50 | 3.07 | 8.37 | 14 | Cele, et al | 10.1038/s41586-021-03471-w |
| Patient | B.1.351 | B.1.1.117 | Beta | Live Virus FRNT50 | 2.05 | 4.14 | 6 | Cele, et al | 10.1038/s41586-021-03471-w |
| SinoVac | A | B.1.1.7 | Alpha | Pseudovirus NT50 | 0.59 | 1.51 | 76 | Chen Y, et al | 10.1016/S1473-3099(21)00287-5 |
| SinoVac | A | B.1.351 | Beta | Pseudovirus NT50 | 2.40 | 5.27 | 76 | Chen Y, et al | 10.1016/S1473-3099(21)00287-5 |
| SinoVac | A | B.1.429 | Epsilon | Pseudovirus NT50 | 0.32 | 1.25 | 76 | Chen Y, et al | 10.1016/S1473-3099(21)00287-5 |
| SinoVac | A | P.1 | Gamma | Pseudovirus NT50 | 1.97 | 3.92 | 76 | Chen Y, et al | 10.1016/S1473-3099(21)00287-5 |
| SinoVac | A | B.1.526 | Iota | Pseudovirus NT50 | 2.01 | 4.03 | 76 | Chen Y, et al | 10.1016/S1473-3099(21)00287-5 |
| SinoVac | A | B.1(D614G) | Non-VOC | Pseudovirus NT50 | 0.28 | 1.21 | 76 | Chen Y, et al | 10.1016/S1473-3099(21)00287-5 |
| Patient | B.1(D614G) | B.1.1.7 | Alpha | Pseudovirus EC50 | 0.43 | 1.35 | 19 | Chen, et al | 10.1038/s41591-021-01294-w |
| BioNTech | B.1(D614G) | B.1.1.7 | Alpha | Pseudovirus EC50 | 1.02 | 2.03 | 24 | Chen, et al | 10.1038/s41591-021-01294-w |
| Patient | B.1(D614G) | B.1.351 | Beta | Pseudovirus EC50 | 2.20 | 4.58 | 19 | Chen, et al | 10.1038/s41591-021-01294-w |
| BioNTech | B.1(D614G) | B.1.351 | Beta | Pseudovirus EC50 | 3.35 | 10.20 | 24 | Chen, et al | 10.1038/s41591-021-01294-w |
| Patient | B.1(D614G) | P.1 | Gamma | Pseudovirus EC50 | 1.33 | 2.52 | 10 | Chen, et al | 10.1038/s41591-021-01294-w |
| BioNTech | B.1(D614G) | P.1 | Gamma | Pseudovirus EC50 | 1.16 | 2.23 | 15 | Chen, et al | 10.1038/s41591-021-01294-w |
| Patient | B.1(D614G) | K417N/D614G | Non-VOC | Pseudovirus EC50 | -0.58 | 0.67 | 19 | Chen, et al | 10.1038/s41591-021-01294-w |
| BioNTech | B.1(D614G) | K417N/D614G | Non-VOC | Pseudovirus EC50 | -0.01 | 0.99 | 24 | Chen, et al | 10.1038/s41591-021-01294-w |
| Patient | B.1(D614G) | E484K/N501Y/D6 | Non-VOC | Pseudovirus EC50 | 2.36 | 5.12 | 19 | Chen, et al | 10.1038/s41591-021-01294-w |
| BioNTech | B.1(D614G) | E484K/N501Y/D6 | Non-VOC | Pseudovirus EC50 | 2.06 | 4.17 | 24 | Chen, et al | 10.1038/s41591-021-01294-w |
| Patient | B.1(D614G) | K417N/E484K/N5 | Non-VOC | Pseudovirus EC50 | 1.83 | 3.55 | 19 | Chen, et al | 10.1038/s41591-021-01294-w |
| Patient | A | B.1.617.1 | Kappa | Live Virus FRNT50 | 2.66 | 6.32 | 24 | Edara, et al | 10.1056/NEJMc2107799 |

| | | | | | | | | | |
|---|---|---|---|---|---|---|---|---|---|
| Moderna | A | B.1.617.1 | Kappa | Live Virus FRNT50 | 2.77 | 6.82 | 15 | Edara, et al | 10.1056/NEJMc2107799 |
| BioNTech | A | B.1.617.1 | Kappa | Live Virus FRNT50 | 2.83 | 7.12 | 10 | Edara, et al | 10.1056/NEJMc2107799 |
| BioNTech | A | B.1.1.7 | Alpha | Pseudovirus NT50 | 1.07 | 2.10 | 30 | Garcia-Beltran, et al | 10.1016/j.cell.2021.03.013 |
| Moderna | A | B.1.1.7 | Alpha | Pseudovirus NT50 | 1.20 | 2.30 | 35 | Garcia-Beltran, et al | 10.1016/j.cell.2021.03.013 |
| BioNTech | A | B.1.351 | Beta | Pseudovirus NT50 | 5.11 | 34.50 | 30 | Garcia-Beltran, et al | 10.1016/j.cell.2021.03.013 |
| Moderna | A | B.1.351 | Beta | Pseudovirus NT50 | 4.79 | 27.70 | 35 | Garcia-Beltran, et al | 10.1016/j.cell.2021.03.013 |
| BioNTech | A | B.1.429 | Epsilon | Pseudovirus NT50 | 1.00 | 2.00 | 30 | Garcia-Beltran, et al | 10.1016/j.cell.2021.03.013 |
| Moderna | A | B.1.429 | Epsilon | Pseudovirus NT50 | 1.00 | 2.00 | 35 | Garcia-Beltran, et al | 10.1016/j.cell.2021.03.013 |
| BioNTech | A | P.1 | Gamma | Pseudovirus NT50 | 2.74 | 6.70 | 30 | Garcia-Beltran, et al | 10.1016/j.cell.2021.03.013 |
| Moderna | A | P.1 | Gamma | Pseudovirus NT50 | 2.17 | 4.50 | 35 | Garcia-Beltran, et al | 10.1016/j.cell.2021.03.013 |
| BioNTech | A | B.1(D614G) | Non-VOC | Pseudovirus NT50 | 0.26 | 1.20 | 30 | Garcia-Beltran, et al | 10.1016/j.cell.2021.03.013 |
| Moderna | A | B.1(D614G) | Non-VOC | Pseudovirus NT50 | 0.26 | 1.20 | 35 | Garcia-Beltran, et al | 10.1016/j.cell.2021.03.013 |
| BioNTech | A | B.1.1.298 | Non-VOC | Pseudovirus NT50 | 0.49 | 1.40 | 30 | Garcia-Beltran, et al | 10.1016/j.cell.2021.03.013 |
| Moderna | A | B.1.1.298 | Non-VOC | Pseudovirus NT50 | 0.38 | 1.30 | 35 | Garcia-Beltran, et al | 10.1016/j.cell.2021.03.013 |
| BioNTech | A | WIV1-CoV | WIV1-CoV | Pseudovirus EC50 | 5.47 | 44.30 | 30 | Garcia-Beltran, et al | 10.1016/j.cell.2021.03.013 |
| Moderna | A | WIV1-CoV | WIV1-CoV | Pseudovirus EC50 | 4.73 | 26.50 | 35 | Garcia-Beltran, et al | 10.1016/j.cell.2021.03.013 |
| BioNTech | A | SARS-CoV | SARS | Pseudovirus EC50 | 5.45 | 43.80 | 30 | Garcia-Beltran, et al | 10.1016/j.cell.2021.03.013 |
| Moderna | A | SARS-CoV | SARS | Pseudovirus EC50 | 5.07 | 33.50 | 35 | Garcia-Beltran, et al | 10.1016/j.cell.2021.03.013 |
| Patient | A | B.1.351 | Beta | Live Virus NT50 | 2.52 | 5.75 | 15 | Hoffman, et al | 10.1016/j.celrep.2021.109415 |
| BioNTech | A | B.1.351 | Beta | Live Virus NT50 | 3.48 | 11.13 | 15 | Hoffman, et al | 10.1016/j.celrep.2021.109415 |
| Patient | A | B.1.617 | Kappa | Live Virus NT50 | 0.97 | 1.96 | 15 | Hoffman, et al | 10.1016/j.celrep.2021.109415 |
| BioNTech | A | B.1.617 | Kappa | Live Virus NT50 | 1.50 | 2.83 | 15 | Hoffman, et al | 10.1016/j.celrep.2021.109415 |
| BioNTech | A | B.1.617.2 | Delta | Live Virus FRNT50 | 0.50 | 1.41 | 20 | Liu J-y, et al | 10.1038/s41586-021-03693-y |
| BioNTech | A | B.1.617.2 | Delta | Live Virus FRNT50 | 0.55 | 1.46 | 20 | Liu J-y, et al | 10.1038/s41586-021-03693-y |
| BioNTech | A | B.1.617.1 | Kappa | Live Virus FRNT50 | 1.68 | 3.20 | 20 | Liu J-y, et al | 10.1038/s41586-021-03693-y |
| BioNTech | A | B.1.525 | Non-VOC | Live Virus FRNT50 | 0.65 | 1.57 | 20 | Liu J-y, et al | 10.1038/s41586-021-03693-y |
| BioNTech | A | B.1.618 | Non-VOC | Live Virus FRNT50 | 0.60 | 1.52 | 20 | Liu J-y, et al | 10.1038/s41586-021-03693-y |
| Patient | A | B.1.1.7 | Alpha | Live Virus FRNT50 | 1.56 | 2.94 | 34 | Liu, et al | 10.1016/j.cell.2021.06.020 |
| Patient | B.1.1.7 | A | Alpha | Live Virus FRNT50 | -0.88 | 0.54 | 18 | Liu, et al | 10.1016/j.cell.2021.06.020 |
| BioNTech | A | B.1.1.7 | Alpha | Live Virus FRNT50 | 1.71 | 3.28 | 25 | Liu, et al | 10.1016/j.cell.2021.06.020 |
| AstraZeneca | A | B.1.1.7 | Alpha | Live Virus FRNT50 | 1.22 | 2.34 | 25 | Liu, et al | 10.1016/j.cell.2021.06.020 |
| Patient | A | B.1.351 | Beta | Live Virus FRNT50 | 3.74 | 13.32 | 34 | Liu, et al | 10.1016/j.cell.2021.06.020 |
| Patient | B.1.351 | A | Beta | Live Virus FRNT50 | 0.93 | 1.91 | 14 | Liu, et al | 10.1016/j.cell.2021.06.020 |

| | | | | | | | | | |
|---|---|---|---|---|---|---|---|---|---|
| BioNTech | A | B.1.351 | Beta | Live Virus FRNT50 | 2.92 | 7.57 | 25 | Liu, et al | 10.1016/j.cell.2021.06.020 |
| AstraZeneca | A | B.1.351 | Beta | Live Virus FRNT50 | 3.17 | 9.00 | 25 | Liu, et al | 10.1016/j.cell.2021.06.020 |
| Patient | A | B.1.617.2 | Delta | Live Virus FRNT50 | 1.42 | 2.67 | 34 | Liu, et al | 10.1016/j.cell.2021.06.020 |
| BioNTech | A | B.1.617.2 | Delta | Live Virus FRNT50 | 1.32 | 2.50 | 25 | Liu, et al | 10.1016/j.cell.2021.06.020 |
| AstraZeneca | A | B.1.617.2 | Delta | Live Virus FRNT50 | 2.11 | 4.31 | 25 | Liu, et al | 10.1016/j.cell.2021.06.020 |
| Patient | A | P.1 | Gamma | Live Virus FRNT50 | 1.65 | 3.13 | 34 | Liu, et al | 10.1016/j.cell.2021.06.020 |
| Patient | P.1 | A | Gamma | Live Virus FRNT50 | 1.96 | 3.89 | 17 | Liu, et al | 10.1016/j.cell.2021.06.020 |
| BioNTech | A | P.1 | Gamma | Live Virus FRNT50 | 1.39 | 2.62 | 25 | Liu, et al | 10.1016/j.cell.2021.06.020 |
| AstraZeneca | A | P.1 | Gamma | Live Virus FRNT50 | 1.52 | 2.86 | 25 | Liu, et al | 10.1016/j.cell.2021.06.020 |
| Patient | A | B.1.617.1-C | Kappa | Pseudovirus FRNT50 | 0.58 | 1.49 | 34 | Liu, et al | 10.1016/j.cell.2021.06.020 |
| Patient | A | B.1.617.1-A | Kappa | Pseudovirus FRNT50 | 1.33 | 2.52 | 34 | Liu, et al | 10.1016/j.cell.2021.06.020 |
| Patient | A | B.1.617.1-B | Kappa | Pseudovirus FRNT50 | 1.97 | 3.91 | 34 | Liu, et al | 10.1016/j.cell.2021.06.020 |
| BioNTech | A | B.1.617.1-B | Kappa | Pseudovirus NT50 | 1.45 | 2.72 | 25 | Liu, et al | 10.1016/j.cell.2021.06.020 |
| AstraZeneca | A | B.1.617.1-B | Kappa | Pseudovirus NT50 | 1.39 | 2.63 | 25 | Liu, et al | 10.1016/j.cell.2021.06.020 |
| Patient | B.1.351 | P.1 | Non-VOC | Live Virus FRNT50 | 1.00 | 2.00 | 14 | Liu, et al | 10.1016/j.cell.2021.06.020 |
| Patient | P.1 | B.1.351 | Non-VOC | Live Virus FRNT50 | 1.16 | 2.24 | 17 | Liu, et al | 10.1016/j.cell.2021.06.020 |
| Patient | B.1.1.7 | B.1.617.1-B | Non-VOC | Pseudovirus NT50 | 2.09 | 4.27 | 34 | Liu, et al | 10.1016/j.cell.2021.06.020 |
| Patient | B.1.1.7 | B.1.351 | Non-VOC | Live Virus FRNT50 | 1.38 | 2.60 | 18 | Liu, et al | 10.1016/j.cell.2021.06.020 |
| Patient | B.1.351 | B.1.1.7 | Non-VOC | Live Virus FRNT50 | 1.72 | 3.30 | 14 | Liu, et al | 10.1016/j.cell.2021.06.020 |
| Patient | B.1.1.7 | P.1 | Non-VOC | Live Virus FRNT50 | 0.76 | 1.69 | 18 | Liu, et al | 10.1016/j.cell.2021.06.020 |
| Patient | P.1 | B.1.1.7 | Non-VOC | Live Virus FRNT50 | 2.03 | 4.09 | 17 | Liu, et al | 10.1016/j.cell.2021.06.020 |
| Patient | B.1.1.7 | B.1.617.2 | Non-VOC | Live Virus FRNT50 | 0.60 | 1.51 | 18 | Liu, et al | 10.1016/j.cell.2021.06.020 |
| Patient | B.1.351 | B.1.617.2 | Non-VOC | Live Virus FRNT50 | 3.53 | 11.54 | 14 | Liu, et al | 10.1016/j.cell.2021.06.020 |
| Patient | P.1 | B.1.617.2 | Non-VOC | Live Virus FRNT50 | 3.51 | 11.37 | 17 | Liu, et al | 10.1016/j.cell.2021.06.020 |
| AstraZeneca | B.1(D614G) | B.1.351 | Beta | Pseudovirus ID50 | 2.00 | 4.01 | 13 | Madhi SA, et al | 10.1056/NEJMoa2102214 |
| Patient | B.1.1.117 | B.1.351 | Beta | Pseudovirus ID50 | 2.19 | 4.58 | 6 | Madhi SA, et al | 10.1056/NEJMoa2102214 |
| AstraZeneca | B.1.1 | B.1.351 | Beta | Live Virus FRNT50 | 3.46 | 11.00 | 13 | Madhi SA, et al | 10.1056/NEJMoa2102214 |
| Patient | B.1.1.117 | B.1.351 | Beta | Live Virus FRNT50 | 2.95 | 7.75 | 6 | Madhi SA, et al | 10.1056/NEJMoa2102214 |
| AstraZeneca | B.1(D614G) | K417N/E484K/N5 | Non-VOC | Pseudovirus ID50 | 1.80 | 3.49 | 13 | Madhi SA, et al | 10.1056/NEJMoa2102214 |
| Patient | B.1.1.117 | K417N/E484K/N5 | Non-VOC | Pseudovirus ID50 | 1.72 | 3.30 | 6 | Madhi SA, et al | 10.1056/NEJMoa2102214 |
| Patient | B.1.351 v2 | B.1(D614G) | Non-VOC | Pseudovirus ID50 | 1.61 | 3.04 | 57 | Moyo-Gwete, et al | 10.1056/NEJMc2104192 |
| BioNTech | A | B.1.1.7 | Alpha | Pseudovirus NT50 | -0.30 | 0.81 | 15 | Schmidt F,et al | 10.1038/s41586-021-04005-0 |
| BioNTech | A | B.1.351.3 | Beta | Pseudovirus NT50 | 1.69 | 3.23 | 15 | Schmidt F,et al | 10.1038/s41586-021-04005-0 |

| | | | | | | | | | |
|---|---|---|---|---|---|---|---|---|---|
| BioNTech | A | B.1.617.2 | Delta | Pseudovirus NT50 | 2.25 | 4.76 | 15 | Schmidt F,et al | 10.1038/s41586-021-04005-0 |
| BioNTech | A | P.1 | Gamma | Pseudovirus NT50 | 0.79 | 1.72 | 15 | Schmidt F,et al | 10.1038/s41586-021-04005-0 |
| BioNTech | A | B.1.526 | Iota | Pseudovirus NT50 | 0.71 | 1.64 | 15 | Schmidt F,et al | 10.1038/s41586-021-04005-0 |
| BioNTech | A | WIV1-CoV | WIV1-CoV | Pseudovirus NT50 | 4.06 | 16.67 | 15 | Schmidt F,et al | 10.1038/s41586-021-04005-0 |
| BioNTech | A | SARS-CoV | SARS | Pseudovirus NT50 | 5.06 | 33.33 | 15 | Schmidt F,et al | 10.1038/s41586-021-04005-0 |
| Patient | B.1(D614G) | B.1.351 | Beta | Pseudovirus ID50 | 3.71 | 13.10 | 14 | Shen X-Y, et al | 10.1056/NEJMc2103740 |
| Moderna | B.1(D614G) | B.1.351 | Beta | Pseudovirus ID50 | 3.28 | 9.70 | 26 | Shen X-Y, et al | 10.1056/NEJMc2103740 |
| Novavax | B.1(D614G) | B.1.351 | Beta | Pseudovirus ID50 | 3.86 | 14.50 | 23 | Shen X-Y, et al | 10.1056/NEJMc2103740 |
| Patient | B.1(D614G) | B.1.429 | Epsilon | Pseudovirus ID50 | 1.63 | 3.10 | 14 | Shen X-Y, et al | 10.1056/NEJMc2103740 |
| Moderna | B.1(D614G) | B.1.429 | Epsilon | Pseudovirus ID50 | 1.00 | 2.00 | 26 | Shen X-Y, et al | 10.1056/NEJMc2103740 |
| Novavax | B.1(D614G) | B.1.429 | Epsilon | Pseudovirus ID50 | 1.32 | 2.50 | 23 | Shen X-Y, et al | 10.1056/NEJMc2103740 |
| Patient | A | B.1.1.7 | Alpha | Live Virus FRNT50 | 1.54 | 2.90 | 34 | Supasa, et al | 10.1016/j.cell.2021.02.033 |
| AstraZeneca | A | B.1.1.7 | Alpha | Live Virus FRNT50 | 1.32 | 2.50 | 15 | Supasa, et al | 10.1016/j.cell.2021.02.033 |
| AstraZeneca | A | B.1.1.7 | Alpha | Live Virus FRNT50 | 1.07 | 2.10 | 10 | Supasa, et al | 10.1016/j.cell.2021.02.033 |
| BioNTech | A | B.1.1.7 | Alpha | Live Virus FRNT50 | 1.72 | 3.30 | 25 | Supasa, et al | 10.1016/j.cell.2021.02.033 |
| Patient | B.1.1.7 | A | Alpha | Live Virus FRNT50 | -0.12 | 0.92 | 13 | Supasa, et al | 10.1016/j.cell.2021.02.033 |
| Patient | A | B.1.617.2 | Delta | Live Virus FRNT50 | 1.26 | 2.40 | 24 | Suthar MS, et al | 10.1056/NEJMc2107799 |
| Moderna | A | B.1.617.2 | Delta | Live Virus FRNT50 | 1.58 | 3.00 | 15 | Suthar MS, et al | 10.1056/NEJMc2107799 |
| BioNTech | A | B.1.617.2 | Delta | Live Virus FRNT50 | 1.72 | 3.30 | 10 | Suthar MS, et al | 10.1056/NEJMc2107799 |
| Patient | A | B.1.617.1 | Kappa | Live Virus FRNT50 | 2.70 | 6.50 | 24 | Suthar MS, et al | 10.1056/NEJMc2107799 |
| Moderna | A | B.1.617.1 | Kappa | Live Virus FRNT50 | 2.81 | 7.00 | 15 | Suthar MS, et al | 10.1056/NEJMc2107799 |
| BioNTech | A | B.1.617.1 | Kappa | Live Virus FRNT50 | 2.81 | 7.00 | 10 | Suthar MS, et al | 10.1056/NEJMc2107799 |
| Patient | B.1(D614G) | B.1.351 | Beta | Pseudovirus IC50 | 2.29 | 4.90 | 8 | Tada, et al | 10.1101/2021.07.02.450959 |
| BioNTech | B.1(D614G) | B.1.351 | Beta | Pseudovirus IC50 | 1.32 | 2.50 | 15 | Tada, et al | 10.1101/2021.07.02.450959 |
| Moderna | B.1(D614G) | B.1.351 | Beta | Pseudovirus IC50 | 2.00 | 4.00 | 6 | Tada, et al | 10.1101/2021.07.02.450959 |
| Patient | B.1(D614G) | C.37 | Lambda | Pseudovirus IC50 | 1.72 | 3.30 | 8 | Tada, et al | 10.1101/2021.07.02.450959 |
| BioNTech | B.1(D614G) | C.37 | Lambda | Pseudovirus IC50 | 1.58 | 3.00 | 15 | Tada, et al | 10.1101/2021.07.02.450959 |
| Moderna | B.1(D614G) | C.37 | Lambda | Pseudovirus IC50 | 1.20 | 2.30 | 6 | Tada, et al | 10.1101/2021.07.02.450959 |
| SARS Patient | SARS-CoV | B.1.1.7 | SARS | Pseudovirus NT50 | 3.62 | 12.33 | 10 | Tan C-W, et al | 10.1056/NEJMoa2108453 |
| Patient | B.1.1.7 | SARS-CoV | SARS | Pseudovirus NT50 | 4.47 | 22.16 | 10 | Tan C-W, et al | 10.1056/NEJMoa2108453 |
| BioNTech | B.1.1.7 | SARS-CoV | SARS | Pseudovirus NT50 | 4.02 | 16.18 | 10 | Tan C-W, et al | 10.1056/NEJMoa2108453 |
| BioNTech | A | B.1.1.7 | Alpha | Live Virus IC50 | 1.38 | 2.60 | 159 | Wall EC, et al | 10.1016/S0140-6736(21)01290-3 |
| BioNTech | A | B.1.351 | Beta | Live Virus IC50 | 2.29 | 4.90 | 159 | Wall EC, et al | 10.1016/S0140-6736(21)01290-3 |

| | | | | | | | | | |
|---|---|---|---|---|---|---|---|---|---|
| BioNTech | A | B.1.617.2 | Delta | Live Virus IC50 | 2.54 | 5.80 | 159 | Wall EC, et al | 10.1016/S0140-6736(21)01290-3 |
| BioNTech | A | B.1(D614G) | Non-VOC | Live Virus IC50 | 1.20 | 2.30 | 159 | Wall EC, et al | 10.1016/S0140-6736(21)01290-3 |
| Patient | A | B.1.1.7 | Alpha | Pseudovirus ID50 | 1.68 | 3.20 | 20 | Wang, et al | 10.1038/s41586-021-03398-2 |
| Moderna | A | B.1.1.7 | Alpha | Pseudovirus ID50 | 0.85 | 1.80 | 12 | Wang, et al | 10.1038/s41586-021-03398-2 |
| BioNTech | A | B.1.1.7 | Alpha | Pseudovirus ID50 | 1.00 | 2.00 | 10 | Wang, et al | 10.1038/s41586-021-03398-2 |
| Patient | A | B.1.351 | Beta | Pseudovirus ID50 | 4.46 | 22.00 | 20 | Wang, et al | 10.1038/s41586-021-03398-2 |
| Moderna | A | B.1.351 | Beta | Pseudovirus ID50 | 3.10 | 8.60 | 12 | Wang, et al | 10.1038/s41586-021-03398-2 |
| BioNTech | A | B.1.351 | Beta | Pseudovirus ID50 | 2.70 | 6.50 | 10 | Wang, et al | 10.1038/s41586-021-03398-2 |
| Patient | B.1(D614G) | B.1.351 | Beta | Pseudovirus ID50 | 2.51 | 5.69 | 44 | Wibmer, et al | 10.1038/s41591-021-01285-x |
| Patient | B.1(D614G) | K417N/E484K/N5 | Non-VOC | Pseudovirus ID50 | 2.14 | 4.42 | 44 | Wibmer, et al | 10.1038/s41591-021-01285-x |
| BioNTech | A | B.1.351 | Beta | Live Virus FRNT50 | 2.92 | 7.57 | 25 | Zhou, et al | 10.1016/j.cell.2021.02.037 |
| AstraZeneca | A | B.1.351 | Beta | Live Virus FRNT50 | 3.17 | 9.00 | 25 | Zhou, et al | 10.1016/j.cell.2021.02.037 |
| Patient | A | B.1.351 | Beta | Live Virus FRNT50 | 3.72 | 13.19 | 34 | Zhou, et al | 10.1016/j.cell.2021.02.037 |
| Patient | B.1.1.7 | B.1.351 | Beta | Live Virus FRNT50 | 1.65 | 3.13 | 14 | Zhou, et al | 10.1016/j.cell.2021.02.037 |
