## Supplementary material for "Computation of Antigenicity Predicts SARS-CoV-2 Vaccine Breakthrough Variants": Table S3

**Table S3. SARS-CoV-2 variants analysed in our study.** We gratefully acknowledge the following Authors from the Originating laboratories responsible for obtaining the specimens and the Submitting laboratories where genetic sequence data were generated and shared via the GISAID Initiative, on which this research is based.

| Lineage | Accession ID | Virus Name | Collection Date | Submitting Lab | Authors |
|---|---|---|---|---|---|
| A | EPI_ISL_529213 | Wuhan/IME-WH01/2019 | 2019/12/30 | Beijing Institute of Microbiology and Epidemiology | Fan et al |
| A.1 | EPI_ISL_2836011 | USA/un-Yale-5643/2020 | 2020/1/19 | Epidemiology of Microbial Diseases, Yale School of Public Health | Vogels et al |
| A.11 | EPI_ISL_420078 | Senegal/620/2020 | 2020/3/20 | Institut Pasteur de Dakar | Ndongo Dia et al |
| A.12 | EPI_ISL_512816 | SierraLeone/KGH-G-8625/2020 | 2020/4/10 | Kenema Government Hospital, Ministry of Health and Sanitation | Goba et al |
| A.15 | EPI_ISL_622356 | Denmark/DCGC-6742/2020 | 2020/4/20 | Albertsen lab, Department of Chemistry and Bioscience, Aalborg University, Denmark | Danish Covid-19 Genome Consortia et al |
| A.16 | EPI_ISL_480042 | Japan/PG-0372/2020 | 2020/2/15 | Pathogen Genomics Center, National Institute of Infectious Diseases | Tsuyoshi Sekizuka et al |
| A.17 | EPI_ISL_660334 | France/ARA-SC338/2020 | 2020/3/2 | CNR Virus des Infections Respiratoires - France SUD | Antonin Bal et al |
| A.18 | EPI_ISL_1662585 | Coted-Ivoire/BKE0714-bc14/2020 | 2020/7/24 | Project group Epidemology of Highly Pathogenic Microorganisms, Robert Koch-Institute | Chantal Akoua-Koffi et al |
| A.19 | EPI_ISL_614351 | Coted-Ivoire/BKE0041/2020 | 2020/6/4 | Project group Epidemology of Highly Pathogenic Microorganisms, Robert Koch-Institute | Chantal Akoua-Koffi et al |
| A.2 | EPI_ISL_496607 | Panama/328719/2020 | 2020/2/22 | Gorgas Memorial Laboratory of Health Studies | Danilo Franco et al |
| A.2.2 | EPI_ISL_483708 | Israel/n14268/2020 | 2020/3/17 | Israel Central Virology laboratory | Neta Zuckerman et al |
| A.2.3 | EPI_ISL_425752 | Scotland/CVR24/2020 | 2020/3/12 | COVID-19 Genomics UK (COG-UK) Consortium | Ana da Silva Filipe et al |
| A.2.4 | EPI_ISL_1225382 | Panama/GMI-PA352445/2020 | 2020/2/4 | Gorgas Memorial Laboratory of Health Studies | Yamilka Diaz et al |
| A.2.5 | EPI_ISL_1502840 | Panama/GMI-PA341357/2020 | 2020/4/15 | Gorgas Memorial Laboratory of Health Studies | Gonzalez Claudia et al |
| A.2.5 | EPI_ISL_2000585 | USA/RI_RKL_001_31745_AM614/2020 | 2020/4/25 | Rami Kantor lab | Josephine Darpolor et al |

| | | | | | |
|---|---|---|---|---|---|
| A.2.5.1 | EPI_ISL_1502839 | Panama/GMI-PA340863/2020 | 2020/4/14 | Gorgas Memorial Laboratory of Health Studies | Gonzalez Claudia et al |
| A.2.5.2 | EPI_ISL_1502836 | Panama/GMI-PA339770/2020 | 2020/4/12 | Gorgas Memorial Laboratory of Health Studies | Gonzalez Claudia et al |
| A.2.5.3 | EPI_ISL_1681051 | USA/FL-CDC-LC0040585/2021 | 2021/4/7 | Centers for Disease Control and Prevention Division of Viral Diseases, Pathogen Discovery | Dakota Howard et al |
| A.21 | EPI_ISL_487456 | Mali/M002675/2020 | 2020/4/10 | Bundeswehr Institut of Microbiology | Kouriba et al |
| A.22 | EPI_ISL_457938 | Oman/RESP-20-1976/2020 | 2020/3/15 | Oman-NIC | Samira Al-Maruqi et al |
| A.23 | EPI_ISL_738016 | Uganda/UG109/2020 | 2020/8/14 | MRC/UVRI & LSHTM Uganda Research Unit | Matthew Cotten et al |
| A.23.1 | EPI_ISL_5143058 | UnitedArabEmirates/BTC-4974/2020 | 2020/9/21 | BTC, Khalifa University | Habiba Al Safar et al |
| A.24 | EPI_ISL_475586 | USA/CA-CSMC110/2020 | 2020/3/22 | Cedars-Sinai Medical Center, Molecular Pathology Laboratory of Department of Pathology & Laboratory Medicine and Genomic Core | Wenjuan Zhang et al |
| A.25 | EPI_ISL_2602919 | Kenya/K10131/2020 | 2020/6/8 | KEMRI-Wellcome Trust Research Programme,Kilifi | Githinji G. et al |
| A.26 | EPI_ISL_539009 | England/LEED-2A89EF/2020 | 2020/3/31 | Wellcome Sanger Institute for the COVID-19 Genomics UK (COG-UK) consortium | Louissa Macfarlane-Smith et al |
| A.27 | EPI_ISL_757470 | Denmark/DCGC-22492/2020 | 2020/12/14 | Albertsen Lab, Department of Chemistry and Bioscience, Aalborg University, Denmark | Danish Covid-19 Genome Consortium et al |
| A.28 | EPI_ISL_812821 | Egypt/CCHE57357-A-48/2020 | 2020/7/28 | Genomics Program, Children Cancer Hospital | Hatem et al |
| A.29 | EPI_ISL_1731548 | Gambia/22178/2020 | 2020/8/29 | MRCG at LSHTM Genomics lab | Abdul Karim sesay et al |
| A.3 | EPI_ISL_438234 | USA/MD-HP00076/2020 | 2020/3/11 | Johns Hopkins Hospital Department of Pathology | Peter M. Thielen et al |
| A.4 | EPI_ISL_417514 | USA/WI-UW-22/2020 | 2020/3/13 | University of Wisconsin-Madison AIDS Vaccine Research Laboratories | Gage Moreno et al |
| A.5 | EPI_ISL_529992 | Spain/MD-H12-61/2020 | 2020/3/1 | Hospital Universitario 12 de Octubre | Sara González et al |
| A.6 | EPI_ISL_437612 | Thailand/SI203285-NT/2020 | 2020/1/23 | Faculty of Medicine | Rodpan et al |
| A.7 | EPI_ISL_455749 | India/OR-RMRC103/2020 | 2020/5/6 | Immunogenomics lab, Institute of Life Sciences, Bhubaneswar | Sunil Raghav et al |

| | | | | | |
|---|---|---|---|---|---|
| A.9 | EPI_ISL_775626 | Germany/HH-hpi-p2134/2020 | 2020/3/21 | Heinrich Pette Institute, Leibniz Institute for Experimental Virology | Alexis Robitaille et al |
| AA.1 | EPI_ISL_540903 | Wales/PHWC-16A540/2020 | 2020/9/6 | COVID-19 Genomics UK (COG-UK) Consortium | Catherine Moore et al |
| AA.2 | EPI_ISL_680608 | Wales/PHWC-48D32C/2020 | 2020/11/14 | COVID-19 Genomics UK (COG-UK) Consortium | Catherine Moore et al |
| AA.3 | EPI_ISL_628267 | Wales/PHWC-485C38/2020 | 2020/10/21 | COVID-19 Genomics UK (COG-UK) Consortium | Catherine Moore et al |
| AA.4 | EPI_ISL_634761 | Wales/ALDP-AA308D/2020 | 2020/10/22 | Wellcome Sanger Institute for the COVID-19 Genomics UK (COG-UK) consortium | Jacquelyn Wynn et al |
| AA.5 | EPI_ISL_742926 | Wales/PHWC-49DECE/2020 | 2020/12/5 | COVID-19 Genomics UK (COG-UK) Consortium | Catherine Moore et al |
| AA.6 | EPI_ISL_637606 | Wales/PHWC-48738C/2020 | 2020/10/13 | COVID-19 Genomics UK (COG-UK) Consortium | Catherine Moore et al |
| AA.7 | EPI_ISL_666232 | Wales/PHWC-48B3D3/2020 | 2020/11/4 | COVID-19 Genomics UK (COG-UK) Consortium | Catherine Moore et al |
| AA.8 | EPI_ISL_814416 | Wales/PHWC-4A6F92/2020 | 2020/11/25 | COVID-19 Genomics UK (COG-UK) Consortium | Catherine Moore et al |
| AB.1 | EPI_ISL_748906 | Denmark/DCGC-19364/2020 | 2020/12/7 | Albertsen Lab, Department of Chemistry and Bioscience, Aalborg University, Denmark | Danish Covid-19 Genome Consortium et al |
| AC.1 | EPI_ISL_629268 | England/MILK-ACC112/2020 | 2020/10/21 | Wellcome Sanger Institute for the COVID-19 Genomics UK (COG-UK) consortium | The Lighthouse Lab in Milton Keynes et al |
| AD.1 | EPI_ISL_669913 | Denmark/DCGC-10343/2020 | 2020/11/2 | Albertsen Lab, Department of Chemistry and Bioscience, Aalborg University, Denmark | Danish Covid-19 Genome Consortium et al |
| AD.2 | EPI_ISL_573313 | England/NORT-286167/2020 | 2020/7/21 | COVID-19 Genomics UK (COG-UK) Consortium | Darren L Smith et al |
| AD.2.1 | EPI_ISL_933914 | Lithuania/ID2614/2020 | 2020/12/4 | Vilnius university hospital Santaros Klinikos, Center of Laboratory Medicine | Ingrida Olendraite et al |
| AE.1 | EPI_ISL_2982533 | DRC/7325/2020 | 2020/5/2 | Pathogen Sequencing Lab, National Institute for Biomedical Research (INRB) | Placide Mbala-Kingebeni et al |
| AE.2 | EPI_ISL_486887 | Bahrain/02/2020 | 2020/6/23 | National Influenza Center, Bahrain | Zaed et al |
| AE.3 | EPI_ISL_977373 | Zambia/ZMB-24445/2020 | 2020/6/4 | UNZAVET and PATH | Mulenga Mwenda-Chimfwembe et al |
| AE.4 | EPI_ISL_486889 | Bahrain/04/2020 | 2020/6/23 | National Influenza Center, Bahrain | Altaif et al |

| | | | | | |
|---|---|---|---|---|---|
| AE.5 | EPI_ISL_977392 | Zambia/ZMB-31145/2020 | 2020/6/28 | UNZAVET and PATH | Mulenga Mwenda-Chimfwembe et al |
| AE.6 | EPI_ISL_682307 | Bahrain/920453170/2020 | 2020/10/11 | Communicable Disease Laboratory, Public Health Directorate | Alwasti et al |
| AE.7 | EPI_ISL_681301 | Bahrain/920396988_S5_L001/2020 | 2020/9/11 | Communicable Disease Laboratory, Public Health Directorate | Alwasti et al |
| AE.8 | EPI_ISL_2163060 | Canada/ABPHL-02509/2020 | 2020/8/5 | Public Health Agency of Canada (PHAC) National Microbiology Laboratory | Buss et al |
| AF.1 | EPI_ISL_1496474 | Switzerland/BL-ETHZ-330007/2020 | 2020/10/16 | Department of Biosystems Science and Engineering, ETH Zürich | Christian Beisel et al |
| AG.1 | EPI_ISL_2886646 | Slovenia/91111/2020 | 2020/9/11 | Institute of Microbiology and Immunology, Faculty of Medicine, University of Ljubljana | Alen Suljič et al |
| AH.1 | EPI_ISL_1059585 | Switzerland/BE-ETHZ-481450/2021 | 2021/1/5 | Department of Biosystems Science and Engineering, ETH Zürich | Chaoran Chen et al |
| AH.2 | EPI_ISL_569103 | France/PAC-IHU-1270/2020 | 2020/8/18 | MEPHI, Aix Marseille University | Anthony LEVASSEUR et al |
| AH.3 | EPI_ISL_759771 | Finland/23LS1E6/2020 | 2020/10/23 | Department of Virology, Faculty of Medicine, University of Helsinki, Helsinki, Finland | Teemu Smura et al |
| AJ.1 | EPI_ISL_732022 | Portugal/PT1795/2020 | 2020/11/8 | Instituto Nacional de Saude (INSA) | Borges et al |
| AK.1 | EPI_ISL_590451 | England/QEUH-9B5511/2020 | 2020/9/3 | Wellcome Sanger Institute for the COVID-19 Genomics UK (COG-UK) consortium | Harper VanSteenhouse et al |
| AK.2 | EPI_ISL_631381 | Germany/NW-HHU-248/2020 | 2020/9/19 | Center of Medical Microbiology, Virology, and Hospital Hygiene, University of Duesseldorf | Maximilian Damagnez et al |
| AL.1 | EPI_ISL_1202078 | USA/TX-HMH-MCoV-12043/2020 | 2020/9/18 | Houston Methodist Hospital | S. Wesley Long et al |
| AM.1 | EPI_ISL_731937 | Portugal/PT1799/2020 | 2020/11/9 | Instituto Nacional de Saude (INSA) | Borges et al |
| AM.3 | EPI_ISL_665814 | England/LOND-1250DAB/2020 | 2020/11/10 | COVID-19 Genomics UK (COG-UK) Consortium | Judith Heaney et al |
| AM.4 | EPI_ISL_1739607 | Canada/ON-SC2233/2020 | 2020/11/5 | McMaster University | Allison McGeer et al |
| AN.1 | EPI_ISL_548256 | Sweden/20-52355/2020 | 2020/7/7 | The Public Health Agency of Sweden | Anna-Malin Linde et al |
| AQ.1 | EPI_ISL_757317 | Finland/17MR7cE12/2020 | 2020/11/17 | Department of Virology, Faculty of Medicine, University of Helsinki, Helsinki, Finland | Teemu Smura et al |

| | | | | | |
|---|---|---|---|---|---|
| AQ.2 | EPI_ISL_616717 | Denmark/DCGC-2956/2020 | 2020/8/24 | Albertsen lab, Department of Chemistry and Bioscience, Aalborg University, Denmark | Danish Covid-19 Genome Consortia et al |
| AS.1 | EPI_ISL_1403144 | Canada/Qc-HSE-5524134965/2020 | 2020/10/6 | Laboratoire de santé publique du Québec | Sandrine Moreira et al |
| AS.2 | EPI_ISL_2900306 | England/ALDP-9E809C/2020 | 2020/9/24 | COVID-19 Genomics UK (COG-UK) Consortium | PHE Covid Sequencing Team et al |
| AU.1 | EPI_ISL_1424680 | PapuaNewGuinea/PNG293/2021 | 2021/1/1 | Melbourne Diagnostic Unit Public Health Laboratory (MDU-PHL) | Palou et al |
| AU.2 | EPI_ISL_2379985 | Malaysia/UNIMAS-B187/2021 | 2021/1/3 | Institute of Health and Community Medicine | Chan Chia Jui et al |
| AU.3 | EPI_ISL_1118214 | Indonesia/JK-NIHRD-MI-21-01773/2021 | 2021/2/9 | National Institute of Health Research and Development | Hana Apsari Pawestri et al |
| AV.1 | EPI_ISL_1386007 | Turkey/HSGM-2150/2021 | 2021/1/27 | Ministry of Health Turkey | Fatma Bayrakdar et al |
| AW.1 | EPI_ISL_974269 | Canada/BC-BCCDC-5890/2020 | 2020/10/12 | BCCDC Public Health Laboratory | Prystajecky Natalie et al |
| AY.1 | EPI_ISL_1838677 | India/AP-CCMB-BN992/2021 | 2021/4/7 | CSIR-Centre for Cellular and Molecular Biology-INSACOG | Payel Mukherjee et al |
| AY.100 | EPI_ISL_2937904 | Mexico/AGU-InDRE_FB18599_S4467/2020 | 2020/9/22 | Instituto de Diagnostico y Referencia Epidemiologicos (INDRE) | Claudia Wong-Arambula et al |
| AY.102 | EPI_ISL_4077684 | Switzerland/BE-IFIK-210906_os-89/2021 | 2021/1/8 | Institute for Infectious Diseases | Stefan Neuenschwander et al |
| AY.102 | EPI_ISL_2716761 | Italy/LAZ-IFO-05103208/2021 | 2021/5/7 | IRCCS Regina Elena National Cancer Institute | Ludovica Ciuffreda et al |
| AY.102 | EPI_ISL_4415500 | India/MH-IISER-Pune-IP01472/2021 | 2021/1/9 | IISER Pune-INSACOG | Krishanpal Karmodiya et al |
| AY.102 | EPI_ISL_4415528 | India/MH-IISER-Pune-IP-01014/2021 | 2021/2/9 | IISER Pune-INSACOG | Krishanpal Karmodiya et al |
| AY.102 | EPI_ISL_4101601 | Turkey/HSGM-E2913/2021 | 2021/3/9 | Ministry of Health Turkey | Fatma Bayrakdar et al |
| AY.102 | EPI_ISL_4415524 | India/MH-IISER-Pune-IP01463/2021 | 2021/1/9 | IISER Pune-INSACOG | Krishanpal Karmodiya et al |
| AY.103 | EPI_ISL_5886224 | India/PY-nCOVb30339_S752_R1_001/2021 | 2021/1/1 | inStem NCBS – INSACOG | Uma Ramakrishnan et al |
| AY.103 | EPI_ISL_4884869 | Italy/SIC_CQRC_21066374/2021 | 2021/3/16 | CQRC_QUALITY CONTROL CHEMICAL BIOLOGICAL RISK_AOOR Villa Sofia Cervello Palermo | Di Gaudio F. et al |
| AY.106 | EPI_ISL_4101782 | Turkey/HSGM-E3132/2021 | 2021/3/9 | Ministry of Health Turkey | Fatma Bayrakdar et al |

| | | | | | |
|---|---|---|---|---|---|
| AY.107 | EPI_ISL_2379462 | India/TN-C_100_S15_R1_001/2021 | 2021/2/5 | inStem NCBS - INSACOG | Uma Ramakrishnan Dasaradhi Palakodeti Aswin SaiNarain et al |
| AY.11 | EPI_ISL_1790701 | England/MILK-1523694/2021 | 2021/4/20 | Wellcome Sanger Institute for the COVID-19 Genomics UK (COG-UK) Consortium | The Lighthouse Lab in Milton Keynes et al |
| AY.110 | EPI_ISL_1907995 | USA/MI-CDC-STM-000056311/2021 | 2021/4/14 | Centers for Disease Control and Prevention Division of Viral Diseases, Pathogen Discovery | Dakota Howard et al |
| AY.111 | EPI_ISL_1652897 | England/RAND-14FD37F/2021 | 2021/4/12 | Wellcome Sanger Institute for the COVID-19 Genomics UK (COG-UK) Consortium | Randox Laboratories et al |
| AY.114 | EPI_ISL_2774897 | India/TG-CCMB-CIA4758/2021 | 2021/4/10 | CSIR-Centre for Cellular and Molecular Biology-INSACOG | Lamuk Zaveri et al |
| AY.115 | EPI_ISL_5502108 | Botswana/R36B65_BHP_000640473/2021 | 2021/5/31 | Botswana Harvard HIV Reference Laboratory | Sikhulile Moyo et al |
| AY.116 | EPI_ISL_6347359 | Norway/24584/2021 | 2021/1/21 | Norwegian Institute of Public Health, Department of Virology | Kathrine Stene-Johansen et al |
| AY.116.1 | EPI_ISL_2545604 | Belgium/UZA-UA-CV2184230236/2021 | 2021/6/7 | Labo Klinische Biologie, UZA | Jasmine Coppens et al |
| AY.118 | EPI_ISL_4484419 | Slovakia/UVZ_PL36_B6_17813/2021 | 2021/2/9 | Laboratory of Genomics and Bioinformatics, Comenius University Science Park | Tatiana Sedláčková et al |
| AY.120.1 | EPI_ISL_5348704 | USA/ID-IBL-760105/2021 | 2021/2/23 | Idaho Bureau of Laboratories | R. Beukelman et al |
| AY.120.2 | EPI_ISL_2227435 | USA/MD-MDH-2355/2021 | 2021/5/6 | MD PHL | Maryland Department of Health Laboratories Administration et al |
| AY.120.2.1 | EPI_ISL_2762159 | Germany/BW-RKI-I-183727/2021 | 2021/6/6 | Robert Koch Institute | ? |
| AY.122 | EPI_ISL_1663563 | India/ILSGS00993/2021 | 2021/3/25 | Institute of Life Sciences - INSACOG | Sunil K. Raghav et al |
| AY.123 | EPI_ISL_5414903 | Belgium/rega-12446/2021 | 2021/4/25 | KU Leuven, Rega Institute, Clinical and Epidemiological Virology | Tony Wawina-Bokalanga et al |
| AY.125 | EPI_ISL_5508422 | Israel/CVL-22316/2021 | 2021/1/8 | Israel Central Virology laboratory | Neta Zuckerman et al |
| AY.125 | EPI_ISL_4271105 | Spain/CT-LabRefCat-751420/2021 | 2021/1/29 | Laboratori de Referencia de Catalunya | Ramirez A. et al |
| AY.13 | EPI_ISL_3215995 | Mexico/BCN-SEARCH-104131/2021 | 2021/2/4 | Andersen lab at Scripps Research | SEARCH Alliance San Diego with Jonathan Gonzalez Garcia et al |

| | | | | | |
|---|---|---|---|---|---|
| AY.14 | EPI_ISL_5778142 | USA/KY-LMDC-20180452/2021 | 2021/3/30 | University of Louisville Sequencing Technology Center | Elizabeth Hudson et al |
| AY.16 | EPI_ISL_4415438 | India/MH-IISER-Pune-IP-01040/2021 | 2021/2/9 | IISER Pune-INSACOG | Krishanpal Karmodiya et al |
| AY.18 | EPI_ISL_3484616 | Canada/BC-BCCDC-93175/2021 | 2021/5/11 | BCCDC Public Health Laboratory | Prystajecky Natalie et al |
| AY.19 | EPI_ISL_2897754 | India/MH-NS2079/2021 | 2021/4/19 | CDFD | Murali Bashyam et al |
| AY.2 | EPI_ISL_3275127 | USA/ID-UNM-IBL720784/2021 | 2021/3/31 | Center for Global Health, University of New Mexico Health Sciences Center | Daryl Domman et al |
| AY.20 | EPI_ISL_5703214 | USA/TN-SPHL-0539/2021 | 2021/1/14 | TN DOH Lab Services | Karen Beasley-Maynard et al |
| AY.20 | EPI_ISL_5639223 | USA/CA-CDPH-500008629/2021 | 2021/8/2 | California Department of Public Health | Emily Smith on behalf of CDPH-COVIDNet et al |
| AY.22 | EPI_ISL_2536066 | Portugal/PT8419/2021 | 2021/5/28 | Instituto Nacional de Saude (INSA) | Borges et al |
| AY.23 | EPI_ISL_5395771 | Indonesia/JA-GS-EIJK-RSRM-0172/2020 | 2020/7/29 | Eijkman Research Center for Molecular Biology, National Research and Innovation Agency; Raden Mattaher Regional General Hospital (RSUD) | Jessica R Saragih et al |
| AY.23.1 | EPI_ISL_3427022 | Singapore/3563/2021 | 2021/8/7 | National Public Health Laboratory, National Centre for Infectious Diseases | Katherine Ching et al |
| AY.24 | EPI_ISL_5395769 | Indonesia/JA-GS-EIJK-RSRM-0170/2020 | 2020/7/29 | Eijkman Research Center for Molecular Biology, National Research and Innovation Agency; Raden Mattaher Regional General Hospital (RSUD) | Willy Agustine et al |
| AY.25 | EPI_ISL_4689452 | USA/WV-WVU-WV121217/2020 | 2020/7/21 | WVU and Marshall University Combined Genomics Core Facilities | James Denvir et al |
| AY.26 | EPI_ISL_4484170 | Slovakia/UVZ_PL36_C5_17806/2021 | 2021/1/9 | Laboratory of Genomics and Bioinformatics, Comenius University Science Park | Tatiana Sedláčková et al |
| AY.28 | EPI_ISL_4469231 | SriLanka/idea_uoc_00065/2021 | 2021/2/16 | IDEA Laboratory | Inoka C. Perera et al |
| AY.29.1 | EPI_ISL_4723864 | Japan/PG-119190/2021 | 2021/7/2 | Pathogen Genomics Center, National Institute of Infectious Diseases | Tsuyoshi Sekizuka et al |
| AY.3 | EPI_ISL_4558284 | USA/WY-WYPHL-21070660/2020 | 2020/9/2 | Wyoming Public Health Laboratory | Jim Mildenberger et al |
| AY.3.1 | EPI_ISL_3355089 | USA/MS-CDC-2-4524366/2021 | 2021/6/1 | Centers for Disease Control and Prevention Division of Viral Diseases, Pathogen Discovery | Mili Sheth et al |

| | | | | | |
|---|---|---|---|---|---|
| AY.30 | EPI_ISL_5030213 | Thailand/DMSc-02114/2021 | 2021/4/6 | Division of Genomic Medicine and Innovation support,Department of Medical Sciences, Ministry of Public Health, Thailand | Surakameth Mahasirimongkol et al |
| AY.31 | EPI_ISL_3026386 | SouthKorea/KDCA6298/2021 | 2021/6/30 | Division of Emerging Infectious Diseases, Bureau of Infectious Diseases Diagnosis Control, Korea Disease Control and Prevention Agency | Jeong-Ah Kim et al |
| AY.33 | EPI_ISL_4101637 | Turkey/HSGM-E2953/2021 | 2021/3/9 | Ministry of Health Turkey | Fatma Bayrakdar et al |
| AY.34 | EPI_ISL_3154890 | Senegal/SN-IR2-1019/2021 | 2021/6/1 | IRESSEF | Souleymane MBOUP et al |
| AY.34 | EPI_ISL_4460844 | India/DL-NCDC-4509323/2021 | 2021/7/13 | NCDC Delhi, Biotechnology Division - INSACOG | Kalaiarasan Ponnusamy et al |
| AY.37 | EPI_ISL_4232132 | Liberia/LIB-0233/2021 | 2021/3/9 | Center for Infection and Immunity, Columbia University | Bode Shobayo et al |
| AY.38 | EPI_ISL_3275133 | USA/ID-UNM-IBL720559/2021 | 2021/3/27 | Center for Global Health, University of New Mexico Health Sciences Center | Daryl Domman et al |
| AY.39 | EPI_ISL_5703453 | USA/TN-SPHL-0979/2021 | 2021/1/14 | TN DOH Lab Services | Karen Beasley-Maynard et al |
| AY.39 | EPI_ISL_4474077 | India/JK-NCDC-3909159/2021 | 2021/6/1 | NCDC Delhi, Biotechnology Division - INSACOG | Mahesh S Dhar et al |
| AY.39.1 | EPI_ISL_2921622 | Australia/NSW-R0509/2021 | 2021/7/2 | Microbiology RPAH | Foster et al |
| AY.39.1.1 | EPI_ISL_3417445 | Australia/NSW-RPAH-0968/2021 | 2021/8/4 | Microbiology RPAH | Foster et al |
| AY.39.2 | EPI_ISL_3112511 | USA/NY-CDC-LC0095673/2021 | 2021/7/6 | Centers for Disease Control and Prevention Division of Viral Diseases, Pathogen Discovery | Dakota Howard et al |
| AY.4 | EPI_ISL_2723338 | England/NORT-1BD830A/2021 | 2021/6/21 | COVID-19 Genomics UK (COG-UK) Consortium | Darren L Smith et al |
| AY.4 | EPI_ISL_2925424 | England/ALDP-188D2DD/2021 | 2021/7/1 | Wellcome Sanger Institute for the COVID-19 Genomics UK (COG-UK) Consortium | Jacquelyn Wynn et al |
| AY.4.2 | EPI_ISL_1030354 | USA/CA-CZB-23359/2021 | 2021/1/14 | Chan-Zuckerberg Biohub | CZB Cliahub Consortium et al |
| AY.4.2.1 | EPI_ISL_2867411 | England/BRBR-1827F0B/2021 | 2021/6/29 | Wellcome Sanger Institute for the COVID-19 Genomics UK (COG-UK) Consortium | Berkshire et al |
| AY.4.5 | EPI_ISL_2703020 | England/MILK-174BF20/2021 | 2021/6/18 | Wellcome Sanger Institute for the COVID-19 Genomics UK (COG-UK) Consortium | The Lighthouse Lab in Milton Keynes et al |

| | | | | | |
|---|---|---|---|---|---|
| AY.41 | EPI_ISL_2897698 | India/MH-NS2016/2021 | 2021/4/9 | CDFD | Murali Bashyam et al |
| AY.43 | EPI_ISL_4533969 | India/GJ-INSACOG-GBRC2227/2021 | 2021/1/9 | Gujarat Biotechnology Research Centre | Bhadreshsinh Gohil et al |
| AY.43 | EPI_ISL_2361845 | India/TN-ICMR-NIRT-INSAGOG-5593/2021 | 2021/5/3 | NIV Influenza | Potdar et al |
| AY.43.1 | EPI_ISL_3246106 | Brazil/SP-IB_121137/2021 | 2021/7/19 | Instituto Butantan | Dimas Tadeu Covas et al |
| AY.43.2 | EPI_ISL_3032334 | Brazil/SP-IB_119544/2021 | 2021/7/5 | Instituto Butantan | Dimas Tadeu Covas et al |
| AY.45 | EPI_ISL_5416409 | SouthAfrica/NICD-N15251/2021 | 2021/1/19 | National Institute for Communicable Diseases of the National Health Laboratory Service | Amoako DG et al |
| AY.46 | EPI_ISL_4483949 | Slovakia/UVZ_PL36_F7_17921/2021 | 2021/1/9 | Laboratory of Genomics and Bioinformatics, Comenius University Science Park | Tatiana Sedláčková et al |
| AY.46.1 | EPI_ISL_4451151 | India/PB-NCDC-3509346/2021 | 2021/5/18 | NCDC Delhi, Biotechnology Division - INSACOG | Robin Marwal et al |
| AY.46.2 | EPI_ISL_4101849 | Turkey/HSGM-E3209/2021 | 2021/3/9 | Ministry of Health Turkey | Fatma Bayrakdar et al |
| AY.46.3 | EPI_ISL_2880332 | India/HR-IGIB-607500312049/2021 | 2021/4/30 | INSACOG at CSIR Institute of Genomics and Integrative Biology | INSACOG et al |
| AY.46.4 | EPI_ISL_6332970 | Uganda/UG974/2021 | 2021/5/27 | MRC/UVRI & LSHTM Uganda Research Unit , Rakai Health Sciences Program | Matthew Cotten Dan Lule Bugembe et al |
| AY.46.5 | EPI_ISL_4307012 | England/NORW-302AE27/2021 | 2021/5/31 | COVID-19 Genomics UK (COG-UK) Consortium | Dave J. Baker et al |
| AY.47 | EPI_ISL_1969244 | Indonesia/SS-NIHRD-WGS00441/2021 | 2021/1/8 | National Institute of Health Research and Development | Vivi Setiawaty et al |
| AY.49 | EPI_ISL_4435563 | India/DL-NCDC-300911/2021 | 2021/4/21 | NCDC Delhi, Biotechnology Division - INSACOG | Radhakrishnan V. S et al |
| AY.5 | EPI_ISL_1697977 | England/MILK-150A709/2021 | 2021/4/14 | Wellcome Sanger Institute for the COVID-19 Genomics UK (COG-UK) Consortium | The Lighthouse Lab in Milton Keynes et al |
| AY.5.1 | EPI_ISL_2695446 | Portugal/PT9522/2021 | 2021/6/4 | Instituto Nacional de Saude (INSA) | Borges et al |
| AY.5.2 | EPI_ISL_2810480 | Portugal/PT10387/2021 | 2021/5/31 | Instituto Nacional de Saude (INSA) | Borges et al |
| AY.5.4 | EPI_ISL_3730676 | Switzerland/BE-UHB-43000628/2021 | 2021/2/13 | Clinical Bacteriology | Tim Roloff et al |
| AY.50 | EPI_ISL_6151639 | India/MH-RFIP01634/2021 | 2021/3/29 | RF-IISER Pune | Krishanpal Karmodiya et al |
| AY.50 | EPI_ISL_2546359 | India/MH-IGIB-NIV-INSACOG-GSEQ-1977/2021 | 2021/3/29 | NIV Influenza | Potdar et al |

| | | | | | |
|---|---|---|---|---|---|
| AY.51 | EPI_ISL_2231615 | India/AP-CCMB-CIA1177/2021 | 2021/4/20 | CSIR-Centre for Cellular and Molecular Biology-INSACOG | Lamuk Zaveri et al |
| AY.54 | EPI_ISL_1663304 | India/ILSGS00608/2021 | 2021/3/15 | Institute of Life Sciences - INSACOG | Sunil K. Raghav et al |
| AY.55 | EPI_ISL_4768098 | Rwanda/CV2200/2021 | 2021/1/29 | Africa Centre for Excellence for Genomics of Infectious Diseases (ACEGID), Redeemer's University | Enatha Mukantwari et al |
| AY.56 | EPI_ISL_6023787 | India/DL-ILBS-104480/2021 | 2021/4/6 | ILBS | Ekta Gupta et al |
| AY.57 | EPI_ISL_4415497 | India/MH-IISER-Pune-IP01466/2021 | 2021/1/9 | IISER Pune-INSACOG | Krishanpal Karmodiya et al |
| AY.59 | EPI_ISL_4101580 | Malaysia/C19UMB104/2021 | 2021/3/17 | UKM Medical Molecular Biology Institute (UMBI) | Nur Alyaa Afifah Md Shahri et al |
| AY.6 | EPI_ISL_4884592 | Italy/SIC_CQRC_2202858/2021 | 2021/3/16 | CQRC_QUALITY CONTROL CHEMICAL BIOLOGICAL RISK_AOOR Villa Sofia Cervello Palermo | Di Gaudio F. et al |
| AY.61 | EPI_ISL_4415492 | India/MH-IISER-Pune-IP01458/2021 | 2021/1/9 | IISER Pune-INSACOG | Krishanpal Karmodiya et al |
| AY.61 | EPI_ISL_2180642 | USA/FL-CDC-ASC210061106/2021 | 2021/5/8 | Centers for Disease Control and Prevention Division of Viral Diseases, Pathogen Discovery | Dakota Howard et al |
| AY.61 | EPI_ISL_3844203 | India/GJ-INSACOG-GBRC1747/2021 | 2021/4/16 | Gujarat Biotechnology Research Centre | Arpit Shukla et al |
| AY.62 | EPI_ISL_2180457 | Bangladesh/IEDCR-OIS-0016/2021 | 2021/5/12 | IEDCR-ideSHi-icddr,b | Hassan Afrad et al |
| AY.62 | EPI_ISL_2042958 | USA/TX-CDC-ASC210034732/2021 | 2021/4/27 | Centers for Disease Control and Prevention Division of Viral Diseases, Pathogen Discovery | Dakota Howard et al |
| AY.63 | EPI_ISL_2673741 | Norway/14196/2021 | 2021/6/10 | Norwegian Institute of Public Health, Department of Virology | Kathrine Stene-Johansen et al |
| AY.65 | EPI_ISL_1660427 | Bahrain/840056464/2021 | 2021/4/10 | Communicable Disease Laboratory, Public Health Directorate | Alwasti et al |
| AY.66 | EPI_ISL_4884877 | Italy/SIC_CQRC_21066357/2021 | 2021/3/16 | CQRC_QUALITY CONTROL CHEMICAL BIOLOGICAL RISK_AOOR Villa Sofia Cervello Palermo | Di Gaudio F. et al |
| AY.66 | EPI_ISL_4884879 | Italy/SIC_CQRC_2202359/2021 | 2021/3/16 | CQRC_QUALITY CONTROL CHEMICAL BIOLOGICAL RISK_AOOR Villa Sofia Cervello Palermo | Di Gaudio F. et al |

| | | | | | |
|---|---|---|---|---|---|
| AY.7 | EPI_ISL_2006067 | England/MILK-15551C5/2021 | 2021/5/1 | Wellcome Sanger Institute for the COVID-19 Genomics UK (COG-UK) Consortium | The Lighthouse Lab in Milton Keynes et al |
| AY.7.1 | EPI_ISL_4483966 | Slovakia/UVZ_PL36_G11_17989/2021 | 2021/2/9 | Laboratory of Genomics and Bioinformatics, Comenius University Science Park | Tatiana Sedláčková et al |
| AY.7.2 | EPI_ISL_3039518 | France/BRE_8600637102/2021 | 2021/6/27 | CHU Pontchaillou | GROLHIER Claire et al |
| AY.71 | EPI_ISL_2881064 | India/UP-IGIB-ES_186/2021 | 2021/4/1 | INSACOG at CSIR Institute of Genomics and Integrative Biology | INSACOG et al |
| AY.71 | EPI_ISL_4101625 | Turkey/HSGM-E2940/2021 | 2021/3/9 | Ministry of Health Turkey | Fatma Bayrakdar et al |
| AY.72 | EPI_ISL_3844219 | India/GJ-INSACOG-GBRC1740/2021 | 2021/4/15 | Gujarat Biotechnology Research Centre | Bhadreshsinh Gohil et al |
| AY.72 | EPI_ISL_3144453 | Italy/APU-POLBA231/2021 | 2021/7/16 | University of Bari Biomedical Sciences and Human Oncology | Maria Chironna et al |
| AY.75 | EPI_ISL_4062648 | Oman/rega-OM-90/2021 | 2021/1/6 | KU Leuven, Rega Institute, Clinical and Epidemiological Virology | Tony Wawina-Bokalanga et al |
| AY.75 | EPI_ISL_2107083 | India/MH-NCCS-ND16337/2021 | 2021/3/21 | National Centre For Cell Science | Dhiraj Paul et al |
| AY.75 | EPI_ISL_5603889 | USA/IL-S21WGS5409/2021 | 2021/1/10 | Illinois Department of Public Health - Springfield Lab | Bryan Sim et al |
| AY.75 | EPI_ISL_2897619 | India/MH-NS1904/2021 | 2021/4/24 | CDFD | Murali Bashyam et al |
| AY.75 | EPI_ISL_2546522 | India/GA-IGIB-NIV-INSACOG-GSEQ-2659/2021 | 2021/4/8 | NIV Influenza | Potdar et al |
| AY.76 | EPI_ISL_2233382 | Malaysia/IMR_26135/2021 | 2021/5/4 | Institute for Medical Research, Infectious Disease Research Centre, National Institutes of Health, Ministry of Health Malaysia | Suppiah J et al |
| AY.78 | EPI_ISL_4435813 | India/DL-NCDC-3009283/2021 | 2021/4/22 | NCDC Delhi, Biotechnology Division - INSACOG | Radhakrishnan V. S et al |
| AY.79 | EPI_ISL_5052019 | Malaysia/UNIMAS-GHML333/2021 | 2021/3/9 | Institute of Health and Community Medicine | Chan Chia Jui et al |
| AY.8 | EPI_ISL_1790928 | England/MILK-1526901/2021 | 2021/4/21 | Wellcome Sanger Institute for the COVID-19 Genomics UK (COG-UK) Consortium | The Lighthouse Lab in Milton Keynes et al |
| AY.80 | EPI_ISL_3843805 | India/GJ-INSACOG-GBRC1806/2021 | 2021/6/1 | Gujarat Biotechnology Research Centre | Nitin Shukla et al |
| AY.82 | EPI_ISL_4820060 | Fiji/FJ075/2021 | 2021/4/25 | Microbiological Diagnostic Unit - Public Health Laboratory (MDU-PHL) | Sahukhan et al |

| | | | | | |
|---|---|---|---|---|---|
| AY.84 | EPI_ISL_2107500 | Australia/NSW-R0239/2021 | 2021/4/30 | Microbiology RPAH | Foster et al |
| AY.84 | EPI_ISL_4483993 | Slovakia/UVZ_PL37_B1_18000/2021 | 2021/4/9 | Laboratory of Genomics and Bioinformatics, Comenius University Science Park | Tatiana Sedláčková et al |
| AY.85 | EPI_ISL_4647511 | Thailand/DMSc-02063/2021 | 2021/1/9 | Division of Genomic Medicine and Innovation support,Department of Medical Sciences, Ministry of Public Health, Thailand | Surakameth Mahasirimongkol et al |
| AY.88 | EPI_ISL_3914982 | Togo/NMIMR-21-TGS-155/2021 | 2021/4/12 | Noguchi Memorial Institute for Medical Research, University of Ghana, Legon, Ghana | Afiwa W. Halatoko et al |
| AY.9 | EPI_ISL_3776795 | England/PHEC-U307UAAE/2021 | 2021/3/27 | COVID-19 Genomics UK (COG-UK) Consortium | PHE Covid Sequencing Team et al |
| AY.9.2.1 | EPI_ISL_4220741 | Canada/QC-L00370038001/2021 | 2021/7/27 | Laboratoire de santé publique du Québec | Sandrine Moreira et al |
| AY.90 | EPI_ISL_2022131 | England/MILK-1557372/2021 | 2021/5/2 | Wellcome Sanger Institute for the COVID-19 Genomics UK (COG-UK) Consortium | The Lighthouse Lab in Milton Keynes et al |
| AY.91.1 | EPI_ISL_5327896 | Namibia/CERI-KRISP-K022511/2021 | 2021/6/7 | CERI, Centre for Epidemic Response and Innvoation, Stellenbosch University and KRISP, KZN Research Innovation and Sequencing Platform, UKZN. | Iyaloo Constantinus et al |
| AY.93 | EPI_ISL_2479874 | Canada/AB-ABPHL-17309/2021 | 2021/4/25 | Public Health Agency of Canada (PHAC) National Microbiology Laboratory | Buss et al |
| AY.94 | EPI_ISL_4884550 | Italy/SIC_CQRC_21070336/2021 | 2021/3/16 | CQRC_QUALITY CONTROL CHEMICAL BIOLOGICAL RISK_AOOR Villa Sofia Cervello Palermo | Di Gaudio F. et al |
| AY.97 | EPI_ISL_1994608 | SouthAfrica/KRISP-K014463_2/2021 | 2021/4/30 | KRISP, KZN Research Innovation and Sequencing Platform | Giandhari J et al |
| AY.98.1 | EPI_ISL_5104615 | Slovenia/17-047667-CE/2020 | 2020/11/13 | NLZOH (National Laboratory for Health, Environment and Food) / CISLD (Clinical Institute of Special Laboratory Diagnostics), University Children's Hospital, University Medical Center Ljubljana | Sandra Janezic et al |
| AY.99.1 | EPI_ISL_2379633 | India/TN-C_83_S69_R1_001/2021 | 2021/4/29 | inStem NCBS - INSACOG | Uma Ramakrishnan Dasaradhi Palakodeti Aswin SaiNarain et al |

| | | | | | |
|---|---|---|---|---|---|
| AZ.1 | EPI_ISL_4026322 | Nigeria/BCVL-21182/2021 | 2021/1/22 | Northwestern University - Center for Pathogen Genomics and Microbial Evolution | Ramon Lorenzo-Redondo et al |
| AZ.2 | EPI_ISL_2370918 | Greece/196448/2021 | 2021/2/5 | Greek Genome Center, Biomedical Research Foundation of the Academy of Athens (BRFAA) | Emmanouil Athanasiadis et al |
| AZ.2.1 | EPI_ISL_1361866 | Germany/ST-MD1536/2021 | 2021/3/15 | Institute of Medical Microbiology and Hospital Hygiene | Prof. Dr. Achim Kaasch et al |
| AZ.3 | EPI_ISL_4964826 | USA/TX-CDC-LC0013987/2021 | 2021/2/8 | Centers for Disease Control and Prevention Division of Viral Diseases, Pathogen Discovery | Dakota Howard et al |
| AZ.4 | EPI_ISL_1180300 | Ireland/D-NVRL-21IRL22786/2021 | 2021/2/8 | National Virus Reference Laboratory | Guerrino Macori et al |
| AZ.5 | EPI_ISL_2156221 | Philippines/PH-PGC-06126/2021 | 2021/2/9 | Philippine Genome Center | Francis A. Tablizo et al |
| B | EPI_ISL_402123 | Wuhan/IPBCAMS-WH-01/2019 | 2019/12/24 | Institute of Pathogen Biology, Chinese Academy of Medical Sciences & Peking Union Medical College | Lili Ren et al |
| B | EPI_ISL_417253 | England/20108006603/2020 | 2020/3/3 | Respiratory Virus Unit, Microbiology Services Colindale, Public Health England | Monica Galiano et al |
| B | EPI_ISL_420796 | USA/TX-CDC-3063244-001/2020 | 2020/2/29 | Pathogen Discovery, Respiratory Viruses Branch, Division of Viral Diseases, Centers for Disease Control and Prevention | Krista Queen et al |
| B | EPI_ISL_457853 | Kenya/P1118/2020 | 2020/4/19 | KEMRI-Wellcome Trust Research Programme/KEMRI-CGMR-C Kilifi | Githinji G. et al 2020 et al |
| B | EPI_ISL_2694591 | Iran/GRC-10/2020 | 2020/5/17 | Genetics Research Center, University of Social Welfare and Rehabilitation Sciences | Zohreh Fattahi et al |
| B.1 | EPI_ISL_4537783 | France/20_B./2020 | 2020/1/1 | APHP Bichat-Claude Bernard | Samuel Lebourgeois et al |
| B.1 | EPI_ISL_452107 | USA/LA-CDC-03069817-001/2020 | 2020/3/10 | Pathogen Discovery, Respiratory Viruses Branch, Division of Viral Diseases, Centers for Disease Control and Prevention | Yan Li et al |
| B.1 | EPI_ISL_3138689 | Belgium/MR71NY1731/2020 | 2020/5/4 | Microbiology Department - University Hospital Brussel | Thomas Demuyser et al |
| B.1 | EPI_ISL_1300948 | USA/NY-MSHSPSP-PV15333/2020 | 2020/7/27 | MSHS Pathogen Surveillance Program | Ana S. Gonzalez-Reiche et al |

| | | | | | |
|---|---|---|---|---|---|
| B.1 | EPI_ISL_2445035 | USA/AZ-ASPHL-1991/2020 | 2020/7/7 | Arizona State Public Health Laboratory | Trung Huynh et al |
| B.1 | EPI_ISL_2443765 | USA/CA-UCI-079/2020 | 2020/4/13 | University of California, Irvine | Amanda N. Pinski et al |
| B.1 | EPI_ISL_2884955 | USA/MD-HP06635-PIDLEHQTMF/2020 | 2020/4/2 | Johns Hopkins Hospital Department of Pathology | C. Paul Morris et al |
| B.1 | EPI_ISL_2219597 | USA/TX-HMH-MCoV-35067/2020 | 2020/8/11 | Houston Methodist Hospital | Randall J. Olsen et al |
| B.1 | EPI_ISL_769328 | England/MILK-C61001/2020 | 2020/12/13 | Wellcome Sanger Institute for the COVID-19 Genomics UK (COG-UK) Consortium | The Lighthouse Lab in Milton Keynes et al |
| B.1 | EPI_ISL_3410636 | USA/MA-UMASSMED-P006E05/2020 | 2020/4/30 | Center for Microbiome Research | Richard T. Ellison III et al |
| B.1 | EPI_ISL_3275378 | SouthAfrica/KRISP-K016875/2021 | 2021/4/9 | KRISP, KZN Research Innovation and Sequencing Platform | Laguda-Akingba O et al |
| B.1 | EPI_ISL_5384449 | USA/TX-HMH-MCoV-49669/2020 | 2020/7/11 | Houston Methodist Hospital | Randall J. Olsen et al |
| B.1 | EPI_ISL_4419636 | Canada/BC-BCCDC-165732/2020 | 2020/10/25 | BCCDC Public Health Laboratory | Prystajecky Natalie et al |
| B.1.1 | EPI_ISL_3568416 | Australia/NSW-SAVID-4085/2020 | 2020/1/8 | Virology Research Laboratory; Area of Virology, Serology and Virology Division (SAViD), New South Wales Health Pathology Randwick | Foster et al |
| B.1.1 | EPI_ISL_2367394 | Switzerland/VD-CHUV-GEN2537/2020 | 2020/3/16 | Laboratory of genomics and metagenomics | Trestan Pillonel et al |
| B.1.1 | EPI_ISL_5248196 | Ireland/D-U-MAY-000235/2020 | 2020/3/18 | Irish Coronavirus Sequencing Consortium - Maynooth University | David A Fitzpatrick et al |
| B.1.1 | EPI_ISL_465738 | England/20122017204/2020 | 2020/3/15 | Respiratory Virus Unit, Microbiology Services Colindale, Public Health England | PHE Covid Sequencing Team et al |
| B.1.1 | EPI_ISL_470842 | Australia/WA24/2020 | 2020/4/1 | PathWest Laboratory Medicine WA | Chisha Sikazwe et al |
| B.1.1 | EPI_ISL_512751 | Australia/WA98/2020 | 2020/3/29 | PathWest Laboratory Medicine WA Microbial Surveillance Unit | PathWest Laboratory Medicine WA Microbial Surveillance Unit et al |
| B.1.1 | EPI_ISL_762994 | NorthMacedonia/ZMC5839/2020 | 2020/6/3 | Laboratory of Genetics and Personalized Medicine, Zan Mitrev Clinic | Kungulovski et al. |
| B.1.1 | EPI_ISL_775881 | Germany/HH-hpi-p2081/2020 | 2020/3/20 | Heinrich Pette Institute, Leibniz Institute for Experimental Virology | Alexis Robitaille et al |
| B.1.1 | EPI_ISL_3730873 | Italy/TAA-APSS-Ph2_RUN2_NB24/2020 | 2020/3/31 | U.O. Microbiologia e Virologia, Azienda Provinciale per i Servizi Sanitari Provincia Autonoma di Trento, Ospedale S.Chiara | Mirko Moser et al |

| | | | | | |
|---|---|---|---|---|---|
| B.1.1 | EPI_ISL_2596117 | France/PAC-IHU-11363_Illu1/2020 | 2020/3/20 | MEPHI, Aix Marseille University | Anthony LEVASSEUR et al |
| B.1.1 | EPI_ISL_1577269 | Italy/CAM-CRGS-634/2021 | 2021/3/3 | 1. Genome Research Center for Health (CRGS) / 2. Laboratory of Molecular Medicine and Genomics(LMMGe) / 3. Center for Research in Pure and Applied Mathematics (CRMPA) | Giorgio Giurato (Corresponding Author) et al |
| B.1.1 | EPI_ISL_685346 | Japan/PG-1740/2020 | 2020/5/2 | Pathogen Genomics Center, National Institute of Infectious Diseases | Tsuyoshi Sekizuka et al |
| B.1.1 | EPI_ISL_730576 | Turkey/Ankara_GUMV_10711/2020 | 2020/6/25 | Gazi University Faculty of Medicine, Medical Virology Laboratory | Erdem Şahin et al |
| B.1.1 | EPI_ISL_1181730 | Italy/VEN-SI_125/2020 | 2020/3/20 | Department of Molecular Medicine,Computational Medicine Group,Univeresity of Padova,Padova,Italy | Elisa Franchin et al |
| B.1.1 | EPI_ISL_684712 | Japan/PG-1102/2020 | 2020/4/22 | Pathogen Genomics Center, National Institute of Infectious Diseases | Tsuyoshi Sekizuka et al |
| B.1.1 | EPI_ISL_3134730 | Brazil/PE-FIOCRUZ-IAM2136/2020 | 2020/9/2 | WallauLab on behalf of Fiocruz COVID-19 Genomic Surveillance Network | Marcelo Henrique dos Santos Paiva et al |
| B.1.1 | EPI_ISL_5074032 | USA/TX-HMH-MCoV-42583/2020 | 2020/7/19 | Houston Methodist Hospital | Randall J. Olsen et al |
| B.1.1 | EPI_ISL_5384262 | USA/TX-HMH-MCoV-49490/2020 | 2020/7/13 | Houston Methodist Hospital | Randall J. Olsen et al |
| B.1.1.1 | EPI_ISL_416140 | Denmark/SSI-05/2020 | 2020/3/2 | Statens Serum Institute | Morten Rasmussen et al |
| B.1.1.1 | EPI_ISL_1534645 | Peru/LIM-INS-869/2020 | 2020/11/30 | Laboratorio de Referencia Nacional de Biotecnología y Biología Molecular. Instituto Nacional de Salud Perú | Carlos Padilla Rojas et al |
| B.1.1.10 | EPI_ISL_602518 | Germany/NW-HHU-148/2020 | 2020/3/1 | Center of Medical Microbiology, Virology, and Hospital Hygiene, University of Duesseldorf | Olympia E. Anastasiou et al |
| B.1.1.100 | EPI_ISL_620841 | Denmark/DCGC-5516/2020 | 2020/9/21 | Albertsen lab, Department of Chemistry and Bioscience, Aalborg University, Denmark | Danish Covid-19 Genome Consortia et al |
| B.1.1.101 | EPI_ISL_454565 | India/MH-NIV-QA-710/2020 | 2020/3/31 | NIV Influenza | Potdar et al |
| B.1.1.107 | EPI_ISL_473446 | England/BIRM-61243/2020 | 2020/6/6 | COVID-19 Genomics UK (COG-UK) Consortium | Institute of Microbiology et al |
| B.1.1.109 | EPI_ISL_559070 | England/ALDP-52A636/2020 | 2020/6/1 | Wellcome Sanger Institute for the COVID-19 Genomics UK (COG-UK) consortium | The Lighthouse Lab in Alderley Park et al |

| | | | | | |
|---|---|---|---|---|---|
| B.1.1.110 | EPI_ISL_1239429 | Finland/24MS1A6/2020 | 2020/3/24 | Department of Virology, Faculty of Medicine, University of Helsinki, Helsinki, Finland | Teemu Smura et al |
| B.1.1.111 | EPI_ISL_644795 | Zimbabwe/ZW-6978/2020 | 2020/5/1 | Quadram Institute Bioscience | Thanh Le Viet et al |
| B.1.1.112 | EPI_ISL_523614 | Netherlands/ZH-EMC-272/2020 | 2020/5/8 | Erasmus Medical Center | Bas Oude Munnink et al |
| B.1.1.113 | EPI_ISL_467608 | USA/NM-UNM-00027/2020 | 2020/3/15 | Center for Global Health, University of New Mexico Health Sciences Center | Daryl Domman et al |
| B.1.1.114 | EPI_ISL_432321 | Wales/PHWC-27B02/2020 | 2020/3/30 | Public Health Wales Microbiology Cardiff | Catherine Moore et al |
| B.1.1.115 | EPI_ISL_580835 | England/ALDP-9EDB5A/2020 | 2020/5/9 | Wellcome Sanger Institute for the COVID-19 Genomics UK (COG-UK) consortium | Jacquelyn Wynn et al |
| B.1.1.116 | EPI_ISL_497204 | USA/WA-S2572/2020 | 2020/6/17 | Seattle Flu Study | Deborah A. Nickerson et al |
| B.1.1.117 | EPI_ISL_500460 | Brazil/PE-FIOCRUZ-IAM08/2020 | 2020/4/7 | WallauLab, Aggeu Magalhaes Institute | Marcelo Henrique Santos Paiva et al |
| B.1.1.118 | EPI_ISL_545087 | USA/TX-HMH-2050/2020 | 2020/5/11 | Houston Methodist Hospital | S. Wesley Long et al |
| B.1.1.119 | EPI_ISL_489205 | NorthernIreland/NIRE-103831/2020 | 2020/4/15 | Wellcome Sanger Institute for the COVID-19 Genomics UK (COG-UK) consortium | Conall McCaughey et al |
| B.1.1.12 | EPI_ISL_559004 | England/ALDP-519746/2020 | 2020/6/2 | Wellcome Sanger Institute for the COVID-19 Genomics UK (COG-UK) consortium | The Lighthouse Lab in Alderley Park et al |
| B.1.1.120 | EPI_ISL_559078 | England/ALDP-52AAA3/2020 | 2020/6/1 | Wellcome Sanger Institute for the COVID-19 Genomics UK (COG-UK) consortium | The Lighthouse Lab in Alderley Park et al |
| B.1.1.121 | EPI_ISL_484153 | England/GSTT-265E0F8/2020 | 2020/3/31 | COVID-19 Genomics UK (COG-UK) Consortium | Chloe Fisher et al |
| B.1.1.122 | EPI_ISL_444168 | England/LOND-D4A07/2020 | 2020/4/10 | COVID-19 Genomics UK (COG-UK) Consortium | Sergi Castellano et al |
| B.1.1.123 | EPI_ISL_589485 | England/ALDP-9ED64A/2020 | 2020/5/18 | Wellcome Sanger Institute for the COVID-19 Genomics UK (COG-UK) consortium | Jacquelyn Wynn et al |
| B.1.1.125 | EPI_ISL_461931 | England/NOTT-11184F/2020 | 2020/5/21 | COVID-19 Genomics UK (COG-UK) Consortium | Gemma Clark et al |
| B.1.1.127 | EPI_ISL_450256 | Russia/StPetersburg-RII7520S/2020 | 2020/4/20 | WHO National Influenza Centre Russian Federation | Andrey Komissarov et al |
| B.1.1.128 | EPI_ISL_515459 | USA/NV-NSPHL-A0205/2020 | 2020/6/2 | Nevada State Public Health Laboratory | Richard Tillett et al |

| | | | | | |
|---|---|---|---|---|---|
| B.1.1.129 | EPI_ISL_430084 | Russia/StPetersburg-RII5017V/2020 | 2020/4/9 | WHO National Influenza Centre Russian Federation | Andrey Komissarov et al |
| B.1.1.13 | EPI_ISL_524622 | England/LOND-126027C/2020 | 2020/3/14 | Wellcome Sanger Institute for the COVID-19 Genomics UK (COG-UK) consortium | Ling Li et al |
| B.1.1.130 | EPI_ISL_536998 | England/QEUH-961848/2020 | 2020/7/31 | Wellcome Sanger Institute for the COVID-19 Genomics UK (COG-UK) consortium | Harper VanSteenhouse et al |
| B.1.1.132 | EPI_ISL_631667 | USA/TX-DSHS-1205/2020 | 2020/3/17 | Texas Department of State Health Services | Rashmi Tuladhar et al |
| B.1.1.133 | EPI_ISL_914979 | Canada/NS-NML-912/2020 | 2020/3/22 | National Microbiology Laboratory (NML) | Anna Majer et al |
| B.1.1.134 | EPI_ISL_432425 | Wales/PHWC-28600/2020 | 2020/4/1 | Public Health Wales Microbiology Cardiff | Catherine Moore et al |
| B.1.1.135 | EPI_ISL_475129 | Sweden/20-50299/2020 | 2020/3/12 | The Public Health Agency of Sweden | Oskar Karlsson Lindsjo et al |
| B.1.1.136 | EPI_ISL_480662 | Australia/VIC2020/2020 | 2020/6/3 | VIDRL and MDU-PHL | Caly L. et al |
| B.1.1.137 | EPI_ISL_420173 | England/SHEF-C08F5/2020 | 2020/3/24 | Department of Infection, Immunity and Cardiovascular Disease, The Florey Institute, The Medical School, University of Sheffield | Thushan de Silva et al |
| B.1.1.138 | EPI_ISL_540217 | Wales/QEUH-9A4506/2020 | 2020/4/4 | Wellcome Sanger Institute for the COVID-19 Genomics UK (COG-UK) consortium | Harper VanSteenhouse et al |
| B.1.1.138 | EPI_ISL_814089 | Turkey/HSGM-2004082/2020 | 2020/5/15 | Ministry of Health Turkey | Fatma Bayrakdar et al |
| B.1.1.139 | EPI_ISL_857000 | USA/CA-CZB-9973/2020 | 2020/6/17 | Chan-Zuckerberg Biohub | CZB Clianub Consortium et al |
| B.1.1.14 | EPI_ISL_433117 | Scotland/EDB1683/2020 | 2020/4/16 | COVID-19 Genomics UK (COG-UK) Consortium | McHugh M et al |
| B.1.1.141 | EPI_ISL_596264 | Russia/NVS-RII-MH1087V/2020 | 2020/7/22 | WHO National Influenza Centre Russian Federation | Andrey Komissarov et al |
| B.1.1.142 | EPI_ISL_475831 | Austria/CeMM0391/2020 | 2020/3/10 | Bergthaler laboratory, CeMM Research Center for Molecular Medicine of the Austrian Academy of Sciences | Alexandra Popa et al |
| B.1.1.144 | EPI_ISL_1495592 | Austria/CeMM5636/2020 | 2020/8/25 | Bergthaler laboratory, CeMM Research Center for Molecular Medicine of the Austrian Academy of Sciences | Lukas Endler et al |
| B.1.1.147 | EPI_ISL_492193 | England/CAMB-1B4820/2020 | 2020/6/4 | Wellcome Sanger Institute for the COVID-19 Genomics UK (COG-UK) consortium | Luke W Meredith et al |

| | | | | | |
|---|---|---|---|---|---|
| B.1.1.148 | EPI_ISL_470028 | NorthernIreland/NIRE-FC368/2020 | 2020/4/12 | Wellcome Sanger Institute for the COVID-19 Genomics UK (COG-UK) consortium | Conall McCaughey et al |
| B.1.1.149 | EPI_ISL_448406 | England/NOTT-110CEA/2020 | 2020/5/2 | COVID-19 Genomics UK (COG-UK) Consortium | Gemma Clark et al |
| B.1.1.15 | EPI_ISL_554634 | England/MILK-957D82/2020 | 2020/6/9 | Wellcome Sanger Institute for the COVID-19 Genomics UK (COG-UK) consortium | The Lighthouse Lab in Milton Keynes et al |
| B.1.1.152 | EPI_ISL_596363 | Russia/SPE-RII-MH953S/2020 | 2020/9/3 | WHO National Influenza Centre Russian Federation | Andrey Komissarov et al |
| B.1.1.153 | EPI_ISL_889356 | CzechRepublic/IAB20_016_031/2020 | 2020/8/6 | Institute of Applied Biotechnologies a.s. | Petr Klempt et al |
| B.1.1.154 | EPI_ISL_432628 | England/SHEF-D13D2/2020 | 2020/3/30 | COVID-19 Genomics UK (COG-UK) Consortium | Thushan de Silva et al |
| B.1.1.155 | EPI_ISL_559032 | England/ALDP-51F142/2020 | 2020/6/2 | Wellcome Sanger Institute for the COVID-19 Genomics UK (COG-UK) consortium | The Lighthouse Lab in Alderley Park et al |
| B.1.1.157 | EPI_ISL_1523399 | Canada/QC-HMR-97204348/2020 | 2020/5/20 | Laboratoire de santé publique du Québec | Sandrine Moreira et al |
| B.1.1.158 | EPI_ISL_496205 | USA/WA-S1779/2020 | 2020/6/1 | Seattle Flu Study | Deborah A. Nickerson et al |
| B.1.1.159 | EPI_ISL_596770 | Australia/WA207/2020 | 2020/8/14 | PathWest Laboratory Medicine WA Microbial Surveillance Unit | PathWest Laboratory Medicine WA Microbial Surveillance Unit et al |
| B.1.1.16 | EPI_ISL_434473 | Greece/136/2020 | 2020/3/11 | Laboratory of Biology, Department of Medicine, Democritus University of Thrace | Kassela K. et al |
| B.1.1.160 | EPI_ISL_489355 | NorthernIreland/NIRE-100184/2020 | 2020/5/3 | Wellcome Sanger Institute for the COVID-19 Genomics UK (COG-UK) consortium | Conall McCaughey et al |
| B.1.1.161 | EPI_ISL_895763 | CzechRepublic/FNHK-03501/2020 | 2020/3/19 | Molecular biology division, Institute of Clinical Biochemistry and Diagnostics, Charles University, Faculty of Medicine in Hradec Králové and University Hospital Hradec Králové | Helena Kovaříková et al |
| B.1.1.162 | EPI_ISL_1586301 | Iceland/1014/2020 | 2020/3/21 | deCODE genetics | Daniel F Gudbjartsson et al |
| B.1.1.163 | EPI_ISL_486829 | Russia/CRIE187963/2020 | 2020/4/1 | Group of Genomics and Postgenomic Technologies of Central Research Institute of Epidemilology | Speranskaya AS et al |

| | | | | | |
|---|---|---|---|---|---|
| B.1.1.164 | EPI_ISL_479933 | Japan/PG-0243/2020 | 2020/3/16 | Pathogen Genomics Center, National Institute of Infectious Diseases | Tsuyoshi Sekizuka et al |
| B.1.1.165 | EPI_ISL_590390 | England/ALDP-9B5C0D/2020 | 2020/8/28 | Wellcome Sanger Institute for the COVID-19 Genomics UK (COG-UK) consortium | Jacquelyn Wynn et al |
| B.1.1.168 | EPI_ISL_458605 | England/NORT-289744/2020 | 2020/3/31 | Wellcome Sanger Institute for the COVID-19 Genomics UK (COG-UK) consortium | Chris Duncan et al |
| B.1.1.169 | EPI_ISL_545081 | USA/TX-HMH-2042/2020 | 2020/5/4 | Houston Methodist Hospital | S. Wesley Long et al |
| B.1.1.17 | EPI_ISL_417757 | Iceland/62/2020 | 2020/3/6 | deCODE genetics | Daniel F Gudbjartsson et al |
| B.1.1.170 | EPI_ISL_532876 | Scotland/QEUH-7B078B/2020 | 2020/7/19 | Wellcome Sanger Institute for the COVID-19 Genomics UK (COG-UK) consortium | Ana da Silva Filipe et al |
| B.1.1.171 | EPI_ISL_420489 | England/20129040602/2020 | 2020/3/20 | Respiratory Virus Unit, Microbiology Services Colindale, Public Health England | Monica Galiano et al |
| B.1.1.172 | EPI_ISL_475663 | USA/CA-CSMC50/2020 | 2020/4/6 | Cedars-Sinai Medical Center, Molecular Pathology Laboratory of Department of Pathology & Laboratory Medicine and Genomic Core | Wenjuan Zhang et al |
| B.1.1.175 | EPI_ISL_605905 | Bangladesh/DNAS-RAB-isl-118/2020 | 2020/4/29 | NGS Lab, DNA SOLUTION LTD. | Khan et al |
| B.1.1.176 | EPI_ISL_535744 | Canada/QC-LSPQ-L00216190/2020 | 2020/3/12 | Laboratoire de santé publique du Québec | Sandrine Moreira et al |
| B.1.1.177 | EPI_ISL_545532 | USA/TX-HMH-2757/2020 | 2020/6/4 | Houston Methodist Hospital | S. Wesley Long et al |
| B.1.1.178 | EPI_ISL_441289 | England/CAMB-7A14A/2020 | 2020/4/2 | Wellcome Sanger Institute for the COVID-19 Genomics UK (COG-UK) consortium | Luke W Meredith et al |
| B.1.1.180 | EPI_ISL_517875 | USA/FL-BPHL-0985/2020 | 2020/6/8 | Florida Bureau of Public Health Laboratories | Sarah Schmedes et al |
| B.1.1.181 | EPI_ISL_586305 | Canada/ON-S1230/2020 | 2020/3/22 | McMaster University | Allison McGeer et al |
| B.1.1.182 | EPI_ISL_582662 | UnitedArabEmirates/skmc-3580730/2020 | 2020/9/9 | Molecular/Surveillance lab Sheikh Khalifa Medical City | Amirtharaj Francis et al |
| B.1.1.184 | EPI_ISL_569735 | Russia/OMS-ORINFI-5710S/2020 | 2020/5/8 | WHO National Influenza Centre Russian Federation | Artem Fadeev et al |
| B.1.1.185 | EPI_ISL_455196 | Netherlands/NA_394/2020 | 2020/3/31 | Erasmus Medical Center | Bas Oude Munnink et al |

| | | | | | |
|---|---|---|---|---|---|
| B.1.1.186 | EPI_ISL_1533465 | USA/GA-CDC-4049061-001/2020 | 2020/6/25 | Genomics and Discovery, Respiratory Viruses Branch, Division of Viral Diseases, Centers for Disease Control and Prevention | Yan Li et al |
| B.1.1.187 | EPI_ISL_778771 | Italy/CAM-TIGEM-452/2020 | 2020/6/24 | TIGEM | Antonio Grimaldi et al |
| B.1.1.189 | EPI_ISL_707793 | Tunisia/9111/2020 | 2020/4/11 | 1-Clinical and Experimental Pharmacology Lab, LR16SP02, National Center of Pharmacovigilance, University of Tunis El Manar, Tunis, Tunisia. 2-Neurodegenerative diseases and psychiatric troubles, LR18SP03, Razi Hospital, University of Tunis El Manar, Tunis, Tunisia. 3- Ministry of Health, National Observatory of New and Emerging Diseases, 1006, Tunis, Tunisia | Ilhem Boutiba-Ben Boubaker et al |
| B.1.1.190 | EPI_ISL_3568108 | Australia/NSW-SAVID-3200/2020 | 2020/3/17 | Virology Research Laboratory; Area of Virology, Serology and Virology Division (SAViD), New South Wales Health Pathology Randwick | Foster et al |
| B.1.1.191 | EPI_ISL_654953 | Sweden/20-23086/2020 | 2020/3/20 | The Public Health Agency of Sweden | Anna-Malin Linde et al |
| B.1.1.192 | EPI_ISL_480297 | Bulgaria/06/2020 | 2020/3/21 | NRL-HIV | Ivan Ivanov et al |
| B.1.1.193 | EPI_ISL_600046 | England/MILK-A0556C/2020 | 2020/9/30 | Wellcome Sanger Institute for the COVID-19 Genomics UK (COG-UK) consortium | The Lighthouse Lab in Milton Keynes et al |
| B.1.1.194 | EPI_ISL_534986 | England/OXON-B49E2/2020 | 2020/3/13 | COVID-19 Genomics UK (COG-UK) Consortium | Tanya Golubchik et al |
| B.1.1.196 | EPI_ISL_551986 | England/MILK-993397/2020 | 2020/8/26 | Wellcome Sanger Institute for the COVID-19 Genomics UK (COG-UK) consortium | The Lighthouse Lab in Milton Keynes et al |
| B.1.1.197 | EPI_ISL_433888 | England/CAMB-7AEAD/2020 | 2020/4/9 | COVID-19 Genomics UK (COG-UK) Consortium | Luke W Meredith et al |
| B.1.1.198 | EPI_ISL_497864 | HongKong/HKU-200723-097/2020 | 2020/3/18 | Department of Microbiology, The University of Hong Kong | Kelvin K.W. To et al |
| B.1.1.200 | EPI_ISL_462195 | Belgium/LL-0328341/2020 | 2020/3/28 | KU Leuven, Rega Institute, Clinical and Epidemiological Virology | Tony Wawina-Bokalanga et al |

| | | | | | |
|---|---|---|---|---|---|
| B.1.1.201 | EPI_ISL_458037 | India/TN-CCMB-C7/2020 | 2020/4/29 | CSIR-Centre for Cellular and Molecular Biology | K.Kaveri et al |
| B.1.1.202 | EPI_ISL_493330 | Italy/LAZ-INMI18/2020 | 2020/5/10 | INMI Lazzaro Spallanzani IRCCS | Cesare E.M. Gruber et al |
| B.1.1.203 | EPI_ISL_520697 | UnitedArabEmirates/L3290250769/2020 | 2020/5/20 | Al Jalila Genomics Center | Ahmad Abou Tayoun et al |
| B.1.1.204 | EPI_ISL_1181818 | Italy/VEN-SI_139/2020 | 2020/3/7 | Department of Molecular Medicine,Computational Medicine Group,Univeresity of Padova,Padova,Italy | Elisa Franchin et al |
| B.1.1.205 | EPI_ISL_672264 | USA/CA-CZB-10225/2020 | 2020/5/16 | Chan-Zuckerberg Biohub | CZB Cliahub Consortium et al |
| B.1.1.207 | EPI_ISL_2425975 | Norway/2439/2020 | 2020/4/14 | Norwegian Institute of Public Health, Department of Virology | Kathrine Stene-Johansen et al |
| B.1.1.208 | EPI_ISL_440072 | England/CAMB-72D11/2020 | 2020/3/22 | Wellcome Sanger Institute for the COVID-19 Genomics UK (COG-UK) consortium | Luke W Meredith et al |
| B.1.1.209 | EPI_ISL_523469 | Netherlands/un-EMC-714/2020 | 2020/5/6 | Erasmus Medical Center | Bas Oude Munnink et al |
| B.1.1.210 | EPI_ISL_560647 | USA/DE-DHSS-F918664/2020 | 2020/3/21 | Delaware Public Health Lab | Gregory Hovan et al |
| B.1.1.213 | EPI_ISL_424033 | England/20129036204/2020 | 2020/3/20 | Respiratory Virus Unit, Microbiology Services Colindale, Public Health England | Monica Galiano et al |
| B.1.1.214 | EPI_ISL_1431487 | Japan/PG-27639/2020 | 2020/2/22 | Pathogen Genomics Center, National Institute of Infectious Diseases | Tsuyoshi Sekizuka et al |
| B.1.1.216 | EPI_ISL_508504 | India/HR-TF38/2020 | 2020/5/6 | National Institute of Biomedical Genomics | Arindam Maitra et al |
| B.1.1.216 | EPI_ISL_667577 | Japan/IC-0148/2020 | 2020/8/3 | Pathogen Genomics Center, National Institute of Infectious Diseases | Tsuyoshi Sekizuka et al |
| B.1.1.217 | EPI_ISL_559696 | England/MILK-482D1A/2020 | 2020/4/26 | Wellcome Sanger Institute for the COVID-19 Genomics UK (COG-UK) consortium | The Lighthouse Lab in Milton Keynes et al |
| B.1.1.217 | EPI_ISL_690984 | Japan/PG-2858/2020 | 2020/4/13 | Pathogen Genomics Center, National Institute of Infectious Diseases | Tsuyoshi Sekizuka et al |
| B.1.1.218 | EPI_ISL_480308 | Bulgaria/37/2020 | 2020/5/11 | NRL-HIV | Ivan Ivanov et al |
| B.1.1.219 | EPI_ISL_549128 | Norway/3104/2020 | 2020/8/4 | Norwegian Institute of Public Health, Department of Virology | Kathrine Stene-Johansen et al |
| B.1.1.220 | EPI_ISL_501573 | England/BRIS-130E0C/2020 | 2020/3/23 | Wellcome Sanger Institute for the COVID-19 Genomics UK (COG-UK) consortium | Stephanie Hutchings et al |

| | | | | | |
|---|---|---|---|---|---|
| B.1.1.221 | EPI_ISL_462211 | Belgium/VHEJ-0329357/2020 | 2020/3/29 | KU Leuven, Rega Institute, Clinical and Epidemiological Virology | Tony Wawina-Bokalanga et al |
| B.1.1.222 | EPI_ISL_837628 | Mexico/CMX-INER-0026/2020 | 2020/4/4 | Instituto Nacional de Enfermedades Respiratorias (INER) | Celia Boukadida et al |
| B.1.1.222 | EPI_ISL_4220071 | Mexico/CMX-INMEGEN-04-07-303/2020 | 2020/8/21 | Instituto Nacional de Medicina Genomica | Cedro-Tanda A et al |
| B.1.1.224 | EPI_ISL_615422 | Denmark/DCGC-3650/2020 | 2020/8/31 | Albertsen lab, Department of Chemistry and Bioscience, Aalborg University, Denmark | Danish Covid-19 Genome Consortia et al |
| B.1.1.225 | EPI_ISL_610002 | USA/TX-DSHS-0866/2020 | 2020/6/16 | Texas Department of State Health Services | Rashmi Tuladhar et al |
| B.1.1.226 | EPI_ISL_1446540 | USA/TX-TCH-TCMC04002/2020 | 2020/4/21 | Texas Children's Microbiome Center | Ruth Ann Luna et al |
| B.1.1.227 | EPI_ISL_554886 | England/ALDP-95AA1F/2020 | 2020/6/13 | Wellcome Sanger Institute for the COVID-19 Genomics UK (COG-UK) consortium | The Lighthouse Lab in Alderley Park et al |
| B.1.1.228 | EPI_ISL_696265 | USA/AZ-TG668340/2020 | 2020/5/13 | TGen North | Jolene Bowers et al |
| B.1.1.229 | EPI_ISL_788954 | Italy/APU-PODV-15925058/2020 | 2020/8/18 | Beaconlab (Bioinformatics, Evolution and Comparative Genomics lab), Dept of Biosciences, University on Milan | Iacobellis M et al |
| B.1.1.230 | EPI_ISL_545216 | USA/TX-HMH-2267/2020 | 2020/5/24 | Houston Methodist Hospital | S. Wesley Long et al |
| B.1.1.231 | EPI_ISL_604962 | USA/UT-QDX-2572/2020 | 2020/3/13 | Quest Diagnostics | Rosenthal et al |
| B.1.1.232 | EPI_ISL_602509 | Germany/NW-HHU-187/2020 | 2020/3/28 | Center of Medical Microbiology, Virology, and Hospital Hygiene, University of Duesseldorf | Olympia E. Anastasiou et al |
| B.1.1.232 | EPI_ISL_2312538 | Greece/69758/2020 | 2020/12/6 | Greek Genome Center, Biomedical Research Foundation of the Academy of Athens (BRFAA) | Emmanouil Athanasiadis et al |
| B.1.1.234 | EPI_ISL_5917781 | Romania/DJ14317/2020 | 2020/7/20 | Medical Genetics Laboratory, Regional Centre of Medical Genetics, Emergency County Hospital Craiova | Anca-Lelia (Riza) Costache et al |
| B.1.1.237 | EPI_ISL_472789 | Wales/PHWC-15E05D/2020 | 2020/5/15 | COVID-19 Genomics UK (COG-UK) Consortium | Catherine Moore et al |
| B.1.1.239 | EPI_ISL_548404 | USA/CA-CZB-3738/2020 | 2020/7/21 | Chan-Zuckerberg Biohub | CZB Cliahub Consortium et al |
| B.1.1.240 | EPI_ISL_540650 | England/MILK-992152/2020 | 2020/8/27 | COVID-19 Genomics UK (COG-UK) Consortium | Gemma Clark et al |
| B.1.1.241 | EPI_ISL_3716388 | Madagascar/IPM-05252/2020 | 2020/4/29 | Institut Pateur de Dakar | Norosoa Razanajatovo et al |

| | | | | | |
|---|---|---|---|---|---|
| B.1.1.242 | EPI_ISL_561164 | Gambia/GC19-3099/2020 | 2020/7/25 | MRCG at LSHTM Genomics lab | Abdul Karim sesay et al |
| B.1.1.243 | EPI_ISL_512640 | Ukraine/203100361/2020 | 2020/4/24 | Respiratory Virus Unit, Microbiology Services Colindale, Public Health England | PHE Covid Sequencing Team et al |
| B.1.1.244 | EPI_ISL_525784 | USA/TX-DSHS-0225/2020 | 2020/4/7 | Texas Department of State Health Services | Jenny Zhang et al |
| B.1.1.249 | EPI_ISL_472490 | Wales/PHWC-15C462/2020 | 2020/3/26 | COVID-19 Genomics UK (COG-UK) Consortium | Catherine Moore et al |
| B.1.1.25 | EPI_ISL_458372 | England/BRIS-129932/2020 | 2020/3/31 | Wellcome Sanger Institute for the COVID-19 Genomics UK (COG-UK) consortium | Stephanie Hutchings et al |
| B.1.1.251 | EPI_ISL_812233 | USA/ND-NDDH-0001/2020 | 2020/6/2 | North Dakota Department of Health, Public Health Laboratory | Lisa Wingerter et al |
| B.1.1.253 | EPI_ISL_580650 | England/ALDP-9EDBD2/2020 | 2020/5/9 | Wellcome Sanger Institute for the COVID-19 Genomics UK (COG-UK) consortium | Jacquelyn Wynn et al |
| B.1.1.254 | EPI_ISL_641552 | France/OCC-SC417/2020 | 2020/3/14 | CNR Virus des Infections Respiratoires - France SUD | Antonin Bal et al |
| B.1.1.255 | EPI_ISL_473392 | England/BIRM-5FC59/2020 | 2020/5/21 | COVID-19 Genomics UK (COG-UK) Consortium | Institute of Microbiology et al |
| B.1.1.256 | EPI_ISL_441892 | England/NOTT-110A95/2020 | 2020/4/30 | COVID-19 Genomics UK (COG-UK) Consortium | Gemma Clark et al |
| B.1.1.257 | EPI_ISL_414444 | Netherlands/ZuidHolland_1/2020 | 2020/3/2 | Erasmus Medical Center | David Nieuwenhuijse et al |
| B.1.1.258 | EPI_ISL_545303 | USA/TX-HMH-2422/2020 | 2020/5/26 | Houston Methodist Hospital | S. Wesley Long et al |
| B.1.1.26 | EPI_ISL_434241 | USA/WA-S481/2020 | 2020/4/2 | Seattle Flu Study | Chu et al |
| B.1.1.261 | EPI_ISL_3085630 | Sweden/01_SE100_21CS504527/2020 | 2020/3/31 | Karolinska University Hospital | Jan Albert et al |
| B.1.1.262 | EPI_ISL_473358 | England/BIRM-5F9BC/2020 | 2020/5/14 | COVID-19 Genomics UK (COG-UK) Consortium | Institute of Microbiology et al |
| B.1.1.263 | EPI_ISL_582627 | UnitedArabEmirates/skmc-35676/2020 | 2020/4/3 | Molecular/Surveillance lab Sheikh Khalifa Medical City | Amirtharaj Francis et al |
| B.1.1.265 | EPI_ISL_1166102 | Italy/LOM-Pavia-41147/2020 | 2020/4/11 | Molecular Virology Unit, Microbiology and Virology Depertment, Fondazione IRCCS Policlinico San Matteo, Pavia | Monica Tallarita et al |
| B.1.1.266 | EPI_ISL_541333 | CzechRepublic/NRL-6834/2020 | 2020/5/23 | State Veterinary Institute Prague | Nagy et al |
| B.1.1.267 | EPI_ISL_596647 | Ireland/D-SVUH-R5315/2020 | 2020/3/24 | St.Vincent's University Hospital | Mary Lucey et al |

| | | | | | |
|---|---|---|---|---|---|
| B.1.1.268 | EPI_ISL_545446 | USA/TX-HMH-2639/2020 | 2020/6/5 | Houston Methodist Hospital | S. Wesley Long et al |
| B.1.1.269 | EPI_ISL_491226 | Portugal/IGC4089/2020 | 2020/6/1 | Instituto Gulbenkian de Ciência | Cathy Paulino et al |
| B.1.1.27 | EPI_ISL_457988 | Oman/RESP-20-4153/2020 | 2020/3/21 | Oman-NIC | Samira Al-Maruqi et al |
| B.1.1.270 | EPI_ISL_440000 | England/BRIS-121D98/2020 | 2020/3/21 | Wellcome Sanger Institute for the COVID-19 Genomics UK (COG-UK) consortium | Stephanie Hutchings et al |
| B.1.1.271 | EPI_ISL_583597 | Austria/CeMM0512/2020 | 2020/3/11 | Bergthaler laboratory, CeMM Research Center for Molecular Medicine of the Austrian Academy of Sciences | Alexandra Popa et al |
| B.1.1.272 | EPI_ISL_615213 | Denmark/DCGC-5282/2020 | 2020/8/17 | Albertsen lab, Department of Chemistry and Bioscience, Aalborg University, Denmark | Danish Covid-19 Genome Consortia et al |
| B.1.1.273 | EPI_ISL_647986 | USA/IN-CDC-3778675-001/2020 | 2020/4/23 | Pathogen Discovery, Respiratory Viruses Branch, Division of Viral Diseases, Centers for Disease Control and Prevention | Yan Li et al |
| B.1.1.274 | EPI_ISL_442562 | England/CAMB-806AC/2020 | 2020/4/10 | Wellcome Sanger Institute for the COVID-19 Genomics UK (COG-UK) consortium | Luke W Meredith et al |
| B.1.1.275 | EPI_ISL_483968 | England/GSTT-265D4DF/2020 | 2020/3/19 | COVID-19 Genomics UK (COG-UK) Consortium | Chloe Fisher et al |
| B.1.1.277 | EPI_ISL_420289 | England/SHEF-C0619/2020 | 2020/3/17 | Department of Infection, Immunity and Cardiovascular Disease, The Florey Institute, The Medical School, University of Sheffield | Thushan de Silva et al |
| B.1.1.279 | EPI_ISL_457989 | Oman/RESP-20-4252/2020 | 2020/3/21 | Oman-NIC | Samira Al-Maruqi et al |
| B.1.1.28 | EPI_ISL_416036 | Brazil/SP-14/2020 | 2020/3/5 | Instituto Adolfo Lutz, Interdiciplinary Procedures Center, Strategic Laboratory | Claudio Tavares Sacchi et al |
| B.1.1.280 | EPI_ISL_636843 | Lithuania/MR-LUHS-Eilnr201/2020 | 2020/7/30 | Lithuanian University of Health Sciences, Molecular cardiology lab. | Lukas Zemaitis et al |
| B.1.1.282 | EPI_ISL_460948 | Netherlands/Gelderland_48/2020 | 2020/3/26 | Erasmus Medical Center | Bas Oude Munnink et al |
| B.1.1.283 | EPI_ISL_479894 | Japan/PG-0189/2020 | 2020/3/16 | Pathogen Genomics Center, National Institute of Infectious Diseases | Tsuyoshi Sekizuka et al |
| B.1.1.284 | EPI_ISL_692598 | Japan/PG-4371/2020 | 2020/5/22 | Pathogen Genomics Center, National Institute of Infectious Diseases | Tsuyoshi Sekizuka et al |
| B.1.1.285 | EPI_ISL_2671842 | Japan/20200409-129/2020 | 2020/1/9 | Juntendo University Hospital | Koji Tuchiya et al |

| | | | | | |
|---|---|---|---|---|---|
| B.1.1.286 | EPI_ISL_2883698 | Canada/SK-NML-686/2020 | 2020/6/4 | National Microbiology Laboratory (NML) | Anna Majer et al |
| B.1.1.289 | EPI_ISL_420926 | Wales/PHWC-24F09/2020 | 2020/3/20 | Public Health Wales Microbiology Cardiff | Catherine Moore et al |
| B.1.1.29 | EPI_ISL_446035 | Wales/PHWC-2DEB8/2020 | 2020/4/8 | Public Health Wales Microbiology Cardiff | Catherine Moore et al |
| B.1.1.290 | EPI_ISL_495690 | USA/WA-S1240/2020 | 2020/5/11 | Seattle Flu Study | Deborah A. Nickerson et al |
| B.1.1.291 | EPI_ISL_2233972 | USA/CA-CDPH383/2020 | 2020/3/9 | California Department of Public Health | CDPH IDLB COVIDNet et al |
| B.1.1.294 | EPI_ISL_462150 | Russia/CRIE162784/2020 | 2020/3/31 | Group of Genomics and Postgenomic Technologies of Central Research Institute of Epidemilology | Speranskaya AS et al |
| B.1.1.296 | EPI_ISL_512410 | England/PORT-2D2041/2020 | 2020/3/20 | COVID-19 Genomics UK (COG-UK) Consortium | Angela Beckett et al |
| B.1.1.297 | EPI_ISL_515061 | Belgium/ULG-10254/2020 | 2020/7/16 | GIGA Medical Genomics | Keith Durkin et al |
| B.1.1.298 | EPI_ISL_618034 | Denmark/DCGC-8476/2020 | 2020/6/8 | Albertsen lab, Department of Chemistry and Bioscience, Aalborg University, Denmark | Danish Covid-19 Genome Consortia et al |
| B.1.1.299 | EPI_ISL_559410 | England/ALDP-4A29A7/2020 | 2020/5/22 | Wellcome Sanger Institute for the COVID-19 Genomics UK (COG-UK) consortium | The Lighthouse Lab in Alderley Park et al |
| B.1.1.3 | EPI_ISL_440368 | England/CAMB-75895/2020 | 2020/3/18 | Wellcome Sanger Institute for the COVID-19 Genomics UK (COG-UK) consortium | Luke W Meredith et al |
| B.1.1.30 | EPI_ISL_560392 | Lithuania/C20-05-R11/2020 | 2020/5/18 | Institute of Biotechnology, Life Sciences Center, Vilnius University and Thermo Fisher Scientific | Justinas Slikas et al |
| B.1.1.301 | EPI_ISL_559630 | England/ALDP-49F5B8/2020 | 2020/5/22 | Wellcome Sanger Institute for the COVID-19 Genomics UK (COG-UK) consortium | The Lighthouse Lab in Alderley Park et al |
| B.1.1.302 | EPI_ISL_534241 | Sweden/20-52386/2020 | 2020/7/6 | The Public Health Agency of Sweden | Anna-Malin Linde et al |
| B.1.1.304 | EPI_ISL_439524 | England/CAMB-77990/2020 | 2020/3/31 | Wellcome Sanger Institute for the COVID-19 Genomics UK (COG-UK) consortium | Luke W Meredith et al |
| B.1.1.305 | EPI_ISL_684600 | Japan/PG-0990/2020 | 2020/4/23 | Pathogen Genomics Center, National Institute of Infectious Diseases | Tsuyoshi Sekizuka et al |
| B.1.1.306 | EPI_ISL_2758212 | India/un-IRSHA-CD210761/2020 | 2020/3/1 | Communicable Diseases, Interactive Research School for Health Affairs (IRSHA) | Shrivastava et al |

| | | | | | |
|---|---|---|---|---|---|
| B.1.1.307 | EPI_ISL_532583 | England/QEUH-94334B/2020 | 2020/7/29 | Wellcome Sanger Institute for the COVID-19 Genomics UK (COG-UK) Consortium | Harper VanSteenhouse et al |
| B.1.1.308 | EPI_ISL_549450 | England/MILK-9AB648/2020 | 2020/8/31 | COVID-19 Genomics UK (COG-UK) Consortium | Tanya Golubchik et al |
| B.1.1.31 | EPI_ISL_569768 | Russia/OMS-ORINFI-5350S/2020 | 2020/6/7 | WHO National Influenza Centre Russian Federation | Artem Fadeev et al |
| B.1.1.310 | EPI_ISL_556972 | England/CAMC-945051/2020 | 2020/7/15 | Wellcome Sanger Institute for the COVID-19 Genomics UK (COG-UK) consortium | Rob Howes et al |
| B.1.1.311 | EPI_ISL_536972 | England/QEUH-9613CF/2020 | 2020/7/29 | Wellcome Sanger Institute for the COVID-19 Genomics UK (COG-UK) consortium | Harper VanSteenhouse et al |
| B.1.1.312 | EPI_ISL_755232 | Jordan/SEARCH-5424/2020 | 2020/8/13 | Andersen lab at Scripps Research | Issa Abu-Dayyeh et al |
| B.1.1.315 | EPI_ISL_859676 | UnitedArabEmirates/2890/2020 | 2020/8/20 | BTC, Khalifa University | Al Safar et al |
| B.1.1.316 | EPI_ISL_1011533 | USA/MA-MGH-03932/2020 | 2020/5/17 | Infectious Disease Program, Broad Institute of Harvard and MIT | Lemieux et al |
| B.1.1.317 | EPI_ISL_455703 | Vietnam/VNHN_3085/2020 | 2020/3/27 | Oxford University Clinical Research Unit, Hanoi, Vietnam | Nguyen Thi Tam et al |
| B.1.1.318 | EPI_ISL_977560 | Nigeria/CV702/2021 | 2021/1/7 | African Centre of Excellence for Genomics of Infectious Diseases (ACEGID), Redeemer’s University | Olawoye I. B. et al |
| B.1.1.318 | EPI_ISL_2166882 | Canada/AB-ABPHL-08599/2021 | 2021/2/21 | Public Health Agency of Canada (PHAC) National Microbiology Laboratory | Buss et al |
| B.1.1.319 | EPI_ISL_417427 | Belgium/VI-03027/2020 | 2020/3/2 | KU Leuven, Clinical and Epidemiological Virology | Joan Marti-Carerras et al |
| B.1.1.320 | EPI_ISL_542068 | USA/NMDOH-2020321601/2020 | 2020/9/1 | New Mexico Department of Health Scientific Laboratory | Ellie Johnson et al |
| B.1.1.322 | EPI_ISL_1301488 | Mexico/CMX-INER-IBT-38/2020 | 2020/5/7 | Instituto de Biotecnología de la UNAM | Authors from IBT et al |
| B.1.1.323 | EPI_ISL_417688 | Iceland/12/2020 | 2020/3/1 | deCODE genetics | Daniel F Gudbjartsson et al |
| B.1.1.324 | EPI_ISL_746784 | Chile/RM-72850/2020 | 2020/5/22 | Instituto de Salud Publica de Chile | Javier Tognarelli et al |
| B.1.1.325 | EPI_ISL_557211 | England/MILK-8274D2/2020 | 2020/7/16 | Wellcome Sanger Institute for the COVID-19 Genomics UK (COG-UK) consortium | The Lighthouse Lab in Milton Keynes et al |
| B.1.1.326 | EPI_ISL_826805 | Ecuador/EC-G-2565/2020 | 2020/3/23 | Instituto de Salud Publica de Chile | Javier Tognarelli et al |
| B.1.1.327 | EPI_ISL_721950 | Switzerland/ZH-ETHZ-380368/2020 | 2020/11/22 | Department of Biosystems Science and Engineering, ETH Zürich | Christian Beisel et al |

| | | | | | |
|---|---|---|---|---|---|
| B.1.1.328 | EPI_ISL_512662 | CostaRica/INC-0056/2020 | 2020/6/12 | Inciensa, Instituto Costarricense de Investigación y Enseñanza en Nutrición y Salud | Francisco Duarte et al |
| B.1.1.329 | EPI_ISL_1301696 | Mexico/ROO-InDRE-IBT-11/2020 | 2020/6/2 | Instituto de Biotecnología de la UNAM | Authors from IBT et al |
| B.1.1.33 | EPI_ISL_457796 | USA/DC-HP00052/2020 | 2020/3/7 | Johns Hopkins Hospital Department of Pathology | Peter M. Thielen et al |
| B.1.1.33 | EPI_ISL_2017478 | Brazil/GO-HLAGYN-814562/2020 | 2020/6/19 | HLAGYN - Laboratorio de Imunologia de Transplantes de Goias | Fernando Antonio Vinhal dos Santos et al |
| B.1.1.330 | EPI_ISL_796017 | Hebei/IVDC-03-05/2021 | 2021/1/2 | Hebei Provincial Center for Disease Control and Prevention, Shijiazhuang, Hebei Province; National Institute for Viral Disease Control and Prevention, China CDC | Shunxiang Qi et al |
| B.1.1.331 | EPI_ISL_419656 | Austria/CeMM0003/2020 | 2020/2/26 | Bergthaler laboratory, CeMM Research Center for Molecular Medicine of the Austrian Academy of Sciences | Alexandra Popa et al |
| B.1.1.332 | EPI_ISL_735423 | Brazil/SP-783/2020 | 2020/7/1 | Instituto Adolfo Lutz, Interdiciplinary Procedures Center, Strategic Laboratory | Claudio Tavares Sacchi et al |
| B.1.1.333 | EPI_ISL_2426018 | Norway/7651/2020 | 2020/1/29 | Norwegian Institute of Public Health, Department of Virology | Kathrine Stene-Johansen et al |
| B.1.1.334 | EPI_ISL_779331 | USA/WI-UW-2499/2020 | 2020/12/30 | University of Wisconsin-Madison AIDS Vaccine Research Laboratories | Gage Moreno et al |
| B.1.1.335 | EPI_ISL_2869911 | Germany/NW-KRO-446/2020 | 2020/4/3 | Center of Medical Microbiology, Virology, and Hospital Hygiene, University of Duesseldorf | Dennis Deschka et al |
| B.1.1.336 | EPI_ISL_733289 | Russia/KDA-RII-MH4936S/2020 | 2020/10/14 | WHO National Influenza Centre Russian Federation | Andrey Komissarov et al |
| B.1.1.337 | EPI_ISL_570516 | USA/WA-UW-12364/2020 | 2020/6/27 | UW Virology Lab | Pavitra Roychoudhury et al |
| B.1.1.338 | EPI_ISL_1195353 | Germany/BW-FR0586/2020 | 2020/12/2 | Institute of Virology, Clinial Virus Genomics, Medical Center, University of Freiburg, Freiburg, Germany | Jonas Fuchs et al |
| B.1.1.339 | EPI_ISL_1390779 | Canada/QC-CHUM-2007902694A/2020 | 2020/3/19 | Laboratoire de santé publique du Québec | Sandrine Moreira et al |
| B.1.1.34 | EPI_ISL_3067991 | SouthAfrica/Tygerberg_66/2020 | 2020/4/16 | Division of Medical Virology, Stellenbosch University and NHLS Tygerberg Hospital | Susan Engelbrecht et al |
| B.1.1.340 | EPI_ISL_591365 | Japan/IC-0020/2020 | 2020/4/6 | Pathogen Genomics Center, National Institute of Infectious Diseases | Tsuyoshi Sekizuka et al |

| | | | | | |
|---|---|---|---|---|---|
| B.1.1.341 | EPI_ISL_589828 | England/QEUH-9D0F03/2020 | 2020/9/14 | Wellcome Sanger Institute for the COVID-19 Genomics UK (COG-UK) consortium | Harper VanSteenhouse et al |
| B.1.1.342 | EPI_ISL_543456 | USA/TX-HMH-4243/2020 | 2020/6/19 | Houston Methodist Hospital | S. Wesley Long et al |
| B.1.1.343 | EPI_ISL_1739654 | Canada/ON-SC2318/2020 | 2020/11/9 | McMaster University | Allison McGeer et al |
| B.1.1.344 | EPI_ISL_1301682 | Mexico/AGU-InDRE-IBT-43/2020 | 2020/4/9 | Instituto de Biotecnología de la UNAM | Authors from IBT et al |
| B.1.1.345 | EPI_ISL_1009196 | USA/MS-UT-1751/2020 | 2020/11/10 | Colleen B. Jonsson | Mariah K. Taylor et al |
| B.1.1.346 | EPI_ISL_621092 | Denmark/DCGC-5983/2020 | 2020/9/21 | Albertsen lab, Department of Chemistry and Bioscience, Aalborg University, Denmark | Danish Covid-19 Genome Consortia et al |
| B.1.1.347 | EPI_ISL_5917671 | Romania/DJ13378/2020 | 2020/7/14 | Medical Genetics Laboratory, Regional Centre of Medical Genetics, Emergency County Hospital Craiova | Anca-Lelia (Riza) Costache et al |
| B.1.1.348 | EPI_ISL_941965 | Colombia/GUR-06575/2020 | 2020/4/30 | Centro de Investigaciones en Microbiología y Biotecnología-UR (CIMBIUR), Facultad de Ciencias Naturales, Universidad del Rosario, Bogotá, Colombia Instituto Nacional de Salud, Bogotá, Colombia Icahn School of Medicine at Mount Sinai, New York, USA | Luz Helena Patiño et al |
| B.1.1.349 | EPI_ISL_524492 | England/BRIS-133838/2020 | 2020/8/4 | Wellcome Sanger Institute for the COVID-19 Genomics UK (COG-UK) consortium | Stephanie Hutchings et al |
| B.1.1.350 | EPI_ISL_437094 | Latvia/015/2020 | 2020/3/23 | Latvian Biomedical Research and Study Centre | Ivars Silamiķelis et al |
| B.1.1.351 | EPI_ISL_567220 | England/QEUH-9C89E2/2020 | 2020/9/10 | Wellcome Sanger Institute for the COVID-19 Genomics UK (COG-UK) consortium | Harper VanSteenhouse et al |
| B.1.1.352 | EPI_ISL_2919826 | USA/AZ-TG955539/2020 | 2020/9/29 | TGen North | "Jolene Bowers et al |
| B.1.1.353 | EPI_ISL_593741 | Australia/NSW1127/2020 | 2020/9/8 | NSW Health Pathology - Institute of Clinical Pathology and Medical Research; Westmead Hospital; University of Sydney | CIDM-PH et al. |
| B.1.1.354 | EPI_ISL_2501532 | India/AS-DIB_10072/2020 | 2020/5/19 | Regional VRDL,Dibrugarh | B Borkakoty et al |
| B.1.1.355 | EPI_ISL_1225757 | USA/IL-IDPH-OGL-S-000928/2020 | 2020/6/2 | Gagnon Lab, Southern Illinois University | Keith Gagnon et al |

| | | | | | |
|---|---|---|---|---|---|
| B.1.1.356 | EPI_ISL_586426 | Canada/ON-S67/2020 | 2020/3/19 | McMaster University | Allison McGeer et al |
| B.1.1.358 | EPI_ISL_2006728 | USA/CA-9200066222/2020 | 2020/9/13 | Moderna Inc. | Yamuna Paila et al |
| B.1.1.359 | EPI_ISL_622565 | Denmark/DCGC-1828/2020 | 2020/7/6 | Albertsen lab, Department of Chemistry and Bioscience, Aalborg University, Denmark | Danish Covid-19 Genome Consortia et al |
| B.1.1.360 | EPI_ISL_474308 | Wales/PHWC-2C314/2020 | 2020/4/3 | COVID-19 Genomics UK (COG-UK) Consortium | Catherine Moore et al |
| B.1.1.361 | EPI_ISL_559283 | England/MILK-4B620F/2020 | 2020/6/4 | Wellcome Sanger Institute for the COVID-19 Genomics UK (COG-UK) consortium | The Lighthouse Lab in Milton Keynes et al |
| B.1.1.362 | EPI_ISL_543882 | USA/TX-HMH-2946/2020 | 2020/6/9 | Houston Methodist Hospital | S. Wesley Long et al |
| B.1.1.363 | EPI_ISL_432211 | Wales/PHWC-2764D/2020 | 2020/3/30 | Public Health Wales Microbiology Cardiff | Catherine Moore et al |
| B.1.1.364 | EPI_ISL_688814 | Japan/PG-5690/2020 | 2020/5/22 | Pathogen Genomics Center, National Institute of Infectious Diseases | Tsuyoshi Sekizuka et al |
| B.1.1.365 | EPI_ISL_558929 | England/ALDP-52B8A8/2020 | 2020/6/1 | Wellcome Sanger Institute for the COVID-19 Genomics UK (COG-UK) consortium | The Lighthouse Lab in Alderley Park et al |
| B.1.1.366 | EPI_ISL_435124 | UnitedArabEmirates/L0879/2020 | 2020/3/16 | Al Jalila Genomics Center | Ahmad Abou Tayoun et al |
| B.1.1.367 | EPI_ISL_544022 | USA/TX-HMH-3143/2020 | 2020/6/11 | Houston Methodist Hospital | S. Wesley Long et al |
| B.1.1.368 | EPI_ISL_5926948 | Guatemala/ASI1123/2020 | 2020/6/30 | Asociación de Salud Integral / Clínica Familiar "Luis Ángel García" | Eduardo Arathon et al |
| B.1.1.369 | EPI_ISL_500085 | England/LIVE-A4D52/2020 | 2020/3/2 | COVID-19 Genomics UK (COG-UK) Consortium | Sam Haldenby et al |
| B.1.1.37 | EPI_ISL_2443745 | USA/CA-UCI-040/2020 | 2020/4/5 | University of California, Irvine | Amanda N. Pinski et al |
| B.1.1.370 | EPI_ISL_427322 | Russia/StPetersburg-RII4655S/2020 | 2020/4/5 | WHO National Influenza Centre Russian Federation | Andrey Komissarov et al |
| B.1.1.371 | EPI_ISL_420451 | Belgium/HC-030760/2020 | 2020/3/7 | KU Leuven, Clinical and Epidemiological Virology | Joan Marti-Carreras et al |
| B.1.1.372 | EPI_ISL_500072 | England/LIVE-A4B0D/2020 | 2020/3/8 | COVID-19 Genomics UK (COG-UK) Consortium | Sam Haldenby et al |
| B.1.1.373 | EPI_ISL_1400558 | Russia/PRI-RII-MH820S/2020 | 2020/8/7 | WHO National Influenza Centre Russian Federation | Andrey Komissarov et al |
| B.1.1.375 | EPI_ISL_629391 | England/MILK-AC841D/2020 | 2020/10/21 | Wellcome Sanger Institute for the COVID-19 Genomics UK (COG-UK) consortium | The Lighthouse Lab in Milton Keynes et al |
| B.1.1.376 | EPI_ISL_672244 | USA/CA-CZB-10172/2020 | 2020/5/11 | Chan-Zuckerberg Biohub | CZB Cliahub Consortium et al |

| | | | | | |
|---|---|---|---|---|---|
| B.1.1.379 | EPI_ISL_442171 | England/CAMB-7CC38/2020 | 2020/4/5 | Wellcome Sanger Institute for the COVID-19 Genomics UK (COG-UK) consortium | Luke W Meredith et al |
| B.1.1.38 | EPI_ISL_490740 | Wales/PHWC-16590D/2020 | 2020/5/25 | COVID-19 Genomics UK (COG-UK) Consortium | Catherine Moore et al |
| B.1.1.380 | EPI_ISL_635381 | USA/CA-ALSR-2190/2020 | 2020/6/12 | Andersen lab at Scripps Research | SEARCH Alliance San Diego with Tracy Basler et al |
| B.1.1.381 | EPI_ISL_536515 | Peru/LIM-INS-117/2020 | 2020/5/4 | Laboratorio de Infecciones Respiratorias Agudas | Eduardo Juscamayta Lopez et al |
| B.1.1.382 | EPI_ISL_3068004 | SouthAfrica/Tygerberg_83/2020 | 2020/4/19 | Division of Medical Virology, Stellenbosch University and NHLS Tygerberg Hospital | Susan Engelbrecht et al |
| B.1.1.383 | EPI_ISL_467477 | SouthAfrica/KRISP-0125/2020 | 2020/5/13 | KRISP, KZN Research Innovation and Sequencing Platform | Giandhari J et al |
| B.1.1.384 | EPI_ISL_847890 | Switzerland/GE-32815676/2020 | 2020/12/28 | HUG, Laboratory of Virology and the Health2030 Genome Center | Samuel Cordey et al |
| B.1.1.385 | EPI_ISL_666647 | Germany/NW-HHU-360/2020 | 2020/9/11 | Center of Medical Microbiology, Virology, and Hospital Hygiene, University of Duesseldorf | Maximilian Damagnez et al |
| B.1.1.386 | EPI_ISL_467498 | SouthAfrica/KRISP-0148/2020 | 2020/5/30 | KRISP, KZN Research Innovation and Sequencing Platform | Giandhari J et al |
| B.1.1.387 | EPI_ISL_733167 | Russia/AST-RII-MH4130S/2020 | 2020/8/4 | WHO National Influenza Centre Russian Federation | Andrey Komissarov et al |
| B.1.1.388 | EPI_ISL_456154 | Colombia/NAR-INS-103303/2020 | 2020/4/26 | Instituto Nacional de Salud, Universidad Cooperativa de Colombia, Instituto Alexander von Humboldt, Imperial College-London, London School of Hygiene & Tropical Medicine | Katherine Laiton-Donato et al |
| B.1.1.389 | EPI_ISL_512665 | CostaRica/INC-0059/2020 | 2020/6/18 | Inciensa, Instituto Costarricense de Investigación y Enseñanza en Nutrición y Salud | Francisco Duarte et al |
| B.1.1.39 | EPI_ISL_751326 | Italy/VEN-UniVR-85/2020 | 2020/3/23 | University of Verona, Department of Biotechnology | Antonio Mori et al |
| B.1.1.391 | EPI_ISL_520724 | UnitedArabEmirates/L920633058/2020 | 2020/3/30 | Al Jalila Genomics Center | Ahmad Abou Tayoun et al |
| B.1.1.392 | EPI_ISL_614060 | USA/VA-DCLS-1809/2020 | 2020/9/16 | Virginia DCLS | Virginia DCLS et al |
| B.1.1.393 | EPI_ISL_1544191 | India/GJ-GBRC518/2020 | 2020/11/24 | Gujarat Biotechnology Research Centre | Ramesh Pandit et al |

| | | | | | |
|---|---|---|---|---|---|
| B.1.1.394 | EPI_ISL_693616 | Portugal/PT1614/2020 | 2020/7/8 | Instituto Nacional de Saude (INSA) | Borges et al |
| B.1.1.395 | EPI_ISL_488032 | England/NORT-2972D9/2020 | 2020/4/27 | Wellcome Sanger Institute for the COVID-19 Genomics UK (COG-UK) consortium | Chris Duncan et al |
| B.1.1.396 | EPI_ISL_569793 | Russia/OMS-ORINFI-9295S/2020 | 2020/7/25 | WHO National Influenza Centre Russian Federation | Artem Fadeev et al |
| B.1.1.397 | EPI_ISL_596287 | Russia/SPE-RII-MH1516S/2020 | 2020/9/11 | WHO National Influenza Centre Russian Federation | Andrey Komissarov et al |
| B.1.1.398 | EPI_ISL_425246 | England/CAMB-7386A/2020 | 2020/3/31 | COVID-19 Genomics UK (COG-UK) Consortium | Luke W Meredith et al |
| B.1.1.399 | EPI_ISL_767807 | Ireland/LH-NVRL-MPZ41611/2020 | 2020/4/15 | Irish Coronavirus Sequencing Consortium - Teagasc Moorepark | Calum Walsh et al |
| B.1.1.4 | EPI_ISL_438391 | England/CAMB-71DA9/2020 | 2020/3/31 | Wellcome Sanger Institute for the COVID-19 Genomics UK (COG-UK) consortium | Luke W Meredith et al |
| B.1.1.40 | EPI_ISL_640080 | SouthAfrica/NHLS-UCT-GS-2440/2020 | 2020/4/24 | NHLS/UCT | Arash Iranzadeh et al |
| B.1.1.400 | EPI_ISL_3730993 | Italy/TAA-APSS-Ph2_RUN3_NB02/2020 | 2020/3/29 | U.O. Microbiologia e Virologia, Azienda Provinciale per i Servizi Sanitari Provincia Autonoma di Trento, Ospedale S.Chiara | Mirko Moser et al |
| B.1.1.401 | EPI_ISL_491225 | Portugal/IGC4033/2020 | 2020/6/1 | Instituto Gulbenkian de Ciência | Cathy Paulino et al |
| B.1.1.402 | EPI_ISL_542813 | USA/TX-HMH-1750/2020 | 2020/5/12 | Houston Methodist Hospital | S. Wesley Long et al |
| B.1.1.403 | EPI_ISL_635806 | USA/CA-ALSR-3037/2020 | 2020/9/3 | Andersen lab at Scripps Research | SEARCH Alliance San Diego with Tracy Basler et al |
| B.1.1.404 | EPI_ISL_660477 | BurkinaFaso/608/2020 | 2020/6/21 | Centre Muraz | Abdoul-Salam Ouedraogo et al |
| B.1.1.405 | EPI_ISL_815356 | Germany/SH-Cento-37955598/2020 | 2020/9/14 | Centogene | Peter Bauer et al |
| B.1.1.406 | EPI_ISL_710490 | Spain/AS-HUCA-232023718/2020 | 2020/3/25 | Virology (microbiology), Hospital Universitario Central de Asturias | Castello et al |
| B.1.1.407 | EPI_ISL_524045 | Russia/SPE-RII-22478V/2020 | 2020/6/9 | WHO National Influenza Centre Russian Federation | Andrey Komissarov et al |
| B.1.1.408 | EPI_ISL_529663 | England/HECH-259A4F5/2020 | 2020/7/11 | COVID-19 Genomics UK (COG-UK) Consortium | Institute of Microbiology et al |
| B.1.1.409 | EPI_ISL_516850 | England/LOND-126092C/2020 | 2020/3/16 | Wellcome Sanger Institute for the COVID-19 Genomics UK (COG-UK) consortium | Ling Li et al |

| | | | | | |
|---|---|---|---|---|---|
| B.1.1.409 | EPI_ISL_1503349 | USA/TX-DSHS-4988/2020 | 2020/7/1 | Texas Department of State Health Services (TXDSHS) | Rashmi Tuladhar et al |
| B.1.1.41 | EPI_ISL_440297 | England/CAMB-73141/2020 | 2020/3/21 | Wellcome Sanger Institute for the COVID-19 Genomics UK (COG-UK) consortium | Luke W Meredith et al |
| B.1.1.410 | EPI_ISL_491227 | Portugal/IGC4092/2020 | 2020/6/1 | Instituto Gulbenkian de Ciência | Cathy Paulino et al |
| B.1.1.411 | EPI_ISL_574294 | Ecuador/2479/2020 | 2020/3/23 | Instituto Nacional de Investigación en Salud Pública | Leandro Patino Patino et al |
| B.1.1.412 | EPI_ISL_515832 | SouthAfrica/KRISP-K001347/2020 | 2020/7/1 | KRISP, KZN Research Innovation and Sequencing Platform | Giandhari J et al |
| B.1.1.413 | EPI_ISL_566603 | England/MILK-9C23DB/2020 | 2020/9/7 | Wellcome Sanger Institute for the COVID-19 Genomics UK (COG-UK) consortium | The Lighthouse Lab in Milton Keynes et al |
| B.1.1.414 | EPI_ISL_699946 | India/MH-ACTREC-294/2020 | 2020/7/14 | Hematopathology Laboratory, ACTREC, TMC | Hematopathology Laboratory et al |
| B.1.1.415 | EPI_ISL_854144 | Austria/CeMM1772/2020 | 2020/11/28 | Bergthaler laboratory, CeMM Research Center for Molecular Medicine of the Austrian Academy of Sciences | Lukas Endler et al |
| B.1.1.416 | EPI_ISL_419887 | Australia/VIC191/2020 | 2020/3/19 | Victorian Infectious Diseases Reference Laboratory and Microbiological Diagnostic Unit Public Health Laboratory, Doherty Institute | Caly L. et al |
| B.1.1.417 | EPI_ISL_570003 | Canada/ON-UHTC-0277/2020 | 2020/6/23 | Ontario Institute for Cancer Research | Ramzi Fattouh et al |
| B.1.1.419 | EPI_ISL_488826 | England/NORT-281418/2020 | 2020/3/14 | Wellcome Sanger Institute for the COVID-19 Genomics UK (COG-UK) consortium | Chris Duncan et al |
| B.1.1.420 | EPI_ISL_3152081 | Senegal/SC20-5366/2020 | 2020/8/2 | LBV Le Dantec | Halimatou Diop Ndiaye et al |
| B.1.1.421 | EPI_ISL_693559 | Portugal/PT1546/2020 | 2020/6/20 | Instituto Nacional de Saude (INSA) | Borges et al |
| B.1.1.422 | EPI_ISL_860817 | Belarus/ChVir21889/2020 | 2020/10/26 | Charité Universitätsmedizin Berlin, Institut für Virologie | Victor M Corman et al |
| B.1.1.423 | EPI_ISL_513872 | USA/CA-CZB-2467/2020 | 2020/5/19 | Chan-Zuckerberg Biohub | CZB Cliahub Consortium et al |
| B.1.1.424 | EPI_ISL_678283 | Russia/SAR-RARI-3573/2020 | 2020/5/2 | Mikrobiologie, RARI | Krasnov et al |
| B.1.1.425 | EPI_ISL_552330 | England/MILK-994E59/2020 | 2020/8/25 | Wellcome Sanger Institute for the COVID-19 Genomics UK (COG-UK) consortium | The Lighthouse Lab in Milton Keynes et al |
| B.1.1.426 | EPI_ISL_2773937 | USA/LA-UW-5500544/2020 | 2020/6/16 | UW Virology Lab | Pavitra Roychoudhury et al |

| | | | | | |
|---|---|---|---|---|---|
| B.1.1.427 | EPI_ISL_732995 | Russia/SVE-RII-MH216S/2020 | 2020/7/25 | WHO National Influenza Centre Russian Federation | Andrey Komissarov et al |
| B.1.1.428 | EPI_ISL_626060 | Denmark/DCGC-8747/2020 | 2020/10/26 | Albertsen lab, Department of Chemistry and Bioscience, Aalborg University, Denmark | Danish Covid-19 Genome Consortia et al |
| B.1.1.429 | EPI_ISL_770056 | Latvia/222/2020 | 2020/9/15 | Latvian Biomedical Research and Study Centre | Ivars Silamiķelis et al |
| B.1.1.43 | EPI_ISL_433327 | Scotland/CVR702/2020 | 2020/3/25 | COVID-19 Genomics UK (COG-UK) Consortium | Ana da Silva Filipe et al |
| B.1.1.431 | EPI_ISL_521499 | Australia/VIC2653/2020 | 2020/7/1 | VIDRL and MDU-PHL | Caly L. et al |
| B.1.1.432 | EPI_ISL_872097 | Mexico/VER-InDRE-104/2020 | 2020/4/24 | Instituto de Diagnostico y Referencia Epidemiologicos (INDRE) | Abril Rodriguez-Maldonado et al |
| B.1.1.433 | EPI_ISL_1498929 | Slovenia/66807/2020 | 2020/8/9 | Institute of Microbiology and Immunology, Faculty of Medicine, University of Ljubljana | Alen Suljič et al |
| B.1.1.434 | EPI_ISL_3098576 | USA/MD-HP07302-PIDOBBJPQM/2020 | 2020/6/1 | Johns Hopkins Hospital Department of Pathology | C. Paul Morris et al |
| B.1.1.435 | EPI_ISL_732663 | Russia/MOS-CRIE-13385391/2020 | 2020/11/6 | Group of Genomics and Postgenomic Technologies of Central Research Institute of Epidemilology | Samoilov AE et al |
| B.1.1.436 | EPI_ISL_420416 | Belgium/FMC-0323159/2020 | 2020/3/23 | KU Leuven, Clinical and Epidemiological Virology | Joan Marti-Carreras et al |
| B.1.1.437 | EPI_ISL_4418817 | USA/MD-HP09200-PIDKCKUCMM/2020 | 2020/5/14 | Johns Hopkins Hospital Department of Pathology | C. Paul Morris et al |
| B.1.1.438 | EPI_ISL_660384 | Sweden/20-22286/2020 | 2020/3/12 | The Public Health Agency of Sweden | Anna-Malin Linde et al |
| B.1.1.44 | EPI_ISL_491093 | CzechRepublic/NRL_4039/2020 | 2020/3/18 | The National Institute of Public Health and State Veterinary Institute Prague | Nagy et al |
| B.1.1.440 | EPI_ISL_522352 | USA/TX-DSHS-0066/2020 | 2020/5/9 | Texas Department of State Health Services | Rashmi Tuladhar et al |
| B.1.1.441 | EPI_ISL_631464 | USA/WI-WSLH-200707/2020 | 2020/6/26 | Wisconsin State Laboratory of Hygiene Communicable Disease Division | Kelsey R. Florek et al |
| B.1.1.442 | EPI_ISL_981038 | Argentina/PAIS-F0040/2020 | 2020/5/16 | Laboratorio Central Mg. Luis Alfredo Pianciola on behalf of 'Proyecto Argentino Interinstitucional de genomica de SARS-CoV-2' (PAIS Consortium) | Liliana Fonseca et al |
| B.1.1.444 | EPI_ISL_515762 | SouthAfrica/KRISP-K002398/2020 | 2020/7/27 | KRISP, KZN Research Innovation and Sequencing Platform | Giandhari J et al |

| | | | | | |
|---|---|---|---|---|---|
| B.1.1.445 | EPI_ISL_733497 | Switzerland/ZH-UZH-IMV127/2020 | 2020/12/14 | Institute of Medical Virology, University of Zurich | Stefan Schmutz et al |
| B.1.1.446 | EPI_ISL_2643656 | Estonia/Cov0223/2020 | 2020/9/14 | Department of Microbiology, Institute of Biomedicine and Translational Medicine, University of Tartu | Radko Avi et al |
| B.1.1.447 | EPI_ISL_516698 | USA/VA-DCLS-0824/2020 | 2020/5/5 | Virginia DCLS | Virginia DCLS et al |
| B.1.1.448 | EPI_ISL_2444003 | SouthAfrica/NICD-N00163/2020 | 2020/5/4 | National Institute for Communicable Diseases of the National Health Laboratory Service | Amoako DG et al |
| B.1.1.449 | EPI_ISL_596250 | Russia/SPE-RII-MH1007S/2020 | 2020/9/4 | WHO National Influenza Centre Russian Federation | Andrey Komissarov et al |
| B.1.1.45 | EPI_ISL_559611 | England/ALDP-49F3BE/2020 | 2020/5/22 | Wellcome Sanger Institute for the COVID-19 Genomics UK (COG-UK) consortium | The Lighthouse Lab in Alderley Park et al |
| B.1.1.450 | EPI_ISL_738746 | USA/CA-CZB-15072/2020 | 2020/11/20 | Chan-Zuckerberg Biohub | CZB Cliahub Consortium et al |
| B.1.1.451 | EPI_ISL_635143 | Norway/4276/2020 | 2020/10/10 | Norwegian Institute of Public Health, Department of Virology | Kathrine Stene-Johansen et al |
| B.1.1.452 | EPI_ISL_639949 | Russia/PER-RII-MH1647S/2020 | 2020/9/10 | WHO National Influenza Centre Russian Federation | Andrey Komissarov et al |
| B.1.1.453 | EPI_ISL_754963 | USA/CA-CDPH090/2020 | 2020/8/5 | California Department of Public Health | CDPH IDLB COVIDNet et al |
| B.1.1.456 | EPI_ISL_487319 | SouthAfrica/KRISP-0368/2020 | 2020/6/18 | KRISP, KZN Research Innovation and Sequencing Platform | Giandhari J et al |
| B.1.1.458 | EPI_ISL_469059 | Sweden/20-14647/2020 | 2020/5/14 | The Public Health Agency of Sweden | Oskar Karlsson Lindsjo et al |
| B.1.1.459 | EPI_ISL_640096 | SouthAfrica/NHLS-UCT-GP-5016/2020 | 2020/4/15 | NHLS/UCT | Arash Iranzadeh et al |
| B.1.1.46 | EPI_ISL_447587 | India/TN-CCMB-C4-15/2020 | 2020/4/20 | CSIR-Centre for Cellular and Molecular Biology | K Kaveri et al |
| B.1.1.462 | EPI_ISL_511770 | Portugal/PT1227/2020 | 2020/3/25 | Instituto Nacional de Saude (INSA) and Instituto Gulbenkian de Ciencia (IGC) | Borges et al |
| B.1.1.463 | EPI_ISL_672258 | USA/CA-CZB-10202/2020 | 2020/5/11 | Chan-Zuckerberg Biohub | CZB Cliahub Consortium et al |
| B.1.1.464 | EPI_ISL_2896976 | USA/WI-MCW-CA085/2020 | 2020/4/1 | Infectious Disease Core Research, Abbott Diagnostics Division | Forberg et al |
| B.1.1.465 | EPI_ISL_4418186 | Japan/CA001/2020 | 2020/4/21 | Department of Emerging Infectious Diseases, Institute of Tropical Medicine, Nagasaki University | Haruka Abe et al |
| B.1.1.466 | EPI_ISL_561105 | Gambia/GC19-2427/2020 | 2020/7/21 | MRCG at LSHTM Genomics lab | Abdul Karim sesay et al |

| | | | | | |
|---|---|---|---|---|---|
| B.1.1.467 | EPI_ISL_633056 | USA/NY-NYCPHL-001044/2020 | 2020/10/8 | New York City Public Health Laboratory | Jade Wang et al |
| B.1.1.47 | EPI_ISL_489977 | Switzerland/VD-ETHZ-180039/2020 | 2020/7/1 | Department of Biosystems Science and Engineering, ETH Zürich | Christian Beisel et al |
| B.1.1.48 | EPI_ISL_685524 | Japan/PG-1918/2020 | 2020/3/27 | Pathogen Genomics Center, National Institute of Infectious Diseases | Tsuyoshi Sekizuka et al |
| B.1.1.480 | EPI_ISL_695809 | USA/AZ-TG640888/2020 | 2020/4/13 | TGen North | Jolene Bowers et al |
| B.1.1.481 | EPI_ISL_733180 | Russia/AST-RII-MH4157S/2020 | 2020/8/18 | WHO National Influenza Centre Russian Federation | Andrey Komissarov et al |
| B.1.1.482 | EPI_ISL_459686 | Scotland/GCVR-17027D/2020 | 2020/3/24 | Wellcome Sanger Institute for the COVID-19 Genomics UK (COG-UK) consortium | Ana da Silva Filipe et al |
| B.1.1.483 | EPI_ISL_745217 | Belgium/UGent-332/2020 | 2020/12/11 | Onderzoeksgroep Virologie | Nick Vereecke et al |
| B.1.1.484 | EPI_ISL_729926 | Nigeria/NA-CV240/2020 | 2020/5/31 | African Centre of Excellence for Genomics of Infectious Diseases (ACEGID), Redeemer's University, Ede, Osun State, Nigeria | Oluniyi P.E. et al |
| B.1.1.485 | EPI_ISL_591210 | Canada/ON-S737/2020 | 2020/4/9 | McMaster University | Allison McGeer et al |
| B.1.1.486 | EPI_ISL_802165 | USA/NY-MSHSPSP-PV16551/2020 | 2020/8/13 | MSHS Pathogen Surveillance Program | Ana S. Gonzalez-Reiche et al |
| B.1.1.487 | EPI_ISL_654794 | Nigeria/NG57839/2020 | 2020/3/22 | Centre for Human Virology & Genomics, Nigerian Institute of Medical Research | Shaibu et al |
| B.1.1.49 | EPI_ISL_432384 | Wales/PHWC-27249/2020 | 2020/3/30 | Public Health Wales Microbiology Cardiff | Catherine Moore et al |
| B.1.1.5 | EPI_ISL_417551 | Iceland/187/2020 | 2020/3/16 | deCODE genetics | Daniel F Gudbjartsson et al |
| B.1.1.50 | EPI_ISL_447450 | Israel/13075703/2020 | 2020/3/29 | Stern Lab | Stern Lab et al |
| B.1.1.500 | EPI_ISL_470903 | Russia/Moscow_PMVL-10/2020 | 2020/4/10 | Pathogenic Microorganisms Variability Laboratory | Alexey Shchetinin et al |
| B.1.1.506 | EPI_ISL_569742 | Russia/OMS-ORINFI-7121S/2020 | 2020/5/14 | WHO National Influenza Centre Russian Federation | Artem Fadeev et al |
| B.1.1.51 | EPI_ISL_538900 | England/LEED-2AA3C1/2020 | 2020/3/30 | Wellcome Sanger Institute for the COVID-19 Genomics UK (COG-UK) consortium | Louissa Macfarlane-Smith et al |
| B.1.1.512 | EPI_ISL_635635 | USA/CA-ALSR-2692/2020 | 2020/7/21 | Andersen lab at Scripps Research | SEARCH Alliance San Diego with Tracy Basler et al |
| B.1.1.513 | EPI_ISL_525622 | USA/NY-Wadsworth-22513-01/2020 | 2020/3/16 | Wadsworth Center, New York State Department of Health | Kirsten St. George et al |
| B.1.1.514 | EPI_ISL_583187 | USA/CA-CZB-6113/2020 | 2020/5/1 | Chan-Zuckerberg Biohub | CZB Clianub Consortium et al |

| | | | | | |
|---|---|---|---|---|---|
| B.1.1.515 | EPI_ISL_428490 | Taiwan/78/2020 | 2020/3/16 | Centers for Disease Control, R.O.C. (Taiwan) | Ji-Rong Yang et al |
| B.1.1.516 | EPI_ISL_1517405 | CostaRica/HNN-0361/2020 | 2020/6/2 | Inciensa, Instituto Costarricense de Investigación y Enseñanza en Nutrición y Salud | Cristian Peréz-Corrales et al |
| B.1.1.517 | EPI_ISL_3536877 | USA/CA-CDPH1435/2020 | 2020/12/7 | California Department of Public Health | CDPH IDLB COVIDNet et al |
| B.1.1.518 | EPI_ISL_4219952 | Mexico/CMX-INMEGEN-04-07-31/2020 | 2020/6/5 | Instituto Nacional de Medicina Genomica | Cedro-Tanda A et al |
| B.1.1.519 | EPI_ISL_2222621 | USA/TX-HMH-MCoV-39891/2020 | 2020/8/6 | Houston Methodist Hospital | Randall J. Olsen et al |
| B.1.1.519 | EPI_ISL_3151148 | Mexico/YUC-NYGC-39037-20/2020 | 2020/8/28 | New York Genome Center | Michael Zody et al |
| B.1.1.519 | EPI_ISL_1841710 | Finland/2342/2021 | 2021/2/14 | Department of Virology, Faculty of Medicine, University of Helsinki, Helsinki, Finland | Teemu Smura et al |
| B.1.1.52 | EPI_ISL_2444005 | SouthAfrica/NICD-N00165/2020 | 2020/4/11 | National Institute for Communicable Diseases of the National Health Laboratory Service | Amoako DG et al |
| B.1.1.528 | EPI_ISL_745141 | SouthAfrica/Tygerberg-460/2020 | 2020/12/7 | National Health Laboratory Service (NHLS), Tygerberg | Susan Engelbrecht et al |
| B.1.1.529 | EPI_ISL_6825395 | SouthAfrica/CERI-KRISP-K032355/2021 | 2021/11/16 | CERI, Centre for Epidemic Response and Innvoation, Stellenbosch University and KRISP, KZN Research Innovation and Sequencing Platform, UKZN. | Amy Strydom et al |
| B.1.1.53 | EPI_ISL_490277 | SouthAfrica/R10427/2020 | 2020/6/4 | National Institute for Communicable Diseases of the National Health Laboratory Service | Allam M et al |
| B.1.1.54 | EPI_ISL_507973 | USA/MN-MDH-1386/2020 | 2020/3/19 | Minnesota Department of Health, Public Health Laboratory | Matt Plumb et al |
| B.1.1.55 | EPI_ISL_441501 | England/NOTT-110330/2020 | 2020/4/12 | COVID-19 Genomics UK (COG-UK) Consortium | Gemma Clark et al |
| B.1.1.56 | EPI_ISL_467493 | SouthAfrica/KRISP-0143/2020 | 2020/5/21 | KRISP, KZN Research Innovation and Sequencing Platform | Giandhari J et al |
| B.1.1.57 | EPI_ISL_455639 | SouthAfrica/KRISP-109/2020 | 2020/3/31 | KRISP, KZN Research Innovation and Sequencing Platform | Giandhari J et al |
| B.1.1.58 | EPI_ISL_1854982 | Italy/LAZ-AMC-2004088031-DS/2020 | 2020/4/8 | Virology Laboratory, Scientific Department, Army Medical Center | Silvia Fillo et al |
| B.1.1.59 | EPI_ISL_479385 | Wales/PHWC-164E0F/2020 | 2020/6/1 | COVID-19 Genomics UK (COG-UK) Consortium | Catherine Moore et al |

| | | | | | |
|---|---|---|---|---|---|
| B.1.1.61 | EPI_ISL_419557 | USA/GA-CDC-03063103-001/2020 | 2020/2/29 | Pathogen Discovery, Respiratory Viruses Branch, Division of Viral Diseases, Centers for Disease Control and Prevention | Ying Tao et al |
| B.1.1.62 | EPI_ISL_3068127 | SouthAfrica/Tygerberg_257/2020 | 2020/5/7 | Division of Medical Virology, Stellenbosch University and NHLS Tygerberg Hospital | Susan Engelbrecht et al |
| B.1.1.63 | EPI_ISL_3105881 | Philippines/PH-RITM-0074/2020 | 2020/6/23 | Research Institute for Tropical Medicine | Lei Lanna Dancel et al |
| B.1.1.67 | EPI_ISL_776018 | Germany/HH-hpi-p1516/2020 | 2020/3/15 | Heinrich Pette Institute, Leibniz Institute for Experimental Virology | Alexis Robitaille et al |
| B.1.1.7 | EPI_ISL_1603195 | Spain/CL-COV00781/2020 | 2020/2/7 | SARS-CoV-2 Sequencing Castilla y Leon-Spain Consortium | Antonio Orduña-Domingo et al |
| B.1.1.7 | EPI_ISL_1100034 | England/RAND-12E0265/2021 | 2021/2/10 | Wellcome Sanger Institute for the COVID-19 Genomics UK (COG-UK) Consortium | Randox Laboratories et al |
| B.1.1.70 | EPI_ISL_856906 | Italy/UMB-ISS-3867/2020 | 2020/3/6 | Virology Laboratory, Scientific Department, Army Medical Center | Paola Stefanelli et al |
| B.1.1.71 | EPI_ISL_419748 | Australia/VIC29/2020 | 2020/3/10 | Victorian Infectious Diseases Reference Laboratory and Microbiological Diagnostic Unit Public Health Laboratory, Doherty Institute | Caly L. et al |
| B.1.1.72 | EPI_ISL_577556 | USA/MI-MDHHS-SC22003/2020 | 2020/8/20 | Michigan Department of Health and Human Services, Bureau of Laboratories | Blankenship HM et al |
| B.1.1.74 | EPI_ISL_448972 | NorthernIreland/NIRE-FB730/2020 | 2020/3/25 | COVID-19 Genomics UK (COG-UK) Consortium | Conall McCaughey et al |
| B.1.1.75 | EPI_ISL_468744 | Belgium/UGent-96/2020 | 2020/3/17 | Onderzoeksgroep Virologie | Nick Vereecke et al |
| B.1.1.77 | EPI_ISL_541158 | USA/FL-BPHL-1234/2020 | 2020/5/28 | Florida Bureau of Public Health Laboratories, Florida Department of Health | Schmedes et al |
| B.1.1.8 | EPI_ISL_604245 | USA/NY-QDX-1894/2020 | 2020/3/15 | Quest Diagnostics | Rosenthal et al |
| B.1.1.82 | EPI_ISL_445688 | Wales/PHWC-2ACA2/2020 | 2020/4/6 | Public Health Wales Microbiology Cardiff | Catherine Moore et al |
| B.1.1.83 | EPI_ISL_441399 | NorthernIreland/NIRE-1022DA/2020 | 2020/3/23 | COVID-19 Genomics UK (COG-UK) Consortium | Conall McCaughey et al |
| B.1.1.84 | EPI_ISL_487316 | SouthAfrica/KRISP-0365/2020 | 2020/6/25 | KRISP, KZN Research Innovation and Sequencing Platform | Giandhari J et al |

| | | | | | |
|---|---|---|---|---|---|
| B.1.1.86 | EPI_ISL_559631 | England/ALDP-49F6A6/2020 | 2020/5/22 | Wellcome Sanger Institute for the COVID-19 Genomics UK (COG-UK) consortium | The Lighthouse Lab in Alderley Park et al |
| B.1.1.87 | EPI_ISL_526215 | Hungary/MH-597/2020 | 2020/4/10 | National Laboratory of Virology, Szentágothai Research Centre | Endre Gábor Tóth et al |
| B.1.1.88 | EPI_ISL_454237 | Portugal/PT0513/2020 | 2020/3/20 | Instituto Nacional de Saude (INSA) | Borges et al |
| B.1.1.89 | EPI_ISL_465000 | England/20118179304/2020 | 2020/3/12 | Respiratory Virus Unit, Microbiology Services Colindale, Public Health England | PHE Covid Sequencing Team et al |
| B.1.1.90 | EPI_ISL_3085643 | Sweden/01_SE100_21CS504562/2020 | 2020/4/24 | Karolinska University Hospital | Jan Albert et al |
| B.1.1.91 | EPI_ISL_3946506 | Sweden/01_SE100_21CS508530/2020 | 2020/4/10 | Karolinska University Hospital | Jan Albert et al |
| B.1.1.92 | EPI_ISL_555640 | England/ALDP-94477B/2020 | 2020/6/7 | Wellcome Sanger Institute for the COVID-19 Genomics UK (COG-UK) consortium | The Lighthouse Lab in Alderley Park et al |
| B.1.1.93 | EPI_ISL_636089 | USA/CA-ALSR-3570/2020 | 2020/6/16 | Andersen lab at Scripps Research | SEARCH Alliance San Diego with Tracy Basler et al |
| B.1.1.95 | EPI_ISL_490775 | Wales/PHWC-165BAD/2020 | 2020/4/3 | COVID-19 Genomics UK (COG-UK) Consortium | Catherine Moore et al |
| B.1.1.97 | EPI_ISL_589454 | England/ALDP-9EE272/2020 | 2020/5/6 | Wellcome Sanger Institute for the COVID-19 Genomics UK (COG-UK) consortium | Jacquelyn Wynn et al |
| B.1.1.98 | EPI_ISL_696373 | USA/AZ-TG670175/2020 | 2020/4/5 | TGen North | Jolene Bowers et al |
| B.1.1.99 | EPI_ISL_493744 | Scotland/CVR1838/2020 | 2020/4/7 | COVID-19 Genomics UK (COG-UK) Consortium | Ana da Silva Filipe et al |
| B.1.103 | EPI_ISL_565968 | USA/MI-MDHHS-SC21907/2020 | 2020/4/9 | Michigan Department of Health and Human Services, Bureau of Laboratories | Blankenship HM et al |
| B.1.104 | EPI_ISL_431080 | USA/CT-Yale-079/2020 | 2020/3/2 | Grubaugh Lab - Yale School of Public Health | Joseph Fauver et al |
| B.1.105 | EPI_ISL_440240 | England/CAMB-73114/2020 | 2020/3/21 | Wellcome Sanger Institute for the COVID-19 Genomics UK (COG-UK) consortium | Luke W Meredith et al |
| B.1.106 | EPI_ISL_575081 | USA/UT-UPHL-02807/2020 | 2020/4/23 | Utah Public Health Laboratory | Erin Young et al |
| B.1.108 | EPI_ISL_418895 | USA/CT-UW328/2020 | 2020/3/14 | UW Virology Lab | Pavitra Roychoudhury et al |

| | | | | | |
|---|---|---|---|---|---|
| B.1.110 | EPI_ISL_424855 | USA/FL-CDC-03067097-001/2020 | 2020/3/2 | Pathogen Discovery, Respiratory Viruses Branch, Division of Viral Diseases, Centers for Disease Control and Prevention | Yan Li et al |
| B.1.110.1 | EPI_ISL_801603 | Chile/Santiago-PUC_MVL_0011/2020 | 2020/3/18 | MSHS Pathogen Surveillance Program | Leonardo I. Almonacid et al |
| B.1.110.2 | EPI_ISL_539017 | England/LEED-2A87C7/2020 | 2020/4/3 | Wellcome Sanger Institute for the COVID-19 Genomics UK (COG-UK) consortium | Louissa Macfarlane-Smith et al |
| B.1.110.3 | EPI_ISL_460047 | USA/MN-MDH-106/2020 | 2020/3/15 | Minnesota Department of Health, Public Health Laboratory | Matt Plumb et al |
| B.1.111 | EPI_ISL_424850 | USA/IL-CDC-03068473-001/2020 | 2020/3/7 | Pathogen Discovery, Respiratory Viruses Branch, Division of Viral Diseases, Centers for Disease Control and Prevention | Yan Li et al |
| B.1.112 | EPI_ISL_623194 | USA/UT-UPHL-201114999/2020 | 2020/4/6 | Utah Public Health Laboratory | Erin Young et al |
| B.1.113 | EPI_ISL_469038 | India/GJ-GBRC193b/2020 | 2020/6/3 | Gujarat Biotechnology Research Centre | Komal Patel et al |
| B.1.115 | EPI_ISL_447309 | Israel/701002314/2020 | 2020/3/21 | Stern Lab | Stern Lab et al |
| B.1.116 | EPI_ISL_683682 | USA/MN-MDH-2053/2020 | 2020/3/29 | Minnesota Department of Health, Public Health Laboratory | Alexandra Lorentz et al |
| B.1.117 | EPI_ISL_466120 | England/20149006704/2020 | 2020/3/30 | Respiratory Virus Unit, Microbiology Services Colindale, Public Health England | PHE Covid Sequencing Team et al |
| B.1.118 | EPI_ISL_614581 | Denmark/DCGC-991/2020 | 2020/3/9 | Albertsen lab, Department of Chemistry and Bioscience, Aalborg University, Denmark | Danish Covid-19 Genome Consortia et al |
| B.1.119 | EPI_ISL_436808 | USA/MI-MDHHS-SC20179/2020 | 2020/3/12 | Michigan Department of Health and Human Services, Bureau of Laboratories | Blankenship HM et al |
| B.1.12 | EPI_ISL_420442 | Belgium/BGM-030444/2020 | 2020/3/4 | KU Leuven, Clinical and Epidemiological Virology | Joan Marti-Carreras et al |
| B.1.120 | EPI_ISL_422216 | Wales/PHWC-26ACA/2020 | 2020/3/27 | Public Health Wales Microbiology Cardiff | Catherine Moore et al |
| B.1.124 | EPI_ISL_434164 | USA/WA-S404/2020 | 2020/3/26 | Seattle Flu Study | Chu et al |
| B.1.126 | EPI_ISL_475232 | USA/CA-11567/2020 | 2020/5/5 | UNMC COVID-19 Response Team | UNMC COVID-19 Response Team et al |
| B.1.127 | EPI_ISL_625897 | Denmark/DCGC-8549/2020 | 2020/10/26 | Albertsen lab, Department of Chemistry and Bioscience, Aalborg University, Denmark | Danish Covid-19 Genome Consortia et al |

| | | | | | |
|---|---|---|---|---|---|
| B.1.128 | EPI_ISL_2367357 | Switzerland/VD-CHUV-GEN2477/2020 | 2020/3/9 | Laboratory of genomics and metagenomics | Trestan Pillonel et al |
| B.1.13 | EPI_ISL_418668 | England/20110136302/2020 | 2020/3/9 | Respiratory Virus Unit, Microbiology Services Colindale, Public Health England | Monica Galiano et al |
| B.1.131 | EPI_ISL_979970 | Ukraine/Vinnytsia_12/2020 | 2020/5/27 | Department of Respiratory and other Viral Infections of L.V.Gromashevsky Institute of Epidemiology & Infectious Diseases NAMS of Ukraine, JSC "Farmak" | Alla Mironenko et al |
| B.1.134 | EPI_ISL_3730960 | Italy/TAA-APSS-Ph2_RUN1_NB15/2020 | 2020/4/3 | U.O. Microbiologia e Virologia, Azienda Provinciale per i Servizi Sanitari Provincia Autonoma di Trento, Ospedale S.Chiara | Mirko Moser et al |
| B.1.137 | EPI_ISL_470670 | USA/UT-02054/2020 | 2020/4/20 | Utah Public Health Laboratory | Erin Young et al |
| B.1.139 | EPI_ISL_4025983 | USA/IL-NM-0293/2020 | 2020/3/26 | Northwestern University - Center for Pathogen Genomics and Microbial Evolution | Ramon Lorenzo-Redondo et al |
| B.1.14 | EPI_ISL_435660 | USA/CA-SCCPHD-UC181/2020 | 2020/3/16 | Chiu Laboratory, University of California, San Francisco | Xianding Deng et al |
| B.1.140 | EPI_ISL_5422012 | USA/NE-MP_NCOV20-9850/2020 | 2020/4/30 | NPHL COVID-19 Response Team | NPHL COVID-19 Response Team et al |
| B.1.142 | EPI_ISL_429160 | Sweden/20-50312/2020 | 2020/3/12 | The Public Health Agency of Sweden | Martin Sundqvist et al |
| B.1.143 | EPI_ISL_429770 | Luxembourg/LNS4530810/2020 | 2020/3/28 | Laboratoire National de Sante, Microbiology, Epidemiology and Microbial Genomics | Anke Wienecke-Baldacchino et al |
| B.1.145 | EPI_ISL_435056 | India/GJ-GBRC9/2020 | 2020/4/21 | Gujarat Biotechnology Research Centre | Maharshi Pandya et al |
| B.1.146 | EPI_ISL_486535 | Switzerland/BE-ETHZ-170023/2020 | 2020/6/23 | Department of Biosystems Science and Engineering, ETH Zürich | Christian Beisel et al |
| B.1.149 | EPI_ISL_2307340 | Japan/YCH0069/2020 | 2020/3/31 | Genome Analysis Center, Yamanashi Central Hospital | Yosuke Hirotsu et al |
| B.1.151 | EPI_ISL_753918 | Germany/BE-ChVir-D762-6625/2020 | 2020/3/13 | Charité Universitätsmedizin Berlin, Institut für Virologie | Victor M Corman et al |
| B.1.153 | EPI_ISL_450108 | USA/NY-PV09431/2020 | 2020/3/11 | MSHS Pathogen Surveillance Program | Ana S. Gonzalez-Reiche et al |
| B.1.157 | EPI_ISL_428679 | Spain/MD-H12-LP23-5852/2020 | 2020/3/8 | Hospital Universitario 12 de Octubre | Elias Dahdouh et al |

| | | | | | |
|---|---|---|---|---|---|
| B.1.158 | EPI_ISL_418390 | Finland/13M33/2020 | 2020/3/13 | Department of Virology, Faculty of Medicine, University of Helsinki, Helsinki, Finland | Teemu Smura et al |
| B.1.159 | EPI_ISL_940432 | France/IDF_HB_112003063212/2020 | 2020/3/17 | IAME UMR1137 Inserm, Université de Paris, Hôpital Bichat | Antoine Bridier-Nahmias et al |
| B.1.160 | EPI_ISL_4899892 | Morocco/INH-106/2020 | 2020/2/2 | Functional Genomic Platform UATRS-biology, CNRST | Hicham Oumzil et al |
| B.1.160.10 | EPI_ISL_3357877 | France/IDF-APHPSorbonneUniversite_632/2020 | 2020/8/27 | Department of Virology, Pitié-Salpêtrière hospital | Valentin Leducq et al |
| B.1.160.11 | EPI_ISL_603549 | Switzerland/BL-ETHZ-310479/2020 | 2020/10/8 | Department of Biosystems Science and Engineering, ETH Zürich | Christian Beisel et al |
| B.1.160.13 | EPI_ISL_578025 | Netherlands/ZH-EMC-597/2020 | 2020/9/18 | Erasmus Medical Center | Bas Oude Munnink et al |
| B.1.160.14 | EPI_ISL_541417 | Switzerland/BE-ETHZ-270048/2020 | 2020/9/1 | Department of Biosystems Science and Engineering, ETH Zürich | Christian Beisel et al |
| B.1.160.15 | EPI_ISL_603297 | Switzerland/UR-ETHZ-300265/2020 | 2020/9/30 | Department of Biosystems Science and Engineering, ETH Zürich | Christian Beisel et al |
| B.1.160.16 | EPI_ISL_603405 | Switzerland/BE-ETHZ-300453/2020 | 2020/9/30 | Department of Biosystems Science and Engineering, ETH Zürich | Christian Beisel et al |
| B.1.160.16 | EPI_ISL_2757556 | France/PAC-IHU-13469_Nova1/2020 | 2020/9/15 | MEPHI, Aix Marseille University | Anthony LEVASSEUR et al |
| B.1.160.17 | EPI_ISL_668504 | Denmark/DCGC-9534/2020 | 2020/11/9 | Albertsen Lab, Department of Chemistry and Bioscience, Aalborg University, Denmark | Danish Covid-19 Genome Consortium et al |
| B.1.160.18 | EPI_ISL_730649 | France/ARA-104848/2020 | 2020/11/10 | CNR Virus des Infections Respiratoires - France SUD | Antonin Bal et al |
| B.1.160.19 | EPI_ISL_2611008 | Switzerland/LU-UHB-42555800/2020 | 2020/11/24 | Clinical Bacteriology | Tim Roloff et al |
| B.1.160.20 | EPI_ISL_737834 | Switzerland/VS-ETHZ-320410/2020 | 2020/10/12 | Department of Biosystems Science and Engineering, ETH Zürich | Chaoran Chen et al |
| B.1.160.21 | EPI_ISL_744845 | Luxembourg/LNS9084900/2020 | 2020/11/24 | Laboratoire national de santé, Microbiology, Microbial Genomics Platform | Anke Wienecke-Baldacchino et al |
| B.1.160.22 | EPI_ISL_603526 | Switzerland/TI-ETHZ-310454/2020 | 2020/10/7 | Department of Biosystems Science and Engineering, ETH Zürich | Christian Beisel et al |
| B.1.160.23 | EPI_ISL_566530 | England/MILK-9C297F/2020 | 2020/9/7 | Wellcome Sanger Institute for the COVID-19 Genomics UK (COG-UK) consortium | The Lighthouse Lab in Milton Keynes et al |

| | | | | | |
|---|---|---|---|---|---|
| B.1.160.24 | EPI_ISL_711390 | Denmark/DCGC-14903/2020 | 2020/11/23 | Albertsen Lab, Department of Chemistry and Bioscience, Aalborg University, Denmark | Danish Covid-19 Genome Consortium et al |
| B.1.160.25 | EPI_ISL_918554 | Brazil/AP-IEC-177390/2020 | 2020/12/16 | Evandro Chagas Institute | Santos et al |
| B.1.160.26 | EPI_ISL_603330 | Switzerland/VD-ETHZ-300298/2020 | 2020/9/25 | Department of Biosystems Science and Engineering, ETH Zürich | Christian Beisel et al |
| B.1.160.27 | EPI_ISL_855362 | Mayotte/IPP00169/2021 | 2021/1/5 | National Reference Center for Viruses of Respiratory Infections, Institut Pasteur, Paris | Marion Barbet et al |
| B.1.160.28 | EPI_ISL_707715 | Belgium/ULG-10947/2020 | 2020/9/5 | GIGA Medical Genomics | Keith Durkin et al |
| B.1.160.29 | EPI_ISL_721807 | Switzerland/GE-ETHZ-340094/2020 | 2020/10/26 | Department of Biosystems Science and Engineering, ETH Zürich | Christian Beisel et al |
| B.1.160.30 | EPI_ISL_537141 | England/QEUH-95DF1D/2020 | 2020/8/7 | Wellcome Sanger Institute for the COVID-19 Genomics UK (COG-UK) consortium | Harper VanSteenhouse et al |
| B.1.160.30 | EPI_ISL_1140271 | Germany/NW-RKI-I-010146/2020 | 2020/9/25 | Robert Koch Institute | ? |
| B.1.160.31 | EPI_ISL_603484 | Switzerland/SZ-ETHZ-310410/2020 | 2020/10/8 | Department of Biosystems Science and Engineering, ETH Zürich | Christian Beisel et al |
| B.1.160.32 | EPI_ISL_2596349 | France/PAC-IHU-11568_Illu1/2020 | 2020/4/22 | MEPHI, Aix Marseille University | Anthony LEVASSEUR et al |
| B.1.160.33 | EPI_ISL_5314801 | France/ARA-HCL990000125202/2020 | 2020/10/26 | CNR Virus des Infections Respiratoires - France SUD | Antonin Bal et al |
| B.1.160.7 | EPI_ISL_595857 | Wales/QEUH-A31057/2020 | 2020/10/3 | COVID-19 Genomics UK (COG-UK) Consortium | Dave J. Baker et al |
| B.1.160.8 | EPI_ISL_619449 | Denmark/DCGC-6549/2020 | 2020/10/5 | Albertsen lab, Department of Chemistry and Bioscience, Aalborg University, Denmark | Danish Covid-19 Genome Consortia et al |
| B.1.160.9 | EPI_ISL_1496478 | Switzerland/BL-ETHZ-330467/2020 | 2020/10/19 | Department of Biosystems Science and Engineering, ETH Zürich | Christian Beisel et al |
| B.1.161 | EPI_ISL_482455 | USA/OR-PROV-220/2020 | 2020/3/27 | Providence St. Joseph Health Molecular Genomics Laboratory | Alexa K Dowdell et al |
| B.1.162 | EPI_ISL_444599 | USA/IL-NM0116/2020 | 2020/3/19 | Ozer Lab | Ramon Lorenzo-Redondo et al |
| B.1.163 | EPI_ISL_468389 | USA/CA-CZB-1414/2020 | 2020/3/16 | Chan-Zuckerberg Biohub | CZB Clinhub Consortium et al |
| B.1.164 | EPI_ISL_456565 | Australia/VIC1664/2020 | 2020/5/10 | Microbiological Diagnostic Unit Public Health Laboratory and Victorian Infectious Diseases Reference Laboratory, Doherty Institute | Caly L. et al |

| | | | | | |
|---|---|---|---|---|---|
| B.1.165 | EPI_ISL_458367 | England/BRIS-12ACFC/2020 | 2020/5/1 | Wellcome Sanger Institute for the COVID-19 Genomics UK (COG-UK) consortium | Stephanie Hutchings et al |
| B.1.166 | EPI_ISL_483194 | USA/CA-ALSR-0515-SAN/2020 | 2020/3/4 | Andersen lab at Scripps Research | SEARCH Alliance San Diego with David Pride et al |
| B.1.167 | EPI_ISL_443943 | England/BRIS-1253DE/2020 | 2020/3/27 | Wellcome Sanger Institute for the COVID-19 Genomics UK (COG-UK) consortium | Stephanie Hutchings et al |
| B.1.168 | EPI_ISL_453122 | Scotland/EDB4335/2020 | 2020/4/23 | COVID-19 Genomics UK (COG-UK) Consortium | McHugh M et al |
| B.1.169 | EPI_ISL_570230 | USA/WA-UW-9195/2020 | 2020/5/15 | UW Virology Lab | Pavitra Roychoudhury et al |
| B.1.170 | EPI_ISL_603907 | USA/CA-QDX-1625/2020 | 2020/3/19 | Quest Diagnostics | Rosenthal et al |
| B.1.173 | EPI_ISL_537248 | England/SHEF-10B2E79/2020 | 2020/5/16 | Wellcome Sanger Institute for the COVID-19 Genomics UK (COG-UK) consortium | Thushan de Silva et al |
| B.1.177 | EPI_ISL_4899898 | Morocco/INH-107/2020 | 2020/2/2 | Functional Genomic Platform UATRS-biology, CNRST | Mohamed Rhajaoui et al |
| B.1.177 | EPI_ISL_3054039 | France/PAC-IHU-15335-N1/2021 | 2020/7/29 | MEPHI, Aix Marseille University | Anthony LEVASSEUR et al |
| B.1.177 | EPI_ISL_2017564 | Spain/IB-HUSE-00072/2020 | 2020/9/26 | HOSPITAL UNIVERSITARIO SON ESPASES | Carla López-Causapé et al |
| B.1.177.10 | EPI_ISL_590540 | England/QEUH-9B7746/2020 | 2020/9/2 | Wellcome Sanger Institute for the COVID-19 Genomics UK (COG-UK) consortium | Harper VanSteenhouse et al |
| B.1.177.11 | EPI_ISL_760151 | SouthKorea/KDCA0367/2020 | 2020/4/4 | Division of Emerging Infectious Diseases, Bureau of Infectious Diseases Diagnosis Control, Korea Disease Control and Prevention Agency | Ae Kyung Park et al |
| B.1.177.12 | EPI_ISL_671267 | Denmark/DCGC-12020/2020 | 2020/3/30 | Albertsen Lab, Department of Chemistry and Bioscience, Aalborg University, Denmark | Danish Covid-19 Genome Consortium et al |
| B.1.177.14 | EPI_ISL_4533657 | Spain/UN-ORC01386/2020 | 2020/7/27 | IAME UMR1137 Inserm, Université de Paris, Hôpital Bichat | Antoine Bridier-Nahmias et al |
| B.1.177.15 | EPI_ISL_560081 | Wales/MILK-9A89BB/2020 | 2020/9/1 | COVID-19 Genomics UK (COG-UK) Consortium | Tanya Golubchik et al |
| B.1.177.16 | EPI_ISL_567654 | England/ALDP-9C5BB2/2020 | 2020/9/10 | Wellcome Sanger Institute for the COVID-19 Genomics UK (COG-UK) consortium | Jacquelyn Wynn et al |

| | | | | | |
|---|---|---|---|---|---|
| B.1.177.17 | EPI_ISL_590323 | England/QEUH-9B4A13/2020 | 2020/8/2 | Wellcome Sanger Institute for the COVID-19 Genomics UK (COG-UK) consortium | Harper VanSteenhouse et al |
| B.1.177.18 | EPI_ISL_573138 | England/CAMB-1B5F2A/2020 | 2020/9/14 | COVID-19 Genomics UK (COG-UK) Consortium | Aminu S. Jahun et al |
| B.1.177.19 | EPI_ISL_590415 | England/QEUH-9B4E35/2020 | 2020/9/3 | Wellcome Sanger Institute for the COVID-19 Genomics UK (COG-UK) consortium | Harper VanSteenhouse et al |
| B.1.177.2 | EPI_ISL_590273 | England/QEUH-9B4BC5/2020 | 2020/9/3 | Wellcome Sanger Institute for the COVID-19 Genomics UK (COG-UK) consortium | Harper VanSteenhouse et al |
| B.1.177.20 | EPI_ISL_611771 | England/PORT-2D2111/2020 | 2020/3/21 | COVID-19 Genomics UK (COG-UK) Consortium | Angela Beckett et al |
| B.1.177.21 | EPI_ISL_616626 | Denmark/DCGC-2791/2020 | 2020/8/24 | Albertsen lab, Department of Chemistry and Bioscience, Aalborg University, Denmark | Danish Covid-19 Genome Consortia et al |
| B.1.177.23 | EPI_ISL_2790847 | France/IDF-IPP5558/2020 | 2020/4/8 | National Reference Center for Viruses of Respiratory Infections, Institut Pasteur, Paris | Marion Barbet et al |
| B.1.177.24 | EPI_ISL_616503 | Denmark/DCGC-2624/2020 | 2020/8/17 | Albertsen lab, Department of Chemistry and Bioscience, Aalborg University, Denmark | Danish Covid-19 Genome Consortia et al |
| B.1.177.25 | EPI_ISL_613639 | Gibraltar/204421176/2020 | 2020/10/27 | Respiratory Virus Unit, Microbiology Services Colindale, Public Health England | PHE Covid Sequencing Team et al |
| B.1.177.26 | EPI_ISL_595162 | England/NORW-EFE2B/2020 | 2020/10/8 | COVID-19 Genomics UK (COG-UK) Consortium | Dave J. Baker et al |
| B.1.177.27 | EPI_ISL_620364 | Denmark/DCGC-6464/2020 | 2020/10/12 | Albertsen lab, Department of Chemistry and Bioscience, Aalborg University, Denmark | Danish Covid-19 Genome Consortia et al |
| B.1.177.28 | EPI_ISL_614910 | Switzerland/BE-ETHZ-330427/2020 | 2020/10/18 | Department of Biosystems Science and Engineering, ETH Zürich | Christian Beisel et al |
| B.1.177.29 | EPI_ISL_871914 | Spain/CT-IBV-98013069/2020 | 2020/12/18 | SeqCOVID-SPAIN consortium/IBV(CSIC) | Elisa Martró et al |
| B.1.177.3 | EPI_ISL_552340 | England/MILK-995201/2020 | 2020/8/24 | Wellcome Sanger Institute for the COVID-19 Genomics UK (COG-UK) consortium | The Lighthouse Lab in Milton Keynes et al |

| | | | | | |
|---|---|---|---|---|---|
| B.1.177.30 | EPI_ISL_649884 | England/204393759/2020 | 2020/10/23 | COVID-19 Genomics UK (COG-UK) Consortium | PHE Covid Sequencing Team et al |
| B.1.177.31 | EPI_ISL_541841 | England/QEUH-999100/2020 | 2020/8/24 | Wellcome Sanger Institute for the COVID-19 Genomics UK (COG-UK) consortium | Harper VanSteenhouse et al |
| B.1.177.32 | EPI_ISL_2323027 | Spain/CL-IBV-99016864/2020 | 2020/9/29 | SeqCOVID-SPAIN consortium/IBV(CSIC) | Ana Carvajal et al |
| B.1.177.33 | EPI_ISL_837273 | Italy/CAM-TIGEM-1017/2020 | 2020/9/2 | TIGEM | Antonio Grimaldi et al |
| B.1.177.34 | EPI_ISL_722792 | Netherlands/ZH-EMC-1052/2020 | 2020/12/6 | Erasmus Medical Center | Bas Oude Munnink et al |
| B.1.177.35 | EPI_ISL_2099982 | Spain/AN-IBV-99009691/2020 | 2020/9/26 | SeqCOVID-SPAIN consortium/IBV(CSIC) | Federico García et al |
| B.1.177.36 | EPI_ISL_1311034 | Netherlands/NB-EMC-754/2020 | 2020/9/14 | Erasmus Medical Center | Bas Oude Munnink et al |
| B.1.177.37 | EPI_ISL_661244 | Belgium/ULG-10821/2020 | 2020/11/16 | GIGA Medical Genomics | Keith Durkin et al |
| B.1.177.38 | EPI_ISL_567667 | England/ALDP-9C5B3A/2020 | 2020/9/11 | Wellcome Sanger Institute for the COVID-19 Genomics UK (COG-UK) consortium | Jacquelyn Wynn et al |
| B.1.177.39 | EPI_ISL_1120921 | Spain/MD-IBV-99020702/2020 | 2020/9/2 | SeqCOVID-SPAIN consortium/IBV(CSIC) | Esther Viedma et al |
| B.1.177.4 | EPI_ISL_950611 | England/NOTT-246F3C/2020 | 2020/6/3 | COVID-19 Genomics UK (COG-UK) Consortium | Gemma Clark et al |
| B.1.177.40 | EPI_ISL_1819069 | Spain/AN-IBV-97014065/2020 | 2020/8/15 | SeqCOVID-SPAIN consortium/IBV(CSIC) | Inmaculada de Toro Peinado et al |
| B.1.177.41 | EPI_ISL_635185 | Norway/3621/2020 | 2020/9/20 | Norwegian Institute of Public Health, Department of Virology | Kathrine Stene-Johansen et al |
| B.1.177.42 | EPI_ISL_617404 | Denmark/DCGC-8449/2020 | 2020/10/26 | Albertsen lab, Department of Chemistry and Bioscience, Aalborg University, Denmark | Danish Covid-19 Genome Consortia et al |
| B.1.177.43 | EPI_ISL_965734 | Netherlands/LI-MUMC-166/2020 | 2020/8/17 | Medical Microbiology, Maastricht University Medical Centre | Jozef Dingemans* et al |
| B.1.177.45 | EPI_ISL_826922 | Iceland/3230/2020 | 2020/9/12 | deCODE genetics | Daniel F Gudbjartsson et al |
| B.1.177.46 | EPI_ISL_648159 | Sweden/20-53276/2020 | 2020/10/29 | The Public Health Agency of Sweden | Anna-Malin Linde et al |
| B.1.177.47 | EPI_ISL_724276 | England/EXET-13DB08/2020 | 2020/12/1 | COVID-19 Genomics UK (COG-UK) Consortium | Ben Temperton et al |
| B.1.177.48 | EPI_ISL_580212 | England/ALDP-9E9F16/2020 | 2020/9/24 | Wellcome Sanger Institute for the COVID-19 Genomics UK (COG-UK) consortium | Jacquelyn Wynn et al |

| | | | | | |
|---|---|---|---|---|---|
| B.1.177.49 | EPI_ISL_831148 | Spain/MD-IBV-99013265/2020 | 2020/9/11 | SeqCOVID-SPAIN consortium/IBV(CSIC) | María Rodríguez-Tejedor et al |
| B.1.177.5 | EPI_ISL_540362 | Scotland/QEUH-98F404/2020 | 2020/8/29 | Wellcome Sanger Institute for the COVID-19 Genomics UK (COG-UK) consortium | Harper VanSteenhouse et al |
| B.1.177.50 | EPI_ISL_539605 | Germany/NW-HHU-89/2020 | 2020/8/17 | Center of Medical Microbiology, Virology, and Hospital Hygiene, University of Duesseldorf | Maximilian Damagnez et al |
| B.1.177.51 | EPI_ISL_3800826 | Italy/LIG-111980-037-01/2020 | 2020/10/1 | Istituto Zooprofilattico Sperimentale della Lombardia e dell'Emilia Romagna (IZSLER), Risk Analysis and Genomic Epidemiology Unit | Bianca Bruzzone et al |
| B.1.177.52 | EPI_ISL_553019 | England/MILK-9714F6/2020 | 2020/8/18 | Wellcome Sanger Institute for the COVID-19 Genomics UK (COG-UK) consortium | The Lighthouse Lab in Milton Keynes et al |
| B.1.177.53 | EPI_ISL_532369 | England/QEUH-95FBF9/2020 | 2020/8/3 | Wellcome Sanger Institute for the COVID-19 Genomics UK (COG-UK) consortium | Harper VanSteenhouse et al |
| B.1.177.54 | EPI_ISL_578257 | Ireland/D-NVRL-72IRL36222/2020 | 2020/8/17 | National Virus Reference Laboratory | Michael Carr et al |
| B.1.177.55 | EPI_ISL_573020 | England/ALDP-9B661D/2020 | 2020/8/28 | COVID-19 Genomics UK (COG-UK) Consortium | Tanya Golubchik et al |
| B.1.177.56 | EPI_ISL_534365 | Scotland/QEUH-964335/2020 | 2020/8/12 | Wellcome Sanger Institute for the COVID-19 Genomics UK (COG-UK) consortium | Ana da Silva Filipe et al |
| B.1.177.57 | EPI_ISL_950607 | England/NOTT-246E4E/2020 | 2020/6/2 | COVID-19 Genomics UK (COG-UK) Consortium | Gemma Clark et al |
| B.1.177.58 | EPI_ISL_580665 | England/MILK-9F0811/2020 | 2020/9/25 | Wellcome Sanger Institute for the COVID-19 Genomics UK (COG-UK) consortium | The Lighthouse Lab in Milton Keynes et al |
| B.1.177.59 | EPI_ISL_578262 | Ireland/D-NVRL-73IRL25053/2020 | 2020/9/5 | National Virus Reference Laboratory | Michael Carr et al |
| B.1.177.6 | EPI_ISL_524856 | USA/UT-UPHL-2008837/2020 | 2020/5/29 | Utah Public Health Laboratory | Erin L. Young et al |
| B.1.177.60 | EPI_ISL_534665 | Scotland/QEUH-96A96A/2020 | 2020/8/17 | Wellcome Sanger Institute for the COVID-19 Genomics UK (COG-UK) consortium | Ana da Silva Filipe et al |
| B.1.177.60 | EPI_ISL_1312496 | Latvia/985/2020 | 2020/9/28 | Latvian Biomedical Research and Study Centre | Janis Pjalkovskis et al |

| | | | | | |
|---|---|---|---|---|---|
| B.1.177.62 | EPI_ISL_2869783 | Germany/NW-KRO-273/2020 | 2020/10/2 | Center of Medical Microbiology, Virology, and Hospital Hygiene, University of Duesseldorf | Dennis Deschka et al |
| B.1.177.63 | EPI_ISL_595582 | Wales/PHWC-47ED20/2020 | 2020/10/13 | COVID-19 Genomics UK (COG-UK) Consortium | Catherine Moore et al |
| B.1.177.64 | EPI_ISL_818534 | Denmark/DCGC-25502/2021 | 2021/1/4 | Albertsen Lab, Department of Chemistry and Bioscience, Aalborg University, Denmark | Danish Covid-19 Genome Consortium et al |
| B.1.177.65 | EPI_ISL_532427 | Wales/QEUH-95E978/2020 | 2020/8/6 | Wellcome Sanger Institute for the COVID-19 Genomics UK (COG-UK) consortium | Harper VanSteenhouse et al |
| B.1.177.66 | EPI_ISL_768786 | Ireland/D-NVRL-AIIDV1268v1/2020 | 2020/10/30 | Irish Coronavirus Sequencing Consortium - National Virus Reference Laboratory | Michael Carr et al |
| B.1.177.67 | EPI_ISL_752524 | Ireland/MH-NVRL-77IRL72701/2020 | 2020/11/5 | National Virus Reference Laboratory | Michael Carr et al |
| B.1.177.68 | EPI_ISL_837373 | Ireland/LK-NVRL-81IRL14907/2020 | 2020/12/27 | National Virus Reference Laboratory | Michael Carr et al |
| B.1.177.69 | EPI_ISL_599678 | England/QEUH-9DCE55/2020 | 2020/9/21 | Wellcome Sanger Institute for the COVID-19 Genomics UK (COG-UK) consortium | Harper VanSteenhouse et al |
| B.1.177.7 | EPI_ISL_693277 | Australia/QLD1276/2020 | 2020/3/21 | Queensland Health Forensic and Scientific Services | Son Nguyen et al |
| B.1.177.70 | EPI_ISL_669121 | Denmark/DCGC-9709/2020 | 2020/11/9 | Albertsen Lab, Department of Chemistry and Bioscience, Aalborg University, Denmark | Danish Covid-19 Genome Consortium et al |
| B.1.177.71 | EPI_ISL_1296403 | France/IDF_HB_112009081106/2020 | 2020/9/21 | IAME UMR1137 Inserm, Université de Paris, Hôpital Bichat | Antoine Bridier-Nahmias et al |
| B.1.177.72 | EPI_ISL_1119698 | Switzerland/SO-ETHZ-501186/2020 | 2020/11/7 | Department of Biosystems Science and Engineering, ETH Zürich | Chaoran Chen et al |
| B.1.177.73 | EPI_ISL_1311716 | Netherlands/ZH-EMC-1816/2020 | 2020/4/12 | Erasmus Medical Center | Bas Oude Munnink et al |
| B.1.177.74 | EPI_ISL_2458149 | Spain/CT-HUVH-R85736/2020 | 2020/10/27 | Hospital Universitari Vall d'Hebron - Vall d'Hebron Institut de Recerca | Cristina Andrés et al |

| | | | | | |
|---|---|---|---|---|---|
| B.1.177.75 | EPI_ISL_1692830 | Italy/CAM-TIGEM-IZSM-COLLI-9497/2020 | 2020/9/2 | Telethon Institute of Genetics and Medicine (TIGEM) | Antonio Grimaldi Patrizia Annunziata Francesco Panariello Teresa Giuliano Michele Cennamo Valentina Bouche Chiara Colantuono Lucio Di Filippo Mariano Fiorenza Anna Manfredi Marcello Salvi Giuseppe Portella Andrea Ballabio Davide Cacchiarelli et al |
| B.1.177.76 | EPI_ISL_722256 | Spain/AR-IBV-98010614/2020 | 2020/10/1 | SeqCOVID-SPAIN consortium/IBV(CSIC) | Antonio Rezusta López et al |
| B.1.177.77 | EPI_ISL_3356319 | Belgium/LHUB11590048/2020 | 2020/3/31 | LHUB-ULB | Charlotte Michel et al |
| B.1.177.8 | EPI_ISL_560031 | Scotland/QEUH-9AF18F/2020 | 2020/9/3 | COVID-19 Genomics UK (COG-UK) Consortium | Tanya Golubchik et al |
| B.1.177.80 | EPI_ISL_661290 | Sweden/20-53262/2020 | 2020/10/21 | The Public Health Agency of Sweden | Department of Microbiology et al |
| B.1.177.81 | EPI_ISL_1311761 | Netherlands/NB-EMC-847/2020 | 2020/8/10 | Erasmus Medical Center | Bas Oude Munnink et al |
| B.1.177.82 | EPI_ISL_552201 | England/MILK-99390E/2020 | 2020/8/26 | Wellcome Sanger Institute for the COVID-19 Genomics UK (COG-UK) consortium | The Lighthouse Lab in Milton Keynes et al |
| B.1.177.83 | EPI_ISL_547965 | Italy/SAR-ATS-246/2020 | 2020/8/25 | Laboratorio Specialistico UOC Ematologia Ospedale San Francesco - ATS ASSL NUORO | Piras Giovanna et al |
| B.1.177.84 | EPI_ISL_566346 | England/ALDP-9CDC5D/2020 | 2020/9/16 | Wellcome Sanger Institute for the COVID-19 Genomics UK (COG-UK) consortium | Jacquelyn Wynn et al |
| B.1.177.85 | EPI_ISL_740149 | Luxembourg/LNS2593581/2020 | 2020/10/23 | Laboratoire national de santé, Microbiology, Microbial Genomics Platform | Anke Wienecke-Baldacchino et al |
| B.1.177.86 | EPI_ISL_621760 | Denmark/DCGC-5222/2020 | 2020/9/21 | Albertsen lab, Department of Chemistry and Bioscience, Aalborg University, Denmark | Danish Covid-19 Genome Consortia et al |
| B.1.177.87 | EPI_ISL_552100 | England/MILK-992EB5/2020 | 2020/8/26 | Wellcome Sanger Institute for the COVID-19 Genomics UK (COG-UK) consortium | The Lighthouse Lab in Milton Keynes et al |
| B.1.177.88 | EPI_ISL_837496 | Italy/CAM-TIGEM-1344/2020 | 2020/12/26 | TIGEM | Antonio Grimaldi et al |

| | | | | | |
|---|---|---|---|---|---|
| B.1.177.89 | EPI_ISL_1658340 | Switzerland/BE-ETHZ-570061/2020 | 2020/12/16 | Department of Biosystems Science and Engineering, ETH Zurich | Christian Beisel et al |
| B.1.177.9 | EPI_ISL_638633 | England/QEUH-9AE33F/2020 | 2020/9/2 | COVID-19 Genomics UK (COG-UK) Consortium | Aminu S. Jahun et al |
| B.1.178 | EPI_ISL_417437 | DRC/KN-0054/2020 | 2020/3/17 | Pathogen Sequencing Lab, National Institute for Biomedical Research (INRB) | Placide Mbala-Kingebeni et al |
| B.1.179 | EPI_ISL_614557 | Denmark/DCGC-961/2020 | 2020/3/9 | Albertsen lab, Department of Chemistry and Bioscience, Aalborg University, Denmark | Danish Covid-19 Genome Consortia et al |
| B.1.180 | EPI_ISL_1501744 | USA/IL-LCH-0052/2020 | 2020/4/6 | Northwestern University - Ozer Lab | Ramon Lorenzo-Redondo et al |
| B.1.181 | EPI_ISL_482362 | USA/OR-PROV-091/2020 | 2020/4/20 | Providence St. Joseph Health Molecular Genomics Laboratory | Alexa K Dowdell et al |
| B.1.182 | EPI_ISL_509724 | USA/FL-BPHL-0666/2020 | 2020/3/13 | Florida Bureau of Public Health Laboratories | Sarah Schmedes et al |
| B.1.184 | EPI_ISL_524739 | India/GJ-GBRC-380a/2020 | 2020/6/18 | Gujarat Biotechnology Research Centre | Pinal Trivedi et al |
| B.1.187 | EPI_ISL_428927 | Poland/PL_P4/2020 | 2020/3/28 | ViroGenetics - BSL3 Laboratory of Virology; Human Genome Variation Research Group & Genomics Centre MCB; Bioinformatics Research Group | Wojciech Branicki et al |
| B.1.188 | EPI_ISL_509203 | USA/OR-OHSU-1447/2020 | 2020/7/17 | Oregon SARS-CoV-2 Genome Sequencing Center | Brendan L. O'Connell et al |
| B.1.189 | EPI_ISL_837604 | Mexico/CMX-INER-0051/2020 | 2020/4/14 | Instituto Nacional de Enfermedades Respiratorias (INER) | Celia Boukadida et al |
| B.1.190 | EPI_ISL_547557 | Netherlands/ZH-RIVM-10050/2020 | 2020/3/13 | National Institute for Public Health and the Environment (RIVM) | Adam Meijer et al |
| B.1.192 | EPI_ISL_648326 | EquatorialGuinea/3350/2020 | 2020/4/24 | University Hospital Basel, Clinical Bacteriology | Carlos Cortes et al |
| B.1.194 | EPI_ISL_545420 | USA/TX-HMH-2603/2020 | 2020/6/4 | Houston Methodist Hospital | S. Wesley Long et al |
| B.1.195 | EPI_ISL_541340 | Brazil/PR-FIOCRUZ-5617/2020 | 2020/3/17 | Laboratory of Respiratory Viruses and Measles, Oswaldo Cruz Institute, FIOCRUZ | Paola Resende et al |
| B.1.198 | EPI_ISL_440738 | England/BRIS-122BAC/2020 | 2020/4/3 | Wellcome Sanger Institute for the COVID-19 Genomics UK (COG-UK) consortium | Stephanie Hutchings et al |
| B.1.199 | EPI_ISL_467811 | USA/CA-QDX-117/2020 | 2020/3/16 | Quest Diagnostics | Anderson et al |
| B.1.2 | EPI_ISL_2835566 | USA/CA-SEARCH-103149/2020 | 2020/1/21 | Andersen lab at Scripps Research | SEARCH Alliance San Diego with Aaron Harding et al |

| | | | | | |
|---|---|---|---|---|---|
| B.1.201 | EPI_ISL_464549 | England/20109028804/2020 | 2020/3/6 | Respiratory Virus Unit, Microbiology Services Colindale, Public Health England | PHE Covid Sequencing Team et al |
| B.1.203 | EPI_ISL_527748 | CostaRica/INC-0080/2020 | 2020/7/5 | Inciensa, Instituto Costarricense de Investigación y Enseñanza en Nutrición y Salud | Francisco Duarte et al |
| B.1.205 | EPI_ISL_482468 | Peru/LIM-01-2/2020 | 2020/3/5 | Laboratorio de Referencia Nacional de Biotecnologia y Biologia Molecular. Instituto Nacional de Salud Peru | Carlos Padilla Rojas et al |
| B.1.206 | EPI_ISL_511903 | India/WB-IK62/2020 | 2020/4/9 | National Institute of Biomedical Genomics - DBT's PAN-INDIA 1000 SARS--CoV-2 RNA Genome Sequencing Consortium | Arindam Maitra et al |
| B.1.208 | EPI_ISL_418414 | France/ARA-09434/2020 | 2020/3/15 | CNR Virus des Infections Respiratoires - France SUD | Antonin Bal et al |
| B.1.210 | EPI_ISL_1055383 | India/MH-ICMR-C300/2020 | 2020/3/10 | Indian Council of Medical Research- National Institute of Virology, Microbial Containment Complex | Pragya D. Yadav et al |
| B.1.211 | EPI_ISL_416749 | France/ARA-532/2020 | 2020/3/4 | CNR Virus des Infections Respiratoires - France SUD | Bal et al |
| B.1.212 | EPI_ISL_414629 | France/HDF-1870/2020 | 2020/3/3 | National Reference Center for Viruses of Respiratory Infections, Institut Pasteur, Paris | Mélnie Albert et al |
| B.1.213 | EPI_ISL_645071 | Poland/PL_MCB_72/2020 | 2020/5/10 | Human Genome Variation Research Group, Malopolska Centre of Biotechnology | Kowalski et al |
| B.1.214 | EPI_ISL_437357 | DRC/3070/2020 | 2020/4/18 | Pathogen Sequencing Lab, National Institute for Biomedical Research (INRB) | Placide Mbala-Kingebeni et al |
| B.1.214.1 | EPI_ISL_912354 | Congo/Q001C088AF_2101-TM-r1-002/2020 | 2020/12/5 | NGS Competence Center Tuebingen, Institut für Medizinische Mikrobiologie und Hygiene, Universitaetsklinikum Tübingen | Angel Angelov et al |
| B.1.214.3 | EPI_ISL_1384313 | Luxembourg/LNS7752025/2020 | 2020/12/14 | Laboratoire national de sante, Microbiology, Microbial Genomics Platform | Anke Wienecke-Baldacchino et al |
| B.1.214.4 | EPI_ISL_971814 | Denmark/DCGC-40137/2021 | 2021/1/25 | Aalborg University | Danish Covid-19 Genome Consortium et al |
| B.1.215 | EPI_ISL_485853 | USA/VA-DCLS-0552/2020 | 2020/3/14 | Virginia DCLS | Virginia DCLS et al |

| | | | | | |
|---|---|---|---|---|---|
| B.1.218 | EPI_ISL_618374 | Denmark/DCGC-2112/2020 | 2020/7/20 | Albertsen lab, Department of Chemistry and Bioscience, Aalborg University, Denmark | Danish Covid-19 Genome Consortia et al |
| B.1.219 | EPI_ISL_536230 | Canada/Qc-LSPQ-L00239009/2020 | 2020/3/22 | Laboratoire de santé publique du Québec | Sandrine Moreira et al |
| B.1.22 | EPI_ISL_523199 | Netherlands/GE-EMC-217/2020 | 2020/3/5 | Erasmus Medical Center | Bas Oude Munnink et al |
| B.1.22.1 | EPI_ISL_636603 | SintEustatius/BQ-RIVM-10373/2020 | 2020/9/7 | National Institute for Public Health and the Environment (RIVM) | Adam Meijer et al |
| B.1.220 | EPI_ISL_457974 | Oman/RESP-20-2282/2020 | 2020/3/14 | Oman-NIC | Samira Al-Maruqi et al |
| B.1.221 | EPI_ISL_2631277 | Poland/KCZ_1.73/2020 | 2020/1/14 | Department of Infectious, Tropical Diseases and Immune Deficiency, Pomeranian Medical University in Szczecin, Szczecin, Poland | Karol Serwin et al |
| B.1.221 | EPI_ISL_1225484 | Panama/GMI-PA395551/2020 | 2020/6/24 | Gorgas Memorial Laboratory of Health Studies | Yamilka Diaz et al |
| B.1.221.1 | EPI_ISL_553089 | England/MILK-970CFF/2020 | 2020/8/18 | Wellcome Sanger Institute for the COVID-19 Genomics UK (COG-UK) consortium | The Lighthouse Lab in Milton Keynes et al |
| B.1.221.2 | EPI_ISL_2448328 | Germany/SL-SU-10446284/2020 | 2020/9/10 | Epigenetics, Saarland University | Kathrin Kattler et al |
| B.1.221.3 | EPI_ISL_615254 | Denmark/DCGC-5323/2020 | 2020/9/14 | Albertsen lab, Department of Chemistry and Bioscience, Aalborg University, Denmark | Danish Covid-19 Genome Consortia et al |
| B.1.221.4 | EPI_ISL_744659 | Luxembourg/LNS7948120/2020 | 2020/11/2 | Laboratoire national de santé, Microbiology, Microbial Genomics Platform | Anke Wienecke-Baldacchino et al |
| B.1.222 | EPI_ISL_779712 | Italy/LOM-UniMI08/2020 | 2020/2/24 | Laboratory of Infectious Diseases, Department of Biomedical and Clinical Sciences L. Sacco, University of Milan | Alessia Lai et al |
| B.1.223 | EPI_ISL_418152 | Wales/PHWC-241A9/2020 | 2020/3/15 | Public Health Wales Microbiology Cardiff | Catherine Moore et al |
| B.1.224 | EPI_ISL_511642 | Portugal/PT1095/2020 | 2020/3/16 | Instituto Nacional de Saude (INSA) | Borges et al |
| B.1.225 | EPI_ISL_549484 | England/NORW-EFADC/2020 | 2020/4/1 | COVID-19 Genomics UK (COG-UK) Consortium | Dave J. Baker et al |
| B.1.227 | EPI_ISL_831975 | Sweden/20-23088/2020 | 2020/3/16 | The Public Health Agency of Sweden | Department of Microbiology et al |
| B.1.229 | EPI_ISL_1620714 | Ireland/KY-NVRL-20G33897/2020 | 2020/3/19 | Irish Coronavirus Sequencing Consortium - Teagasc Moorepark | Jose Sanchez-Morgado et al |

| | | | | | |
|---|---|---|---|---|---|
| B.1.23 | EPI_ISL_794712 | Australia/WA540/2020 | 2020/3/18 | PathWest Laboratory Medicine WA Microbial Surveillance Unit | PathWest Laboratory Medicine WA Microbial Surveillance Unit et al |
| B.1.231 | EPI_ISL_478763 | England/OXON-AF6C5/2020 | 2020/4/8 | COVID-19 Genomics UK (COG-UK) Consortium | Tanya Golubchik et al |
| B.1.232 | EPI_ISL_487893 | Scotland/EDB3155/2020 | 2020/4/6 | Wellcome Sanger Institute for the COVID-19 Genomics UK (COG-UK) consortium | McHugh M et al |
| B.1.233 | EPI_ISL_1017380 | USA/OR-PROV-398/2020 | 2020/6/16 | Providence St. Joseph Health Molecular Genomics Laboratory | Alexa K Dowdell et al |
| B.1.234 | EPI_ISL_694614 | USA/AZ-TG376210/2020 | 2020/6/15 | TGen North | Jolene Bowers et al |
| B.1.235 | EPI_ISL_553969 | England/ALDP-9562B1/2020 | 2020/6/8 | Wellcome Sanger Institute for the COVID-19 Genomics UK (COG-UK) consortium | The Lighthouse Lab in Alderley Park et al |
| B.1.237 | EPI_ISL_700567 | SouthAfrica/NHLS-UCT-GP-5183/2020 | 2020/4/24 | NHLS/UCT | Arash Iranzadeh et al |
| B.1.238 | EPI_ISL_533364 | England/QEUH-544733/2020 | 2020/5/14 | Wellcome Sanger Institute for the COVID-19 Genomics UK (COG-UK) consortium | Harper VanSteenhouse et al |
| B.1.239 | EPI_ISL_2712642 | Mexico/BCN-SEARCH-102575/2020 | 2020/4/9 | Andersen lab at Scripps Research | SEARCH Alliance with Gisela Barrera Badillo et al |
| B.1.240 | EPI_ISL_422555 | USA/NY-PV09200/2020 | 2020/3/22 | MSHS Pathogen Surveillance Program | Ana S. Gonzalez-Reiche et al |
| B.1.240 | EPI_ISL_1760240 | PuertoRico/PR-CDC-S298/2021 | 2021/1/19 | Centers for Disease Control and Prevention, Dengue Branch | Gilberto A. Santiago et al |
| B.1.240.1 | EPI_ISL_636514 | Aruba/AW-RIVM-10388/2020 | 2020/8/4 | National Institute for Public Health and the Environment (RIVM) | Adam Meijer et al |
| B.1.240.2 | EPI_ISL_969220 | Canada/BC-BCCDC-4087/2020 | 2020/8/24 | BCCDC Public Health Laboratory | Prystajecky Natalie et al |
| B.1.241 | EPI_ISL_2712687 | Mexico/BCN-SEARCH-102621/2020 | 2020/4/15 | Andersen lab at Scripps Research | SEARCH Alliance with Gisela Barrera Badillo et al |
| B.1.241 | EPI_ISL_1383041 | Spain/NC-CHN-01000218/2020 | 2020/3/24 | Centro de Secuenciación NASERTIC | Carmen Ezpeleta Baquedano et al |
| B.1.242 | EPI_ISL_3031179 | Norway/2783/2020 | 2020/6/12 | Norwegian Institute of Public Health, Department of Virology | Kathrine Stene-Johansen et al |
| B.1.243 | EPI_ISL_494714 | USA/CA-ALSR-1649/2020 | 2020/3/23 | Andersen lab at Scripps Research | SEARCH Alliance San Diego with Tracy Basler et al |
| B.1.243.1 | EPI_ISL_5628643 | USA/AZ-ASU18869/2021 | 2021/1/18 | Arizona State University | Ajeet Bains et al |

| | | | | | |
|---|---|---|---|---|---|
| B.1.245 | EPI_ISL_475618 | USA/CA-CSMC14/2020 | 2020/3/24 | Cedars-Sinai Medical Center, Molecular Pathology Laboratory of Department of Pathology & Laboratory Medicine and Genomic Core | Wenjuan Zhang et al |
| B.1.247 | EPI_ISL_444484 | India/GJ-GBRC53/2020 | 2020/5/4 | Gujarat Biotechnology Research Centre | Pranay Shah et al |
| B.1.248 | EPI_ISL_489620 | Scotland/GCVR-17176E/2020 | 2020/3/28 | Wellcome Sanger Institute for the COVID-19 Genomics UK (COG-UK) consortium | Ana da Silva Filipe et al |
| B.1.250 | EPI_ISL_425483 | England/NOTT-10E001/2020 | 2020/3/16 | COVID-19 Genomics UK (COG-UK) Consortium | Gemma Clark et al |
| B.1.251 | EPI_ISL_559053 | England/ALDP-52B8D5/2020 | 2020/6/1 | Wellcome Sanger Institute for the COVID-19 Genomics UK (COG-UK) consortium | The Lighthouse Lab in Alderley Park et al |
| B.1.252 | EPI_ISL_443908 | England/BRIS-1254AE/2020 | 2020/3/27 | Wellcome Sanger Institute for the COVID-19 Genomics UK (COG-UK) consortium | Stephanie Hutchings et al |
| B.1.254 | EPI_ISL_442230 | England/CAMB-7CEC9/2020 | 2020/4/6 | Wellcome Sanger Institute for the COVID-19 Genomics UK (COG-UK) consortium | Luke W Meredith et al |
| B.1.256 | EPI_ISL_489646 | Scotland/GCVR-16FD16/2020 | 2020/3/22 | Wellcome Sanger Institute for the COVID-19 Genomics UK (COG-UK) consortium | Ana da Silva Filipe et al |
| B.1.258 | EPI_ISL_423656 | England/20130069504/2020 | 2020/3/22 | Respiratory Virus Unit, Microbiology Services Colindale, Public Health England | Monica Galiano et al |
| B.1.258.10 | EPI_ISL_612125 | Scotland/CVR4698/2020 | 2020/9/20 | COVID-19 Genomics UK (COG-UK) Consortium | Ana da Silva Filipe et al |
| B.1.258.11 | EPI_ISL_615212 | Denmark/DCGC-5281/2020 | 2020/9/14 | Albertsen lab, Department of Chemistry and Bioscience, Aalborg University, Denmark | Danish Covid-19 Genome Consortia et al |
| B.1.258.14 | EPI_ISL_882747 | Italy/CAM-CRGS-241/2020 | 2020/10/6 | 1. Genome Research Center for Health (CRGS) / 2. Laboratory of Molecular Medicine and Genomics(LMMGe) / 3. Center for Research in Pure and Applied Mathematics (CRMPA) | Giorgio Giurato et al |
| B.1.258.15 | EPI_ISL_6230322 | Denmark/DCGC-207339/2020 | 2020/11/10 | Statens Serum Institut Bioinformatics and Microbial Genomics | Danish Covid-19 Genome Consortium et al |

| | | | | | |
|---|---|---|---|---|---|
| B.1.258.16 | EPI_ISL_711874 | Denmark/DCGC-14621/2020 | 2020/11/23 | Albertsen Lab, Department of Chemistry and Bioscience, Aalborg University, Denmark | Danish Covid-19 Genome Consortium et al |
| B.1.258.17 | EPI_ISL_1499000 | Slovenia/77414/2020 | 2020/8/28 | Institute of Microbiology and Immunology, Faculty of Medicine, University of Ljubljana | Alen Suljič et al |
| B.1.258.18 | EPI_ISL_722425 | Netherlands/ZH-EMC-1068/2020 | 2020/10/5 | Erasmus Medical Center | Bas Oude Munnink et al |
| B.1.258.19 | EPI_ISL_711361 | Denmark/DCGC-14599/2020 | 2020/11/23 | Albertsen Lab, Department of Chemistry and Bioscience, Aalborg University, Denmark | Danish Covid-19 Genome Consortium et al |
| B.1.258.2 | EPI_ISL_558199 | England/ALDP-672355/2020 | 2020/7/3 | Wellcome Sanger Institute for the COVID-19 Genomics UK (COG-UK) consortium | The Lighthouse Lab in Alderley Park et al |
| B.1.258.20 | EPI_ISL_815374 | Germany/BY-Cento-32458760/2020 | 2020/9/23 | Centogene | Peter Bauer et al |
| B.1.258.21 | EPI_ISL_632445 | Netherlands/FL-EMC-35/2020 | 2020/10/8 | Erasmus Medical Center | Bas Oude Munnink et al |
| B.1.258.22 | EPI_ISL_2898873 | Sweden/01_SE100_21CS503818/2020 | 2020/12/7 | Karolinska University Hospital | Jan Albert et al |
| B.1.258.23 | EPI_ISL_849870 | USA/UT-UPHL-2101980997/2020 | 2020/12/19 | Utah Public Health Laboratory | Erin L. Young et al |
| B.1.258.24 | EPI_ISL_804053 | Italy/FVG-ASUGI-152122/2020 | 2020/12/21 | ARGO Laboratorio Genomica ed Epigenomica | Licastro D et al |
| B.1.258.3 | EPI_ISL_557984 | England/MILK-622EBD/2020 | 2020/7/6 | Wellcome Sanger Institute for the COVID-19 Genomics UK (COG-UK) consortium | The Lighthouse Lab in Milton Keynes et al |
| B.1.258.4 | EPI_ISL_590381 | England/QEUH-9B5520/2020 | 2020/9/3 | Wellcome Sanger Institute for the COVID-19 Genomics UK (COG-UK) consortium | Harper VanSteenhouse et al |
| B.1.258.5 | EPI_ISL_572850 | England/ALDP-9BCEA6/2020 | 2020/9/9 | COVID-19 Genomics UK (COG-UK) Consortium | Tanya Golubchik et al |
| B.1.258.6 | EPI_ISL_567431 | England/QEUH-9C9F2B/2020 | 2020/9/10 | Wellcome Sanger Institute for the COVID-19 Genomics UK (COG-UK) consortium | Harper VanSteenhouse et al |
| B.1.258.7 | EPI_ISL_549868 | England/MILK-99D096/2020 | 2020/8/29 | Wellcome Sanger Institute for the COVID-19 Genomics UK (COG-UK) consortium | The Lighthouse Lab in Milton Keynes et al |

| | | | | | |
|---|---|---|---|---|---|
| B.1.258.9 | EPI_ISL_616087 | Denmark/DCGC-3333/2020 | 2020/8/31 | Albertsen lab, Department of Chemistry and Bioscience, Aalborg University, Denmark | Danish Covid-19 Genome Consortia et al |
| B.1.260 | EPI_ISL_678037 | SaudiArabia/KAUST-MADINAH665/2020 | 2020/3/26 | Pathogen Genomics Lab King Abdullah University of Science and Technology(KAUST) | Sara Mfarrej et al |
| B.1.263 | EPI_ISL_452032 | Denmark/ALAB-HH-157/2020 | 2020/3/19 | Albertsen lab, Department of Chemistry and Bioscience, Aalborg University, Denmark | Rasmus Kirkegaard et al |
| B.1.264 | EPI_ISL_604309 | USA/CA-QDX-2021/2020 | 2020/3/17 | Quest Diagnostics | Rosenthal et al |
| B.1.264.1 | EPI_ISL_733209 | Russia/AST-RII-MH4485S/2020 | 2020/8/26 | WHO National Influenza Centre Russian Federation | Andrey Komissarov et al |
| B.1.265 | EPI_ISL_571780 | USA/MN-QDX-981/2020 | 2020/3/15 | Quest Diagnostics | Rosenthal et al |
| B.1.267 | EPI_ISL_494727 | USA/CA-ALSR-1667/2020 | 2020/3/24 | Andersen lab at Scripps Research | SEARCH Alliance San Diego with Tracy Basler et al |
| B.1.268 | EPI_ISL_444022 | USA/TX_GCID_192000003/2020 | 2020/3/18 | Baylor College of Medicine: HGSC | Vasanthi Avadhanula et al |
| B.1.270 | EPI_ISL_586299 | Canada/ON-S1222/2020 | 2020/3/18 | McMaster University | Allison McGeer et al |
| B.1.273 | EPI_ISL_940393 | France/IDF_HB_112003050544/2020 | 2020/3/13 | IAME UMR1137 Inserm, Université de Paris, Hôpital Bichat | Antoine Bridier-Nahmias et al |
| B.1.274 | EPI_ISL_1001043 | Cameroon/Yaounde-20V-6189/2020 | 2020/4/13 | Institut Pasteur de Dakar | Njouom Richard et al |
| B.1.276 | EPI_ISL_434852 | USA/TX-HMH0153/2020 | 2020/3/22 | Houston Methodist Hospital | S. Wesley Long et al |
| B.1.277 | EPI_ISL_414637 | France/HDF-1993/2020 | 2020/3/4 | National Reference Center for Viruses of Respiratory Infections, Institut Pasteur, Paris | Mélnie Albert et al |
| B.1.280 | EPI_ISL_512564 | USA/FL-BPHL-0758/2020 | 2020/6/27 | Florida Bureau of Public Health Laboratories | Sarah Schmedes et al |
| B.1.281 | EPI_ISL_483563 | Bahrain/BAH-22/2020 | 2020/4/8 | Erasmus Medical Center | Bas Oude Munnink et al |
| B.1.282 | EPI_ISL_558971 | England/ALDP-52B87B/2020 | 2020/6/1 | Wellcome Sanger Institute for the COVID-19 Genomics UK (COG-UK) consortium | The Lighthouse Lab in Alderley Park et al |
| B.1.284 | EPI_ISL_420643 | England/20136094704/2020 | 2020/3/26 | Respiratory Virus Unit, Microbiology Services Colindale, Public Health England | Monica Galiano et al |
| B.1.284 | EPI_ISL_604936 | USA/LA-QDX-2624/2020 | 2020/3/16 | Quest Diagnostics | Rosenthal et al |
| B.1.285 | EPI_ISL_544039 | USA/TX-HMH-3163/2020 | 2020/6/10 | Houston Methodist Hospital | S. Wesley Long et al |

| | | | | | |
|---|---|---|---|---|---|
| B.1.287 | EPI_ISL_569689 | USA/TX-DSHS-0577/2020 | 2020/6/2 | Texas Department of State Health Services | Rashmi Tuladhar et al |
| B.1.289 | EPI_ISL_914632 | USA/AZ-TG655061/2020 | 2020/3/29 | TGen North | "Jolene Bowers et al |
| B.1.291 | EPI_ISL_1517403 | CostaRica/HNN-0359/2020 | 2020/5/31 | Inciensa, Instituto Costarricense de Investigación y Enseñanza en Nutrición y Salud | Cristian Peréz-Corrales et al |
| B.1.292 | EPI_ISL_427490 | USA/NY-NYUMC195/2020 | 2020/4/1 | Departments of Pathology and Medicine, New York University School of Medicine | Maria Aguero-Rosenfeld et al |
| B.1.293 | EPI_ISL_603938 | USA/FL-QDX-1620/2020 | 2020/3/16 | Quest Diagnostics | Rosenthal et al |
| B.1.294 | EPI_ISL_484840 | USA/WI-UW-465/2020 | 2020/6/16 | University of Wisconsin-Madison AIDS Vaccine Research Laboratories | Gage Moreno et al |
| B.1.298 | EPI_ISL_676923 | USA/NY-Wadsworth-96601-01/2020 | 2020/3/20 | Wadsworth Center, New York State Department.of Health | Kirsten St. George et al |
| B.1.3 | EPI_ISL_4025997 | USA/IL-NM-8510/2020 | 2020/4/18 | Northwestern University - Center for Pathogen Genomics and Microbial Evolution | Ramon Lorenzo-Redondo et al |
| B.1.301 | EPI_ISL_493063 | USA/WY-WYPHL-20007007/2020 | 2020/4/1 | Wyoming Public Health Laboratory | Noah Hull et al |
| B.1.302 | EPI_ISL_631571 | USA/NY-NYCPHL-000078/2020 | 2020/3/12 | New York City Public Health Laboratory | Jade Wang et al |
| B.1.304 | EPI_ISL_484886 | USA/WI-UW-511/2020 | 2020/6/5 | University of Wisconsin-Madison AIDS Vaccine Research Laboratories | Gage Moreno et al |
| B.1.305 | EPI_ISL_1321842 | USA/TX-DSHS-4209/2020 | 2020/6/9 | TXDSHS | Rashmi Tuladhar et al |
| B.1.306 | EPI_ISL_594313 | USA/FL-BPHL-1623/2020 | 2020/6/3 | Florida Bureau of Public Health Laboratories | Sarah Schmedes et al |
| B.1.308 | EPI_ISL_805923 | Canada/AB-35896/2020 | 2020/3/22 | Alberta Precision Labs (APL) | Gordon P et al |
| B.1.309 | EPI_ISL_576529 | USA/CA-IGI-0340/2020 | 2020/6/29 | Innovative Genomics Institute, UC Berkeley | Stacia Wyman et al |
| B.1.310 | EPI_ISL_447404 | Israel/2046434/2020 | 2020/4/1 | Stern Lab | Stern Lab et al |
| B.1.311 | EPI_ISL_578151 | USA/MI-UM-10033882288/2020 | 2020/4/8 | Lauring Lab, University of Michigan, Department of Microbiology and Immunology | Valesano et al |
| B.1.313 | EPI_ISL_696019 | USA/AZ-TG654719/2020 | 2020/4/1 | TGen North | Jolene Bowers et al |
| B.1.314 | EPI_ISL_421354 | USA/NY-PV08125/2020 | 2020/3/13 | MSHS Pathogen Surveillance Program | Ana S. Gonzalez-Reiche et al |
| B.1.315 | EPI_ISL_428202 | USA/NE-875/2020 | 2020/3/25 | UNMC COVID-19 Response Team | UNMC COVID-19 Response Team et al |
| B.1.316 | EPI_ISL_586325 | Canada/ON-S1274/2020 | 2020/4/29 | McMaster University | Allison McGeer et al |

| | | | | | |
|---|---|---|---|---|---|
| B.1.318 | EPI_ISL_514188 | USA/FL-BPHL-0842/2020 | 2020/6/22 | Florida Bureau of Public Health Laboratories | Sarah Schmedes et al |
| B.1.319 | EPI_ISL_482469 | Canada/NS_40/2020 | 2020/3/5 | National Microbiology Laboratory | Anna Majer et al |
| B.1.320 | EPI_ISL_424890 | USA/UT-CDC-03068786-001/2020 | 2020/3/7 | Pathogen Discovery, Respiratory Viruses Branch, Division of Viral Diseases, Centers for Disease Control and Prevention | Yan Li et al |
| B.1.321 | EPI_ISL_418985 | Belgium/LT-030956/2020 | 2020/3/9 | KU Leuven, Clinical and Epidemiological Virology | Bert Vanmechelen et al |
| B.1.323 | EPI_ISL_450111 | USA/NY-PV09434/2020 | 2020/3/13 | MSHS Pathogen Surveillance Program | Ana S. Gonzalez-Reiche et al |
| B.1.324 | EPI_ISL_467906 | USA/TX-QDX-25/2020 | 2020/3/15 | Quest Diagnostics | Anderson et al |
| B.1.325 | EPI_ISL_545312 | USA/TX-HMH-2445/2020 | 2020/6/1 | Houston Methodist Hospital | S. Wesley Long et al |
| B.1.326 | EPI_ISL_542539 | USA/TX-HMH-1113/2020 | 2020/4/25 | Houston Methodist Hospital | S. Wesley Long et al |
| B.1.328 | EPI_ISL_509478 | USA/MD-MDH-0074/2020 | 2020/5/23 | Maryland Department of Health | Keller et al |
| B.1.329 | EPI_ISL_2869924 | Germany/NW-KRO-463/2020 | 2020/4/9 | Center of Medical Microbiology, Virology, and Hospital Hygiene, University of Duesseldorf | Dennis Deschka et al |
| B.1.330 | EPI_ISL_483013 | USA/MN-MDH-1293/2020 | 2020/3/24 | Minnesota Department of Health, Public Health Laboratory | Matt Plumb et al |
| B.1.332 | EPI_ISL_632050 | USA/NY-NYCPHL-000592/2020 | 2020/3/10 | New York City Public Health Laboratory | Jade Wang et al |
| B.1.333 | EPI_ISL_604460 | USA/MI-QDX-2087/2020 | 2020/3/17 | Quest Diagnostics | Rosenthal et al |
| B.1.334 | EPI_ISL_632146 | USA/NY-NYCPHL-000766/2020 | 2020/3/20 | New York City Public Health Laboratory | Jade Wang et al |
| B.1.335 | EPI_ISL_467407 | USA/NY-NYUMC914/2020 | 2020/5/8 | Departments of Pathology and Medicine, New York University School of Medicine | Maria Aguero-Rosenfeld et al |
| B.1.336 | EPI_ISL_468562 | USA/CA-QDX-110/2020 | 2020/3/13 | Quest Diagnostics | Anderson et al |
| B.1.337 | EPI_ISL_2727987 | USA/WY-WYPHL-20037947/2020 | 2020/7/5 | Wyoming Public Health Laboratory | Jim Mildenberger et al |
| B.1.338 | EPI_ISL_604872 | USA/WA-QDX-2548/2020 | 2020/3/16 | Quest Diagnostics | Rosenthal et al |
| B.1.340 | EPI_ISL_3726054 | USA/TN-VUMC-000080/2020 | 2020/3/13 | Dr. Suman Das Lab - Vanderbilt University Medical Center (VUMC) (https://my.vanderbilt.edu/daslab/) | Grant Vestal et al |
| B.1.341 | EPI_ISL_570200 | USA/WA-UW-8165/2020 | 2020/4/25 | UW Virology Lab | Pavitra Roychoudhury et al |
| B.1.342 | EPI_ISL_510834 | Sweden/20-51697/2020 | 2020/5/23 | The Public Health Agency of Sweden | Oskar Karlsson Lindsjo et al |
| B.1.343 | EPI_ISL_429376 | Denmark/ALAB-SSI160/2020 | 2020/3/9 | Albertsen lab, Department of Chemistry and Bioscience, Aalborg University, Denmark | Rasmus Kirkegaard et al |

| | | | | | |
|---|---|---|---|---|---|
| B.1.344 | EPI_ISL_2549656 | USA/CO-CDPHE-2003120688/2020 | 2020/3/12 | Colorado Department of Public Health and Environment | Laura Bankers et al |
| B.1.346 | EPI_ISL_1973129 | Japan/Donner270/2020 | 2020/4/13 | Keio University School of Medicine | Kenjiro Kosaki et al |
| B.1.349 | EPI_ISL_1241045 | Canada/QC-JEW-Q8292157/2020 | 2020/4/29 | Laboratoire de santé publique du Québec | Sandrine Moreira et al |
| B.1.35 | EPI_ISL_514566 | Wales/PHWC-169A15/2020 | 2020/3/20 | COVID-19 Genomics UK (COG-UK) Consortium | Catherine Moore et al |
| B.1.350 | EPI_ISL_452327 | Beijing/DT-travelAT01/2020 | 2020/3/13 | Laboratory of Infectious Diseases Center of Beijing Ditan Hospital | Siyuan Yang et al |
| B.1.350.1 | EPI_ISL_512433 | England/PORT-2F5DE7/2020 | 2020/5/25 | COVID-19 Genomics UK (COG-UK) Consortium | Angela Beckett et al |
| B.1.351 | EPI_ISL_2493065 | Zimbabwe/CERI-KRISP-K011662/2020 | 2020/5/27 | CERI, Centre for Epidemic Response and Innvoation, Stellenbosch University and KRISP, KZN Research Innovation and Sequencing Platform, UKZN. | Air Comodor Dr J. Chimedza et al |
| B.1.351.2 | EPI_ISL_736966 | SouthAfrica/KRISP-MDSH920866/2020 | 2020/12/6 | KRISP, KZN Research Innovation and Sequencing Platform | Giandhari J et al |
| B.1.351.3 | EPI_ISL_943561 | Bangladesh/BCSIR-NILMRC-448/2021 | 2021/1/24 | Genomic Research Lab, BCSIR | Tanjina Akhtar Banu et al |
| B.1.351.5 | EPI_ISL_1700678 | EquatorialGuinea/81769/2021 | 2021/1/14 | Swiss Tropical and Public Health Institute | Salome Hosch et al |
| B.1.354 | EPI_ISL_418228 | France/HDF-2405/2020 | 2020/3/12 | National Reference Center for Viruses of Respiratory Infections, Institut Pasteur, Paris | Mélanie Albert et al |
| B.1.355 | EPI_ISL_449800 | Chile/RM-ISPCH-53/2020 | 2020/4/1 | Instituto de Salud Publica de Chile | Andrés E Castillo et al |
| B.1.356 | EPI_ISL_418206 | Senegal/003/2020 | 2020/2/28 | Institut Pasteur de Dakar | Ndongo Dia et al |
| B.1.357 | EPI_ISL_1017469 | USA/OR-PROV-535/2020 | 2020/5/21 | Providence St. Joseph Health Molecular Genomics Laboratory | Alexa K Dowdell et al |
| B.1.358 | EPI_ISL_437630 | Denmark/ALAB-SSI-1046/2020 | 2020/4/1 | Albertsen lab, Department of Chemistry and Bioscience, Aalborg University, Denmark | Rasmus Kirkegaard et al |
| B.1.359 | EPI_ISL_525612 | USA/NY-Wadsworth-14422-01/2020 | 2020/3/9 | Wadsworth Center, New York State Department of Health | Kirsten St. George et al |
| B.1.36 | EPI_ISL_490003 | SaudiArabia/562/2020 | 2020/2/16 | King Fahad Medical City | Alosaimi et al |
| B.1.36.1 | EPI_ISL_490042 | Australia/NSW633/2020 | 2020/7/1 | NSW Health Pathology - Institute of Clinical Pathology and Medical Research; Westmead Hospital; University of Sydney | CIDM-PH et al. |

| | | | | | |
|---|---|---|---|---|---|
| B.1.36.10 | EPI_ISL_513168 | SaudiArabia/KAUST-MADINAH599/2020 | 2020/4/30 | Pathogen Genomics Lab King Abdullah University of Science and Technology(KAUST) | Sara Mfarrej et al |
| B.1.36.12 | EPI_ISL_558361 | England/MILK-566C5A/2020 | 2020/6/12 | Wellcome Sanger Institute for the COVID-19 Genomics UK (COG-UK) consortium | The Lighthouse Lab in Milton Keynes et al |
| B.1.36.16 | EPI_ISL_450344 | Bangladesh/BARJ-CVASU-CTG-517/2020 | 2020/5/3 | Basic and Applied Research on Jute Project | Rasel Ahmed et al |
| B.1.36.17 | EPI_ISL_533016 | Scotland/QEUH-789563/2020 | 2020/7/16 | Wellcome Sanger Institute for the COVID-19 Genomics UK (COG-UK) consortium | Ana da Silva Filipe et al |
| B.1.36.18 | EPI_ISL_3276881 | India/DL-ILBS-17864/2020 | 2020/6/5 | ILBS | Ekta Gupta et al |
| B.1.36.19 | EPI_ISL_956311 | Indonesia/JI-ITDua-17047NTv/2020 | 2020/7/16 | Institute of Tropical Disease, Universitas Airlangga | Kazufumi Shimizu et al |
| B.1.36.2 | EPI_ISL_638446 | England/CAMC-A65DDB/2020 | 2020/10/11 | COVID-19 Genomics UK (COG-UK) Consortium | Aminu S. Jahun et al |
| B.1.36.20 | EPI_ISL_534634 | Scotland/QEUH-96D396/2020 | 2020/8/20 | Wellcome Sanger Institute for the COVID-19 Genomics UK (COG-UK) Consortium | Ana da Silva Filipe et al |
| B.1.36.21 | EPI_ISL_775318 | Norway/4694/2020 | 2020/10/7 | Norwegian Institute of Public Health, Department of Virology | Kathrine Stene-Johansen et al |
| B.1.36.22 | EPI_ISL_1586054 | Iceland/926/2020 | 2020/4/14 | deCODE genetics | Daniel F Gudbjartsson et al |
| B.1.36.23 | EPI_ISL_4482150 | Netherlands/ZH-EMC-3480/2020 | 2020/5/8 | Erasmus Medical Center | Bas Oude Munnink et al |
| B.1.36.24 | EPI_ISL_1751685 | Germany/BY-MVP-000002132/2020 | 2020/10/27 | Laboratory for Functional Genome Analysis; Dept. Genomics; Gene Center of the LMU Munich | Max Muenchhoff et al |
| B.1.36.25 | EPI_ISL_618931 | Denmark/DCGC-7478/2020 | 2020/10/12 | Albertsen lab, Department of Chemistry and Bioscience, Aalborg University, Denmark | Danish Covid-19 Genome Consortia et al |
| B.1.36.26 | EPI_ISL_876599 | Canada/ON-SC0258/2020 | 2020/8/26 | McMaster University | Allison McGeer et al |
| B.1.36.27 | EPI_ISL_760032 | HongKong/VM20123554/2020 | 2020/9/18 | School of Public Health, The University of Hong Kong | Daniel Chu et al |
| B.1.36.28 | EPI_ISL_566961 | England/MILK-9C3E9D/2020 | 2020/9/9 | Wellcome Sanger Institute for the COVID-19 Genomics UK (COG-UK) consortium | The Lighthouse Lab in Milton Keynes et al |
| B.1.36.29 | EPI_ISL_862465 | India/AP-CS0392/2020 | 2020/6/27 | CSIR Institute of Genomics and Integrative Biology | Pallavali Roja Rani et al |

| | | | | | |
|---|---|---|---|---|---|
| B.1.36.31 | EPI_ISL_1229177 | Australia/SA0633/2020 | 2020/3/8 | SA Pathology | Lex Leong et al |
| B.1.36.33 | EPI_ISL_596752 | Australia/WA189/2020 | 2020/5/23 | PathWest Laboratory Medicine WA Microbial Surveillance Unit | PathWest Laboratory Medicine WA Microbial Surveillance Unit et al |
| B.1.36.34 | EPI_ISL_569969 | Canada/ON-UHTC-0199/2020 | 2020/6/10 | Ontario Institute for Cancer Research | Ramzi Fattouh et al |
| B.1.36.35 | EPI_ISL_1009016 | Lebanon/LAU-R87/2020 | 2020/9/1 | School of Pharmacy | Ahmed Kandeil et al |
| B.1.36.36 | EPI_ISL_483663 | Switzerland/GE-ETHZ-150035/2020 | 2020/6/3 | Department of Biosystems Science and Engineering, ETH Zürich | Christian Beisel et al |
| B.1.36.37 | EPI_ISL_4421463 | Canada/BC-BCCDC-35581/2020 | 2020/10/6 | BCCDC Public Health Laboratory | Prystajecky Natalie et al |
| B.1.36.38 | EPI_ISL_1109628 | Egypt/MASRI-C5-013/2020 | 2020/6/8 | Medical Ain Shams Research Institute (MASRI), Ain Shams University | Hesham Elghazaly et al |
| B.1.36.39 | EPI_ISL_584847 | England/QEUH-9E3495/2020 | 2020/9/21 | COVID-19 Genomics UK (COG-UK) Consortium | Darren L Smith et al |
| B.1.36.7 | EPI_ISL_1014687 | Iran/Tehran-055M/2020 | 2020/8/17 | National Influenza Center | J Yavarian et al |
| B.1.36.8 | EPI_ISL_450783 | India/GJ-GBRC95b/2020 | 2020/4/11 | Gujarat Biotechnology Research Centre | Shirish Patel et al |
| B.1.36.9 | EPI_ISL_513222 | SaudiArabia/KAUST-MAKKAH301/2020 | 2020/4/21 | Pathogen Genomics Lab King Abdullah University of Science and Technology(KAUST) | Sharif Hala et al |
| B.1.360 | EPI_ISL_437943 | Austria/CeMM0074/2020 | 2020/3/17 | Bergthaler laboratory, CeMM Research Center for Molecular Medicine of the Austrian Academy of Sciences | Alexandra Popa et al |
| B.1.361 | EPI_ISL_602312 | USA/FL-B2TM/2020 | 2020/8/2 | University of Miami Immunology and Histocompatibility Laboratory | Emilio Margolles-Clark et al |
| B.1.362 | EPI_ISL_447427 | Israel/13077723/2020 | 2020/4/3 | Stern Lab | Stern Lab et al |
| B.1.362.1 | EPI_ISL_601611 | England/CAMC-9D2095/2020 | 2020/9/19 | Wellcome Sanger Institute for the COVID-19 Genomics UK (COG-UK) consortium | Rob Howes et al |
| B.1.362.2 | EPI_ISL_575332 | Israel/CVL-s2049/2020 | 2020/9/1 | Israel Central Virology laboratory | Neta Zuckerman et al |
| B.1.363 | EPI_ISL_3009337 | USA/AZ-TG968075/2020 | 2020/9/3 | TGen North | Jolene Bowers et al |
| B.1.366 | EPI_ISL_494543 | USA/TX-QDX-188/2020 | 2020/4/27 | Quest Diagnostics | Anderson et al |
| B.1.367 | EPI_ISL_510700 | Switzerland/GE-ETHZ-200011/2020 | 2020/7/13 | Department of Biosystems Science and Engineering, ETH Zürich | Christian Beisel et al |
| B.1.369 | EPI_ISL_1591225 | USA/NE-CDC-3960290-001/2020 | 2020/3/11 | Pathogen Discovery, Respiratory Viruses Branch, Division of Viral Diseases, Centers for Disease Control and Prevention | Yan Li et al |

| | | | | | |
|---|---|---|---|---|---|
| B.1.369.1 | EPI_ISL_961578 | Canada/NB-NML-3272/2020 | 2020/12/29 | National Microbiology Laboratory (NML) | Anna Majer et al |
| B.1.37 | EPI_ISL_452112 | USA/TX-CDC-03070108-001/2020 | 2020/3/10 | Pathogen Discovery, Respiratory Viruses Branch, Division of Viral Diseases, Centers for Disease Control and Prevention | Yan Li et al |
| B.1.370 | EPI_ISL_417971 | USA/UT-028/2020 | 2020/3/19 | Utah Public Health Laboratory | Erin Young et al |
| B.1.371 | EPI_ISL_414616 | USA/WA-UW29/2020 | 2020/3/8 | UW Virology Lab | Pavitra Roychoudhury et al |
| B.1.372 | EPI_ISL_622596 | Denmark/DCGC-1846/2020 | 2020/3/30 | Albertsen lab, Department of Chemistry and Bioscience, Aalborg University, Denmark | Danish Covid-19 Genome Consortia et al |
| B.1.375 | EPI_ISL_976921 | USA/MA-Broad_UMMS-00165/2020 | 2020/9/14 | Infectious Disease Program, Broad Institute of Harvard and MIT | Lemieux et al |
| B.1.377 | EPI_ISL_1392875 | USA/MO-MSPHL-000515/2020 | 2020/4/19 | Missouri State Public Health Laboratory | Matthew Sinn et al |
| B.1.378 | EPI_ISL_571600 | USA/IL-QDX-831/2020 | 2020/3/14 | Quest Diagnostics | Rosenthal et al |
| B.1.379 | EPI_ISL_440220 | England/BRIS-12193A/2020 | 2020/3/16 | Wellcome Sanger Institute for the COVID-19 Genomics UK (COG-UK) consortium | Stephanie Hutchings et al |
| B.1.38 | EPI_ISL_418954 | USA/WA-UW389/2020 | 2020/3/17 | UW Virology Lab | Pavitra Roychoudhury et al |
| B.1.380 | EPI_ISL_960302 | Rwanda/NRLNAT2054/2020 | 2020/5/27 | GIGA Medical Genomics | Yvan Butera et al |
| B.1.381 | EPI_ISL_3068006 | SouthAfrica/Tygerberg_60/2020 | 2020/4/14 | Division of Medical Virology, Stellenbosch University and NHLS Tygerberg Hospital | Susan Engelbrecht et al |
| B.1.382 | EPI_ISL_590777 | USA/MN-MDH-1774/2020 | 2020/4/13 | Minnesota Department of Health, Public Health Laboratory | Matt Plumb et al |
| B.1.383 | EPI_ISL_413588 | Netherlands/Utrecht_1363564/2020 | 2020/3/1 | Erasmus Medical Center | David Nieuwenhuijse et al |
| B.1.384 | EPI_ISL_509587 | USA/UT-03775/2020 | 2020/4/1 | Utah Public Health Laboratory | Heidi Butz et al |
| B.1.385 | EPI_ISL_676618 | USA/TX-DSHS-1485/2020 | 2020/6/22 | Texas Department of State Health Services | Rashmi Tuladhar et al |
| B.1.387 | EPI_ISL_416756 | France/ARA-06531/2020 | 2020/3/6 | CNR Virus des Infections Respiratoires - France SUD | Bal et al |
| B.1.388 | EPI_ISL_614355 | Coted-Ivoire/BKE0125/2020 | 2020/6/10 | Project group Epidemology of Highly Pathogenic Microorganisms, Robert Koch-Institute | Chantal Akoua-Koffi et al |

| | | | | | |
|---|---|---|---|---|---|
| B.1.389 | EPI_ISL_618138 | Denmark/DCGC-1568/2020 | 2020/6/15 | Albertsen lab, Department of Chemistry and Bioscience, Aalborg University, Denmark | Danish Covid-19 Genome Consortia et al |
| B.1.39 | EPI_ISL_614577 | Denmark/DCGC-979/2020 | 2020/3/9 | Albertsen lab, Department of Chemistry and Bioscience, Aalborg University, Denmark | Danish Covid-19 Genome Consortia et al |
| B.1.390 | EPI_ISL_460122 | USA/MA-MGH-00080/2020 | 2020/3/25 | Infectious Disease Program, Broad Institute of Harvard and MIT | Lemieux et al |
| B.1.391 | EPI_ISL_464737 | England/201140585/2020 | 2020/3/10 | Respiratory Virus Unit, Microbiology Services Colindale, Public Health England | PHE Covid Sequencing Team et al |
| B.1.393 | EPI_ISL_2779421 | Kenya/BUS-406/2020 | 2020/5/29 | USAMRD-A, Basic Science Laboratory | Gathii Kimita et al |
| B.1.395 | EPI_ISL_415435 | Wales/PHW06/2020 | 2020/3/6 | Public Health Wales Microbiology Cardiff | Catherine Moore et al |
| B.1.396 | EPI_ISL_2596864 | France/PAC-IHU-10794_Illu1/2020 | 2020/4/6 | MEPHI, Aix Marseille University | Anthony LEVASSEUR et al |
| B.1.397 | EPI_ISL_2712631 | Mexico/BCN-SEARCH-102564/2020 | 2020/4/7 | Andersen lab at Scripps Research | SEARCH Alliance with Gisela Barrera Badillo et al |
| B.1.398 | EPI_ISL_512998 | SaudiArabia/KAUST-JEDDAH755/2020 | 2020/4/2 | Pathogen Genomics Lab King Abdullah University of Science and Technology(KAUST) | Amit Kumar Subudhi et al |
| B.1.399 | EPI_ISL_483449 | USA/CA-ALSR-0993-SAN/2020 | 2020/4/26 | Andersen lab at Scripps Research | SEARCH Alliance San Diego with David Pride et al |
| B.1.40 | EPI_ISL_425906 | Scotland/EDB124/2020 | 2020/3/18 | COVID-19 Genomics UK (COG-UK) Consortium | McHugh M et al |
| B.1.400 | EPI_ISL_483241 | USA/CA-ALSR-0585-SAN/2020 | 2020/4/15 | Andersen lab at Scripps Research | SEARCH Alliance San Diego with David Pride et al |
| B.1.400.1 | EPI_ISL_2423567 | USA/AZ-TG812013/2020 | 2020/11/9 | TGen North | "Jolene Bowers et al |
| B.1.401 | EPI_ISL_636164 | USA/CA-ALSR-3720/2020 | 2020/6/4 | Andersen lab at Scripps Research | SEARCH Alliance San Diego with Tracy Basler et al |
| B.1.403 | EPI_ISL_1137163 | USA/CA-CZB-16547/2020 | 2020/7/13 | Chan-Zuckerberg Biohub | CZB Clihub Consortium et al |
| B.1.404 | EPI_ISL_1523526 | USA/OR-OSPHL00196/2020 | 2020/3/16 | Oregon State Public Health Laboratory | LaDonna Grenz et al |
| B.1.405 | EPI_ISL_483281 | USA/CA-ALSR-0653-IPL/2020 | 2020/4/30 | Andersen lab at Scripps Research | SEARCH Alliance San Diego with David Pride et al |
| B.1.406 | EPI_ISL_618273 | Denmark/DCGC-1957/2020 | 2020/7/6 | Albertsen lab, Department of Chemistry and Bioscience, Aalborg University, Denmark | Danish Covid-19 Genome Consortia et al |

| | | | | | |
|---|---|---|---|---|---|
| B.1.407 | EPI_ISL_554831 | England/MILK-9586FF/2020 | 2020/6/9 | Wellcome Sanger Institute for the COVID-19 Genomics UK (COG-UK) consortium | The Lighthouse Lab in Milton Keynes et al |
| B.1.408 | EPI_ISL_455469 | Romania/283584/2020 | 2020/5/12 | Cantacuzino Institute | M.Lazar et al |
| B.1.409 | EPI_ISL_5384435 | USA/TX-HMH-MCoV-49655/2020 | 2020/7/13 | Houston Methodist Hospital | Randall J. Olsen et al |
| B.1.411 | EPI_ISL_602574 | SriLanka/GQC12/2020 | 2020/10/23 | Centre for Dengue Research, Department of Immunology and Molecular Medicine | Chandima Jeewandara et al |
| B.1.413 | EPI_ISL_604591 | USA/UT-QDX-2230/2020 | 2020/3/12 | Quest Diagnostics | Rosenthal et al |
| B.1.415 | EPI_ISL_801588 | Chile/Santiago-op7d1/2020 | 2020/3/16 | MSHS Pathogen Surveillance Program | Leonardo I. Almonacid et al |
| B.1.416 | EPI_ISL_480788 | Senegal/1938/2020 | 2020/3/31 | Institut Pasteur de Dakar | Ndongo Dia et al |
| B.1.416.1 | EPI_ISL_560598 | France/BRE-8930/2020 | 2020/7/24 | National Reference Center for Viruses of Respiratory Infections, Institut Pasteur, Paris | Sylvie Behillil et al |
| B.1.417 | EPI_ISL_509289 | SouthAfrica/KRISP-K000640/2020 | 2020/7/13 | KRISP, KZN Research Innovation and Sequencing Platform | Giandhari J et al |
| B.1.418 | EPI_ISL_439237 | Scotland/EDB2599/2020 | 2020/4/24 | COVID-19 Genomics UK (COG-UK) Consortium | McHugh M et al |
| B.1.420 | EPI_ISL_498154 | Colombia/CUN-INS-88496/2020 | 2020/3/25 | Instituto Nacional de Salud, Bogotá, Colombia | Katherine Laiton-Donato et al |
| B.1.421 | EPI_ISL_2628353 | USA/CA-SEARCH-101735/2020 | 2020/7/28 | Andersen lab at Scripps Research | SEARCH Alliance San Diego with Ashleigh Murphy et al |
| B.1.422 | EPI_ISL_464051 | Canada/ON-UHTC_0004/2020 | 2020/3/18 | Ontario Institute for Cancer Research | Ramzi Fattouh et al |
| B.1.423 | EPI_ISL_579013 | USA/LA-EVTL609/2020 | 2020/7/1 | Microbial Genome Sequencing Center | Jeremy P. Kamil et al |
| B.1.424 | EPI_ISL_604071 | USA/MN-QDX-1722/2020 | 2020/3/16 | Quest Diagnostics | Rosenthal et al |
| B.1.425 | EPI_ISL_524337 | USA/UT-08586/2020 | 2020/5/7 | Utah Public Health Laboratory | Erin L. Young et al |
| B.1.426 | EPI_ISL_449837 | USA/UT-01607/2020 | 2020/4/6 | Utah Public Health Laboratory | Erin Young et al |
| B.1.427 | EPI_ISL_2304313 | USA/CA-ACPHD-00310/2020 | 2020/4/11 | Alameda County Public Health Department | Kristna Hsieh et al |
| B.1.428 | EPI_ISL_415648 | Denmark/SSI-104/2020 | 2020/3/3 | ViFU | Morten Rasmussen et al |
| B.1.428.1 | EPI_ISL_812258 | Iraq/USAFSAM-S061/2020 | 2020/6/6 | United States Air Force School of Aerospace Medicine | Anthony Fries et al |

| | | | | | |
|---|---|---|---|---|---|
| B.1.428.2 | EPI_ISL_803817 | Tunisia/12-8274/2020 | 2020/4/7 | Clinical and Experimental Pharmacology Lab, LR16SP02, National Center of Pharmacovigilance, University of Tunis El Manar, Tunis, Tunisia. 2- Neurodegenerative diseases and psychiatric troubles, LR18SP03, Razi Hospital, University of Tunis El Manar, Tunis, Tunisia. 3- Ministry of Health, National Observatory of New and Emerging Diseases, 1006, Tunis, Tunisia | Ilhem Boutiba-Ben Boubaker et al |
| B.1.428.3 | EPI_ISL_1714216 | Qatar/QA-WCMQ_FD17462393/2020 | 2020/6/4 | Weill Cornell Medical College - Qatar (WCM-Q), Genomics Core Laboratory / Qatar Genome Project (QGP) | WCMQ: Ayeda A. Ahmed et al |
| B.1.429 | EPI_ISL_1675148 | USA/NV-NSPHL-357909/2020 | 2020/4/15 | Nevada State Public Health Laboratory | Andrew Gorzalski et al |
| B.1.429.1 | EPI_ISL_1708775 | USA/NY-MSHSPSP-PV24036/2020 | 2020/12/2 | MSHS Pathogen Surveillance Program | Ana S. Gonzalez-Reiche et al |
| B.1.431 | EPI_ISL_489039 | Scotland/EDB2810/2020 | 2020/3/30 | Wellcome Sanger Institute for the COVID-19 Genomics UK (COG-UK) consortium | McHugh M et al |
| B.1.432 | EPI_ISL_5254069 | USA/CO-CDPHE-2004270037/2020 | 2020/4/25 | Colorado Department of Public Health and Environment | Laura Bankers et al |
| B.1.433 | EPI_ISL_2223502 | USA/TX-HMH-MCoV-40320/2020 | 2020/8/3 | Houston Methodist Hospital | Randall J. Olsen et al |
| B.1.434 | EPI_ISL_426677 | Australia/VIC369/2020 | 2020/3/22 | Microbiological Diagnostic Unit Public Health Laboratory and Victorian Infectious Diseases Reference Laboratory, Doherty Institute | Caly L. et al |
| B.1.435 | EPI_ISL_695939 | USA/AZ-TG648768/2020 | 2020/4/10 | TGen North | Jolene Bowers et al |
| B.1.436 | EPI_ISL_524895 | USA/MD-MDH-0130/2020 | 2020/7/8 | MD PHL | Maryland Department of Health Laboratories Administration et al |
| B.1.437 | EPI_ISL_653200 | USA/FL-BPHL-2047/2020 | 2020/6/23 | Florida Bureau of Public Health Laboratories | Sarah Schmedes et al |
| B.1.438 | EPI_ISL_478710 | Australia/NSW598/2020 | 2020/6/8 | NSW Health Pathology - Institute of Clinical Pathology and Medical Research; Westmead Hospital; University of Sydney | CIDM-PH et al. |

| | | | | | |
|---|---|---|---|---|---|
| B.1.438.2 | EPI_ISL_855367 | Mayotte/IPP00174/2021 | 2021/1/6 | National Reference Center for Viruses of Respiratory Infections, Institut Pasteur, Paris | Marion Barbet et al |
| B.1.438.3 | EPI_ISL_1303308 | USA/SC-DHEC-0307/2021 | 2021/2/1 | Microbiology Division, South Carolina Department of Health and Environmental Control | Flores et al |
| B.1.438.4 | EPI_ISL_5696417 | USA/CA-CDPH-MC1205035/2020 | 2020/10/19 | California Department of Public Health | Emily Smith on behalf of CDPH-COVIDNet et al |
| B.1.439 | EPI_ISL_604495 | USA/UT-QDX-2129/2020 | 2020/3/13 | Quest Diagnostics | Rosenthal et al |
| B.1.44 | EPI_ISL_421005 | Wales/PHWC-25560/2020 | 2020/3/23 | Public Health Wales Microbiology Cardiff | Catherine Moore et al |
| B.1.441 | EPI_ISL_3726044 | USA/TN-VUMC-000070/2020 | 2020/3/13 | Dr. Suman Das Lab - Vanderbilt University Medical Center (VUMC) (https://my.vanderbilt.edu/daslab/) | Grant Vestal et al |
| B.1.442 | EPI_ISL_485841 | USA/VA-DCLS-0540/2020 | 2020/3/12 | Virginia DCLS | Virginia DCLS et al |
| B.1.443 | EPI_ISL_1168366 | USA/NH-Yale-1640/2020 | 2020/4/18 | Grubaugh Lab - Yale School of Public Health | Mary Petrone et al |
| B.1.444 | EPI_ISL_525343 | USA/UT-UPHL-04478/2020 | 2020/5/21 | Utah Public Health Laboratory | Erin Young et al |
| B.1.445 | EPI_ISL_696360 | USA/AZ-TG670160/2020 | 2020/4/5 | TGen North | Jolene Bowers et al |
| B.1.446 | EPI_ISL_424174 | USA/CT-UW-1365/2020 | 2020/3/13 | UW Virology Lab | Pavitra Roychoudhury et al |
| B.1.448 | EPI_ISL_826613 | USA/NY-AECOM_031/2020 | 2020/4/15 | Albert Einstein College of Medicine, Dept. of Microbiology & Immunology, Chandran lab | J. Maximilian Fels et al |
| B.1.450 | EPI_ISL_434718 | USA/TX-HMH07/2020 | 2020/3/15 | Houston Methodist Hospital | S. Wesley Long et al |
| B.1.451 | EPI_ISL_500846 | USA/VA-DCLS-0684/2020 | 2020/4/18 | Virginia DCLS | Virginia DCLS et al |
| B.1.452 | EPI_ISL_513791 | USA/CA-CZB-2341/2020 | 2020/5/12 | Chan-Zuckerberg Biohub | CZB Cliahub Consortium et al |
| B.1.452 | EPI_ISL_2224736 | USA/TX-HMH-MCoV-41000/2020 | 2020/7/21 | Houston Methodist Hospital | Randall J. Olsen et al |
| B.1.453 | EPI_ISL_571702 | USA/CA-QDX-1037/2020 | 2020/3/17 | Quest Diagnostics | Rosenthal et al |
| B.1.456 | EPI_ISL_490007 | SaudiArabia/610/2020 | 2020/2/20 | King Fahad Medical City | Alosaimi et al |
| B.1.458 | EPI_ISL_471590 | India/TG-CCMB-L1451/2020 | 2020/5/25 | CSIR-Centre for Cellular and Molecular Biology | Lamuk Zaveri et al |
| B.1.459 | EPI_ISL_428715 | Turkey/HSGM-5624/2020 | 2020/3/18 | Ministry of Health Turkey | Fatma Bayrakdar et al |
| B.1.460 | EPI_ISL_567528 | England/QEUH-9C9F1C/2020 | 2020/9/10 | Wellcome Sanger Institute for the COVID-19 Genomics UK (COG-UK) consortium | Harper VanSteenhouse et al |

| | | | | | |
|---|---|---|---|---|---|
| B.1.462 | EPI_ISL_729920 | Nigeria/KD-CV233/2020 | 2020/4/20 | African Centre of Excellence for Genomics of Infectious Diseases (ACEGID), Redeemer's University, Ede, Osun State, Nigeria | Oluniyi P.E. et al |
| B.1.463 | EPI_ISL_737061 | Finland/S12A1/2020 | 2020/9/10 | Department of Virology, Faculty of Medicine, University of Helsinki, Helsinki, Finland | Teemu Smura et al |
| B.1.465 | EPI_ISL_1184553 | Finland/10MS2E4/2020 | 2020/3/10 | Department of Virology, Faculty of Medicine, University of Helsinki, Helsinki, Finland | Teemu Smura et al |
| B.1.466 | EPI_ISL_482765 | Egypt/MASRI-014/2020 | 2020/4/30 | Medical Ain Shams Research Institute (MASRI), Ain Shams University | Hesham Elghazaly et al |
| B.1.466.1 | EPI_ISL_1015121 | France/IDF_PSL_384/2020 | 2020/6/9 | Department of Virology, Pitié-Salpêtrière hospital | Valentin Leducq et al |
| B.1.466.2 | EPI_ISL_2868921 | Indonesia/BT-NIHRD-WGS03289/2020 | 2020/8/6 | National Institute of Health Research and Development | Vivi Setiawaty et al |
| B.1.467 | EPI_ISL_730012 | Nigeria/BO-CV347/2020 | 2020/5/17 | African Centre of Excellence for Genomics of Infectious Diseases (ACEGID), Redeemer's University, Ede, Osun State, Nigeria | Oluniyi P.E. et al |
| B.1.468 | EPI_ISL_490001 | SaudiArabia/545/2020 | 2020/2/13 | King Fahad Medical City | Alosaimi et al |
| B.1.469 | EPI_ISL_552561 | England/MILK-97551D/2020 | 2020/8/21 | Wellcome Sanger Institute for the COVID-19 Genomics UK (COG-UK) consortium | The Lighthouse Lab in Milton Keynes et al |
| B.1.470 | EPI_ISL_759965 | Indonesia/JI-ITDua-2858NTv/2020 | 2020/4/9 | Institute of Tropical Disease, Universitas Airlangga | Rima R Prasetya et al |
| B.1.471 | EPI_ISL_437496 | SaudiArabia/KAUST-Jeddah59/2020 | 2020/3/25 | Pathogen Genomics Lab King Abdullah University of Science and Technology(KAUST) | Sara Mfarrej et al |
| B.1.473 | EPI_ISL_426428 | USA/NC-CDC-03068803-001/2020 | 2020/3/8 | Pathogen Discovery, Respiratory Viruses Branch, Division of Viral Diseases, Centers for Disease Control and Prevention | Krista Queen et al |
| B.1.474 | EPI_ISL_1015160 | France/IDF_PSL_415/2020 | 2020/9/21 | Department of Virology, Pitié-Salpêtrière hospital | Valentin Leducq et al |
| B.1.475 | EPI_ISL_753805 | Germany/BE-ChVir-W1302-8173/2020 | 2020/4/29 | Charité Universitätsmedizin Berlin, Institut für Virologie | Victor M Corman et al |

| | | | | | |
|---|---|---|---|---|---|
| B.1.476 | EPI_ISL_2596774 | France/PAC-IHU-10734_Illu1/2020 | 2020/4/2 | MEPHI, Aix Marseille University | Anthony LEVASSEUR et al |
| B.1.478 | EPI_ISL_2157568 | Haiti/34003/2020 | 2020/5/15 | Laboratory of Respiratory Viruses and Measles, Oswaldo Cruz Institute, FIOCRUZ | Paola Resende et al |
| B.1.479 | EPI_ISL_460364 | USA/MA-MGH-00136/2020 | 2020/3/25 | Infectious Disease Program, Broad Institute of Harvard and MIT | Lemieux et al |
| B.1.480 | EPI_ISL_512722 | Australia/WA69/2020 | 2020/7/3 | PathWest Laboratory Medicine WA Microbial Surveillance Unit | PathWest Laboratory Medicine WA Microbial Surveillance Unit et al |
| B.1.482 | EPI_ISL_420045 | France/GES-2843/2020 | 2020/3/17 | National Reference Center for Viruses of Respiratory Infections, Institut Pasteur, Paris | Mélanie Albert et al |
| B.1.483 | EPI_ISL_428254 | USA/WI-UW-158/2020 | 2020/3/15 | University of Wisconsin-Madison AIDS Vaccine Research Laboratories | Gage Moreno et al |
| B.1.485 | EPI_ISL_2246395 | USA/MI-MDHHS-SC26958/2020 | 2020/4/27 | Michigan Department of Health and Human Services, Bureau of Laboratories | Blankenship HM et al |
| B.1.486 | EPI_ISL_593478 | USA/MA-JLL-D18/2020 | 2020/4/28 | Jonathan Li Laboratory | Manish C. Choudhary et al |
| B.1.487 | EPI_ISL_545210 | USA/TX-HMH-2260/2020 | 2020/5/24 | Houston Methodist Hospital | S. Wesley Long et al |
| B.1.488 | EPI_ISL_534228 | Sweden/20-08938/2020 | 2020/7/13 | The Public Health Agency of Sweden | Anna-Malin Linde et al |
| B.1.489 | EPI_ISL_422164 | Wales/PHWC-26620/2020 | 2020/3/26 | Public Health Wales Microbiology Cardiff | Catherine Moore et al |
| B.1.490 | EPI_ISL_481595 | Finland/20A16S7/2020 | 2020/4/20 | Department of Virology, Faculty of Medicine, University of Helsinki, Helsinki, Finland | Teemu Smura et al |
| B.1.491 | EPI_ISL_438718 | England/CAMB-83FBD/2020 | 2020/4/22 | COVID-19 Genomics UK (COG-UK) Consortium | Luke W Meredith et al |
| B.1.492 | EPI_ISL_677289 | USA/CO-CDPHE-2004230036/2020 | 2020/4/21 | Colorado Department of Puplic Health and Environment | Laura Bankers et al |
| B.1.493 | EPI_ISL_508860 | USA/FL-BPHL-0656/2020 | 2020/4/7 | Florida Bureau of Public Health Laboratories | Sarah Schmedes et al |
| B.1.494 | EPI_ISL_2691720 | USA/MD-HP13053-PIDCLFMKUI/2020 | 2020/3/31 | Johns Hopkins Hospital Department of Pathology | C. Paul Morris et al |
| B.1.495 | EPI_ISL_482357 | USA/OR-PROV-086/2020 | 2020/4/19 | Providence St. Joseph Health Molecular Genomics Laboratory | Alexa K Dowdell et al |
| B.1.496 | EPI_ISL_444550 | USA/IL-NM054/2020 | 2020/3/15 | Ozer Lab | Ramon Lorenzo-Redondo et al |
| B.1.498 | EPI_ISL_494523 | USA/NC-QDX-171/2020 | 2020/4/27 | Quest Diagnostics | Anderson et al |

| | | | | | |
|---|---|---|---|---|---|
| B.1.499 | EPI_ISL_4405694 | Argentina/PAIS-A1026/2020 | 2020/1/1 | Área de Secuenciación del Laboratorio de Virología del Hospital de Niños Dr. Ricardo Gutierrez on behalf of 'Proyecto Argentino Interinstitucional de genomica de SARS-CoV-2' (PAIS Consortium) | Nabaes Jodar et al |
| B.1.499 | EPI_ISL_1396437 | Argentina/PAIS-H0010/2020 | 2020/8/25 | Laboratorio Mixto de Biotecnología Acuática (LMBA) on behalf of 'Proyecto Argentino Interinstitucional de genomica de SARS-CoV-2' (PAIS Consortium) | Joaquín Ezpeleta et al |
| B.1.501 | EPI_ISL_494526 | USA/NC-QDX-174/2020 | 2020/4/27 | Quest Diagnostics | Anderson et al |
| B.1.502 | EPI_ISL_544032 | USA/TX-HMH-3156/2020 | 2020/6/11 | Houston Methodist Hospital | S. Wesley Long et al |
| B.1.503 | EPI_ISL_526828 | USA/VA-DCLS-0612/2020 | 2020/4/1 | Virginia DCLS | Virginia DCLS et al |
| B.1.504 | EPI_ISL_474971 | Israel/CVL-n-9560/2020 | 2020/4/19 | Israel Central Virology laboratory | Neta Zuckerman et al |
| B.1.505 | EPI_ISL_447381 | Israel/2086004/2020 | 2020/4/1 | Stern Lab | Stern Lab et al |
| B.1.506 | EPI_ISL_525686 | USA/NY-Wadsworth-57118-01/2020 | 2020/4/20 | Wadsworth Center, New York State Department of Health | Kirsten St. George et al |
| B.1.507 | EPI_ISL_450050 | USA/NY-PV09362/2020 | 2020/3/15 | MSHS Pathogen Surveillance Program | Ana S. Gonzalez-Reiche et al |
| B.1.508 | EPI_ISL_754930 | USA/CA-CDPH041/2020 | 2020/4/28 | California Department of Public Health | CDPH IDLB COVIDNet et al |
| B.1.509 | EPI_ISL_653139 | USA/FL-BPHL-1986/2020 | 2020/4/18 | Florida Bureau of Public Health Laboratories | Sarah Schmedes et al |
| B.1.510 | EPI_ISL_2426012 | Norway/1766-2/2020 | 2020/3/6 | Norwegian Institute of Public Health, Department of Virology | Kathrine Stene-Johansen et al |
| B.1.511 | EPI_ISL_542678 | USA/TX-HMH-1371/2020 | 2020/5/2 | Houston Methodist Hospital | S. Wesley Long et al |
| B.1.513 | EPI_ISL_451999 | Denmark/ALAB-HH-115/2020 | 2020/3/9 | Albertsen lab, Department of Chemistry and Bioscience, Aalborg University, Denmark | Rasmus Kirkegaard et al |
| B.1.515 | EPI_ISL_462706 | USA/MI-MDHHS-SC20624/2020 | 2020/4/17 | Michigan Department of Health and Human Services, Bureau of Laboratories | Blankenship HM et al |
| B.1.516 | EPI_ISL_631862 | USA/NY-NYCPHL-000356/2020 | 2020/3/11 | New York City Public Health Laboratory | Jade Wang et al |
| B.1.517 | EPI_ISL_2976222 | USA/ID-IBL-614702/2020 | 2020/4/6 | Idaho Bureau of Laboratories | "R. Beukelman et al |
| B.1.517.1 | EPI_ISL_710125 | Australia/NSW1227/2020 | 2020/12/2 | CIDM-PH et al. | CIDM-PH et al. |
| B.1.518 | EPI_ISL_426054 | USA/CT-UW-4347/2020 | 2020/3/30 | UW Virology Lab | Pavitra Roychoudhury et al |
| B.1.520 | EPI_ISL_582289 | Canada/MB-NML-617/2020 | 2020/3/25 | National Microbiology Laboratory (NML) | Anna Majer et al |
| B.1.521 | EPI_ISL_508793 | USA/FL-BPHL-0589/2020 | 2020/3/21 | Florida Bureau of Public Health Laboratories | Sarah Schmedes et al |

| | | | | | |
|---|---|---|---|---|---|
| B.1.523 | EPI_ISL_437733 | SaudiArabia/KAUST-Madinah238/2020 | 2020/3/29 | Pathogen Genomics Lab King Abdullah University of Science and Technology(KAUST) | Sharif Hala et al |
| B.1.524 | EPI_ISL_644671 | Malaysia/IMR_CV140342/2020 | 2020/9/1 | Institute for Medical Research, Infectious Disease Research Centre, National Institutes of Health, Ministry of Health Malaysia | Suppiah J et al |
| B.1.525 | EPI_ISL_4739567 | Sudan/N6667/2020 | 2020/10/4 | National Institute for Communicable Diseases of the National Health Laboratory Service | Elham Rizgalla et al |
| B.1.526 | EPI_ISL_3364539 | USA/NY-GEO-0231/2020 | 2020/1/28 | Biotia | Dorottya Nagy-Szakal et al |
| B.1.527 | EPI_ISL_636467 | Italy/VEN-IZSVe-14-21/2020 | 2020/3/24 | Istituto Zooprofilattico Sperimentale delle Venezie | Adelaide Milani et al |
| B.1.528 | EPI_ISL_451400 | Morocco/OUA677-19/2020 | 2020/4/23 | Laboratoire de Recherche et d'Analyse Médicale de la Gendarmerie Royale | Sanaâ LEMRISS et al |
| B.1.529 | EPI_ISL_645201 | France/OCC-SC545/2020 | 2020/4/2 | CNR Virus des Infections Respiratoires - France SUD | Antonin Bal et al |
| B.1.529 | EPI_ISL_1133148 | Italy/LOM-Pavia-48855/2020 | 2020/4/28 | Molecular Virology Unit, Microbiology and Virology Depertment, Fondazione IRCCS Policlinico San Matteo, Pavia | Monica Tallarita et al |
| B.1.530 | EPI_ISL_3049453 | Kenya/C57901/2020 | 2020/9/15 | KEMRI-Wellcome Trust Research Programme,Kilifi | Githinji G. et al |
| B.1.531 | EPI_ISL_783074 | England/MILK-BAB136/2020 | 2020/11/18 | Wellcome Sanger Institute for the COVID-19 Genomics UK (COG-UK) Consortium | The Lighthouse Lab in Milton Keynes et al |
| B.1.532 | EPI_ISL_556109 | England/MILK-6422BB/2020 | 2020/7/4 | Wellcome Sanger Institute for the COVID-19 Genomics UK (COG-UK) consortium | The Lighthouse Lab in Milton Keynes et al |
| B.1.533 | EPI_ISL_512937 | SaudiArabia/KAUST-JEDDAH494/2020 | 2020/4/9 | Pathogen Genomics Lab King Abdullah University of Science and Technology(KAUST) | Fadwa Alofi et al |
| B.1.534 | EPI_ISL_1876157 | Denmark/DCGC-67586/2020 | 2020/11/2 | Aalborg University | Danish Covid-19 Genome Consortium et al |
| B.1.535 | EPI_ISL_776198 | Germany/HH-hpi-p1847/2020 | 2020/3/22 | Heinrich Pette Institute, Leibniz Institute for Experimental Virology | Alexis Robitaille et al |
| B.1.536 | EPI_ISL_616955 | Denmark/DCGC-8136/2020 | 2020/10/26 | Albertsen lab, Department of Chemistry and Bioscience, Aalborg University, Denmark | Danish Covid-19 Genome Consortia et al |

| | | | | | |
|---|---|---|---|---|---|
| B.1.537 | EPI_ISL_528810 | India/TG-CCMB-GC1/2020 | 2020/5/24 | CSIR-Centre for Cellular and Molecular Biology | Thrilok Chander Bingi et al |
| B.1.538 | EPI_ISL_476890 | India/MP-DRDE-4022/2020 | 2020/5/16 | Defence Research & Development Establishment (DRDE) | Shashi Sharma et al |
| B.1.539 | EPI_ISL_1624329 | USA/VA_NIDDL_27852/2020 | 2020/10/25 | Naval Medical Research Center Biological Defense Research Directorate | Logan Voegtly et al |
| B.1.540 | EPI_ISL_455640 | India/WB-S4/2020 | 2020/3/27 | National Institute of Biomedical Genomics | Arindam Maitra et al |
| B.1.541 | EPI_ISL_579016 | USA/LA-EVTL612/2020 | 2020/7/1 | Microbial Genome Sequencing Center | Jeremy P. Kamil et al |
| B.1.543 | EPI_ISL_801878 | USA/NY-MSHSPSP-PV17906/2020 | 2020/8/30 | MSHS Pathogen Surveillance Program | Ana S. Gonzalez-Reiche et al |
| B.1.544 | EPI_ISL_581493 | Congo/UKT-015/2020 | 2020/6/25 | NGS Competence Center Tübingen, Institut für Medizinische Mikrobiologie und Hygiene, Universitätsklinikum Tübingen | Angel Angelov et al |
| B.1.545 | EPI_ISL_579469 | NewZealand/20VR3181/2020 | 2020/3/25 | Institute of Environmental Science and Research (ESR) | Xiaoyun Ren et al |
| B.1.546 | EPI_ISL_545546 | USA/TX-HMH-2776/2020 | 2020/6/3 | Houston Methodist Hospital | S. Wesley Long et al |
| B.1.547 | EPI_ISL_535765 | Canada/QC-LSPQ-L00217574/2020 | 2020/3/13 | Laboratoire de santé publique du Québec | Sandrine Moreira et al |
| B.1.548 | EPI_ISL_471623 | India/TG-CCMB-L1693/2020 | 2020/5/29 | CSIR-Centre for Cellular and Molecular Biology | Payel Mukherjee et al |
| B.1.549 | EPI_ISL_2779501 | Kenya/MAL-5042/2020 | 2020/6/9 | USAMRD-A, Basic Science Laboratory | Gathii Kimita et al |
| B.1.550 | EPI_ISL_872402 | USA/AL-Morgan-JDTB/2020 | 2020/10/3 | Hudsonalpha Genome Sequencing Center | Jane Grimwood et al |
| B.1.551 | EPI_ISL_837802 | Mexico/CMX-INER-0208/2020 | 2020/8/20 | Instituto Nacional de Enfermedades Respiratorias (INER) | Celia Boukadida et al |
| B.1.552 | EPI_ISL_467971 | USA/CA-SR0447/2020 | 2020/3/30 | Andersen lab at Scripps Research | SEARCH Alliance San Diego with Tracy Basler et al |
| B.1.554 | EPI_ISL_2678230 | USA/UT-UPHL-2105275840/2020 | 2020/4/21 | Utah Public Health Laboratory | Erin L. Young et al |
| B.1.555 | EPI_ISL_445221 | Sweden/20-05444/2020 | 2020/3/11 | The Public Health Agency of Sweden | Frida Ahlfors et al |
| B.1.556 | EPI_ISL_648834 | USA/CA-ALSR-4125/2020 | 2020/6/10 | Andersen lab at Scripps Research | SEARCH Alliance San Diego with Tracy Basler et al |
| B.1.557 | EPI_ISL_459336 | NorthernIreland/NIRE-FA7AA/2020 | 2020/4/6 | Wellcome Sanger Institute for the COVID-19 Genomics UK (COG-UK) consortium | Conall McCaughey et al |

| | | | | | |
|---|---|---|---|---|---|
| B.1.558 | EPI_ISL_437575 | USA/CA-SR0225/2020 | 2020/4/6 | Andersen lab at Scripps Research | SEARCH Alliance San Diego with Michael Quigley et al |
| B.1.559 | EPI_ISL_439142 | Scotland/CVR989/2020 | 2020/3/29 | COVID-19 Genomics UK (COG-UK) Consortium | Ana da Silva Filipe et al |
| B.1.560 | EPI_ISL_1383217 | Spain/NC-CHN-01000049/2020 | 2020/3/20 | Centro de Secuenciación NASERTIC | Carmen Ezpeleta Baquedano et al |
| B.1.561 | EPI_ISL_1531819 | Mexico/BCN-ALSR-8354/2020 | 2020/6/18 | Andersen lab at Scripps Research | SEARCH Alliance San Diego with Idanya Rubi Serafin Higuera et al |
| B.1.562 | EPI_ISL_761042 | England/ALDP-CB2E16/2020 | 2020/12/1 | Wellcome Sanger Institute for the COVID-19 Genomics UK (COG-UK) Consortium | Jacquelyn Wynn et al |
| B.1.563 | EPI_ISL_709546 | England/ALDP-C06DCB/2020 | 2020/12/6 | Wellcome Sanger Institute for the COVID-19 Genomics UK (COG-UK) Consortium | Jacquelyn Wynn et al |
| B.1.564 | EPI_ISL_1165135 | USA/OH-ODH-SC149737/2020 | 2020/3/11 | Ohio Department of Health Laboratory | Holmes et al |
| B.1.564.1 | EPI_ISL_591187 | Canada/ON-S1416/2020 | 2020/8/23 | McMaster University | Allison McGeer et al |
| B.1.565 | EPI_ISL_746393 | USA/UT-UPHL-2012514374/2020 | 2020/6/18 | Utah Public Health Laboratory | Erin Young et al |
| B.1.566 | EPI_ISL_571800 | USA/TX-QDX-984/2020 | 2020/3/16 | Quest Diagnostics | Rosenthal et al |
| B.1.567 | EPI_ISL_460055 | USA/MN-MDH-152/2020 | 2020/3/23 | Minnesota Department of Health, Public Health Laboratory | Matt Plumb et al |
| B.1.568 | EPI_ISL_872412 | USA/AL-Shelby-JDTM/2020 | 2020/10/5 | Hudsonalpha Genome Sequencing Center | Jane Grimwood et al |
| B.1.570 | EPI_ISL_5253966 | USA/CO-CDPHE-2007100817/2020 | 2020/7/6 | Colorado Department of Public Health and Environment | Laura Bankers et al |
| B.1.571 | EPI_ISL_543079 | USA/TX-HMH-3627/2020 | 2020/6/15 | Houston Methodist Hospital | S. Wesley Long et al |
| B.1.572 | EPI_ISL_434942 | USA/TX-HMH0348/2020 | 2020/4/2 | Houston Methodist Hospital | S. Wesley Long et al |
| B.1.573 | EPI_ISL_968263 | Canada/BC-BCCDC-3565/2020 | 2020/6/24 | BCCDC Public Health Laboratory | Prystajecky Natalie et al |
| B.1.574 | EPI_ISL_1138489 | USA/IL-IDPH-COO-C-000663/2020 | 2020/5/1 | Gagnon Lab, Southern Illinois University | Keith Gagnon et al |
| B.1.575 | EPI_ISL_786939 | USA/TX-HMH-MCoV-15455/2020 | 2020/10/23 | Houston Methodist Hospital | S. Wesley Long et al |
| B.1.575.1 | EPI_ISL_4425102 | Spain/MD-ISCIII-215564/2020 | 2020/11/30 | Instituto de Salud Carlos III | Vázquez-Morón et al |
| B.1.575.2 | EPI_ISL_2776189 | DominicanRepublic/Yale-5507/2021 | 2021/1/14 | Grubaugh Lab - Yale School of Public Health | Joseph Fauver et al |

| | | | | | |
|---|---|---|---|---|---|
| B.1.576 | EPI_ISL_2877403 | France/HDF-IPP3652/2020 | 2020/3/25 | National Reference Center for Viruses of Respiratory Infections, Institut Pasteur, Paris | Marion Barbet et al |
| B.1.577 | EPI_ISL_542702 | USA/TX-HMH-1404/2020 | 2020/4/17 | Houston Methodist Hospital | S. Wesley Long et al |
| B.1.578 | EPI_ISL_604296 | USA/CA-QDX-1984/2020 | 2020/3/17 | Quest Diagnostics | Rosenthal et al |
| B.1.579 | EPI_ISL_1113373 | USA/AZ-TG724120/2020 | 2020/4/24 | TGen North | "Jolene Bowers et al |
| B.1.581 | EPI_ISL_543334 | USA/TX-HMH-4060/2020 | 2020/6/18 | Houston Methodist Hospital | S. Wesley Long et al |
| B.1.582 | EPI_ISL_3275380 | Maldives/MAV00499/2020 | 2020/4/11 | Indira Gandhi Memorial Hospital | Dr. Milza Abdul Muhsin et al |
| B.1.585 | EPI_ISL_786330 | USA/TX-HMH-MCoV-14846/2020 | 2020/10/13 | Houston Methodist Hospital | S. Wesley Long et al |
| B.1.586 | EPI_ISL_468582 | USA/NY-QDX-87/2020 | 2020/3/12 | Quest Diagnostics | Anderson et al |
| B.1.587 | EPI_ISL_667079 | USA/OR-OHSU-0070/2020 | 2020/4/23 | Oregon SARS-CoV-2 Genome Sequencing Center | Brendan L. O'Connell et al |
| B.1.588 | EPI_ISL_1168693 | PuertoRico/PR-CDC-S172/2020 | 2020/8/2 | Centers for Disease Control and Prevention, Dengue Branch | Gilberto A. Santiago et al |
| B.1.588.1 | EPI_ISL_738317 | Romania/USAFSAM-S556/2020 | 2020/11/28 | United States Air Force School of Aerospace Medicine | Anthony Fries et al |
| B.1.589 | EPI_ISL_572109 | USA/MN-QDX-1320/2020 | 2020/3/15 | Quest Diagnostics | Rosenthal et al |
| B.1.590 | EPI_ISL_572166 | USA/MN-QDX-1377/2020 | 2020/3/15 | Quest Diagnostics | Rosenthal et al |
| B.1.591 | EPI_ISL_1494524 | USA/IL-IDPH-WIN-S-0002001/2020 | 2020/8/27 | Gagnon Lab, Southern Illinois University | Keith Gagnon et al |
| B.1.592 | EPI_ISL_889857 | Canada/QC-JUS-V6011098/2020 | 2020/3/30 | Laboratoire de santé publique du Québec | Sandrine Moreira et al |
| B.1.593 | EPI_ISL_5440637 | USA/CO-CDPHE-2007070535/2020 | 2020/7/3 | Colorado Department of Public Health and Environment | Laura Bankers et al |
| B.1.595 | EPI_ISL_424868 | USA/LA-CDC-03068669-001/2020 | 2020/3/9 | Pathogen Discovery, Respiratory Viruses Branch, Division of Viral Diseases, Centers for Disease Control and Prevention | Yan Li et al |
| B.1.595.1 | EPI_ISL_804868 | USA/DC-DFS-PHL-0076/2020 | 2020/5/16 | DC Public Health Lab/ Dept. of Forensic Sciences | Scott Nguyen et al |
| B.1.595.2 | EPI_ISL_914336 | USA/AZ-TG375489/2020 | 2020/6/14 | TGen North | "Jolene Bowers et al |
| B.1.595.3 | EPI_ISL_1482724 | USA/SC-MUSC-0476/2020 | 2020/6/27 | MUSC Molecular Pathology Laboratory | Julie W. Hirschhorn et al |
| B.1.595.4 | EPI_ISL_477276 | USA/MN-MDH-1186/2020 | 2020/4/20 | Minnesota Department of Health, Public Health Laboratory | Matt Plumb et al |
| B.1.596 | EPI_ISL_590772 | USA/MN-MDH-1768/2020 | 2020/4/11 | Minnesota Department of Health, Public Health Laboratory | Matt Plumb et al |

| | | | | | |
|---|---|---|---|---|---|
| B.1.596.1 | EPI_ISL_3049403 | Kenya/C54633/2020 | 2020/9/7 | KEMRI-Wellcome Trust Research Programme,Kilifi | Githinji G. et al |
| B.1.597 | EPI_ISL_699656 | Tunisia/5008/2020 | 2020/3/24 | 1-Clinical and Experimental Pharmacology Lab, LR16SP02, National Center of Pharmacovigilance, University of Tunis El Manar, Tunis, Tunisia. 2-Neurodegenerative diseases and psychiatric troubles, LR18SP03, Razi Hospital, University of Tunis El Manar, Tunis, Tunisia. 3- Ministry of Health, National Observatory of New and Emerging Diseases, 1006, Tunis, Tunisia | Ilhem Boutiba-Ben Boubaker et al |
| B.1.598 | EPI_ISL_444679 | USA/NY-NYUMC562/2020 | 2020/3/30 | Departments of Pathology and Medicine, New York University School of Medicine | Maria Aguero-Rosenfeld et al |
| B.1.599 | EPI_ISL_582465 | Canada/MB-NML-1134/2020 | 2020/7/20 | National Microbiology Laboratory (NML) | Anna Majer et al |
| B.1.6 | EPI_ISL_475866 | Austria/CeMM0449/2020 | 2020/3/5 | Bergthaler laboratory, CeMM Research Center for Molecular Medicine of the Austrian Academy of Sciences | Alexandra Popa et al |
| B.1.600 | EPI_ISL_1167817 | Chile/VS-39494/2020 | 2020/4/7 | Instituto de Salud Publica de Chile | Javier Tognarelli et al |
| B.1.601 | EPI_ISL_5736723 | Italy/EMR-UA-02_00003/2020 | 2020/3/4 | Laboratory of Medical Microbiology, University of Antwerp | Mathias Smet et al |
| B.1.601 | EPI_ISL_2923967 | USA/NY-PRL-2021_0706_51F19/2020 | 2020/10/28 | Pandemic Response Lab, R&D | Henry Lee et al |
| B.1.602 | EPI_ISL_525951 | USA/OR-OHSU-1023/2020 | 2020/7/7 | Oregon SARS-CoV-2 Genome Sequencing Center | Brendan L. O'Connell et al |
| B.1.603 | EPI_ISL_683880 | USA/NY-NYCPHL-001318/2020 | 2020/11/10 | New York City Public Health Laboratory | Jade Wang et al |
| B.1.604 | EPI_ISL_876038 | USA/NY-AECOM_109/2020 | 2020/7/28 | Albert Einstein College of Medicine, Dept. of Microbiology & Immunology, Chandran lab | J. Maximilian Fels et al |
| B.1.605 | EPI_ISL_522781 | USA/VA-DCLS-0908/2020 | 2020/5/12 | Virginia DCLS | Virginia DCLS et al |
| B.1.606 | EPI_ISL_545165 | USA/TX-HMH-2178/2020 | 2020/5/21 | Houston Methodist Hospital | S. Wesley Long et al |
| B.1.607 | EPI_ISL_420221 | England/SHEF-C03E5/2020 | 2020/3/29 | Department of Infection, Immunity and Cardiovascular Disease, The Florey Institute, The Medical School, University of Sheffield | Thushan de Silva et al |

| | | | | | |
|---|---|---|---|---|---|
| B.1.609 | EPI_ISL_419680 | Spain/VC-FISABIO-16/2020 | 2020/3/10 | Sequencing and Bioinformatics Service and Molecular Epidemiology Research Group. FISABIO-Public Health | Maria Alma Bracho et al |
| B.1.609 | EPI_ISL_2835331 | Mexico/BCN-SEARCH-102851/2020 | 2020/4/28 | Andersen lab at Scripps Research | SEARCH Alliance with Gisela Barrera Badillo et al |
| B.1.610 | EPI_ISL_529513 | England/ALDP-98EAC7/2020 | 2020/7/11 | COVID-19 Genomics UK (COG-UK) Consortium | Sam Haldenby et al |
| B.1.611 | EPI_ISL_825829 | Canada/Qc-CHUM-2016103625A/2020 | 2020/6/9 | Laboratoire de santé publique du Québec | Sandrine Moreira et al |
| B.1.612 | EPI_ISL_590748 | USA/MI-UM-MH27455/2020 | 2020/3/11 | Lauring Lab, University of Michigan, Department of Microbiology and Immunology | Valesano et al |
| B.1.613 | EPI_ISL_618805 | Denmark/DCGC-1386/2020 | 2020/4/6 | Albertsen lab, Department of Chemistry and Bioscience, Aalborg University, Denmark | Danish Covid-19 Genome Consortia et al |
| B.1.614 | EPI_ISL_510871 | Sweden/20-52071/2020 | 2020/6/15 | The Public Health Agency of Sweden | Oskar Karlsson Lindsjo et al |
| B.1.615 | EPI_ISL_825696 | Canada/Qc-JUS-V927155399/2020 | 2020/7/26 | Laboratoire de santé publique du Québec | Sandrine Moreira et al |
| B.1.616 | EPI_ISL_1118892 | France/BRE-IPP03808/2021 | 2021/2/10 | National Reference Center for Viruses of Respiratory Infections, Institut Pasteur, Paris | Marion Barbet et al |
| B.1.617.1 | EPI_ISL_2758215 | India/un-IRSHA-CD210871/2020 | 2020/3/3 | Communicable Diseases, Interactive Research School for Health Affairs (IRSHA) | Shrivastava et al |
| B.1.617.2 | EPI_ISL_1914592 | India/CT-AIIMS-Raipur-L-1010/2021 | 2021/3/30 | State Virus Research and Diagnostic Laboratory (VRDL), AIIMS Raipur | Pushpendra Singh et al |
| B.1.617.2 | EPI_ISL_2879009 | India/MH-IGIB-GSEQ_1954/2021 | 2021/3/23 | INSACOG at CSIR Institute of Genomics and Integrative Biology | INSACOG et al |
| B.1.617.2 | EPI_ISL_5644723 | India/GJ-INSACOG-GBRC2563/2021 | 2021/1/10 | Gujarat Biotechnology Research Centre | Sonal Sharma et al |
| B.1.617.2 | EPI_ISL_4768101 | Rwanda/CV2197/2020 | 2020/9/10 | Africa Centre for Excellence for Genomics of Infectious Diseases (ACEGID), Redeemer's University | Enatha Mukantwari et al |
| B.1.617.2 | EPI_ISL_2341920 | India/CT-ILSGS01119/2021 | 2021/3/21 | Institute of Life Sciences - INSACOG | Sunil K. Raghav et al |
| B.1.617.2 | EPI_ISL_3614352 | India/MN-1425500076304/2021 | 2021/4/25 | National Institute of Biomedical Genomics – INSACOG | Arindam Maitra et al |
| B.1.617.2 | EPI_ISL_4415533 | India/MH-IISER-Pune-IP-00986/2021 | 2021/2/9 | IISER Pune-INSACOG | Krishanpal Karmodiya et al |

| | | | | | |
|---|---|---|---|---|---|
| B.1.617.2 | EPI_ISL_6025382 | India/DL-ILBS-B_79/2021 | 2021/1/28 | ILBS | Ekta Gupta et al |
| B.1.617.2 | EPI_ISL_4415621 | India/MH-IISER-Pune-IP-01021/2021 | 2021/2/9 | IISER Pune-INSACOG | Krishanpal Karmodiya et al |
| B.1.617.2 | EPI_ISL_3614353 | India/MN-1425500076321/2021 | 2021/4/25 | National Institute of Biomedical Genomics – INSACOG | Arindam Maitra et al |
| B.1.617.2 | EPI_ISL_3215467 | USA/CA-SEARCH-18281/2021 | 2021/4/11 | Andersen lab at Scripps Research | Chip Schooley et al |
| B.1.617.2 | EPI_ISL_4415418 | India/MH-IISER-Pune-IP-00995/2021 | 2021/2/9 | IISER Pune-INSACOG | Krishanpal Karmodiya et al |
| B.1.617.2 | EPI_ISL_2842285 | India/KA-NIMH-SEQ-1757/2021 | 2021/4/10 | INSACOG-KA, NIMHANS | Chitra Pattabiraman et al |
| B.1.617.3 | EPI_ISL_1415164 | India/MH-NCCS-86945/2021 | 2021/2/13 | National Centre For Cell Science – INSACOG | Dhiraj Paul et al |
| B.1.618 | EPI_ISL_1419395 | India/WB-1931300237615/2020 | 2020/10/27 | National Institute of Biomedical Genomics – INSACOG | Arindam Maitra et al |
| B.1.619 | EPI_ISL_3458197 | Canada/QC-L00333705001A/2020 | 2020/3/10 | Laboratoire de santé publique du Québec | Sandrine Moreira et al |
| B.1.619.1 | EPI_ISL_2361093 | SouthKorea/KDCA4323/2021 | 2021/2/22 | Division of Emerging Infectious Diseases, Bureau of Infectious Diseases Diagnosis Control, Korea Disease Control and Prevention Agency | Ae Kyung Park et al |
| B.1.621 | EPI_ISL_6512535 | Colombia/ANT-LDSP461/2020 | 2020/10/14 | Laboratorio Departamental de Salud Publica de Antioquia | Idabely Betancur Ortiz et al |
| B.1.621.1 | EPI_ISL_4361670 | USA/OH-CDC-ASC210055228/2021 | 2021/4/9 | Centers for Disease Control and Prevention Division of Viral Diseases, Pathogen Discovery | Dakota Howard et al |
| B.1.622 | EPI_ISL_2245960 | Guinea/Conakry_IPG11718/2020 | 2020/9/16 | Institut Pateur de Dakar | Grayo Solene et al |
| B.1.625 | EPI_ISL_2339873 | Colombia/COR-U27/2020 | 2020/11/16 | Centro de Investigaciones en Microbiología y Biotecnología-UR (CIMBIUR), Facultad de Ciencias Naturales, Universidad del Rosario, Bogotá, Colombia | Salim Mattar et al |
| B.1.626 | EPI_ISL_2125133 | Germany/BY-RKI-I-141456/2021 | 2021/4/16 | Robert Koch Institute | ? |
| B.1.627 | EPI_ISL_3568367 | Australia/NSW-SAVID-3904/2021 | 2021/1/9 | Virology Research Laboratory; Area of Virology, Serology and Virology Division (SAViD), New South Wales Health Pathology Randwick | Foster et al |
| B.1.630 | EPI_ISL_2274018 | DominicanRepublic/21915/2021 | 2021/3/17 | Laboratory of Respiratory Viruses and Measles, Oswaldo Cruz Institute, FIOCRUZ | Paola Resende et al |

| | | | | | |
|---|---|---|---|---|---|
| B.1.631 | EPI_ISL_2628735 | USA/SEARCH-102258/2020 | 2020/12/16 | Andersen lab at Scripps Research | SEARCH Alliance San Diego with Ashleigh Murphy et al |
| B.1.632 | EPI_ISL_2343008 | Mexico/TAM-InDRE_FB14701_S2256/2021 | 2021/4/28 | Instituto de Diagnostico y Referencia Epidemiologicos (INDRE) | Claudia Wong-Arambula et al |
| B.1.633 | EPI_ISL_2461371 | India/DL-NCDC-2509105/2021 | 2021/4/14 | NCDC Delhi, Biotechnology Division INSACOG | Manoj K Singh et al |
| B.1.634 | EPI_ISL_1184562 | USA/AZ-ASPHL-3709/2021 | 2021/3/2 | Arizona State Public Health Laboratory | Trung Huynh et al |
| B.1.635 | EPI_ISL_1168492 | Mexico/VER-InDRE_414/2021 | 2021/1/16 | Instituto de Diagnostico y Referencia Epidemiologicos (INDRE) | Claudia Wong-Arambula et al |
| B.1.637 | EPI_ISL_801973 | USA/NY-MSHSPSP-PV21166/2020 | 2020/10/9 | MSHS Pathogen Surveillance Program | Ana S. Gonzalez-Reiche et al |
| B.1.639 | EPI_ISL_2443791 | USA/NY-NYGC-787-VTM1-OUGT620/2021 | 2021/1/19 | New York Genome Center | Michael Zody et al |
| B.1.640 | EPI_ISL_5592661 | Congo/FCRM-100-A32.28.09.21/2021 | 2021/9/28 | Fondation Congolaise pour la Recherche Médicale | Mfoutou Mapanguy Claujens Chastel et al |
| B.1.67 | EPI_ISL_420985 | Wales/PHWC-252B4/2020 | 2020/3/23 | Public Health Wales Microbiology Cardiff | Catherine Moore et al |
| B.1.69 | EPI_ISL_425808 | Scotland/CVR78/2020 | 2020/3/6 | COVID-19 Genomics UK (COG-UK) Consortium | Ana da Silva Filipe et al |
| B.1.70 | EPI_ISL_425759 | Scotland/CVR30/2020 | 2020/3/13 | COVID-19 Genomics UK (COG-UK) Consortium | Ana da Silva Filipe et al |
| B.1.76 | EPI_ISL_464467 | England/201080091/2020 | 2020/3/5 | Respiratory Virus Unit, Microbiology Services Colindale, Public Health England | PHE Covid Sequencing Team et al |
| B.1.77 | EPI_ISL_532277 | England/QEUH-52DB8F/2020 | 2020/5/18 | Wellcome Sanger Institute for the COVID-19 Genomics UK (COG-UK) consortium | Harper VanSteenhouse et al |
| B.1.78 | EPI_ISL_523523 | Netherlands/un-EMC-767/2020 | 2020/3/12 | Erasmus Medical Center | Bas Oude Munnink et al |
| B.1.8 | EPI_ISL_414530 | Netherlands/NoordBrabant_21/2020 | 2020/3/4 | Erasmus Medical Center | David Nieuwenhuijse et al |
| B.1.81 | EPI_ISL_440218 | England/BRIS-121C9B/2020 | 2020/3/18 | Wellcome Sanger Institute for the COVID-19 Genomics UK (COG-UK) consortium | Stephanie Hutchings et al |
| B.1.83 | EPI_ISL_1029971 | Belgium/MBLG51240/2020 | 2020/3/20 | UCLouvain/IREC/MBLG | Jean Ruelle et al |
| B.1.84 | EPI_ISL_940256 | France/IDF_HB_112003109999/2020 | 2020/3/31 | IAME UMR1137 Inserm, Université de Paris, Hôpital Bichat | Antoine Bridier-Nahmias et al |
| B.1.88.1 | EPI_ISL_1586845 | Iceland/1483/2020 | 2020/3/26 | deCODE genetics | Daniel F Gudbjartsson et al |

| | | | | | |
|---|---|---|---|---|---|
| B.1.9 | EPI_ISL_1499727 | Togo/BMC-4-20/2020 | 2020/3/4 | Unité Mixte Internationale TransVIHMI (UMI 233 IRD – U1175 INSERM - Université de Montpellier)IRD (Institut de recherche pour le développement) | Mounerou SALOU et al |
| B.1.9.1 | EPI_ISL_453188 | Scotland/EDB5108/2020 | 2020/3/31 | COVID-19 Genomics UK (COG-UK) Consortium | McHugh M et al |
| B.1.9.2 | EPI_ISL_467793 | USA/VA-DCLS-0398/2020 | 2020/4/20 | Virginia DCLS | Virginia DCLS et al |
| B.1.9.3 | EPI_ISL_462180 | Belgium/JBE-0317326/2020 | 2020/3/17 | KU Leuven, Rega Institute, Clinical and Epidemiological Virology | Tony Wawina-Bokalanga et al |
| B.1.9.4 | EPI_ISL_500920 | Switzerland/BL-ETHZ-190059/2020 | 2020/7/9 | Department of Biosystems Science and Engineering, ETH Zürich | Christian Beisel et al |
| B.1.9.5 | EPI_ISL_417944 | DRC/81/2020 | 2020/3/18 | Pathogen Sequencing Lab, National Institute for Biomedical Research (INRB) | Placide Mbala-Kingebeni et al |
| B.1.91 | EPI_ISL_451307 | Italy/LOM-INMI-5314-N/2020 | 2020/2/21 | Laboratory of Virology, INMI Lazzaro Spallanzani IRCCS | Fausto Baldanti et al |
| B.1.93 | EPI_ISL_644683 | France/OCC-SC467/2020 | 2020/3/6 | CNR Virus des Infections Respiratoires - France SUD | Antonin Bal et al |
| B.1.94 | EPI_ISL_419564 | Luxembourg/LNS0366116/2020 | 2020/3/15 | Laboratoire National de Santé, Microbiology, Epidemiology and Microbial Genomics | Anke Wienecke-Baldacchino et al |
| B.1.96 | EPI_ISL_422601 | Netherlands/NA_292/2020 | 2020/4/1 | Erasmus Medical Center | Bas Oude Munnink et al |
| B.1.97 | EPI_ISL_439884 | England/CAMB-726F8/2020 | 2020/3/24 | Wellcome Sanger Institute for the COVID-19 Genomics UK (COG-UK) consortium | Luke W Meredith et al |
| B.10 | EPI_ISL_492694 | England/BRIS-1309ED/2020 | 2020/3/12 | Wellcome Sanger Institute for the COVID-19 Genomics UK (COG-UK) consortium | Stephanie Hutchings et al |
| B.11 | EPI_ISL_413586 | Netherlands/Tilburg_1363354/2020 | 2020/2/27 | Erasmus Medical Center | David Nieuwenhuijse et al |
| B.12 | EPI_ISL_479799 | Japan/PG-0015/2020 | 2020/1/20 | Pathogen Genomics Center, National Institute of Infectious Diseases | Tsuyoshi Sekizuka et al |
| B.13 | EPI_ISL_436572 | USA/WI-UW-278/2020 | 2020/4/3 | University of Wisconsin-Madison AIDS Vaccine Research Laboratories | Gage Moreno et al |
| B.15 | EPI_ISL_425480 | England/NOTT-10DFD8/2020 | 2020/3/16 | COVID-19 Genomics UK (COG-UK) Consortium | Gemma Clark et al |
| B.18 | EPI_ISL_417553 | Iceland/220/2020 | 2020/3/16 | deCODE genetics | Daniel F Gudbjartsson et al |
| B.19 | EPI_ISL_436672 | USA/CA-CZB-1031/2020 | 2020/3/29 | Chan-Zuckerberg Biohub | CZB Cliahub Consortium et al |

| | | | | | |
|---|---|---|---|---|---|
| B.20 | EPI_ISL_437044 | USA/CA-CZB-1044/2020 | 2020/3/31 | Chan-Zuckerberg Biohub | CZB CliaHub Consortium et al |
| B.23 | EPI_ISL_464328 | England/20104013504/2020 | 2020/2/28 | Respiratory Virus Unit, Microbiology Services Colindale, Public Health England | PHE Covid Sequencing Team et al |
| B.26 | EPI_ISL_425701 | Scotland/CVR162/2020 | 2020/3/17 | COVID-19 Genomics UK (COG-UK) Consortium | Ana da Silva Filipe et al |
| B.28 | EPI_ISL_464563 | England/20109045404/2020 | 2020/3/6 | Respiratory Virus Unit, Microbiology Services Colindale, Public Health England | PHE Covid Sequencing Team et al |
| B.29 | EPI_ISL_415144 | England/20102000106/2020 | 2020/2/27 | Respiratory Virus Unit, Microbiology Services Colindale, Public Health England | Monica Galiano et al |
| B.3 | EPI_ISL_414497 | Germany/NW-HHU-02-1/2020 | 2020/2/25 | Center of Medical Microbiology, Virology, and Hospital Hygiene, University of Duesseldorf | Ortwin Adams et al |
| B.3.1 | EPI_ISL_416389 | Shanghai/SH0093/2020 | 2020/1/21 | National Research Center for Translational Medicine (Shanghai), Ruijin Hospital affiliated to Shanghai Jiao Tong University School of Medicine & Shanghai Public Health Clinical Center | Shengyue Wang et al |
| B.30 | EPI_ISL_482451 | USA/OR-PROV-216/2020 | 2020/3/9 | Providence St. Joseph Health Molecular Genomics Laboratory | Alexa K Dowdell et al |
| B.31 | EPI_ISL_456190 | NewZealand/20VR1109/2020 | 2020/3/3 | Institute of Environmental Science and Research (ESR) | Matt Storey et al |
| B.32 | EPI_ISL_425776 | Scotland/CVR47/2020 | 2020/3/13 | COVID-19 Genomics UK (COG-UK) Consortium | Ana da Silva Filipe et al |
| B.33 | EPI_ISL_408010 | USA/CA-CDC-03040142-001/2020 | 2020/1/29 | Pathogen Discovery, Respiratory Viruses Branch, Division of Viral Diseases, Centers for Disease Control and Prevention | Ying Tao et al |
| B.34 | EPI_ISL_425768 | Scotland/CVR40/2020 | 2020/3/13 | COVID-19 Genomics UK (COG-UK) Consortium | Ana da Silva Filipe et al |
| B.35 | EPI_ISL_420104 | Singapore/27/2020 | 2020/3/5 | Programme in Emerging Infectious Diseases, Duke-NUS Medical School | Danielle E Anderson et al |
| B.36 | EPI_ISL_1712699 | Qatar/QA-QU_02-2-44/2020 | 2020/3/13 | Biomedical Research Center (BRC), Qatar University / Qatar Genome Project (QGP) | BRC: Fatiha M. Benslimane et al |
| B.37 | EPI_ISL_631528 | USA/NY-NYCPHL-000032/2020 | 2020/3/12 | New York City Public Health Laboratory | Jade Wang et al |

| | | | | | |
|---|---|---|---|---|---|
| B.38 | EPI_ISL_417008 | Belgium/ULG-3662/2020 | 2020/3/7 | GIGA Medical Genomics | Durkin Keith et al |
| B.39 | EPI_ISL_464192 | England/20098017104/2020 | 2020/2/27 | Respiratory Virus Unit, Microbiology Services Colindale, Public Health England | PHE Covid Sequencing Team et al |
| B.4 | EPI_ISL_412981 | Wuhan/HBCDC-HB-05/2020 | 2020/1/18 | Hubei Provincial Center for Disease Control and Prevention | Bin Fang et al |
| B.4.1 | EPI_ISL_435045 | Kazakhstan/NCB-1/2020 | 2020/3/22 | RSE "National Center for Biotechnology" | Alexandr Shevtsov et al |
| B.4.2 | EPI_ISL_435135 | UnitedArabEmirates/L4682/2020 | 2020/2/25 | Al Jalila Genomics Center | Ahmad Abou Tayoun et al |
| B.4.4 | EPI_ISL_419753 | Australia/VIC34/2020 | 2020/3/10 | Victorian Infectious Diseases Reference Laboratory and Microbiological Diagnostic Unit Public Health Laboratory, Doherty Institute | Caly L. et al |
| B.4.5 | EPI_ISL_419749 | Australia/VIC30/2020 | 2020/3/10 | Victorian Infectious Diseases Reference Laboratory and Microbiological Diagnostic Unit Public Health Laboratory, Doherty Institute | Caly L. et al |
| B.4.6 | EPI_ISL_435121 | UnitedArabEmirates/L0184/2020 | 2020/2/25 | Al Jalila Genomics Center | Ahmad Abou Tayoun et al |
| B.4.7 | EPI_ISL_435120 | UnitedArabEmirates/L068/2020 | 2020/3/14 | Al Jalila Genomics Center | Ahmad Abou Tayoun et al |
| B.40 | EPI_ISL_412116 | England/09c/2020 | 2020/2/9 | Respiratory Virus Unit, Microbiology Services Colindale, Public Health England | Monica Galiano et al |
| B.41 | EPI_ISL_1007665 | SouthKorea/Chosun-CoV01_NP/2020 | 2020/2/6 | Department of Internal Medicine, College of Medicine, Chosun University | Dong-Min Kim et al |
| B.42 | EPI_ISL_497771 | HongKong/HKU-200723-004/2020 | 2020/1/29 | Department of Microbiology, The University of Hong Kong | Kelvin K.W. To et al |
| B.43 | EPI_ISL_417187 | HongKong/HKPU29_0102/2020 | 2020/2/8 | Department of Health Technology and Informatics, Faculty of Health and Social Science, The Hong Kong Polytechnic University | Kenneth Siu-Sing LEUNG et al |
| B.44 | EPI_ISL_418025 | Portugal/PT0040/2020 | 2020/3/17 | Instituto Nacional de Saude (INSA) | Guiomar et al |
| B.45 | EPI_ISL_425762 | Scotland/CVR33/2020 | 2020/3/13 | COVID-19 Genomics UK (COG-UK) Consortium | Ana da Silva Filipe et al |
| B.46 | EPI_ISL_604297 | USA/CA-QDX-1986/2020 | 2020/3/18 | Quest Diagnostics | Rosenthal et al |
| B.47 | EPI_ISL_417296 | England/20109098906/2020 | 2020/3/6 | Respiratory Virus Unit, Microbiology Services Colindale, Public Health England | Monica Galiano et al |

| | | | | | |
|---|---|---|---|---|---|
| B.49 | EPI_ISL_614437 | Denmark/DCGC-978/2020 | 2020/3/9 | Albertsen lab, Department of Chemistry and Bioscience, Aalborg University, Denmark | Danish Covid-19 Genome Consortia et al |
| B.5 | EPI_ISL_2544696 | Japan/KNG0073/2020 | 2020/2/5 | Kanagawa Prefectural Institute of Public Health | Takayuki Hishiki et al |
| B.50 | EPI_ISL_984277 | Indonesia/KI-NIHRDC004160/2020 | 2020/3/14 | National Institute of Health Research and Development | Kindi Adam et al |
| B.51 | EPI_ISL_571259 | USA/WI-QDX-439/2020 | 2020/3/18 | Quest Diagnostics | Rosenthal et al |
| B.52 | EPI_ISL_440349 | England/CAMB-74061/2020 | 2020/3/18 | Wellcome Sanger Institute for the COVID-19 Genomics UK (COG-UK) consortium | Luke W Meredith et al |
| B.53 | EPI_ISL_582609 | UnitedArabEmirates/skmc-0381517/2020 | 2020/2/7 | Molecular/Surveillance lab Sheikh Khalifa Medical City | Amirtharaj Francis et al |
| B.55 | EPI_ISL_451336 | Sichuan/SC-NC-077/2020 | 2020/1/29 | State Key Laboratory of Biotherapy of Sichuan University | Baowen Du et al |
| B.56 | EPI_ISL_435142 | UnitedArabEmirates/L9766/2020 | 2020/3/14 | Al Jalila Genomics Center | Ahmad Abou Tayoun et al |
| B.57 | EPI_ISL_416371 | Shanghai/SH0068/2020 | 2020/2/5 | National Research Center for Translational Medicine (Shanghai), Ruijin Hospital affiliated to Shanghai Jiao Tong University School of Medicine & Shanghai Public Health Clinical Center | Shengyue Wang et al |
| B.58 | EPI_ISL_464404 | England/201061034/2020 | 2020/3/4 | Respiratory Virus Unit, Microbiology Services Colindale, Public Health England | PHE Covid Sequencing Team et al |
| B.6 | EPI_ISL_422407 | Taiwan/NTU04/2020 | 2020/3/4 | Microbial Genomics Core Lab, National Taiwan University Centers of Genomic and Precision Medicine | Shiou-Hwei Yeh et al |
| B.6.1 | EPI_ISL_2001656 | Malaysia/NPHL_19105/2020 | 2020/3/11 | National Public Health Laboratory Malaysia | Noorliza MN et al |
| B.6.2 | EPI_ISL_490090 | Malaysia/IMR_WC80031/2020 | 2020/5/23 | Institute for Medical Research, Infectious Disease Research Centre, National Institutes of Health, Ministry of Health Malaysia | Suppiah J et al |
| B.6.3 | EPI_ISL_429807 | Canada/MB_2/2020 | 2020/3/10 | National Microbiology Laboratory | Anna Majer et al |
| B.6.4 | EPI_ISL_435689 | Singapore/84/2020 | 2020/4/18 | National Public Health Laboratory, National Centre for Infectious Diseases | Mak Tze Minn et al |

| | | | | | |
|---|---|---|---|---|---|
| B.6.5 | EPI_ISL_500639 | Australia/NSW2168/2020 | 2020/3/19 | Area of Virology, Serology and Virology Division (SAViD), New South Wales Health Pathology Randwick | Rawlinson et al |
| B.6.6 | EPI_ISL_5051470 | Philippines/PH-RITM-0108/2020 | 2020/3/10 | Research Institute for Tropical Medicine | Inez Andrea Medado et al |
| B.6.8 | EPI_ISL_6463883 | PapuaNewGuinea/PNG3146/2020 | 2020/7/13 | Microbiological Diagnostic Unit - Public Health Laboratory (MDU-PHL) | Kumbu et al |
| B.60 | EPI_ISL_418677 | England/20122087702/2020 | 2020/3/15 | Respiratory Virus Unit, Microbiology Services Colindale, Public Health England | Monica Galiano et al |
| B.61 | EPI_ISL_464172 | England/20079000404/2020 | 2020/2/14 | Respiratory Virus Unit, Microbiology Services Colindale, Public Health England | PHE Covid Sequencing Team et al |
| C.1 | EPI_ISL_3537060 | USA/CA-CDPH1181/2020 | 2020/1/3 | California Department of Public Health | CDPH IDLB COVIDNet et al |
| C.1.1 | EPI_ISL_887438 | Mozambique/INS-K007883/2020 | 2020/11/25 | KRISP, KZN Research Innovation and Sequencing Platform | Nalia Ismael et al |
| C.1.2 | EPI_ISL_2770450 | SouthAfrica/KRISP-K018739/2021 | 2021/6/5 | KRISP, KZN Research Innovation and Sequencing Platform | Florette Treurnicht et al |
| C.10 | EPI_ISL_451646 | Poland/Pom5/2020 | 2020/4/22 | Laboratory of Recombinant Vaccines | Lukasz Rabalski et al |
| C.11 | EPI_ISL_445350 | Chile/RM-ISPCH-54/2020 | 2020/4/1 | Instituto de Salud Publica de Chile | Andrés E Castillo et al |
| C.12 | EPI_ISL_547977 | NewZealand/20CV0068/2020 | 2020/8/12 | Institute of Environmental Science and Research (ESR) | Xiaoyun Ren et al |
| C.13 | EPI_ISL_1111163 | Peru/ICA-INS-703/2020 | 2020/3/29 | Laboratorio de Referencia Nacional de Enteropatógenos. Instituto Nacional de Salud del Perú | Ronnie Gavilan Chavez et al |
| C.14 | EPI_ISL_1111162 | Peru/ICA-INS-702/2020 | 2020/3/20 | Laboratorio de Referencia Nacional de Enteropatógenos. Instituto Nacional de Salud del Perú | Ronnie Gavilan Chavez et al |
| C.17 | EPI_ISL_479723 | Egypt/CUNCI-HGC6I033/2020 | 2020/6/2 | Egyptian National Cancer Institute (ENCI) | Zekri et al |

| | | | | | |
|---|---|---|---|---|---|
| C.18 | EPI_ISL_1692799 | Italy/CAM-TIGEM-IZSM-COLLI-9470/2020 | 2020/7/1 | Telethon Institute of Genetics and Medicine (TIGEM) | Antonio Grimaldi Patrizia Annunziata Francesco Panariello Teresa Giuliano Michele Cennamo Valentina Bouche Chiara Colantuono Lucio Di Filippo Mariano Fiorenza Anna Manfredi Marcello Salvi Giuseppe Portella Andrea Ballabio Davide Cacchiarelli et al |
| C.19 | EPI_ISL_747927 | Denmark/DCGC-20010/2020 | 2020/12/7 | Albertsen Lab, Department of Chemistry and Bioscience, Aalborg University, Denmark | Danish Covid-19 Genome Consortium et al |
| C.2 | EPI_ISL_490282 | SouthAfrica/R10517/2020 | 2020/6/9 | National Institute for Communicable Diseases of the National Health Laboratory Service | Allam M et al |
| C.2.1 | EPI_ISL_1014550 | Curacao/CW-RIVM-21733/2020 | 2020/12/18 | National Institute for Public Health and the Environment (RIVM) | Adam Meijer et al |
| C.20 | EPI_ISL_617252 | Denmark/DCGC-8213/2020 | 2020/10/26 | Albertsen lab, Department of Chemistry and Bioscience, Aalborg University, Denmark | Danish Covid-19 Genome Consortia et al |
| C.21 | EPI_ISL_804964 | Spain/ML-ISCIII-202059850/2020 | 2020/9/15 | Instituto de Salud Carlos III | Iglesias-Caballero et al |
| C.22 | EPI_ISL_430808 | Argentina/PAIS-A0017/2020 | 2020/4/3 | Área de Secuenciación del Laboratorio de Virología del Hospital de Niños Dr. Ricardo Gutierrez on behalf of 'Proyecto Argentino Interinstitucional de genomica de SARS-CoV-2' (PAIS Consortium) | Nabaes Jodar et al |
| C.23 | EPI_ISL_548944 | Pakistan/UN-UVAS-Lahore-III/2020 | 2020/5/11 | Institute of Microbiology, University of Veterinary and Animal sciences | Yaqub et al |
| C.25 | EPI_ISL_529068 | Peru/LIM-UPCH-0011/2020 | 2020/3/20 | Laboratorio de Genómica Microbiana, Universidad Peruana Cayetano Heredia | Pablo Tsukayama et al |
| C.26 | EPI_ISL_1167833 | Chile/RM-44567/2020 | 2020/4/28 | Instituto de Salud Publica de Chile | Javier Tognarelli et al |
| C.27 | EPI_ISL_444426 | England/CAMB-7FE7B/2020 | 2020/4/19 | COVID-19 Genomics UK (COG-UK) Consortium | Luke W Meredith et al |
| C.28 | EPI_ISL_537505 | USA/CA-UCLA-17/2020 | 2020/4/30 | Kruglyak Lab | Guo et al. |
| C.29 | EPI_ISL_746564 | Chile/RM-110525/2020 | 2020/6/25 | Instituto de Salud Publica de Chile | Javier Tognarelli et al |

| | | | | | |
|---|---|---|---|---|---|
| C.3 | EPI_ISL_559590 | England/ALDP-49E20D/2020 | 2020/5/20 | Wellcome Sanger Institute for the COVID-19 Genomics UK (COG-UK) consortium | The Lighthouse Lab in Alderley Park et al |
| C.30 | EPI_ISL_532000 | England/QEUH-960946/2020 | 2020/8/4 | Wellcome Sanger Institute for the COVID-19 Genomics UK (COG-UK) consortium | Harper VanSteenhouse et al |
| C.30.1 | EPI_ISL_619440 | Denmark/DCGC-6540/2020 | 2020/10/5 | Albertsen lab, Department of Chemistry and Bioscience, Aalborg University, Denmark | Danish Covid-19 Genome Consortia et al |
| C.31 | EPI_ISL_1494152 | USA/IL-IDPH-COO-S-0001755/2020 | 2020/8/11 | Gagnon Lab, Southern Illinois University | Keith Gagnon et al |
| C.32 | EPI_ISL_1111276 | Peru/LIM-INS-378/2020 | 2020/5/7 | Laboratorio de Referencia Nacional de Enteropatógenos. Instituto Nacional de Salud del Perú | Ronnie Gavilan Chavez et al |
| C.33 | EPI_ISL_1532156 | Peru/ICA-INS-757/2020 | 2020/5/18 | Laboratorio de Referencia Nacional de Biotecnología y Biología Molecular. Instituto Nacional de Salud Perú | Carlos Padilla Rojas et al |
| C.34 | EPI_ISL_968510 | Canada/BC-BCCDC-6020/2020 | 2020/8/6 | BCCDC Public Health Laboratory | Prystajecky Natalie et al |
| C.35 | EPI_ISL_1111080 | Peru/ANC-INS-620/2020 | 2020/3/31 | Laboratorio de Referencia Nacional de Enteropatógenos. Instituto Nacional de Salud del Perú | Ronnie Gavilan Chavez et al |
| C.36 | EPI_ISL_509726 | USA/FL-BPHL-0668/2020 | 2020/3/13 | Florida Bureau of Public Health Laboratories | Sarah Schmedes et al |
| C.36.1 | EPI_ISL_1366018 | Canada/Qc-L00292853/2020 | 2020/9/24 | Laboratoire de santé publique du Québec | Sandrine Moreira et al |
| C.36.2 | EPI_ISL_930860 | Switzerland/SO-UHB-42489126/2020 | 2020/10/16 | Clinical Bacteriology | Tim Roloff et al |
| C.36.3 | EPI_ISL_3274160 | Egypt/CPHL-A4/2021 | 2021/1/6 | CPHL/MOH/EGYPT | Wael H. Roshdy/ Mohamed Kamal / Shymaa s. Ahmed/ Ramy Galal/Nancy el guindy/ Amel nagiub/yasser el hady/salma sayed/ Abd Monaem Adel/Galal Mahmoud/Dalia Ramadan/Rabeh ..R. El/Shesheny/ Mohamed A Ali/Mohamed Hassany et al |
| C.36.3.1 | EPI_ISL_1464522 | Germany/HH-HPI-p4792/2021 | 2021/3/4 | Heinrich Pette Institute, Leibniz Institute for Experimental Virology | Alexis Robitaille et al |

| | | | | | |
|---|---|---|---|---|---|
| C.37.1 | EPI_ISL_3023616 | Peru/MDD-INS-1923/2021 | 2021/3/15 | Laboratorio de Referencial Nacional de Virus Respiratorios | Carlos Padilla Rojas et al |
| C.38 | EPI_ISL_1357071 | Germany/BY-RKI-I-049645/2021 | 2021/2/16 | Robert Koch Institute | ? |
| C.39 | EPI_ISL_2245832 | Guinea/Conakry_IPG12108/2020 | 2020/9/23 | Institut Pateur de Dakar | Grayo Solene et al |
| C.39 | EPI_ISL_3401511 | Peru/LAL-INS-2325/2021 | 2021/2/10 | Laboratorio de Referencia Nacional de Virus Respiratorios. Centro Nacional de Salud Publica. Instituto Nacional de Salud Peru. | Carlos Padilla Rojas et al |
| C.4 | EPI_ISL_1111257 | Peru/LIM-INS-359/2020 | 2020/5/19 | Laboratorio de Referencia Nacional de Enteropatógenos. Instituto Nacional de Salud del Perú | Ronnie Gavilan Chavez et al |
| C.5 | EPI_ISL_541472 | Switzerland/VD-ETHZ-270056/2020 | 2020/9/2 | Department of Biosystems Science and Engineering, ETH Zürich | Christian Beisel et al |
| C.6 | EPI_ISL_3068080 | SouthAfrica/Tygerberg_79/2020 | 2020/4/18 | Division of Medical Virology, Stellenbosch University and NHLS Tygerberg Hospital | Susan Engelbrecht et al |
| C.7 | EPI_ISL_614513 | Denmark/DCGC-1088/2020 | 2020/5/11 | Albertsen lab, Department of Chemistry and Bioscience, Aalborg University, Denmark | Danish Covid-19 Genome Consortia et al |
| C.8 | EPI_ISL_961669 | Canada/NB-NML-240/2020 | 2020/3/31 | National Microbiology Laboratory (NML) | Anna Majer et al |
| C.9 | EPI_ISL_2444017 | SouthAfrica/NICD-N00184/2020 | 2020/5/20 | National Institute for Communicable Diseases of the National Health Laboratory Service | Amoako DG et al |
| D.2 | EPI_ISL_480772 | Australia/VIC2165/2020 | 2020/3/19 | MDU-PHL | Seemann T. et al |
| D.3 | EPI_ISL_521953 | Australia/VIC2213/2020 | 2020/6/14 | VIDRL and MDU-PHL | Caly L. et al |
| D.4 | EPI_ISL_578302 | Ireland/KK-NVRL-72IRL24802/2020 | 2020/8/13 | National Virus Reference Laboratory | Michael Carr et al |
| D.5 | EPI_ISL_1008424 | Sweden/21-20401/2020 | 2020/10/12 | The Public Health Agency of Sweden | Anna-Malin Linde et al |
| G.1 | EPI_ISL_566478 | England/MILK-9C2766/2020 | 2020/9/10 | Wellcome Sanger Institute for the COVID-19 Genomics UK (COG-UK) consortium | The Lighthouse Lab in Milton Keynes et al |
| K.1 | EPI_ISL_526708 | SouthKorea/KCDC2774/2020 | 2020/7/23 | Division of Viral Diseases, Center for Laboratory Control of Infectious Diseases, Korea Centers for Diseases Control and Prevention | Jeong-Min Kim et al |
| K.2 | EPI_ISL_596354 | Russia/PRI-RII-MH744S/2020 | 2020/8/7 | WHO National Influenza Centre Russian Federation | Andrey Komissarov et al |

| | | | | | |
|---|---|---|---|---|---|
| K.3 | EPI_ISL_590909 | Norway/3271/2020 | 2020/8/29 | Norwegian Institute of Public Health, Department of Virology | Kathrine Stene-Johansen et al |
| L.1 | EPI_ISL_579578 | Canada/NS-NML-948/2020 | 2020/3/26 | National Microbiology Laboratory (NML) | Anna Majer et al |
| L.2 | EPI_ISL_454766 | Netherlands/Utrecht_10017/2020 | 2020/3/23 | National Institute for Public Health and the Environment (RIVM) | Adam Meijer et al |
| L.3 | EPI_ISL_729932 | Nigeria/FC-CV247/2020 | 2020/7/8 | African Centre of Excellence for Genomics of Infectious Diseases (ACEGID), Redeemer's University, Ede, Osun State, Nigeria | Oluniyi P.E. et al |
| L.4 | EPI_ISL_1017303 | USA/OR-PROV-301/2020 | 2020/6/29 | Providence St. Joseph Health Molecular Genomics Laboratory | Alexa K Dowdell et al |
| M.1 | EPI_ISL_516935 | Moldova/ICGEB_MD4/2020 | 2020/6/18 | International Centre for Genetic Engineering and Biotechnology (ICGEB) and ARGO Open Lab Platform for Genome Sequencing | Ulinici M et al |
| M.2 | EPI_ISL_829812 | Iceland/6134/2020 | 2020/10/26 | deCODE genetics | Daniel F Gudbjartsson et al |
| M.3 | EPI_ISL_551169 | England/MILK-9A9C2A/2020 | 2020/9/1 | Wellcome Sanger Institute for the COVID-19 Genomics UK (COG-UK) consortium | The Lighthouse Lab in Milton Keynes et al |
| N.1 | EPI_ISL_460202 | USA/MA-MGH-00076/2020 | 2020/3/25 | Infectious Disease Program, Broad Institute of Harvard and MIT | Lemieux et al |
| N.2 | EPI_ISL_6507361 | FrenchGuiana/IPP8886/2020 | 2020/6/23 | National Reference Center for Viruses of Respiratory Infections, Institut Pasteur, Paris | Mélanie Albert et al |
| N.3 | EPI_ISL_430814 | Argentina/PAIS-A0023/2020 | 2020/4/17 | Área de Secuenciación del Laboratorio de Virología del Hospital de Niños Dr. Ricardo Gutierrez on behalf of 'Proyecto Argentino Interinstitucional de genomica de SARS-CoV-2' (PAIS Consortium) | Nabaes Jodar et al |
| N.4 | EPI_ISL_586379 | Canada/ON-S201/2020 | 2020/3/25 | McMaster University | Allison McGeer et al |
| N.5 | EPI_ISL_792335 | Argentina/PAIS-A0255/2020 | 2020/4/25 | Área de Secuenciación del Laboratorio de Virología del Hospital de Niños Dr. Ricardo Gutierrez on behalf of 'Proyecto Argentino Interinstitucional de genomica de SARS-CoV-2' (PAIS Consortium) | Nabaes Jodar et al |
| N.6 | EPI_ISL_1167830 | Chile/AR-265171/2020 | 2020/2/16 | Instituto de Salud Publica de Chile | Javier Tognarelli et al |
| N.7 | EPI_ISL_749151 | Uruguay/TYT-M57/2020 | 2020/6/18 | Institut Pasteur de Montevideo | Daiana Mir et al |

| | | | | | |
|---|---|---|---|---|---|
| N.8 | EPI_ISL_806623 | Kenya/C24022/2020 | 2020/6/23 | KEMRI-Wellcome Trust Research Programme/KEMRI-CGMR-C Kilifi | Githinji et al |
| N.9 | EPI_ISL_2241601 | Brazil/PI-BA1200-584437/2020 | 2020/10/12 | Coordenação Geral de Laboratórios de Saúde Pública (CGLAB/DAEVS/SVS/MS) | Vagner Fonseca et al |
| None | EPI_ISL_1423385 | USA/NY-NYULH1008/2021 | 2021/2/23 | Departments of Pathology and Medicine, New York University School of Medicine | Adriana Heguy et al |
| P.1 | EPI_ISL_2249957 | USA/IL-RED-FUL-39828238/2020 | 2020/4/7 | Reditus Laboratories | Joshua J. Geltz et al |
| P.1 | EPI_ISL_1404618 | USA/MA-MASPHL-02518/2021 | 2021/3/18 | Massachusetts State Public Health Laboratory | Andrew Lang et al |
| P.1.1 | EPI_ISL_2775487 | Brazil/PR-FIOCRUZ-33170-TG-165/2020 | 2020/12/30 | Instituto Carlos Chagas - Fiocruz | Mauro de Medeiros Oliveira et al |
| P.1.1 | EPI_ISL_5155963 | USA/NY-P16G7/2021 | 2021/4/23 | 505 Irving Avenue, Syracuse, NY 13210 - Frank Middleton Lab | Frank Middleton et al |
| P.1.10 | EPI_ISL_3912382 | Brazil/CE-FIOCRUZ-00730/2021 | 2021/2/13 | Analytical Competence Molecular Epidemiology Lab/ACME, Oswaldo Cruz Foundation, Ceara (FIOCRUZ CE) | Fabio Miyajima et al |
| P.1.10.2 | EPI_ISL_2674658 | Mexico/HID-InDRE_FB17208_S3171/2021 | 2021/5/23 | Instituto de Diagnostico y Referencia Epidemiologicos (INDRE) | Claudia Wong-Arambula et al |
| P.1.11 | EPI_ISL_2445049 | Brazil/SP-IB_103317/2021 | 2021/5/10 | Instituto Butantan | Dimas Tadeu Covas et al |
| P.1.12 | EPI_ISL_3376385 | Peru/LOR-INS-2202/2021 | 2021/1/2 | Laboratorio de Referencial Nacional de Virus Respiratorios | Carlos Padilla Rojas et al |
| P.1.13 | EPI_ISL_1420865 | USA/FL-CDC-STM-000023348/2021 | 2021/2/24 | Centers for Disease Control and Prevention Division of Viral Diseases, Pathogen Discovery | Peter W. Cook et al |
| P.1.14 | EPI_ISL_1821208 | Brazil/SP-2389/2020 | 2020/11/3 | Instituto Adolfo Lutz, Interdiciplinary Procedures Center, Strategic Laboratory | Claudio Tavares Sacchi et al |
| P.1.15 | EPI_ISL_2233906 | Brazil/AM-CD1739/2020 | 2020/12/21 | Scientific Platform Pasteur USP, University of Sao Paulo | Cunha et al |
| P.1.16 | EPI_ISL_984622 | Belgium/ULG-11925/2021 | 2021/1/31 | GIGA Medical Genomics | Keith Durkin et al |
| P.1.17.1 | EPI_ISL_3334239 | Luxembourg/LNS8192483/2021 | 2021/4/2 | Laboratoire national de sante, Microbiology, Microbial Genomics Platform | Anke Wienecke-Baldacchino et al |
| P.1.2 | EPI_ISL_3664121 | Brazil/PE-NXBS509GENOV827539448543/2021 | 2021/1/3 | DASA | Rodrigo Guarischi et al |

| | | | | | |
|---|---|---|---|---|---|
| P.1.3 | EPI_ISL_2777689 | Brazil/AM-FIOCRUZ-21891957JM/2021 | 2021/3/12 | Laboratorio de Ecologia de Doencas Transmissiveis na Amazonia, Instituto Leonidas e Maria Deane - Fiocruz Amazonia | Valdinete Nascimento et al |
| P.1.4 | EPI_ISL_2777824 | Brazil/AM-FIOCRUZ-21896338DCF/2021 | 2021/3/19 | Laboratorio de Ecologia de Doencas Transmissiveis na Amazonia, Instituto Leonidas e Maria Deane - Fiocruz Amazonia | Valdinete Nascimento et al |
| P.1.5 | EPI_ISL_2777660 | Brazil/AM-FIOCRUZ-21891575VSC/2021 | 2021/4/3 | Laboratorio de Ecologia de Doencas Transmissiveis na Amazonia, Instituto Leonidas e Maria Deane - Fiocruz Amazonia | Valdinete Nascimento et al |
| P.1.6 | EPI_ISL_2777596 | Brazil/AM-FIOCRUZ-21890984TSM/2021 | 2021/3/26 | Laboratorio de Ecologia de Doencas Transmissiveis na Amazonia, Instituto Leonidas e Maria Deane - Fiocruz Amazonia | Valdinete Nascimento et al |
| P.1.8 | EPI_ISL_3664270 | Brazil/RJ-NXBS768GENOV827517602806/2021 | 2021/5/3 | DASA | Rodrigo Guarischi et al |
| P.2 | EPI_ISL_2157561 | Brazil/RJ-FIOCRUZ-2188-1P/2020 | 2020/4/13 | Laboratory of Respiratory Viruses and Measles, Oswaldo Cruz Institute, FIOCRUZ | Paola Resende et al |
| P.3 | EPI_ISL_1597203 | Netherlands/FL-RIVM-24324/2021 | 2021/2/27 | National Institute for Public Health and the Environment (RIVM) | Adam Meijer et al |
| P.4 | EPI_ISL_1625982 | Brazil/SP-2281/2021 | 2021/2/17 | Instituto Adolfo Lutz, Interdiciplinary Procedures Center, Strategic Laboratory | Claudio Tavares Sacchi et al |
| P.5 | EPI_ISL_1445155 | Brazil/SP-765479-NBD14/2021 | 2021/3/9 | Instituto Butantan / Mendelics | Dimas Tadeu Covas et al |
| P.6 | EPI_ISL_2965563 | Uruguay/RT16-UYMO/2020 | 2020/12/2 | Centro de Innovación en Vigilancia Epidemiológica (CiVE), Institut Pasteur Montevideo, Uruguay | Natalia Rego et al |
| P.7 | EPI_ISL_5416087 | Brazil/MG-LBIb1128/2020 | 2020/6/24 | Laboratório de Biologia Integrativa / UFMG | Ana Valesca Fernandes Gilson Silva et al |
| Q.1 | EPI_ISL_2903380 | England/OXON-F8A254/2020 | 2020/12/19 | COVID-19 Genomics UK (COG-UK) Consortium | Tanya Golubchik et al |

| | | | | | |
|---|---|---|---|---|---|
| Q.2 | EPI_ISL_2020611 | Italy/CAM-TIGEM-11334/2020 | 2020/12/15 | TIGEM | Antonio Grimaldi Patrizia Annunziata Francesco Panariello Teresa Giuliano Michele Cennamo Valentina Bouche Chiara Colantuono Lucio Di Filippo Mariano Fiorenza Anna Manfredi Marcello Salvi Giuseppe Portella Andrea Ballabio Davide Cacchiarelli et al |
| Q.3 | EPI_ISL_2628482 | USA/CA-ALSR-101991/2020 | 2020/7/8 | Andersen lab at Scripps Research | SEARCH Alliance San Diego with Ashleigh Murphy et al |
| Q.4 | EPI_ISL_1730391 | England/LOND-E652F/2020 | 2020/12/13 | COVID-19 Genomics UK (COG-UK) Consortium | Sergi Castellano et al |
| Q.6 | EPI_ISL_1262942 | France/PDL-IPP04848/2021 | 2021/3/2 | National Reference Center for Viruses of Respiratory Infections, Institut Pasteur, Paris | Marion Barbet et al |
| Q.7 | EPI_ISL_1135025 | France/ARA-HCL021022962101/2021 | 2021/1/29 | CNR Virus des Infections Respiratoires - France SUD | Antonin Bal et al |
| Q.8 | EPI_ISL_917989 | Scotland/QEUH-10E71AF/2021 | 2021/1/21 | Wellcome Sanger Institute for the COVID-19 Genomics UK (COG-UK) Consortium | Harper VanSteenhouse et al |
| R.1 | EPI_ISL_2716627 | SierraLeone/SL10/2020 | 2020/1/14 | State Key Laboratory of Pathogen and Biosecurity, Beijing Institute of Microbiology and Epidemiology | Wurie et al |
| R.2 | EPI_ISL_2284073 | USA/MA-Yale-5158/2020 | 2020/12/22 | Yale Center for Genomic Analysis | Shrikant Mane et al |
| S.1 | EPI_ISL_552514 | England/MILK-975243/2020 | 2020/8/21 | Wellcome Sanger Institute for the COVID-19 Genomics UK (COG-UK) consortium | The Lighthouse Lab in Milton Keynes et al |
| U.1 | EPI_ISL_838504 | England/LIVE-DCEE25/2020 | 2020/12/15 | COVID-19 Genomics UK (COG-UK) Consortium | Sam Haldenby et al |
| U.2 | EPI_ISL_1469208 | Sweden/RVFOI21-00682/2020 | 2020/11/11 | CBRN Defence and Security, Swedish Defence Research Agency | Andreas Sjödin et al |
| U.3 | EPI_ISL_1317736 | Norway/2001/2021 | 2021/1/11 | Norwegian Institute of Public Health, Department of Virology | Kathrine Stene-Johansen et al |
| V.1 | EPI_ISL_822311 | NorthernIreland/CAMC-A41A92/2020 | 2020/10/5 | Wellcome Sanger Institute for the COVID-19 Genomics UK (COG-UK) Consortium | Rob Howes et al |

| | | | | | |
|---|---|---|---|---|---|
| V.2 | EPI_ISL_650923 | NorthernIreland/NIRE-10926A/2020 | 2020/10/20 | COVID-19 Genomics UK (COG-UK) Consortium | Conall McCaughey et al |
| W.1 | EPI_ISL_678350 | Australia/NSW2802/2020 | 2020/10/2 | Virology Research Laboratory; Area of Virology, Serology and Virology Division (SAViD), New South Wales Health Pathology Randwick | Foster et al |
| W.2 | EPI_ISL_3426611 | Netherlands/NB-MVD-CWGS2101771/2020 | 2020/9/23 | Microvida | Suzan D. Pas et al |
| W.3 | EPI_ISL_530531 | England/QEUH-96DE86/2020 | 2020/8/18 | Wellcome Sanger Institute for the COVID-19 Genomics UK (COG-UK) consortium | Harper VanSteenhouse et al |
| W.4 | EPI_ISL_551840 | England/CAMC-990C35/2020 | 2020/8/27 | Wellcome Sanger Institute for the COVID-19 Genomics UK (COG-UK) consortium | Rob Howes et al |
| XA | EPI_ISL_989697 | Wales/ALDP-11CF93B/2021 | 2021/1/30 | Wellcome Sanger Institute for the COVID-19 Genomics UK (COG-UK) Consortium | Jacquelyn Wynn et al |
| XB | EPI_ISL_2628473 | USA/SEARCH-101966/2020 | 2020/7/8 | Andersen lab at Scripps Research | SEARCH Alliance San Diego with Ashleigh Murphy et al |
| Y.1 | EPI_ISL_651724 | England/CAMC-B23224/2020 | 2020/11/3 | COVID-19 Genomics UK (COG-UK) Consortium | Thushan de Silva et al |
| Z.1 | EPI_ISL_950606 | England/NOTT-246E3F/2020 | 2020/6/2 | COVID-19 Genomics UK (COG-UK) Consortium | Gemma Clark et al |
