## Supplementary material for "Computation of Antigenicity Predicts SARS-CoV-2 Vaccine Breakthrough Variants": Table S5

**Table S5 Data sources for Efficacy Data**

| Manufacturer | Vaccine | Test ID | Study Type | VE | Ref(DOI/Link) |
|---|---|---|---|---|---|
| AstraZeneca | ChAdOx1 nCoV-19 | AZ | Phase3 | 0.81 | 10.1016/S0140-6736(21)00432-3 |
| | | US-AZ | Phase3 | 0.74 | 10.1056/NEJMoa2105290 |
| | | CL-AZ | RealWorld | 0.71 | https://cdn.who.int/media/docs/default-source/blue-print/chile_rafael-araos_who-vr-call_25oct2021.pdf?sfvrsn=7a7ca72a_7 |
| | | Scotland-Lancet-AZ | RealWorld | 0.59 | 10.1016/S0140-6736(21)00677-2 |
| | | England-AZ | RealWorld | 0.89 | https://assets.publishing.service.gov.uk/government/uploads/system/uploads/attachment_data/file/990089/Vaccine_surveillance_report_-_week_20.pdf |
| BioNTech | BNT162b2 | BNT | Phase3 | 0.95 | 10.1056/NEJMoa2034577 |
| | | CL-BNT | RealWorld | 0.84 | https://cdn.who.int/media/docs/default-source/blue-print/chile_rafael-araos_who-vr-call_25oct2021.pdf?sfvrsn=7a7ca72a_7 |
| | | Scotland-Lancet-BNT | RealWorld | 0.77 | 10.1016/S0140-6736(21)00677-2 |
| | | IL-NEJM-BNT | RealWorld | 0.94 | 10.1056/NEJMoa2101765 |
| | | IL-Lancet-BNT | RealWorld | 0.97 | 10.1016/S0140-6736(21)00947-8 |
| | | URY-BNT | RealWorld | 0.78 | https://www.gub.uy/ministerio-salud-publica/comunicacion/noticias/segundo-estudio-efectividad-vacunacion-anti-sars-cov-2-uruguay-8-junio-2021 |
| | | England-BNT | RealWorld | 0.85 | https://assets.publishing.service.gov.uk/government/uploads/system/uploads/attachment_data/file/990089/Vaccine_surveillance_report_-_week_20.pdf |
| | | US-BNT | RealWorld | 0.90 | 10.15585/mmwr.mm7013e3 |
| Cadila | ZyCoV-D | Cadila | Phase3 | 0.67 | https://zyduscadila.com/public/pdf/pressrelease/ZyCoV_D_Press_Release_1_7_2021.pdf |
| Covaxin | BBV152 | Covaxin | Phase3 | 0.78 | 10.1016/S0140-6736(21)02000-6 |
| Johnson | Ad26.COV2.S | Johnson | Phase3 | 0.66 | 10.1056/NEJMoa2101544 |
| | | US-Johnson | Phase3 | 0.72 | 10.1056/NEJMoa2101544 |
| | | BR-Johnson | Phase3 | 0.68 | 10.1056/NEJMoa2101544 |
| | | SA-Johnson | Phase3 | 0.64 | 10.1056/NEJMoa2101544 |

| | | | | | |
|---|---|---|---|---|---|
| Longcom | ZF2001 | Longcom | Phase3 | 0.82 | https://www.scmp.com/coronavirus/greater-china/article/3146729/chinese-3-shot-covid-19-vaccine-maker-says-trials-show-it |
| Moderna | mRNA-1273 | Moderna | Phase3 | 0.94 | 10.1056/NEJMoa2035389 |
| | | US-Moderna | RealWorld | 0.90 | 10.15585/mmwr.mm7013e3 |
| Novavax | NVX-CoV2373 | Novavax | Phase3 | 0.89 | 10.1126/science.abg8101 |
| Sinopharm | BBIBP-CorV | Sinopharm | Phase3 | 0.78 | 10.1001/jama.2021.8565 |
| SinoVac | CoronaVac | BR-SinoVac | Phase3 | 0.51 | 10.2139/ssrn.3822780 |
| | | TR-SinoVac | Phase3 | 0.84 | 10.1016/S0140-6736(21)01429-X |
| | | BR2-SinoVac | RealWorld | 0.80 | https://apnews.com/article/caribbean-brazil-coronavirus-pandemic-business-health-20bd94d28ac7b373d7a8f3f9c557e5b6 |
| | | CL-SinoVac | RealWorld | 0.54 | https://cdn.who.int/media/docs/default-source/blue-print/chile_rafael-araos_who-vr-call_25oct2021.pdf?sfvrsn=7a7ca72a_7 |
| | | CL-NEJM-SinoVac | RealWorld | 0.66 | 10.1056/NEJMoa2107715 |
| | | URY-SinoVac | RealWorld | 0.60 | https://www.gub.uy/ministerio-salud-publica/comunicacion/noticias/segundo-estudio-efectividad-vacunacion-anti-sars-cov-2-uruguay-8-junio-2021 |
| | | ID-SinoVac | RealWorld | 0.94 | https://sehatnegeriku.kemkes.go.id/baca/berita-utama/20210512/1937767/kajian-cepat-kemenkes-vaksin-sinovac-efektif-cegah-kematian/ |
| Sputnik | rAd26-S+rAd5-S | Sputnik | Phase3 | 0.92 | 10.1016/S0140-6736(21)00234-8 |
| | | RU-Sputnik | RealWorld | 0.98 | https://rdif.ru/Eng_fullNews/6722/ |
| | | UAE-Sputnik | RealWorld | 0.98 | https://rdif.ru/Eng_fullNews/6919/ |
